# Tertiary structure of single-instant RNA molecule reveals folding landscape

**DOI:** 10.1101/2023.05.19.541511

**Authors:** Jianfang Liu, Ewan K.S. McRae, Meng Zhang, Cody Geary, Ebbe Sloth Andersen, Gang Ren

## Abstract

The folding of RNA and protein molecules during their synthesis is a crucial self-assembly process that nature employs to convert genetic information into the complex molecular machinery that supports life. Misfolding events are the cause of several diseases, and the folding pathway of central biomolecules, such as the ribosome, is strictly regulated by programmed maturation processes and folding chaperones. However, the dynamic folding processes are challenging to study because current structure determination methods heavily rely on averaging, and existing computational methods do not efficiently simulate non-equilibrium dynamics. Here we utilize individual-particle cryo-electron tomography (IPET) to investigate the folding landscape of a rationally designed RNA origami 6-helix bundle that undergoes slow maturation from a “young” to “mature” conformation. By optimizing the IPET imaging and electron dose conditions, we obtain 3D reconstructions of 120 individual particles at resolutions ranging from 23-35 Å, enabling us first-time to observe individual RNA helices and tertiary structures without averaging. Statistical analysis of 120 tertiary structures confirms the two main conformations and suggests a possible folding pathway driven by helix-helix compaction. Studies of the full conformational landscape reveal both trapped states, misfolded states, intermediate states, and fully compacted states. The study provides novel insight into RNA folding pathways and paves the way for future studies of the energy landscape of molecular machines and self-assembly processes.

## Introduction

“*No man ever steps in the same river twice, for it’s not the same river and he’s not the same man*,” stated the ancient Greek philosopher Heraclitus. Similarly, macromolecules in aqueous environments exhibit dynamic behavior characterized by continuous conformational changes due to thermal vibrations, meaning that they are never exactly in the same state at a given timepoint. Cryogenic electron microscopy (cryo-EM) is a technique that enables the examination of the composition and kinetic behavior of large molecules in solution by rapidly freezing the aqueous sample to capture a momentary image of its conformation at a specific temperature and time. However, due to the acquisition of a single projection of each molecule, the cryo-EM single-particle analysis (SPA) method^1^ must select similar conformations from a large population of heterogeneous molecules and average them to obtain a 3D structure estimate. Although SPA can generate high-resolution structures, the determined structures are structural averages, resulting in an anisotropic distribution of resolution^2^ and absence of flexible domains^3^. While methods for analyzing conformationally heterogeneous populations of biomolecules in cryo-EM data are being developed to elucidate dynamical features and the structure of flexible domains^4–6^, determining the structure with small populations or rare high-energy states is difficult using SPA. This is particularly challenging for molecules with “floppy” and “disordered” structures that undergo continuous conformational changes, such as DNA, RNA, antibodies, lipoproteins, intrinsically disordered proteins (IDPs)^7^, and intermediates of chemical reactions or folding.

Natural RNAs, such as ribosomal RNA, are difficult to be studied due to their highly complex folding paths, with a significant contribution from noncanonical base-pair interactions and folding chaperones ^8^, the rationally designed RNA nanostructures, such as cotranscriptional RNA ^9^, represent simpler systems that capture non-equilibrium elements, including folding traps and maturation processes ^10, 11^. Unlike to DNA origami nanostructure ^12^ that folds by heat annealing to thermodynamic equilibrium, RNA origami was developed as a single-stranded architecture to enable isothermal folding during transcription by RNA polymerase ^9^. Cotranscriptional folding is a non-equilibrium self-assembly process that leads to the formation of structures that are often far from the global minimum free energy state ^13^.

RNA origami can be computationally designed ^10^ with specific base pairings to gradually fold into secondary structures, such as stems, loops, and junctions. Subsequently, the structure further condenses into tertiary structures through the formation of pseudoknots, which involve kissing-loop (KL) interactions (**Fig. 1a**). Using a combination of cryo-EM SPA and small-angle X-ray scattering (SAXS) ^11^, an RNA origami 6-helix bundle with a clasp helix (6HBC) (**Extended Data Fig. 1**) was found undergoing a slow conformational maturation process, where a “young” open conformation changes into a “mature” closed conformation ∼10 hours after transcription initiation (**Fig. 1a**). Cryo-EM SPA of a 6HBC sample at this transition point allowed sorting and reconstruction of the two major conformations to sub-nanometer resolution, which together with the observation of a structural transition by SAXS, led to the hypothesis of a transition state where a central KL is broken and reformed (**Fig. 1b**). Further attempts to identify and reconstruct intermediate conformations using conventional cryo-EM SPA methods were not readily achievable (**Supplementary Fig. 1**).

**Fig. 1:**
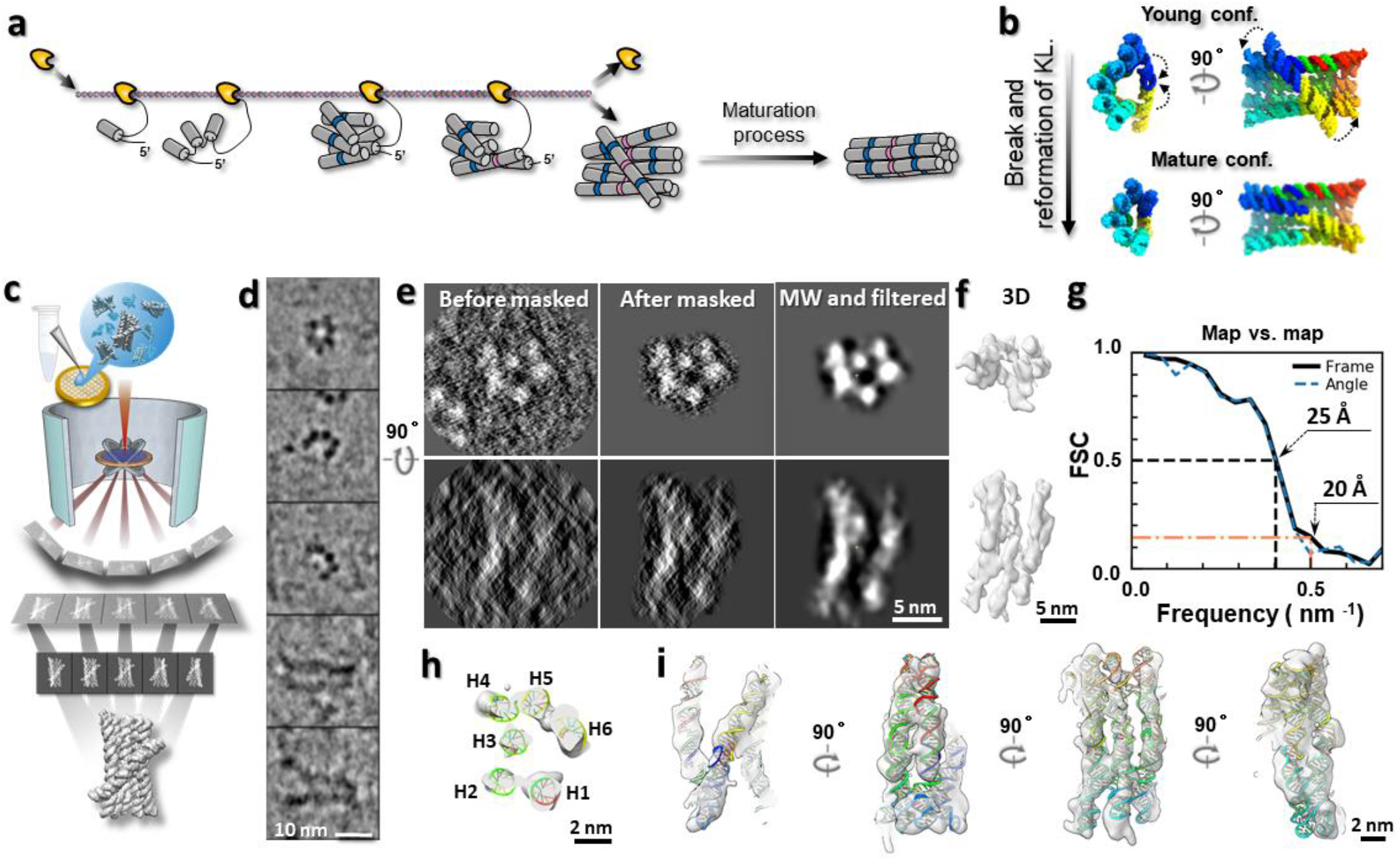
RNA origami folding process and cryo-ET 3D reconstruction. **a**, A schematic of the cotranscriptional folding and maturation processes of an RNA origami 6-helix bundle with clasp (6HBC). **b,** Atomic models of “young” (7PTK) and “mature” (7PTL) conformations of 6HBC determined by cryo-EM SAP method^11^, colored from 5’ (blue) to 3’ (red) with arrows indicating the proposed maturation mechanism. **c,** A schematic introducing the cryo-ET and 3D reconstruction methods. **d,** Five representative cryo-EM images of particles, with examples of helix features, acquired at an electron dose of 50 e^−^Å^-2^. **e,** The 3D reconstruction of an individual particle by cryo-ET and IPET from a total electron dose of 168 e^−^Å^-2^, displayed from two perpendicular directions. The central slices before and after applying the soft-masks and noise reduction for 3D reconstruction are compared to that after missing-wedge (MW) correction and low-pass filtered to 8 Å. **f,** The final 3D map view from the corresponding view directions. **g,** The resolution measured by FSC curves two half-maps reconstructed using the even-odd frames (solid line) and tilt angle (dash line) at FSC=0.5 and 0.143, respectively. **h,** The fitting model demonstrated by its central cross-section, **i,** The 3D map with the fitting model view from the four directions.

Cryogenic electron tomography (cryo-ET) has the potential to be an ideal tool for capturing non-averaged intermediate conformations of individual RNA molecules, although it has traditionally been used to study the ultrastructure of large biological objects, such as bacteria^14^ and cell sections^15^, which are imbedded in vitrified thin ice and imaged from a series of tilt angles. The cryo-ET instrument has the resolution capability to achieve near-atomic resolution structure of ribosome in bacterial cell^16^ using subtomogram averaging. However, determining the non-averaged 3D conformation of an individual biomolecule remains challenging^17^ due to the small size of the target, low image contrast, and radiation damage related dose limit^18^, especially under the condition of predicted cryo-ET theoretical resolution limit of ∼15-20 Å^19, 20^.

To expand the cryo-ET capability for studying small biological objects, the individual-particle electron tomography (IPET) method was developed by optimizing several experimental and image processing protocols^21–24^. This method has been used to determine the non-averaged conformations of individual biomolecules, characterize the structural varieties of several floppy or flexible molecules, including lipid-related proteins^25, 26^ and DNA nanostructures^27, 28^, at low to intermediate resolution.

To further push the cryo-ET capability near its theoretical resolution limit, we continuously refine the experimental conditions, such as optimized electron dose, image tilting step, and sample supporting film, to resolve nucleic acid double helices and dynamic processes of RNA folding.

### Cryo-EM image of RNA particles

To gain more insight on the transition, the purified 6HBC RNA sample was studied by cryo-ET (**Fig. 1c**). The sample was frozen ∼10 hours after the start of transcription in order to capture the molecules in the transition between the young and mature states^11^. Survey cryo-EM images show particles that were evenly distributed (**Extended Data Fig. 2a**). Representative zoomed-in images revealed that some particles contain ∼20 Å spots arranged in rings or half-rings with diameters of ∼60-100 Å, while other particles show fibers of ∼20 Å width and ∼150-200 Å length arranged in parallel bundles of ∼60-100 Å width (**Fig. 1d, Extended Data Fig. 2b**). These features were confirmed by negative staining (NS) EM (**Extended Data Fig. 2c,d**). The observation of dot- and fiber-type features suggests that single RNA helixes with a diameter of ∼20 Å was visible and the general shapes were consistent with the reported top and side views of the 6HBC^11^.

### Assessment of electron-dose limit

To assess the electron-dose damage limit to this RNA sample, we acquired untilted cryo-EM images of the same specimen area at a constant dose of 50 e^−^Å^-2^ for a total of 20 times (**Extended Data Fig. 3a**). Although all images were acquired at the same dose (50 e^−^Å^-2^), the first image contains the radiation damage caused by a dose range of 0 -50 e^−^Å^-2^, while the last image contains that of 0 – 1,000 e^−^Å^-2^. Tracking particles from the series revealed that most particles retained their outer shape at doses up to ∼200 e^−^Å^-2^ and some even up to ∼350 e^−^Å^-2^. However, the structural details and image contrast of particles gradually degenerated as the dose increased (**Extended Data Fig. 3b**). To quantitatively measure the degradation, we used the first images (50 e^−^Å^-2^) of 144 particles as a reference standard to calculate the FSC curves against the corresponding images taken after higher dose had caused damage (**Extended Data Fig. 3c**). The FSC curves showed a decay in correlation (**Extended Data Fig. 3d**), with higher frequencies degrading faster than lower frequencies. Specifically, the 10 Å information was lost after an accumulated dose of 100 e^−^Å^-2^, while the 20 Å information could tolerate an accumulated dose up to ∼350 e^−^Å^-2^, indicating that high-resolution details are more susceptible to damage than low-resolution features. A similar dose-resolution degradation relationship was also observed in a protein study^29^.

The measured electron-dose limit to this RNA sample seems much higher than that used for protein study. A possible explanation is that RNA is at least 4 times less susceptible to radiation damage compared to protein due to its unique physical properties of the aromatic bases and phosphate backbone^30, 31^, which has been highlighted by cryo-EM advances in RNA structural determination^32^.

### Structure of an individual RNA particle at ∼25 Å resolution

To test the IPET capability for the 3D structure of an individual particle, we used cryo-ET to track and image a molecule during specimen tilting (**Extended Data Fig. 4a**). We then used IPET to iteratively align the tilt images for an *ab initio* 3D reconstruction of this targeted particle at the resolution of 25 Å (**Fig. 1e-g, Extended Data Fig. 4b-g, Supplementary Video 1**), measured based on the frame map-map Fourier Shell Correlation (FSC) analysis at FSC=0.5. The achieved resolution is not related to the soft-boundary masks that were used (**Extended Data Fig. 4h,i**). After low-pass filtering to 8 Å followed by Gaussian and median filtering, the 3D map displayed the particle with dimensions of approximately 170 × 100 × 100 Å, composed of six rod-shaped densities with diameter of ∼20 Å (**Fig. 1h**). The latter dimension matched the diameter of the RNA helix, indicating that the helix is visible, and its orientation can be determined though rigid-body docking of each of the four helical segments within each coaxially stacked helix of the 6HBC model into this density map (**Fig. 1i, Extended Data Fig. 4j**).

### Resolution distribution of 170 RNA particles

Next, we investigated the effect of electron dose related damage on the cryo-ET 3D maps reconstructed by the IPET method. We achieved a total of 170 individual-particle 3D maps from the tilt series using a total dose ranging from 54 to 597 e^−^Å^-2^ (**Fig. 2a-d, Extended Data Fig. 5-7, Supplementary Fig. 6-175**). Our examination of the 3D maps and cross-sections revealed that, although helix elements were still discernible at all doses, the fitting of models to densities became less reliable at highest dose, i.e. 597 e^−^Å^-2^ (**Extended Data Fig. 6**). Moreover, the low dose of 54 e^−^Å^-2^ produced slightly noisy, while the high dose appeared slightly blurry (**Fig. 3a-c**). Notably, the 3D reconstruction achieved from the dose of 168 e^−^Å^-2^ showed a near identical structure to that obtained by removing a portion of the higher dose data (**Supplementary Fig. 2**), the so-called dose-filtering method, suggesting that the low-dose portion of the data (corresponding to the dose of 136 e^−^Å^-2^) dominates the final reconstruction, which is consistent with a study of proteins^29^.

**Fig. 2:**
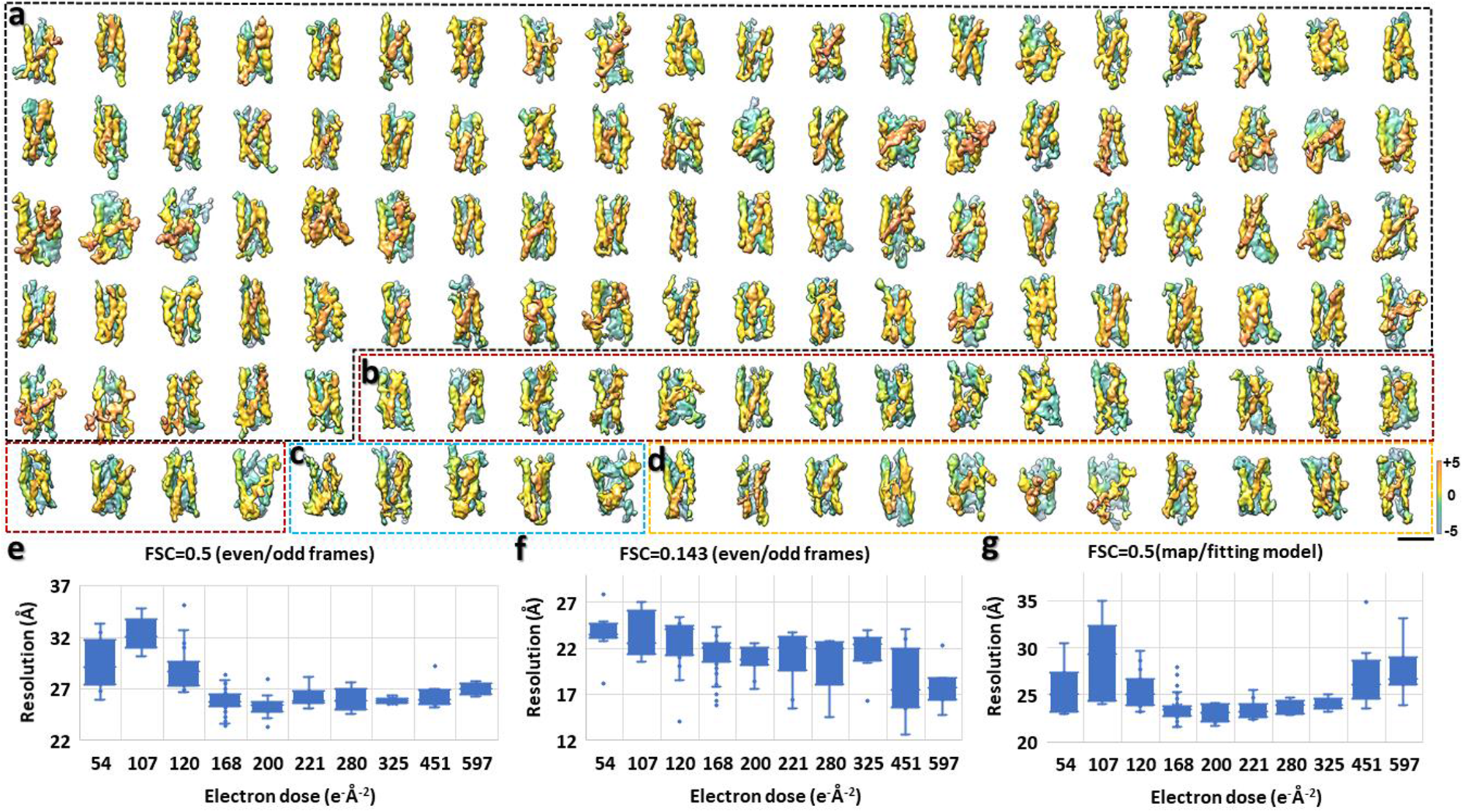
A gallery of 120 individual-particle 3D reconstructions of RNA origami 6HBC and the resolution distribution against electron dose. **a,** A total of 85 IPET 3D density maps reconstructed at a total electron dose of 168 e^-^Å^-2^. **b,** 19 maps reconstructed at a dose of 120 e^-^Å^-2^. c, 5 maps reconstructed at a dose of 107 e^-^Å^-2^. **d,** 11 maps reconstructed at a dose of 54 e^-^Å^-2^. **e,f,g,** Plots of the resolutions of IPET 3D maps against their acquired electron doses. The resolutions are estimated by three methods/criteria. The 3D maps with a dose of 200 – 597 e^-^Å^-2^ are shown in **Extended Data Fig. 6**, and the data of the plots are shown in **Supplementary data table 1**. Scale bars are 100 Å.

**Fig. 3:**
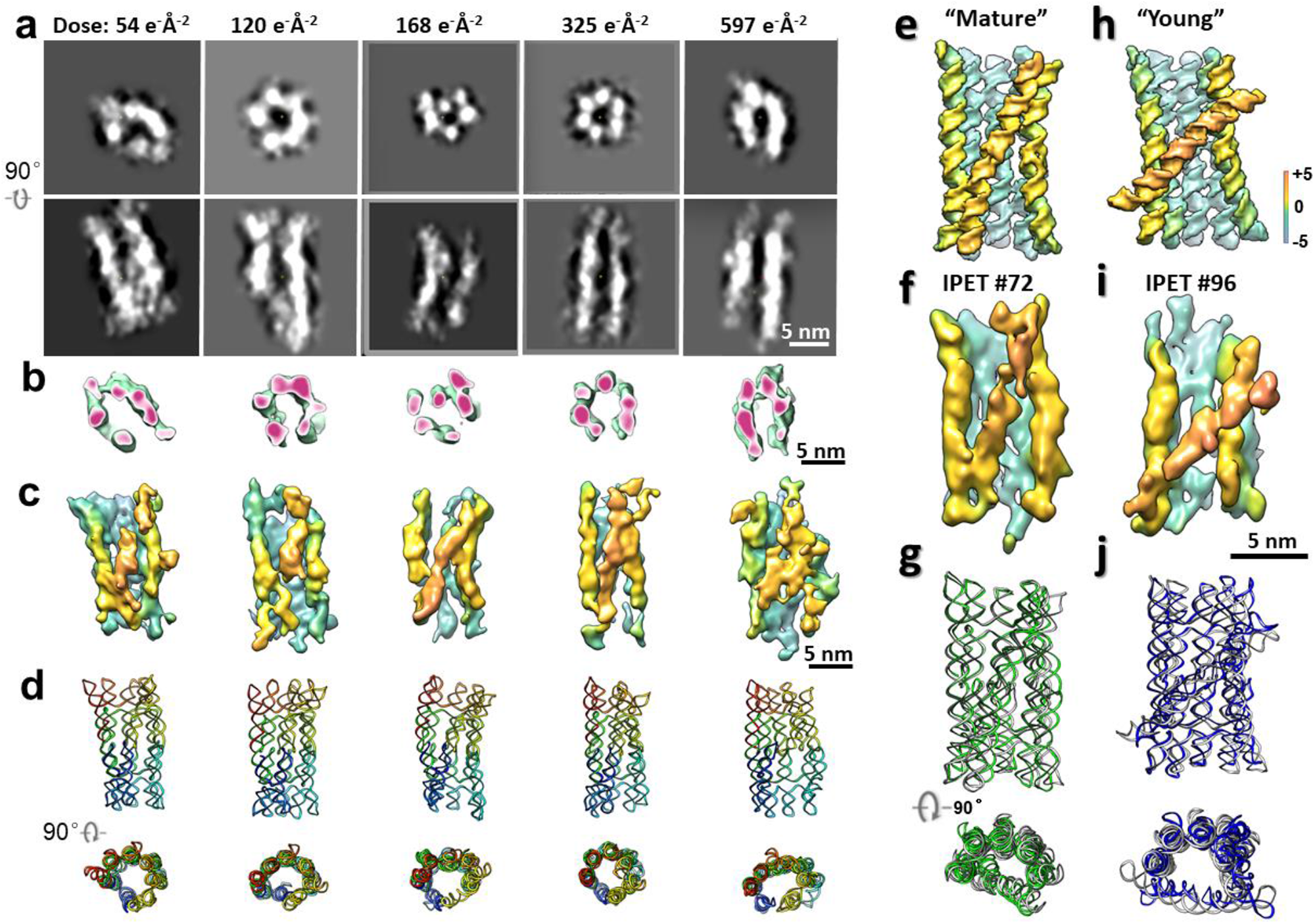
The effect of electron dose on IPET 3D reconstructions. **a,** Examples of IPET 3D reconstructions achieved at various total electron doses ranging from 54 to 597 e^−^Å^-2^, with corresponding central slices shown (particle # 10, 27, 67, 148, 165 from left to right). **b,** The cross-sections of 3D reconstructions. **c,** The 3D reconstructions. **d,** The fitting models in two perpendicular views, with rainbow-colored sequences indicating the order from 5’ to 3’ end. **e,** The 3D map of the SPA “mature” conformation reported by conventional SPA method (obtained at a dose of 60 e^-^Å^-2^)^11^. **f,** The IPET representative structure (particle #72, obtained at a dose of 168 e^-^Å^-2^) with a conformation similar to the SPA “mature” conformation. **g,** The superimposed “mature” and IPET structures from two perpendicular directions, with RMSD of 7.2 Å. **h-j,** A similar comparison of the SPA “young” conformation and IPET structure (particle #96, achieved at the dose of 168 e^-^Å^-2^), which showed an RMSD of 9.4 Å.

To assess the resolution of the reconstructed maps, we employed several methods that include recommended metrics^11^. i) The frame map-map FSC analyses (**Fig. 2e, 2f**): The FSC curves between two half-maps reconstructed, respectively, from the even and odd frames at each tilt angle were computed, in which the, frequencies at FSC=0.5 and 0.143, respectively were used as the assessed resolutions; ii) The tilt map-map FSC analyses (**Supplementary Data Table 1**): it is same as above, except two half-maps were reconstructed from the even and odd indexes of tilt images; The results showed above two methods provides near identical resolutions. iii) The map-model FSC analyses (**Fig. 2g**): the FSC curve between the map and its fitting model generated map was calculated, and the frequency at FSC=0.5 (**Fig. 2g**) was used as the evaluation of resolution; iv) The visible structural features (**Supplementary Fig. 3**); v) Comparison to the conventional SPA reconstruction: the “mature” SPA structure determined at a dose of 60 e^−^Å^-2^ was superimposed to the IPET maps for comparison (**Supplementary Fig. 4**); and vi) The reconstructions by a third-party cryo-ET 3D reconstruction software (*i.e.* IMOD^33^) (**Supplementary Fig. 5**).

Statistical analysis of the resolution defined by the map-map FSC=0.5 criteria revealed a U-shaped distribution of resolution against dose (**Fig. 2e**). The highest resolution of ∼23 Å was achieved at a dose of 168 e^−^Å^-2^ and 200 e^−^Å^-2^, respectively, in which the dose of 200 e^−^Å^-2^ had the highest averaged resolution (∼25.4 Å), and the dose of 168 e^−^Å^-2^ had the second-highest averaged resolution (∼25.9 Å). Lower doses (ranging from 120 to 54 e^−^Å^-2^) resulted in lower averaged resolutions (in the range of 28.9-31.6 Å), possibly due to decreased signal-to-noise ratio (SNR) and increased errors in alignment. Higher doses (ranging from 221 to 597 e^−^Å^-2^) resulted in slightly lower resolution (in the range of 26.0-26.9 Å), which could be due to the increased attribution of the high-dose portion of the data *i.e.,* a higher percentage of damaged images.

The analysis of the resolution measured by the map-model FSC=0.5 metric (**Fig. 2g**) agreed with this U-shaped distribution and confirmed that doses of 168 to 325 e^−^Å^-2^ had relatively higher average resolutions in the range of ∼23-24 Å. However, the distribution of resolution measured based on the gold-standard metric ^18^ (the map-map FSC=0.143 criteria, **Fig. 2f**) did not align well with this observation. Although a U-shaped distribution of resolution (20-23 Å) remained within the lower dose range of 54-325 e^−^Å^-2^, the unexpectedly increasing resolution from 22 to 17 Å at higher dose (i.e. 325 and 597 e^−^Å^-2^) broken the U-shaped distribution and destroyed confidence in the measured resolution by this criterion.

The above analyses demonstrated that the highest resolution can be achieved within the dose range of 120-325 e^−^Å^-2^. Although it is well established that RNA is less susceptible to radiation damage compared to protein ^30, 31^ and protein structures at near-atomic resolution have been successfully solved using a dose within the range of 120-140 e^−^Å^-2^ ^16^, a dose exceeding 200 e^−^Å^-2^ seldom been employed in the field of cryo-EM/cryo-ET^34, 35^, primarily due to early measured dose limits that were significantly lower than those used today. Consequently, in order to address potential concerns regarding radiation damage in the data, only reconstructions obtained using low doses (up to 168 e^−^Å^-2^) were utilized for subsequent structural analyses.

### Mapping of the conformational landscape

To investigate the structural intermediates during the maturation folding process of the 6HBC RNA, we analyzed the 120 3D reconstructions achieved at a dose of 168 e^−^Å^-2^ and below (**Fig. 2a-d**, **Supplementary Video 1**). These low-dose reconstructions displayed a wide range of morphologies (**Extended Data Fig. 8**), including structures that were very similar to the “mature” conformation (**Fig. 3e,f**) and “young” conformation (**Fig. 3h,i**) previously determined by conventional cryo-EM SPA ^11^. Superimposing the SPA structures and IPET models showed a root-mean-square deviation (RMSD) of 7.0-9.4 Å, indicating a very close resemblance (**Fig. 3g,j**). The averaged resolutions of all 3D reconstructions were ∼26.8 Å for map-map FSC=0.5 metric (**Fig. 2e**) and ∼24.2 Å for map-model FSC=0.5 metric (**Fig. 2g**), respectively.

To further understand the structural variety of 6HBC, the 120 low-dose models were aligned and superimposed, revealing a wide range of conformations (**Fig. 4a**). To quantitatively measure the structural difference, RMSD was calculated for each of the two models (including the “young” and “mature” SPA models) for comparison. The resulting variety was displayed in a dendrogram by hierarchical sorting based on RMSD (**Fig. 4b, Extended Data Fig. 8**). Notably, the reported “young” and “mature” conformations were categorized into two primary classes (with a mean RMSD of ∼69 Å between the conformations of these two classes) (**Fig. 4b**), suggesting a major energy barrier between these two conformations. More particles were found in the “mature” conformation, suggesting that the majority of molecules in the sample had passed this major energy barrier, *i.e.*, breakage and reformation of KL ^11^. Interestingly, several sub-classes existed within both major classes, with RMSD ∼28 Å, which could be related to additional energy barriers.

**Fig. 4:**
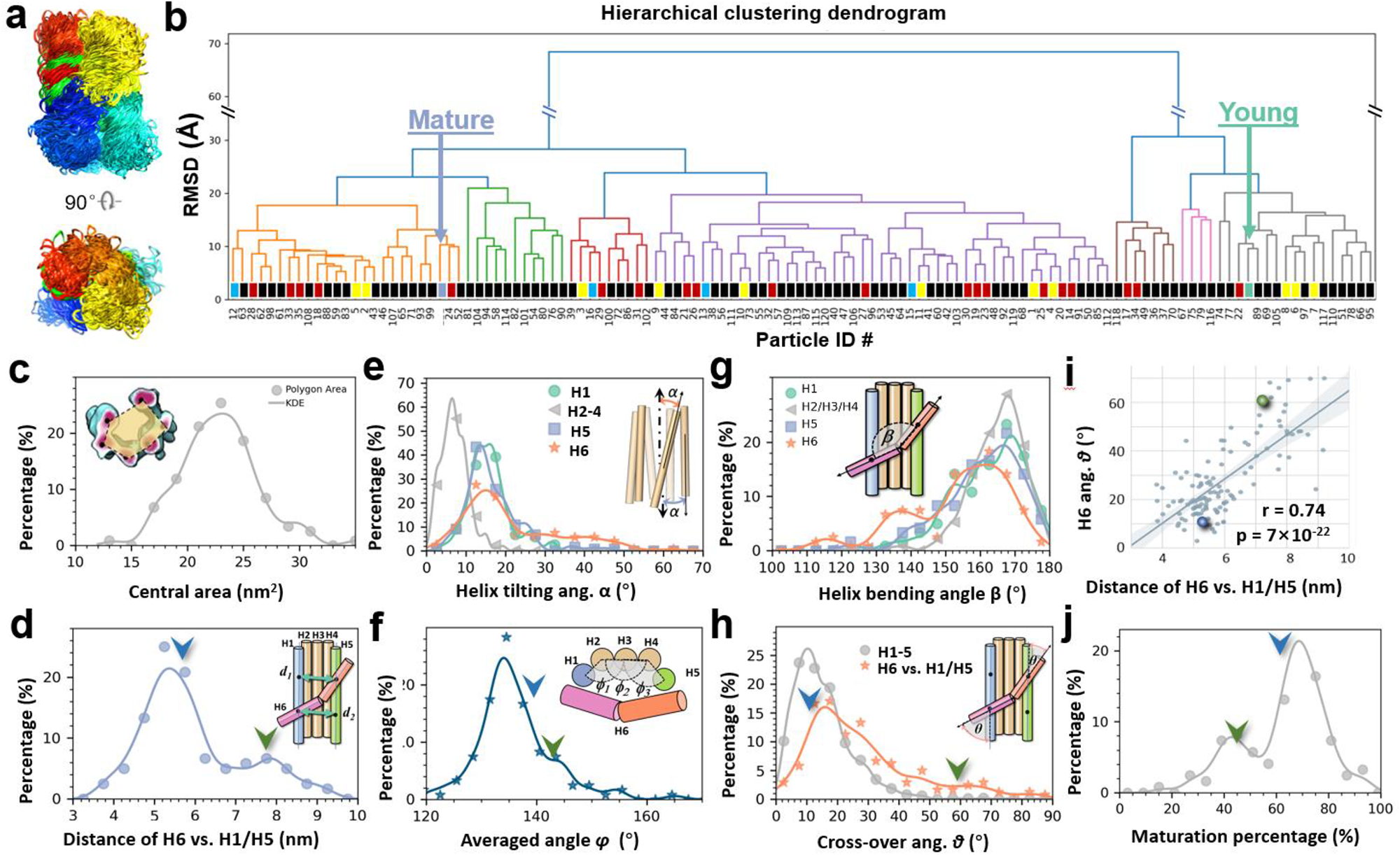
Statistical analyses of the structural variety. **a,** The superimposition of 120 IPET structural models in two perpendicular views, with the order of sequences colored by rainbow colors from the 5’ to 3’ end. **b,** A dendrogram of the Hierarchical clustering analysis of RMSD values, with structures on the X-axis and cluster distance on the Y-axis. The structures of the “young” and “mature” conformations determined by the SPA method^11^ are included and labeled by the colors of “medium aquamarine” and “polo blue,” respectively, for comparison. The structures of 168 e^−^Å^-2^, 120 e^−^Å^-2^, 107 e^−^Å^-2^, and 54 e^−^Å^-2^ are marked by rectangle boxes in black, red, cyan, and yellow, respectively. **c,** A histogram of the helical bundle center area. **d,** A histogram of the averaged distance *d̄* of H6 against the H1 and H5, indicated by the *d_1_* and *d_2_*. **e,** A histogram of the helix tilting angle *α* against the bundle center axis. **f,** A histogram of the helical bundle angle *φ*, formed within H1-5. **g,** Characterization of the curvature of the H6 helix by the bending angle β between its two halves. **h,** A histogram of the crossover angle *θ* of H6 against its connected H1 or H5, which is compared to the crossover angle formed among the rest of the helices. **i,** A correlation analysis of the H6 crossover angle *θ* against the averaged H6 distance, *d̄*. **j,** A histogram of the maturation percentage (MP), with the green and blue arrows/balls indicating the SPA determined “young” and “mature” conformations, respectively^11^. All measured data are fitted by kernel density estimation (KDE).

To validate whether the classification of RMSDs was affected by the electron dose, the RMSD of the particle achieved at each dose was annotated (colors in **Fig. 4b**). The uniform distribution across the observed conformational space indicated that the RMSD had no correlation to the dose. This validation was further conducted by using the two SPA structures as references, computing the RMSDs against IPET models, and then plotting the distribution against the dose (**Extended Data Fig. 9**). The result confirmed that the RMSDs had a poor correlation to the electron dose.

### Statistical analysis of structural parameters

To analyze the detailed structural variety within the molecules, the distances and angles among the helices were measured (**Fig. 4c-j**). The distribution of the molecular central area defined by the six helical centers showed a range from ∼1,400 to ∼3,400 Å^2^ (**Fig. 4c**), suggesting an average distance between two adjacent helices in the range of ∼23 to ∼35 Å. Among the helices, Helix (H) 6 (H6) had the longest distance to H1 and H5, which contributed to the increase in the averaged distance between other helices. Nevertheless, the distances indicate a co-existence of loose conformation (with a one-helix width gap between two nearby helices) and a compact conformation (where helices are packed tightly), in which the compact structure is the lowest free-energy state as designed (**Fig. 1a,b**) and studied by fitting SAXS curves ^11^.

Among the six helices, H6 is the last helix to form the KL and join the helical bundle. Thus, the conformation of H6 relative to other helices is one of the most sensitive parameters to detect the maturation folding process. The H6 conformation can be quantitively measured by the averaged distance between H6 and H1/H5, i.e., *d̄*, which is the averaged distances of *d*_1_ (measured between the crossover points of H1/H6 and H5/H4) and the *d*_2_ (measured by the crossover points of H5/H6 and H1/H2) (**Fig. 4d**). The distribution of *d̄* is found to have a range of ∼30 to ∼100 Å with two major peak populations at ∼52 Å and ∼79 Å. Notably, the two cryo-EM SPA structures were located close to these two major population peaks, suggesting that they represent low-energy states ^36^. This analysis confirms that the distance *d̄* is a sensitive parameter to probe the conformational change of 6HBC during the maturation process.

The tilt of H6 was measured by the angle *α*, which is defined as the angle between H6 and the bundle axis (**Fig. 4e**). H6 was found to have a wider distribution of *α* in a range of 0° to ∼60° compared to the other helices, which had an *α* range of ∼0° to ∼30°. The major peak of the H6 *α* distribution was at ∼15°, which is close to the tilting angles reported in the “mature” conformations, *i.e.* ∼17° ^11^. H6 also shows a broader, flatter distribution in a range from ∼30° to ∼55°, which includes the 47° tilting angle reported in the “young” conformations ^11^. The histogram of *α* measurements shows that the “mature” conformation was about ∼6-10 times more populated than the “young” conformation, indicating that the majority of the molecules were released from the energy trap of H6 KL.

We next measured two curvatures inside each molecule. One was the helical bundle curvature that is formed by H1-5 and measured by the average of three φ angles (**Fig. 4f**). The distribution showed a narrow range of ∼120° to 150°, except for a few φ values up to ∼165°. The locations of the young and mature conformations in the distribution reveled that the sample was closer to the mature conformation. The other curvature was the H6 curvature, which was measured by the bending angle *β* at the KL connecting the helix segments (**Fig. 4g**). As expected, the distribution showed H6 is the most bent helix with population peaks at ∼160° and ∼130°. In this distribution, the “young” and “mature” conformations had nearly identical H6 *β* values (166° vs 165°), and thus, it is an insensitive parameter for distinguishing the maturation process.

The crossover angle θ at the crossover points of H6/H1 and H6/H5, respectively showed a widely distributed range, in which the “mature” and “young” conformations were significantly different, suggesting that *θ* is a sensitive parameter for measuring the maturation process (**Fig. 4h**). The cross-correlation analysis of *θ* and the H6 distance *d̄* showed a certain level of correlation (r=0.74) suggests that the tilting of H6 controls the distance between H1 and H5 (**Fig. 4i**). The presence of more particle around the “mature” conformation than the “young” indicates the sample is closer to the mature state.

Based on the statistical analysis, we found that the distance *d̄*, the H6 tilting angle *α* and the H1-H5 bundle curvature angle φ are the most sensitive parameters characterizing the maturation process. We therefore introduce the “maturation percentage” (MP) as a measure of maturation by equally weighing these three parameters, followed by scaling to a percentage (see the definition and formula in **Methods**). The histogram of MPs shows that the 120 structures are within a range of ∼17% to ∼95%, with two population peaks at ∼45% and ∼70%, respectively (**Fig. 4j**). These peaks were close to the MPs of “young” and “mature” structures, *i.e.* ∼47% and ∼64%, respectively, suggesting that both cryo-EM-derived structures were in low energy states.

### Exploring the maturation folding path

The MP was utilized as a simple parameter to sort RNA conformations along a hypothetical folding pathway. Extracting representative structures from this pathway aids in visualizing the primary structural changes that occur during the maturation process (**Fig. 5a**). Alternatively, all structural intermediates can be presented as snapshots of a movie, which provides insight into the flexibility and structural changes along this trajectory (**Supplementary Video 1**). To obtain a better understanding of the folding path, MP values were plotted against two of their three parameters, i.e. the angles *α* and *φ*, and then fitted with a curve surface to represent a pseudo free-energy landscape ^37^ (**Fig. 5b**). The 3D surface comprises local mountains, hills, valleys and canyons, where a higher MP corresponds to a lower energy state, while a lower MP indicates a higher energy state.

**Fig. 5:**
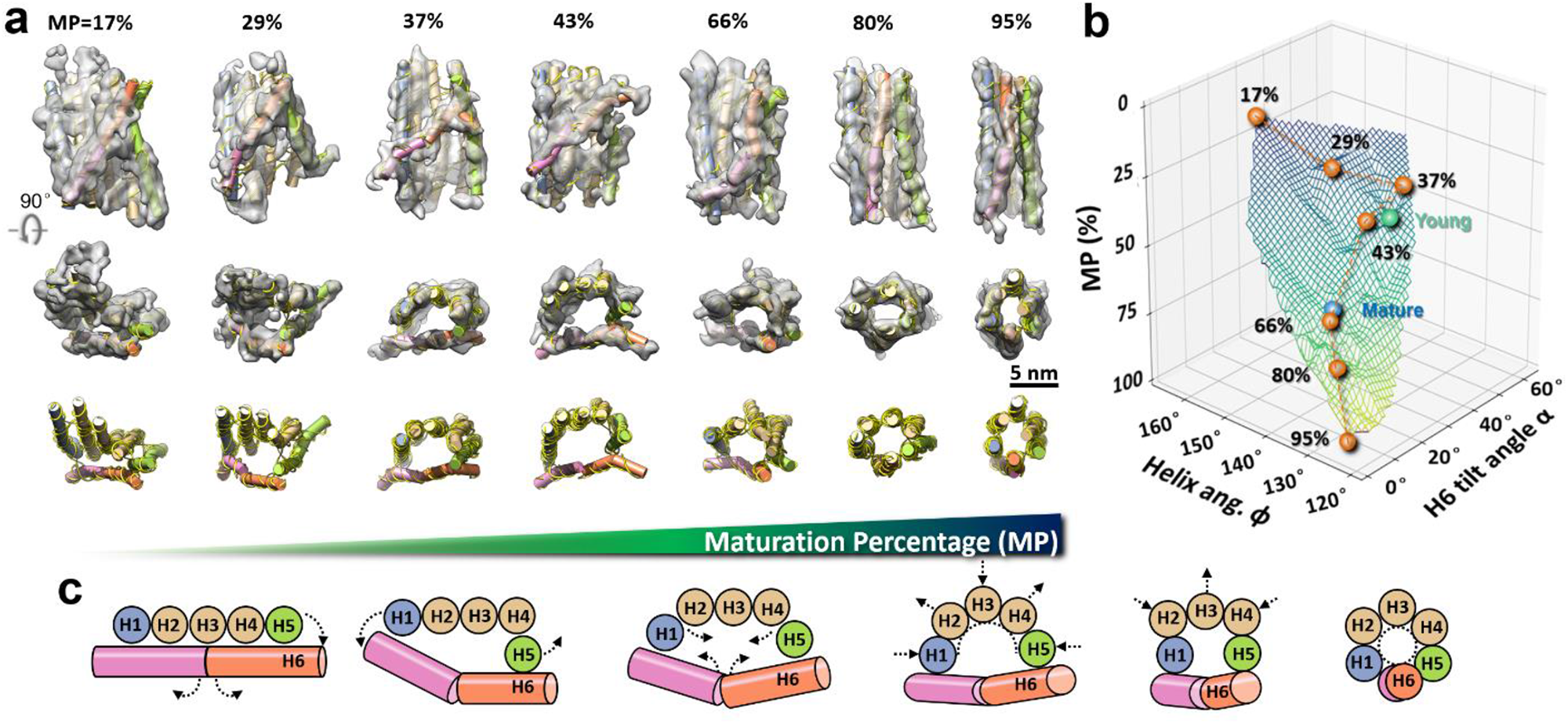
A hypothesis of the 6HBC maturation folding process. **a,** Seven representative particles shown as 3D maps and tertiary models in the order of their maturation percentage (MP) values. The map is displayed from two perpendicular directions with their fitting models, and the corresponding tertiary structures are shown in the bottom row. **b,** A fitted curvy surface of the MP values distributed against bending angle *α* and *φ*. One possible folding pathway is presented by seven representative particles shown as orange balls. The segmentations connect the conformations by the order of their MP values, which passed through the “young” and “mature” conformations indicated by green and blue balls, respectively. **c,** Schematics of a folding landscapes that show a maturation pathway/process of 6HBC from “a paddle with a stick”-shaped conformation to a “helix bundle”-shaped compact conformation.

To clarify the folding mechanism, seven representative particles were labeled on the MP surface, illustrating a potential folding pathway or energy minimizing trajectory (indicated by orange balls and lines in **Fig. 5b**), which passed through the “young” to “mature” conformations (indicated by green and blue balls in **Fig. 5b**). It is observed that there is a significant increase in MP from the “young” to “mature” conformation, and the primary conformational change occurs in the H6 *α* angle, while the helix angle *φ* does not change substantially – in agreement with the observation that H6 becomes parallel to the other helices in the mature state.

We can now examine the structural changes occurring along this representative maturation pathway. At MP = ∼17%, the five helices (H1-5) form a flat sheet, with H6 positioned close to the sheet surface, potentially representing a rare, trapped state. At MP = ∼29%, H5 tilts away from the sheet formed by H1-4, resulting in an increase in the central cavity. At MP = ∼37%, H1-5 orient in parallel to form a bundle with a central cavity. At MP = ∼43%, H1-5 form a semicircle bridged by H6, and the central cavity reaches its maximum area. This conformation is comparable to the “young” conformation ^11^. At MP = ∼66%, H1-5 assume a more compact “zig-zag” conformation, reducing the area of the central cavity. This conformation is similar to the “mature” conformation, and its transition requires the temporary dissociation of the H6 KL, as proposed previously^11^. At MP = ∼80%, the central cavity is further reduced as H6 joins the helical bundle by aligning itself parallelly with the other helices. At MP = ∼95%, all helices are completely parallel and in close proximity to each other, indicating that the final stage of the maturation process has been achieved. These landmark structures provide a hypothetical model of the dynamic self-folding process (**Fig. 5c**). A video was created by sequentially morphing between the structures by their MPs to visualize the pseudo dynamic process of flexibility and self-folding along the MP trajectory (**Supplementary Video 1**).

## Discussion and conclusions

In this study, we investigated the folding and maturation process of an RNA molecule using cryo-ET of individual particles without the need for selecting a homogeneous population, averaging different particles or staining the sample. This approach allowed us to achieve a resolution of up to 23 Å (measured based on the map-map FSC=0.5 criteria), which is close to the theoretical limit of cryo-ET of 20 Å (without subtomogram averaging) ^18^. To achieve this resolution, we utilized advanced equipment, including a state-of-the-art cryo-EM instrument, a direct-detection camera, a graphene oxide (GO) film, and optimized electron doses and tilt angle steps. Importantly we show that for this RNA sample, total doses of up to 325 e^−^Å^-2^ did not result in loss of helical features from the cross section of our reconstructions. Indeed, the best map-map FSC was achieved with a total dose between 168-200 e^−^Å^-2^. It has been long known that nucleotides are less sensitive than amino acids to electron induced damage ^30–32^, with electron diffraction spots from adenosine crystals fading 10 times slower than those from valine crystals^38^. This is likely due to the conjugated pi orbitals in the adenosine ring being more capable of absorbing and distributing the excess energy from inelastically scattered electrons than sigma bonds. We hypothesize that the extended pi stacking between adjacent bases in RNA, particularly in the nearly 100% double stranded RNA origami, further hardens these structures to electron-induced radiation damage.

The statistical analysis of the structural parameters of RNA origami has facilitated the development of a metric for measuring the maturation process along a hypothetical folding pathway. The IPET data capture a snapshot of the 3D structure of folding intermediates in the sample at the given time. The 6HBC is a unique case where the KL of H6 forms a kinetic trap, preventing the rest of the bundle (H1-H5) from assuming its preferred 120° bundle shape.

However, the helix packing forces, driven by the hydrogen bonds between sugar-phosphate backbones of the helices and the stacking strain in the junctions, squeeze H6, causing its KL to break and reform by gradually rotating 180° until the bundle matures ^11^. This global behavior of the bundle compaction process is observable in the IPET data, which is displayed by two primary peaks. During this slow transition process, the two observed peak populations can be roughly interpreted as two energy wells based on the Boltzmann distribution in thermodynamics^36^. The state with lower energy will always have a higher probability of being occupied.

Furthermore, the data revealed several unusual or exotic conformations, including the connection of two particles through a helix (potentially indicating intermolecular KL interaction), particles exhibiting a five-helix structure (possibly transcription termination products leaving H1 partially single-stranded), and the presence of loop structures (where KLs may misconnect). Given that the sample underwent maturation for several hours, these conformations are likely dead-end folding products that have become trapped in a deep energy well, rendering their escape impossible.

The self-assembly of RNA origami structures after cotranscriptional folding^9^ provides a simple model system for investigating the complex folding of natural RNAs, such as the ribosomal RNAs. Utilizing simple model systems can help disentangle complex RNA folding pathways and identify the contributions of tertiary contacts and protein chaperones to folding^39^. We propose that the IPET approach described in this study can be applied to complex natural RNAs to elucidate folding pathways occurring in distinct phases^40^, achieved through the compaction of secondary and tertiary interactions. Moreover, earlier stages of folding processes can be explored by freezing samples at earlier stages or during the transcription process itself to analyze the population of species and conformations at these timepoints. The IPET method can complement conventional SPA analysis by providing a global analysis of conformational diversity via the structures of individual particles. IPET can also be used to study snapshots of dynamic processes in response to temperature, pH, and polymerization reactions, such as folding during transcription and translation and can benefit from companion techniques such as time-resolved cryo-EM^41^, chemical probing^42, 43^, and single-molecule FRET microscopy^44–46^. As IPET data acquisition and analysis become more automated, a large database of structural conformations for each individual macromolecular species can be produced. This database of dynamic structures will be a rich dataset for machine learning methods to predict dynamical properties beyond the single static structure of a macromolecule^47, 48^. By acquiring time-resolved IPET data of multi-stage dynamic structures and complex folding pathways, machine learning methods may allow for realistic predictions and design of the kinetics of biomolecular machines and self-assembly processes.

## Acknowledgements

We would like to express our gratitude to Drs. Mingdong Dong and Jinghui Tao and Ms. Amy Ren for their insightful discussions and comments. We also extend our appreciation to Dr. Dan Toso at Cal-Cryo-EM center of QB3-Berkeley for his supporting in cryo-ET imaging. The work at the molecular foundry, LBNL was supported by the Office of Science, Office of Basic Energy Sciences of the United States Department of Energy (contract no. DE-AC02-05CH11231). The single particle analysis data set was collected at EMBION Cryo-EM Facility at iNANO, Aarhus University, Denmark. We acknowledge following research grants: the US National Institutes of Health grants of R01HL115153, R01GM104427, R01MH077303 and R01DK042667 (GR, JL, MZ); Independent Research Fund Denmark grant 9040-00425B (ESA, EKSM); Canadian Natural Sciences and Engineering Research Council grant 532417 (EKSM); European Research Council (ERC) Consolidator grant 683305 (ESA, CG, EKSM) and Novo Nordisk Foundation Ascending Investigator grant 0060694 (ESA, CG).

## Author contributions

Experiments were designed by J.L., G.R., and E.S.A. Sample preparation was carried out by E.K.S.M. IPET cryo-ET experiments, including data acquisition and processing, were performed by J.L. SPA cryo-EM data collection and analysis performed by E.K.S.M. The templates for modeling were provided by E.K.S.M. and E.S.A. IPET cryo-ET models were built by G.R. and J.L., and the models were refined by M.Z. and J.L. Data was interpreted and analyzed by all authors. J.L. and G.R. created the figures and video. The initial draft of the manuscript was written by J.L. and G.R., and all authors contributed to revising the manuscript.

## Data and material availability

The RNA sequence can be found in the **Supplementary Materials**. We have deposited the 3D density maps of 170 cryo-ET IPET reconstructed RNA origami (6HBC) particles in the EMDB under accession codes EMD-25230 to EMD-25261, EMD-25270 to EMD-25357, EMD-40353, and EMD-40356 to EMD-40404. The density maps and fitting models were also deposited to Zenodo.org (doi: 10.5281/zenodo.7811352). The related experimental parameters and measured resolutions of these 170 density maps were listed in **Supplementary Data Table 1**.

## Methods

### Transcription and purification of RNA origami

The RNA origami 6HBC sample was provided by Dr. Ebbe S. Andersen’s group, as reported in^11^. In brief, synthetic DNA containing the a t7 promoter, the 6HBC design, 3’ BsaI linearization site and flanking restriction enzyme sites for cloning was ordered from IDT in the form of a G-block, which was then cloned into a modified pUC vector. This plasmid was transformed into DH5-alpha cells and individual clones were sequence verified. Large-scale production of plasmid was performed using a MaxiPrep kit from Machery Nagel. Plasmid was subsequently linearized using BsaI (NEB), three times phenol chloroform extracted and then ethanol precipitated. Linear plasmid was re-suspended in RNase-free water and diluted to 0.5mg/mL.

Transcription reactions were setup in a buffer containing 40mM Tris-Cl (pH 8.0) at 37°C, 1 mM spermidine, 0.001% Triton X-100, 100 mM DTT, 12 mM MgCl2, 8 mM NTP mix and 0.05 mg ml–1 template DNA. Transcriptions are started on the addition of in-house prepared T7 polymerase and transcription reactions are carried out for 3 h at 37 °C. Precipitated inorganic pyrophosphate is pelleted by centrifugation at 17,000g for 5 min at room temperature. The transcription reaction was then loaded onto a Superose 6 column (Cytiva) equilibrated with 25 mM HEPES buffer (pH 8.0), 50 mM KCl and 5 mM MgCl2. The major RNA peak is then collected and concentrated in 10-kDa-cutoff Amicon spin concentrators. Purified 6HBC was at room temperature for approximately 6 hours before grid preparation for single particle analysis cryo-EM. After which the sample was frozen to -20°C, for weeks before being shipped on dry ice to Berkely for IPET analysis.

### TEM specimen preparation by Cryo-ET and OpNS EM

The sample was prepared for both cryo-EM and optimized negative-staining (OpNS)^49, 50^. Briefly, a 4 μl aliquot of sample (diluted concentration: 0.3 mg/ml) was placed on a single layer graphene oxide support film on lacey carbon, 300 mesh copper grid (Ted Pella Inc., Redding, CA, USA) that had been glow-discharged for 15 seconds prior to cryo-EM specimen preparation. After blotting with filter paper from one side for 3.5 seconds, the grids were flash-frozen in liquid ethane at ∼90 % humidity and 4 °C with a Leica EM GP rapid-plunging device (Leica, Buffalo Grove, IL, USA) before being transferred into liquid nitrogen for storage. In the meantime, OpNS-EM specimens were prepared as follows: a 4 μl aliquot of the sample (diluted concentration: 0.25 mg/ml) was placed on ultra-thin carbon film grids (CF-200-Cu-UL, Electron Microscopy Sciences, Hatfield, PA; Cu-200CN, Pacific Grid-Tech, San Francisco, CA USA) that were prior glow-discharged for 15 seconds. After 1 min. of incubation, the excess solution on the grid was removed by filter paper blotting. The grid was then washed with water and stained with 1 % (w/v) uranyl formate (UF). The excess UF solution was removed by blotting the grid with filter paper from the opposite side of the carbon film before air-drying with nitrogen.

### Cryo-EM and cryo-ET data acquisition

The cryo-EM specimens were imaged using a Titan Krios G2 and G3i TEM (ThermoFisher Scientific) equipped with a Gatan energy filter (Gatan, Inc., Pleasanton, CA, USA) and operated at 300 keV. Micrographs were acquired using a Gatan K3 direct electron detector in correlated double sampling (CDS) mode ^51^ and super-resolution mode with a defocus of ∼2 μm, controlled by SerialEM^52^. Un-tilted micrographs were acquired at a nominal magnification of 81 k× (corresponding to ∼0.94 Å/pixel) with an exposure time of ∼5.5 s (∼0.11 s for each frame) and a dose rate of ∼8 e^-^Å^-2^s^-1^. The tilt series were acquired using a symmetric tilt scheme starting from 0° to the highest angle. The tilt series comprised three rounds of imaging sessions.

In the first round, acquired on a Krios G2 at 81 k× magnification (corresponding to ∼0.94 Å/pixel) and a dose rate of ∼8 e^-^Å^-2^s^-1^, four tilt series were obtained in a tilting range of ±60° in a tilting step of 5° and total dose of ∼325 e^-^ Å^-2^ (at exposure time of ∼1.43 s per tilt image); two series were collected in a tilting range of ±50° in a tilting step of 5° and a total dose of ∼168 e^-^Å^-2^ (at exposure time of ∼0.88 s per tilt image); one series was obtained in a tilting range of ±51° in a tilting step of 3° and total dose of ∼280 e^-^Å^-2^ (at exposure time of ∼0.88 s per tilt image); one series was acquired in a tilting range of ±60° in a tilting step of 5° and total dose of ∼200 e^-^Å^-2^ (at exposure time of ∼0.88 s per tilt image); one series was collected in a tilting range of ±56° in a tilting step of 7° and total dose of ∼221 e^-^Å^-2^ (at exposure time of ∼1.43 s per tilt image).

In the second round, acquired on a Krios G3i at a nominal magnification of 53 k× (corresponding to ∼1.67 Å/pixel) and a dose rate of ∼2.91 e^-^ Å^-2^s^-1^, one tilt series was obtained in a tilting range of ±50° in a tilting step of 5° and total dose ∼54 e^-^Å^-2^ (at exposure time of ∼0.88 s per tilt image); one series was acquired in a tilting range of ±50° in a tilting step of 5° and total dose of ∼107 e^-^Å^-2^ (at exposure time of ∼1.75 s per tilt image); one series was collected with a tilting range of ±50° in a tilting step of 5° and a total dose of ∼451 e^-^Å^-2^ (at an exposure time of ∼7.39 s per tilt image); one series was collected in a tilting range of ±50° in a tilting step of 5° and total dose of ∼597 e^-^Å^-2^ (at exposure time of ∼9.76 s per tilt image).

In the third round, acquired with a Krios G3i at a nominal magnification of 53 k× (corresponding to ∼1.67 Å/pixel) and a dose rate of ∼3.07 e^-^Å^-2^s^-1^, two tilt series were collected with a tilting range of ±49° in a tilting step of 7° and a total dose of ∼120 e^-^Å^-2^ (at exposure time of ∼2.61 s per tilt image). The dose at 60° was 1.41 times higher than the dose at 0°, and the exposures were proportional to the inverse cosine of the tilt angle to the 1/2 power.

### Negative-staining EM data acquisition

The OpNS EM specimens were examined using a Zeiss Libra 120 Plus TEM (Carl Zeiss NTS, Overkochen, Germany), equipped with a LaB_6_ gun operating at 120 kV, an in-column energy filter, and a 4 k × 4 k Gatan UltraScan 4000 charge-coupled device (CCD) camera. The un-tilt micrographs were acquired at near Scherzer defocus and at a magnification of 80 k× (corresponding to 1.48 Å/pixel) with a dose of ∼100 e^−^Å^-^_2._

### IPET image preprocessing

Motion correction of the multi-frame from the K3 camera was conducted by MotionCor2^53^. The tilt series of whole micrographs were first aligned using IMOD^33^. The Contrast Transfer Function (CTF) was determined using the GCTF software package^54^ and then corrected by TOMOCTF^55^. To reduce the image noise, tilt series were processed using an in-house developed machine learning software (manuscript in preparation), a median-filter software and a contrast enhancement method^23^.

### IPET 3D reconstruction

In the pipeline of IPET 3D reconstruction^21^, a CTF-corrected tilt series containing a single or few RNA origami 6HBC particle was extracted from the full-sized tilt series. This allows us to perform “focused” 3D reconstruction, in which the reconstruction is less sensitive to image distortion, tilt-axis variation with respect to tilt angle, and tilt angle offset. Initially, each targeted 6HBC particle was windowed and extracted from the whole-micrograph tilt series into a small-size tilt series measuring ∼256 × 256 pixel (1.88 Å/pixel) for first round of data at 81 k× magnification, ∼320 × 320 pixel (1.67 Å/pixel) for the second round of data at 53 k× magnification, or ∼288 × 288 pixel (1.67 Å/pixel) for third round of data at 53 k× magnification.

To start the 3D reconstruction, an *ab initio* 3D map was generated as an initial model by back-projecting the small-size tilt series. During the iteration and refinement processes, a set of Gaussian low-pass filters and soft-boundary circular and particle-shaped masks were automatically generated and sequentially applied to the tilt series and projections of the references to increase their signal-to-noise ratio (SNR). To reduce the missing-wedge artifact caused by the tilt limit, the final 3D maps were subjected to a low-tilt tomographic 3D reconstruction method (LoTToR)^21^. Finally, all IPET 3D reconstructions were low-pass filtered to 8 Å using EMAN software^56^, followed by Gaussian filtering (the standard deviation is 3.0), median filter (3×3×3) and rendered in UCSF Chimera software^57^.

### Resolution analysis of IPET map

The resolution was estimated using following four methods. i) Map-to-Map frame-based FSC analysis: the FSC between two reconstructed 3D maps from two-half frames at each tilt angle (based on their odd and even tilt frames at each tilt angle) was generated using IPET aligned particle tilt series^8^. The frequencies at which the FSC curve begin falls below the values of 0.5 and 0.143 were used to represent the resolution. It should be notably that the resolution estimated by this method was under-estimated since the reconstructions were based on only the half of tilt-series, especially the super low numbers of images compared to SPA, where the total number of images is hundreds to thousands higher. As a result, the resolution measured from the two reconstructions achieved from the half data/dose should be significantly lower than that of the final reconstruction.

ii) Map-to-Map angle-based FSC analysis: The FSC between two reconstructed 3D maps from two-half tilt-angles (based on their odd and even tilt angles) was generated. The frequencies at which the FSC curve begin falls below the values of 0.5 and 0.143 were used to represent the resolution.

iii) Map-to-model based FSC analysis: the FSC curve between the final IPET reconstruction and its fitting model-generated density map was computed, and the frequencies at which the FSC curve constantly fell below 0.5 were used as the estimated resolution. The density map of the fitting model was generated by the *molmap* command in UCSF Chimera^57^.

iv) IPET-to-SPA-based FSC analysis: The FSC between each of two selected particles and McRae’s 4.9 Å cryo-EM SPA map^11^ was computed, and the frequency at which the FSC curve constantly fell below 0.5 was used as the estimated resolution.

### Structural modeling of each individual 6HBC particle

A PDB model (6HBC-mature^11^) was used as a template model. The four helical segments of each helix from the model were rigid-body dockde manually into each helix of the 120 IPET reconstructed 3D maps using Chimera^57^. The loop structure between segments were then repaired using Phenix software^58^.

### Statistical analysis

Each helix of 6HBC-mature was divided into four short-helices based on the KL and crossover. The orientations of a total of 24 short-helices in each particle were defined by 24 vectors, which were measured using DSSR software^59^. These vectors were then used for statistical analysis of the particle structural variety. For tilting angle measurement, the tilting angle of each helix was defined by averaging its two central vectors. Crossover angles were measured by two averaged vectors, each representing half of the helix containing two short-helices. The helix curvature angle was measured by the angles between two averaged vectors within each half of the helix. The bending angle was measured by the two conjunctive vectors at the KL. The coupling angle was measured by two vectors within each half of the helix. The corresponding histograms were plotted and fitted with kernel density estimation in Python.

### Maturation percentage (MP) definition

The MP was determined by averaging the contribution of these three parameters (the distance between H1-H5, the H6 tilting angle α and the H1-H5 bundle curvature angle φ. This was expressed as a percentage, *i.e*.

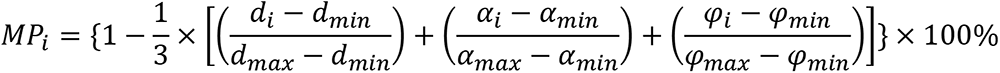

### RNA templates

The 6HBC RNA origami was designed using 720 bases^11^, as shown in **Extended Data Fig. 1a**. The corresponding secondary structure, including the high probability base pairs and single-stranded regions, was predicted using RNA structure software^60^ (**Extended Data Fig. 1b**). The blueprint of the design is presented in **Extended Data Fig. 1c.**

### Sample preparation for single particle analysis cryo-EM

A Leica GP2 was prepared with the sample application chamber set to 100% humidity and 21°C. The Leica GP2 was set with a manually calibrated blotting distance onto two Whatman no. 1 filter paper layers with 4 seconds of delay after sample application, 6 seconds of blot time followed directly by plunging into liquid ethane at -184°C. ProtoChips Au-FLAT 1.2/1.3 300 mesh grids were glow discharged for 45 seconds at 15mA in a PELCA easiGlow immediately before sample application.

### Single particle analysis cryo-EM data collection

Data was acquired on a Titan Krios G1 equipped with Cs corrector and a K2 camera (Gatan/Ametek). Data were collected with targeted defocus range of −0.5 to −2.0 μm and a targeted dose of 60 e^−^Å^−2^ and a pixel size of 0.860 Å px–1. Automated data collection was performed with EPU and data saved as MRC files.

### Single particle analysis data processing

All SPA analysis was performed in cryoSPARC, with the pre-processing occurring in cryoSPARC V2 and the final refinements and 3D variability analysis in cryoSPARC V4 ^61^. Patch motion correction was performed with Fourier cropping by ½ and otherwise default settings. Patch CTF was used with default parameters for CTF correction. Micrographs were curated to remove bad ice, and micrographs with CTF fits worse than 10 Å, leaving 2393 curated exposures.

Particles were picked with a pre-trained Topaz ^62^ model (ResNet16 (64 units), with a downsampling factor of 4. Particles were extracted with a box size of 256 pixels (430 Å) and down sampled to 128 pixels, resulting in 264,890 particles. 2D classification was performed with 100 classes and a circular mask with a diameter of 200 Angstrom. 25 Junk classes were excluded from further analysis, leaving 201,745 particles in the working stack. 4-class ab initio reconstruction was performed using 40k random particles, followed by heterogeneous refinement using all 4 volumes and all 201,745 particles. This resulted in two distinct classes each resembling a 6-helix bundle, with 69,577 and 69,948 particles, respectively.

These particles were re-extracted (256px box) and re-centered based on aligned shifts. Further refinement of these classes revealed some duplicate picks, which were then removed from the stack, leaving 113,076 particles.

To explore the possibility of further sub-populations of alternate conformers we performed 3D variability analysis ^5^ using a mask that was low-pass filtered to 50 Å, dilated 5 voxels with a threshold set to 0.5 and soft-padded with a width of 15 voxels. 3DVA was performed solving for 3 modes of variability, 1 of which appears to resemble real differences in the data, the other 2 appear to contain noise. We then isolated particles along the reaction coordinate that showed the transition from conformer 1 to conformer 2 and made independent reconstructions from these intermediary particle stacks.

To further explore the possibility of additional sub-populations of conformers we performed 3D classification without alignment into 2, 5 and 10 classes using standard parameters.

## Extended Data

**Extended Data Fig. 1:**
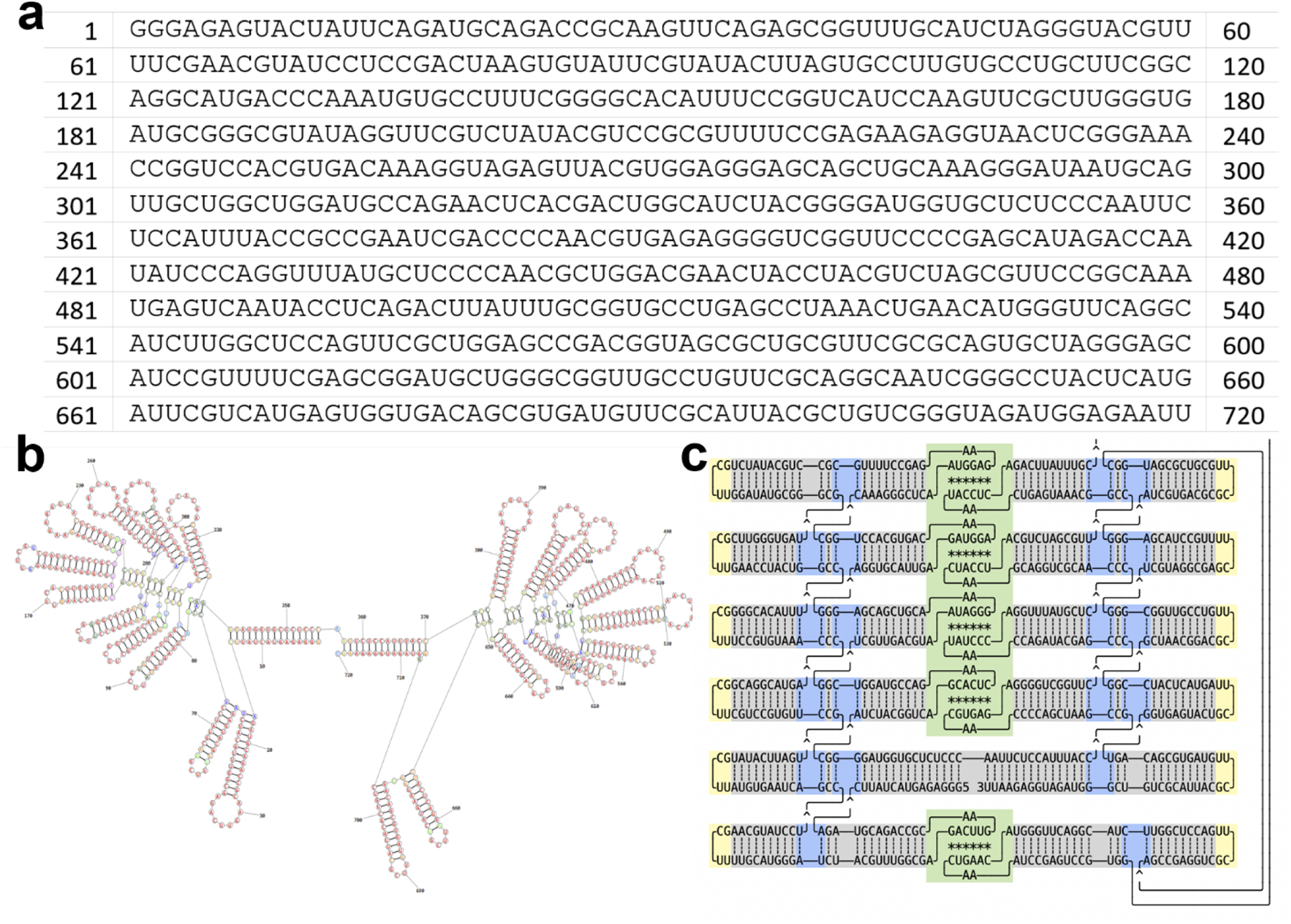
Schematic of 6HBC. **a,** Primary and **b** and **c,** secondary structure of 6HBC. A diagram of the related tertiary structure is also shown, featuring the A-form helix (grey), kissing-loop (KL) interactions (green), coaxial stacking at junctions (blue) and tetra loops (yellow).

**Extended Data Fig. 2:**
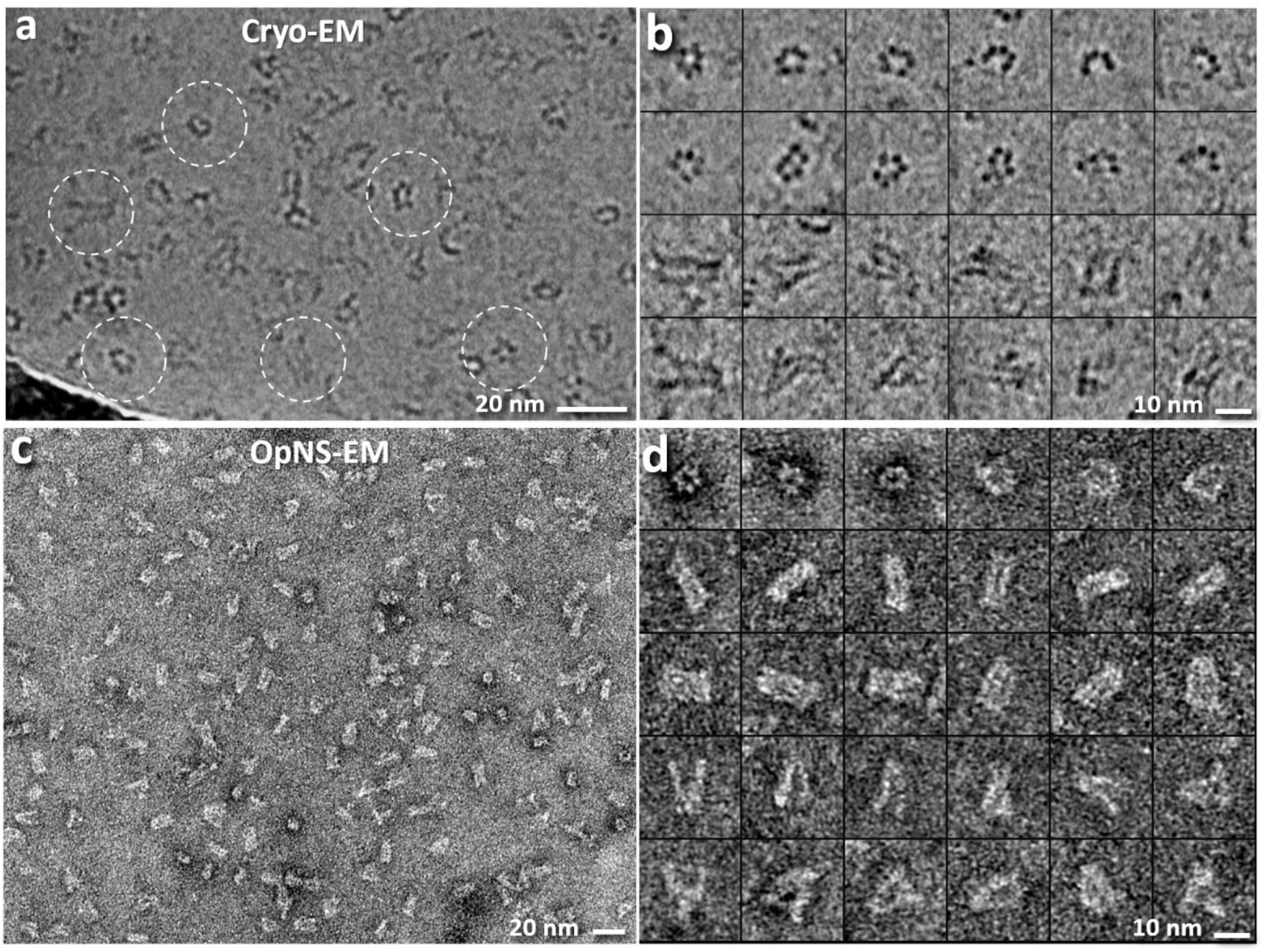
Cryo-EM and NS-EM images of RNA origami 6HBC. **a,** A survey cryo-EM EM image. **b,** 24 representative particles, **c,** A survey NS EM image. **d,** 30 representative particles.

**Extended Data Fig. 3:**
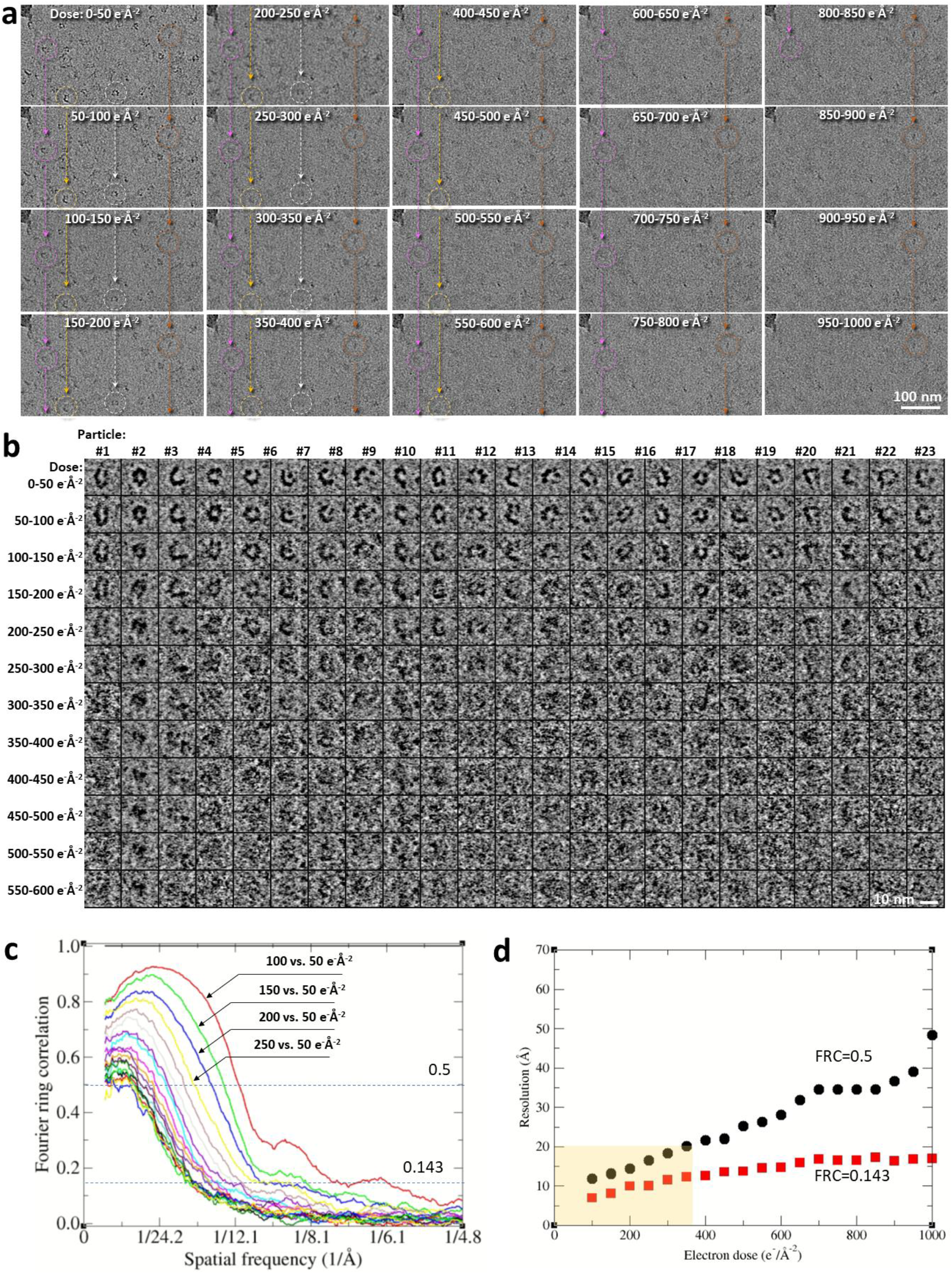
The effect of electron dose on the cryo-EM images of 6HBC RNA particles. **a,** A sample area is sequentially imaged at a dose of 50 e^-^Å^-2^ for 20 times. Thus, the images correspond to the radiation damage with an accumulated dose of 50 e^-^Å^-2^, 100 e^-^Å^-2^, 150 e^-^Å^-2^, up to 1,000 e^-^Å^-2^, respectively. Representative particles were tracked and marked by circles. **b,** Zoomed-in images of 23 representative areas that contain particles. **c,** The FSC curves affected by the radiation damage, with each curve calculated by comparing the first image (50 e^-^Å^-2^) against the rest of images. **d,** The plot of the FSC frequencies at FSC=0.5 and 0.143, respectively, against their corresponding damaged doses.

**Extended Data Fig. 4:**
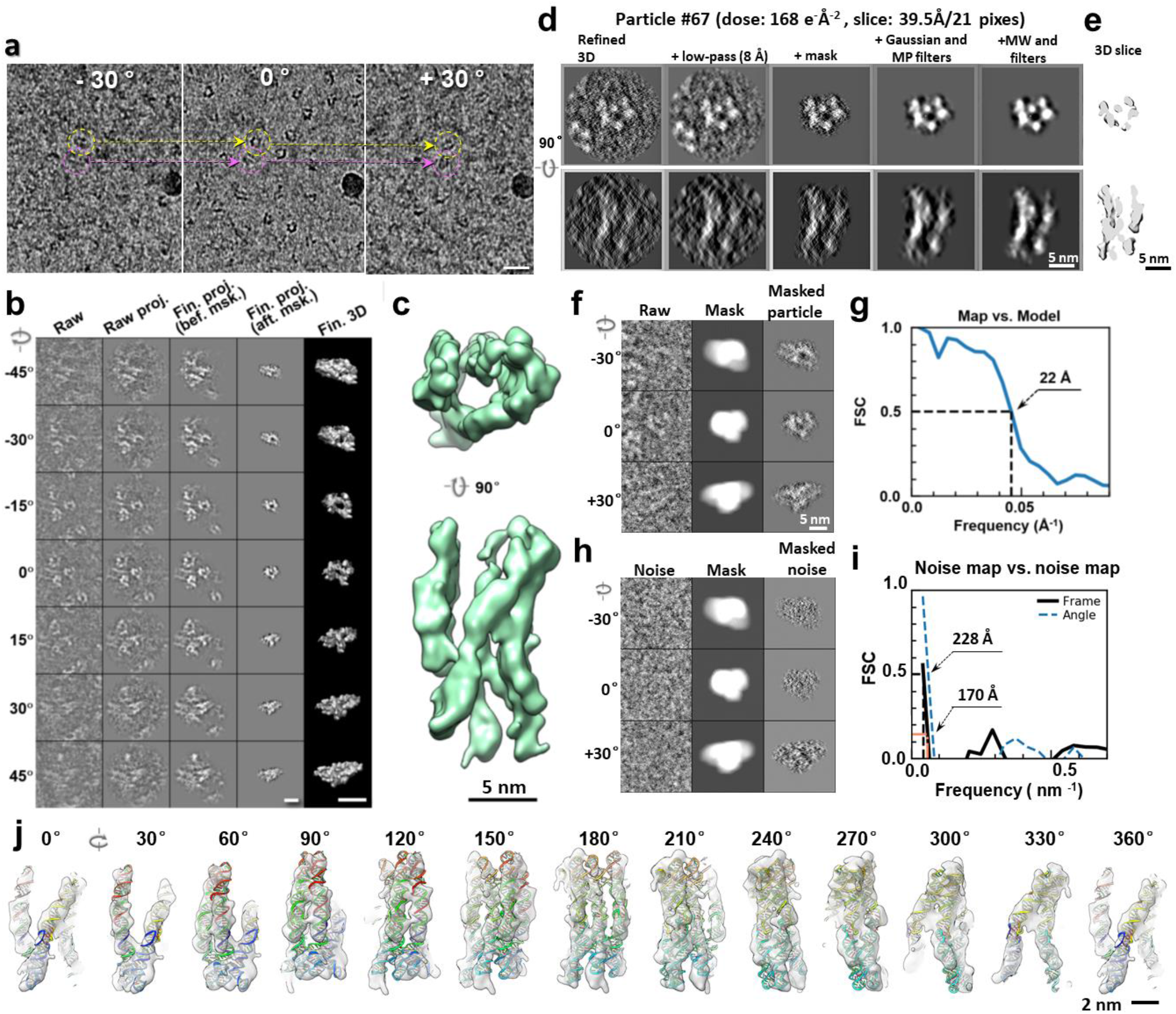
IPET 3D reconstruction and fitting model of an individual particle achieved at a dose of 168 e^-^Å^-2^. **a,** Cryo-ET images at three representative tilt angles, with two exemplary particles marked in the tilt images. **b,** The process of 3D reconstruction of a targeted particle using IPET. Seven representative tilt images of the particle are progressively aligned through iterative refinement for 3D reconstruction. **c,** The final 3D reconstruction of the targeted particle at 25-Å resolution, as measured by the frame-based map-map FSC analysis at FSC=0.5. **d,** Two perpendicular views of the central slices (39.5-Å thickness) of the 3D reconstructions before and after masking and denoising. **e,** 3D views of the central cross-sections of the final reconstruction, displayed from two perpendicular directions. **f,** Three representative tilt raw-images of the particle, the images of the automatically generated soft-boundary masks, and the particle images after mask application. **j,** FSC curve calculated between the IPET reconstruction and the density map generated from the fitting model. **h,** A control test was performed by applying the same masks on a noise area in the sample, which contains no particles. **i,** The FSC analysis of the two-half reconstructions generated from the masked noise. **j,** The detailed fitting model shown in a series of rotated views in 30° increments along the vertical axis.

**Extended Data Fig. 5:**
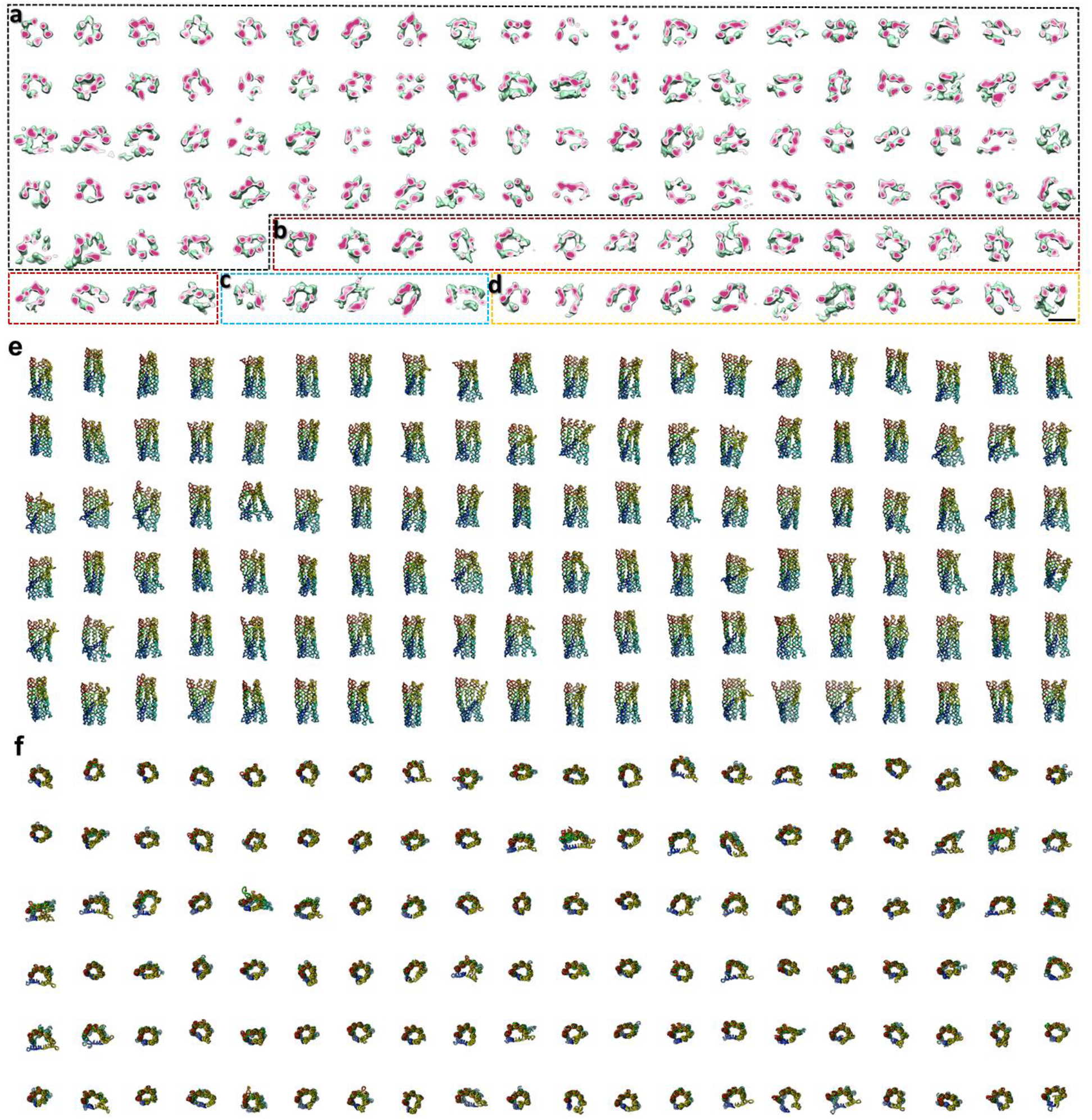
Gallery of 120 individual-particle 3D reconstructions and corresponding fitting models of RNA origami 6HBC. **a-d**, The central cross-sections of the 120 IPET 3D reconstructions, corresponding to those shown in Fig. 2a. **e,f**, The corresponding fitting models from two perpendicular views. The order of the sequences is color-coded by a rainbow.

**Extended Data Fig. 6:**
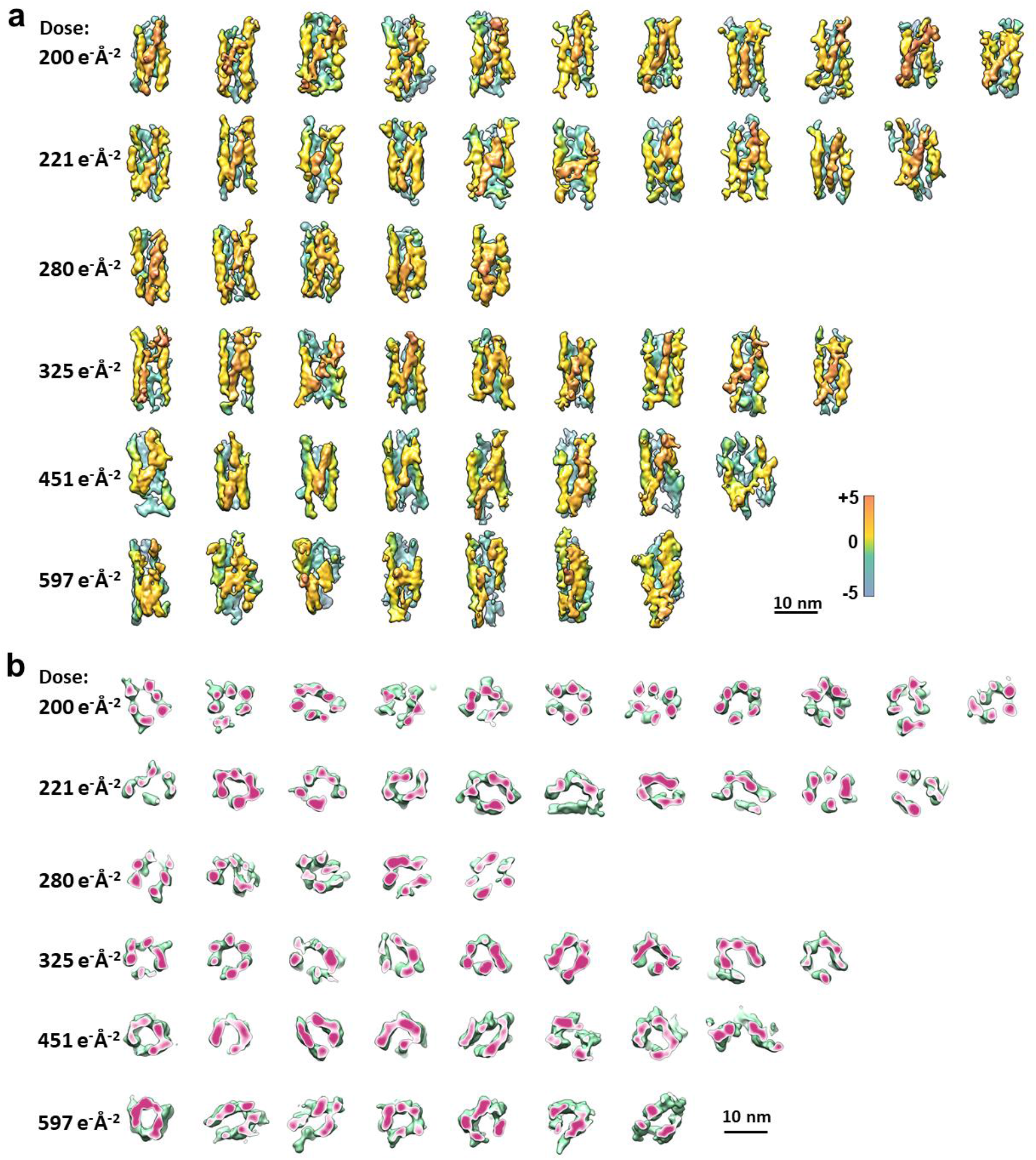
Gallery of fifty-two individual-particle 3D reconstructions of RNA origami 6HBC achieved at a high dose in the range of 200 – 597 e^-^Å^-2^. **a,** Eleven reconstructions achieved at a dose of 200 e^−^Å^-2^, ten reconstructions at a dose of 221 e^−^Å^-2^, five reconstructions at a dose of 280 e^−^Å^-2^, nine reconstructions at a dose of 325 e^−^Å^-2^, eight reconstructions at a dose of 451 e^−^Å^-2^ and seven reconstructions at a dose of 597 e^−^Å^-2^. b, Central cross-sections of the above fifty-two IEPT 3D reconstructions.

**Extended Data Fig. 7:**
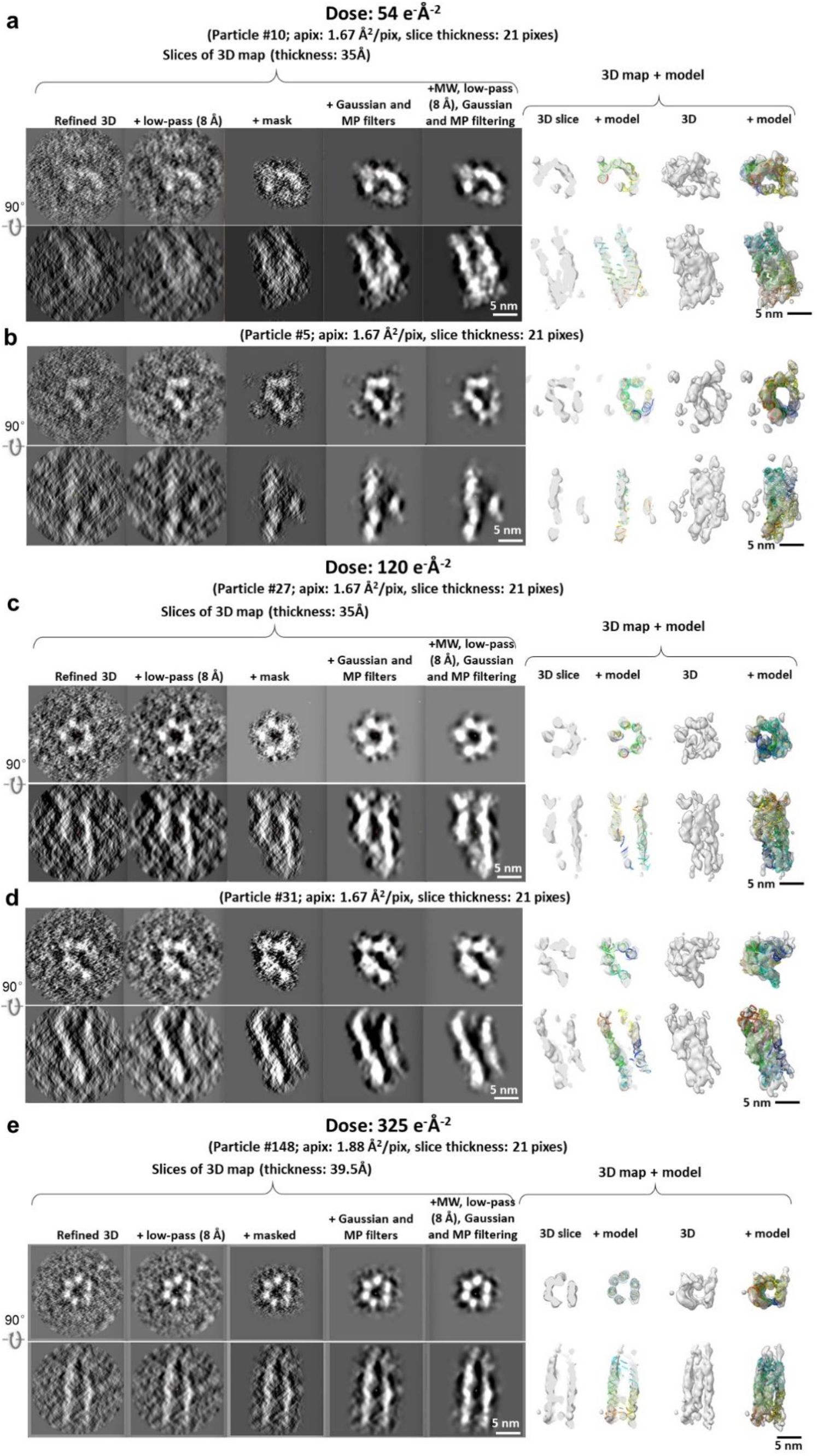
Intermediate procedure of 3D reconstruction of representative individual particle of 6HBC. **a,b,** Comparison of the central slices of 3D reconstructions before and after applying masks and filters of two representative individual particle achieved at a dose of 54 e^-^Å^-2^ (left 5 panels), viewing from two perpendicular directions. The right four panels show the cross-section of the final 3D map and its fitting model. **c,d,** The procedures of two representative individual particles achieved at a dose of 120 e^-^ Å^-2^. **e,** The procedures of a representative individual particle achieved at a dose of 325 e^-^Å^-2^.

**Extended Data Fig. 8:**
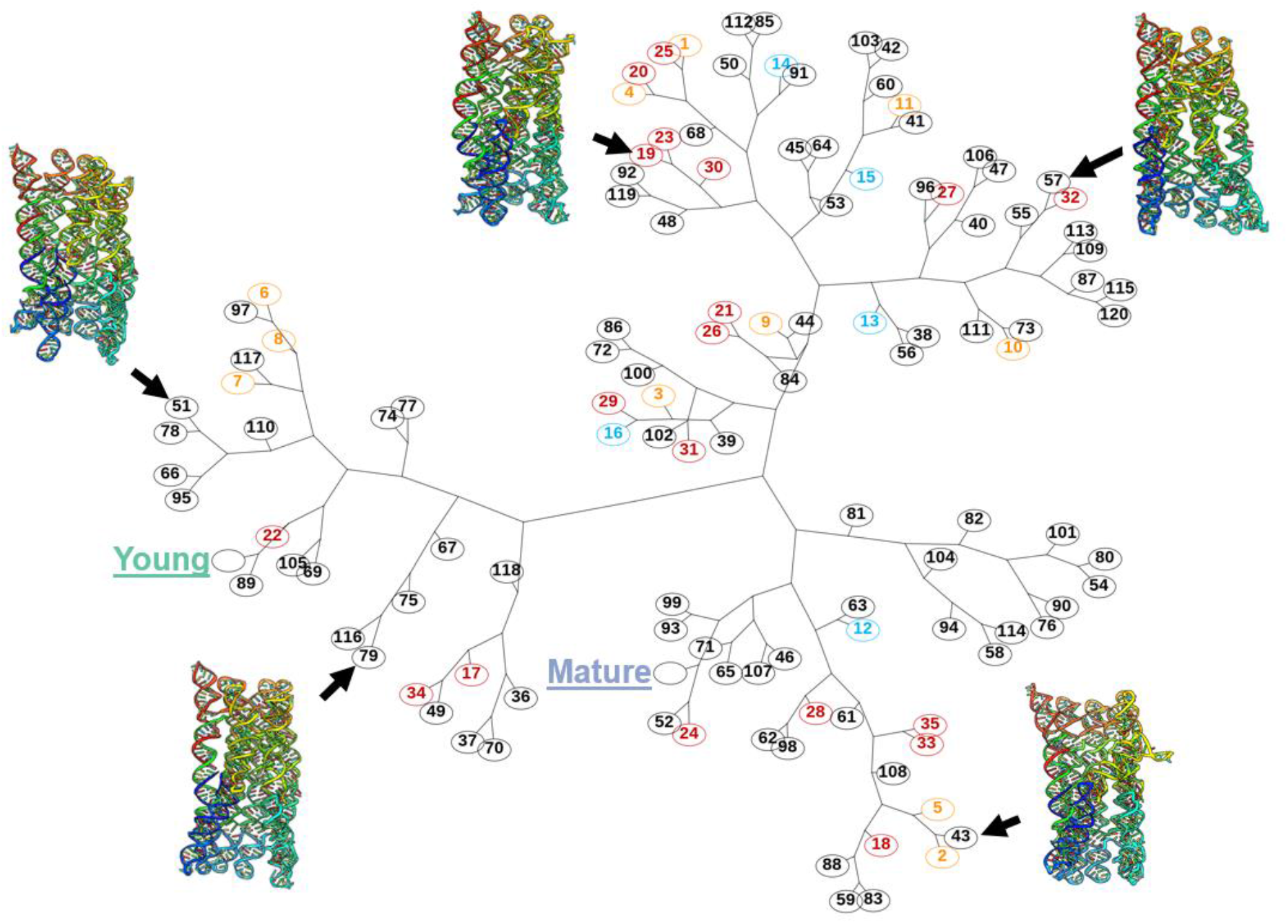
2D tree diagram of the Hierarchical clustering analysis of RMSD values. The RMSD is calculated between each pair of structures, in which the two SPA structures, *i.e.* the “young” and “mature” conformations^11^, are included for comparison. The structures obtained at a dose of 168 e^−^Å^-2^ are marked in black circles, 120 e^−^Å^-2^ in red, 107 e^−^Å^-2^ in cyan and 54 e^−^Å^-2^ in orange. The tree map presents the distances by RMSD values, and the structures of five representative particle are showed beside their corresponding indexes.

**Extended Data Fig. 9:**
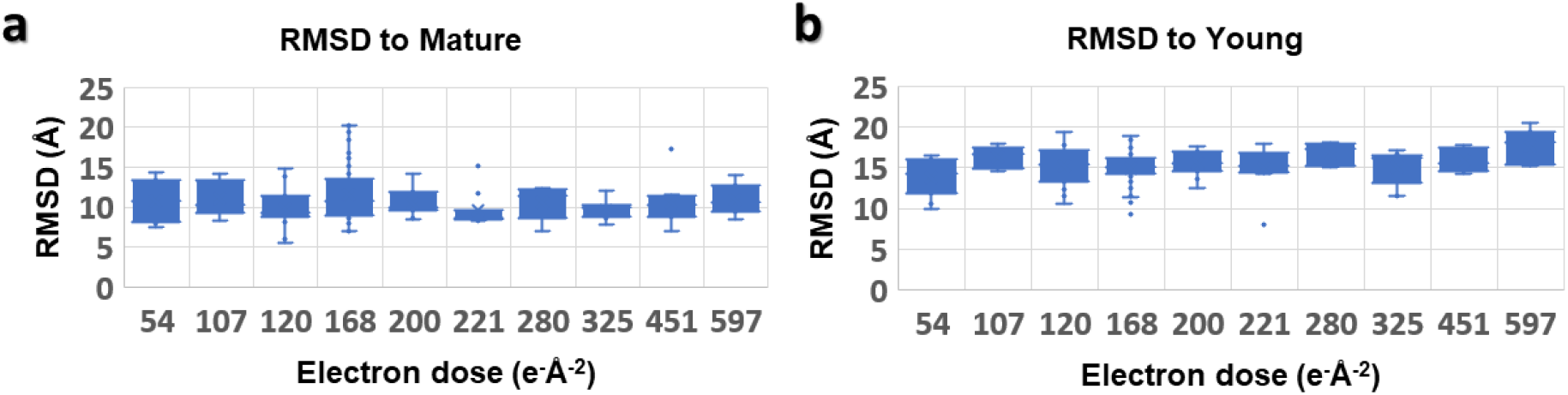
The box plot of the mean of RMSDs against the corresponding dose. **a,** The mean of RMSDs against the “mature” structure distributed based on the achieved electron dose. **b,** The plot of the RMSD against the “young” structure. The data for the plots is shown in the **Supplementary data table 1.**

