## Supplemental information table and figures for "Tertiary structure of single-instant RNA molecule reveals folding landscape"

**Extended methods:**

**Resolution assessment of the 3D reconstructions.** The resolution of the 3D reconstructions was assessed using following four recommended metrics[63]: i) Fourier shell correlation (FSC) between two halves of 3D maps from even-to-odd frames, with frequencies at FSC=0.5[64,65] and 0.143[18]. As exemplary particle shown in Fig. 1, the measured resolutions are 25 Å and 20 Å, respectively (solid line showed in **Fig. 1g**); ii) FSC from even-to-odd tilt images, which showed a nearly identical FSC curve and resolutions to the previous metric (dash line showed in **Fig. 1g**). Both assessments are independent of the applied soft-masks (**Extended Data Fig. 3i**); iii) FSC curve between the map and model[63], where the density map generated from the fitted model showed a frequency of 22 Å at FSC=0.5 for the particle showed in Fig. 1 (**Extended Data Fig. 3g**); iv) Visible structure features, such as the ~20 Å RNA helix and major grooves. To make a comparison, a density map generated from the PDB structure of the "mature" conformation[11] and its cryo-EM SPA density map were low-pass filtered to resolutions of 15 Å, 20 Å, 25 Å, and 30 Å, respectively (**Supplementary Fig. 3**). The views of the filtered maps showed that both the individual helix and its surface large groove became difficult to discern at resolutions of 25 Å and above. In comparison, four IPET maps (particle #67, #60, #40, and #82) with similar "mature" conformation showed, although not all large grooves were visible, nearly all helices were visible in the IPET maps (**Supplementary Fig. 3h**). Cross-sectional views showed a similar contrast of helices to that of the low-passed filtered SPA model/map at 20 and 25 Å (**Supplementary Fig. 3j**), suggesting a resolution of 20-25 Å. The consistency of the resolutions assessed by multiple metrics confirmed that the achieved resolution is suitable for tertiary structure determination.

**Validation of the 3D reconstructions.**

To assess whether the achieved tertiary structures are affected by radiation damage from the electron dose (*i.e.*, 168 $e^{-}Å^{-2}$) or the image processing protocols, we conducted four validation analyses:

i) Validated by dose-filtering method[29]: To test whether the 3D reconstruction is related to the acquired dose, we removed the last four images of the tilt series, acquired at the highest-tilt angles (+50° +45°, -45°, and -50° in our case), before 3D reconstruction. This is equivalent to reducing the total electron dose for the reconstruction to 136 $e^{-}Å^{-2}$ (the similar dose has been used to determine an atomic resolution structure by cryo-ET sub-volume average method [16]). As an example, using particle #072, the central slices of the dose-reduced final 3D reconstruction showed near-identical features to the original one when viewed from two perpendicular directions (**Supplementary Fig. 2a**). Superimposing the two 3D maps showed a near-perfect match to their rod-shaped helix densities, except for the surrounding noise and distal end noise (**Supplementary Fig. 2b**). The FSC curves confirmed their similarity (**Supplementary Fig. 2c-e**). All comparisons suggest that the reconstruction did not significantly improve by removing the high-dose portion of the data (*i.e.*, dose-filtered to 136 $e^{-}Å^{-2}$), suggesting that the high-dose portion of the data did not significantly affect the final 3D reconstruction, confirming that the final 3D map is not related to the high-dose portion of the data. This result is consistent with previous studies, in which the resolution improved slightly (from 2.9 Å to 2.6 Å) at a dose of 100 $e^{-}Å^{-2}$ by dose-filtering[29].

ii) Validated by SPA reconstructions: We have studied the same 6HBC sample using conventional cryo-EM SPA method[11] at a dose of 60 $e^{-}Å^{-2}$, which has resulted in the determination of "young" and "mature" conformations at 3.4 - 4.9 Å resolution. By comparing the IPET structures to SPA structures, we have observed that some particles exhibit a conformation similar to the "mature" structure (**Fig. 3e-g**), while others resemble the "young" structure (**Fig. 3h-j**). For example, when comparing particle #072 to the "mature" structure (**Supplementary Fig. 4a,b**), we found that the frequencies at FSC=0.5 and 0.143 are 27 Å and 21 Å, respectively (**Supplementary Fig. 4c**). By rigid-body docking the "mature" structure directly into this particle #072 map, we observed that the superimposed images match the density map and fitted model very well, with only a H5 shift of ~1 nm (about helix half-width) toward the inside of the molecule and one distal end of H6 tilted toward the outside of the map (indicated by orange arrows in **Supplementary Fig. 4d**). The root-mean-square deviation (RMSD) between the two structures is 7.2 Å (**Supplementary Fig. 4e**). Considering that the two structures were obtained using different software (cryoSPARC vs. IPET), different numbers of particles (thousands vs. one), and different approaches for reconstruction (un-tilt vs. tilt series), and were imaged at different electron doses (60 vs. 168 $e^{-}Å^{-2}$), using different cryo-EM grids and cryo-EM instruments (Titan Krios G2 vs. G1), the similarity between the structures suggests that the IPET structure is unlikely related to the artifact from the radiation damage or reconstruction method.

iii) Low-dose validation: To demonstrate that the determined structures are not artifacts resulting from radiation damage caused by the dose of 168 $e^{-}Å^{-2}$, we have repeated the cryo-ET experiment using three lower doses and have obtained an additional 35 density maps, including 19 maps with average resolution ranging from 22.7 to 28.5 Å, achieved from an electron dose of 120 $e^{-}Å^{-2}$, 5 maps with average resolution of 23.5 – 32.3 Å, achieved from 107 $e^{-}Å^{-2}$, and 11 maps with average resolution of 23.6 - 29.1 Å, achieved from 54 $e^{-}Å^{-2}$ (**Fig. 2b-d, Extended Data Fig. 5b-d**). All 3D reconstructions and central cross-sections clearly show the helical structures (**Extended Data Fig. 7a-d**). Zoomed-in images of two representative reconstructions acquired at the doses of 120 $e^{-}Å^{-2}$ and 54 $e^{-}Å^{-2}$, respectively, indicate that the helical structure is not dependent on the use of soft-masks and filters (**Extended Data Fig. 7a-d**). Additionally, these low-dose structures also contain both "young" and "mature" conformations, as seen in the "high-dose" structure (**Extended Data Fig. 7e**), further supporting the idea that the dose is not related to the observation of structural variability in this study.

iv) Reconstruction using IMOD software: In order to determine whether the tertiary structure achieved is an artifact of our IPET software[21], we reprocessed the 3D reconstruction using a third-party software, IMOD[33] on the cryo-ET tilt series acquired at a doses of 168 $e^{-}Å^{-2}$, 120 $e^{-}Å^{-2}$, and 54 $e^{-}Å^{-2}$, respectively. The central slices of the large micrograph reconstructions and zoomed-in particles revealed that the helical structures were still visible in each individual particle, although the reconstructions at 54 $e^{-}Å^{-2}$ were noisier than the others (**Supplementary Fig. 5**). This test confirmed that the IPET software is not a key parameter for achieving the helical structure but rather improves the signal-to-noise ratio by reducing the missing-wedge effects.

v) RMSD analysis: To determine whether the tertiary structure obtained by IPET is related to radiation damage, we investigate the RMSD distribution against the achieved dose from following three perspectives. First, the RMSD obtained at different doses was displayed in a Hierarchical clustering dendrogram using various colors (**Fig. 4b**). Second, the RMSD obtained at different doses was displayed in a 2D tree diagram of the Hierarchical clustering circle marked by different colors (**Extended Data Fig. 8**). Lastly, the RMSD of each IPET model was compared against each of SPA[11] structures (i.e. the "mature" and "young" conformations achieved at a dose of 60 $e^{-}Å^{-2}$). A box plot of the RMSD against the achieved dose was then displayed (**Extended Data Fig. 9**). All three analyses showed poor correlation between the RMSD and the dose, suggesting that the IPET structure is less likely to be related to radiation damage.

**Supplementary data table 1: Detailed information on IPET 3D reconstructions of 170 particles.** The MPs for the "young" and "mature" structures are approximately 47% and 64%, respectively. The maxima and minima of MPs are highlighted in bold.

[a] Method 1: The resolution is defined based on the map-map FSC analysis, where two-half maps are generated from the even and odd frames at each tilt angle, respectively, and the frequency at FSC=0.5 is defined as the resolution.

[b] Method 2: Same as Method 1, except the frequency at FSC = 0.143 is defined as the resolution.

[c] Method 3: Same as Method 1, except two-half maps are generated from the even and odd tilt images, respectively, and the frequency at FSC=0.5 is defined as the resolution.

[d] Method 4: Same as Method 3, except the frequency at FSC=0.143 is defined as the resolution.

[e] Method 5: The resolution is defined based on the map-model FSC analysis, where the model map is generated from the fitted model, and the frequency at FSC=0.5 is defined as the resolution.

| Figure | Part. ID | Total dose $(e^-/Å^2)$ | TEM | Magnification | Angle range | Tilt step | Exposure time (s) | Resolution (Å) Meth. 1[a] | Meth. 2[b] | Meth. 3[c] | Meth. 4[d] | Meth. 5[e] | Contour level | MP (%) | EMDB ID |
|---|---|---|---|---|---|---|---|---|---|---|---|---|---|---|---|
| S7 | 036 | 168 | G2 | 81K | -50° to +50° | 5° | 0.88 | 26 | 21 | 26 | 24 | 23 | 1.080 | 62.3 | 25273 |
| S8 | 037 | 168 | G2 | 81K | -50° to +50° | 5° | 0.88 | 26 | 22 | 26 | 22 | 23 | 1.000 | 75.7 | 25274 |
| S9 | 038 | 168 | G2 | 81K | -50° to +50° | 5° | 0.88 | 26 | 23 | 26 | 22 | 24 | 1.000 | 85.0 | 25275 |
| S10 | 039 | 168 | G2 | 81K | -50° to +50° | 5° | 0.88 | 25 | 17 | 26 | 22 | 24 | 1.000 | 61.0 | 25276 |
| S11 | 040 | 168 | G2 | 81K | -50° to +50° | 5° | 0.88 | 25 | 21 | 27 | 22 | 22 | 1.000 | 72.8 | 25277 |
| S12 | 041 | 168 | G2 | 81K | -50° to +50° | 5° | 0.88 | 26 | 23 | 28 | 24 | 23 | 1.040 | 67.4 | 25278 |
| S13 | 042 | 168 | G2 | 81K | -50° to +50° | 5° | 0.88 | 27 | 23 | 27 | 24 | 23 | 1.000 | 80.9 | 25356 |
| S14 | 043 | 168 | G2 | 81K | -50° to +50° | 5° | 0.88 | 26 | 22 | 26 | 22 | 24 | 1.000 | 64.8 | 25279 |
| S15 | 044 | 168 | G2 | 81K | -50° to +50° | 5° | 0.88 | 28 | 23 | 28 | 25 | 26 | 1.000 | 74.3 | 25280 |
| S16 | 045 | 168 | G2 | 81K | -50° to +50° | 5° | 0.88 | 26 | 23 | 27 | 22 | 22 | 1.000 | 76.5 | 25281 |
| S17 | 046 | 168 | G2 | 81K | -50° to +50° | 5° | 0.88 | 26 | 22 | 28 | 24 | 23 | 1.100 | 60.6 | 25357 |
| S18 | 047 | 168 | G2 | 81K | -50° to +50° | 5° | 0.88 | 26 | 23 | 27 | 24 | 24 | 1.100 | 75.7 | 25282 |
| S19 | 048 | 168 | G2 | 81K | -50° to +50° | 5° | 0.88 | 28 | 23 | 30 | 23 | 24 | 0.900 | 41.8 | 25283 |
| S20 | 049 | 168 | G2 | 81K | -50° to +50° | 5° | 0.88 | 26 | 23 | 26 | 23 | 23 | 1.000 | 66.2 | 25284 |
| S21 | 050 | 168 | G2 | 81K | -50° to +50° | 5° | 0.88 | 27 | 21 | 29 | 24 | 23 | 1.000 | 39.5 | 25285 |
| S22 | 051 | 168 | G2 | 81K | -50° to +50° | 5° | 0.88 | 26 | 23 | 27 | 23 | 22 | 1.100 | 68.8 | 25286 |
| S23 | 052 | 168 | G2 | 81K | -50° to +50° | 5° | 0.88 | 28 | 23 | 28 | 24 | 24 | 1.000 | 67.3 | 25287 |
| S24 | 053 | 168 | G2 | 81K | -50° to +50° | 5° | 0.88 | 26 | 23 | 27 | 23 | 24 | 1.000 | 63.1 | 25288 |
| S25 | 054 | 168 | G2 | 81K | -50° to +50° | 5° | 0.88 | 26 | 22 | 27 | 24 | 24 | 1.000 | 55.5 | 25289 |

| | | | | | | | | | | | | | | | |
|---|---|---|---|---|---|---|---|---|---|---|---|---|---|---|---|
| S26 | 055 | 168 | G2 | 81K | -50° to +50° | 5° | 0.88 | 27 | 22 | 26 | 23 | 22 | 1.000 | 70.3 | 25290 |
| S27 | 056 | 168 | G2 | 81K | -50° to +50° | 5° | 0.88 | 26 | 23 | 28 | 23 | 24 | 1.000 | 69.6 | 25291 |
| S28 | 057 | 168 | G2 | 81K | -50° to +50° | 5° | 0.88 | 27 | 22 | 26 | 23 | 23 | 1.000 | 68.7 | 25292 |
| S29 | 058 | 168 | G2 | 81K | -50° to +50° | 5° | 0.88 | 25 | 22 | 27 | 24 | 23 | 1.000 | 67.4 | 25293 |
| S30 | 059 | 168 | G2 | 81K | -50° to +50° | 5° | 0.88 | 27 | 23 | 28 | 24 | 23 | 1.000 | 63.0 | 25294 |
| S31 | 060 | 168 | G2 | 81K | -50° to +50° | 5° | 0.88 | 28 | 23 | 27 | 23 | 24 | 1.110 | 77.3 | 25295 |
| S32 | 061 | 168 | G2 | 81K | -50° to +50° | 5° | 0.88 | 26 | 17 | 26 | 23 | 24 | 1.040 | 76.1 | 25296 |
| S33 | 062 | 168 | G2 | 81K | -50° to +50° | 5° | 0.88 | 26 | 22 | 26 | 24 | 23 | 1.000 | 71.8 | 25297 |
| S34 | 063 | 168 | G2 | 81K | -50° to +50° | 5° | 0.88 | 25 | 21 | 25 | 23 | 23 | 1.000 | 72.0 | 25298 |
| S35 | 064 | 168 | G2 | 81K | -50° to +50° | 5° | 0.88 | 26 | 16 | 26 | 22 | 23 | 1.000 | 59.7 | 25299 |
| S36 | 065 | 168 | G2 | 81K | -50° to +50° | 5° | 0.88 | 26 | 18 | 28 | 21 | 24 | 0.900 | 36.8 | 25300 |
| S37 | 066 | 168 | G2 | 81K | -50° to +50° | 5° | 0.88 | 26 | 21 | 29 | 25 | 28 | 1.000 | 16.7 | 25301 |
| S6 | 067 | 168 | G2 | 81K | -50° to +50° | 5° | 0.88 | 25 | 20 | 25 | 21 | 22 | 1.000 | 64.0 | 25302 |
| S38 | 068 | 168 | G2 | 81K | -50° to +50° | 5° | 0.88 | 26 | 22 | 25 | 23 | 25 | 1.000 | 43.1 | 25303 |
| S39 | 069 | 168 | G2 | 81K | -50° to +50° | 5° | 0.88 | 25 | 17 | 26 | 23 | 24 | 0.926 | 46.8 | 25304 |
| S40 | 070 | 168 | G2 | 81K | -50° to +50° | 5° | 0.88 | 26 | 22 | 24 | 22 | 23 | 1.000 | 60.0 | 25305 |
| S41 | 071 | 168 | G2 | 81K | -50° to +50° | 5° | 0.88 | 25 | 20 | 26 | 22 | 24 | 1.000 | 90.3 | 25306 |
| S42 | 072 | 168 | G2 | 81K | -50° to +50° | 5° | 0.88 | 26 | 23 | 26 | 22 | 23 | 1.000 | 66.6 | 25307 |
| S43 | 073 | 168 | G2 | 81K | -50° to +50° | 5° | 0.88 | 28 | 22 | 27 | 24 | 28 | 0.985 | 26.6 | 25308 |
| S44 | 074 | 168 | G2 | 81K | -50° to +50° | 5° | 0.88 | 27 | 21 | 28 | 20 | 23 | 0.900 | 29.0 | 25309 |
| S45 | 075 | 168 | G2 | 81K | -50° to +50° | 5° | 0.88 | 25 | 20 | 25 | 21 | 23 | 1.000 | 66.0 | 25310 |
| S46 | 076 | 168 | G2 | 81K | -50° to +50° | 5° | 0.88 | 27 | 23 | 27 | 18 | 24 | 0.722 | 32.8 | 25311 |
| S47 | 077 | 168 | G2 | 81K | -50° to +50° | 5° | 0.88 | 27 | 22 | 27 | 23 | 24 | 0.774 | 28.2 | 25312 |
| S48 | 078 | 168 | G2 | 81K | -50° to +50° | 5° | 0.88 | 29 | 16 | 27 | 23 | 25 | 0.970 | 33.6 | 25313 |
| S49 | 079 | 168 | G2 | 81K | -50° to +50° | 5° | 0.88 | 26 | 22 | 26 | 22 | 23 | 1.000 | 68.8 | 25314 |
| S50 | 080 | 168 | G2 | 81K | -50° to +50° | 5° | 0.88 | 26 | 22 | 26 | 21 | 25 | 0.940 | 43.4 | 25315 |
| S51 | 081 | 168 | G2 | 81K | -50° to +50° | 5° | 0.88 | 27 | 23 | 27 | 21 | 25 | 1.000 | 43.1 | 25316 |
| S52 | 082 | 168 | G2 | 81K | -50° to +50° | 5° | 0.88 | 25 | 23 | 26 | 24 | 22 | 1.100 | 75.2 | 25317 |
| S53 | 083 | 168 | G2 | 81K | -50° to +50° | 5° | 0.88 | 27 | 22 | 27 | 23 | 23 | 1.000 | 52.3 | 25355 |
| S54 | 084 | 168 | G2 | 81K | -50° to +50° | 5° | 0.88 | 25 | 22 | 28 | 22 | 23 | 1.000 | 70.9 | 25318 |
| S55 | 085 | 168 | G2 | 81K | -50° to +50° | 5° | 0.88 | 26 | 23 | 26 | 22 | 22 | 1.100 | 95.0 | 25319 |
| S56 | 086 | 168 | G2 | 81K | -50° to +50° | 5° | 0.88 | 26 | 23 | 27 | 24 | 23 | 1.000 | 63.6 | 25320 |
| S57 | 087 | 168 | G2 | 81K | -50° to +50° | 5° | 0.88 | 24 | 19 | 26 | 22 | 23 | 0.890 | 66.1 | 25321 |
| S58 | 088 | 168 | G2 | 81K | -50° to +50° | 5° | 0.88 | 27 | 21 | 28 | 22 | 23 | 0.863 | 56.2 | 25322 |
| S59 | 089 | 168 | G2 | 81K | -50° to +50° | 5° | 0.88 | 25 | 18 | 25 | 21 | 24 | 1.000 | 51.2 | 25323 |
| S60 | 090 | 168 | G2 | 81K | -50° to +50° | 5° | 0.88 | 25 | 21 | 27 | 24 | 25 | 1.000 | 72.5 | 25354 |
| S61 | 091 | 168 | G2 | 81K | -50° to +50° | 5° | 0.88 | 25 | 22 | 25 | 23 | 23 | 1.000 | 72.7 | 25324 |
| S62 | 092 | 168 | G2 | 81K | -50° to +50° | 5° | 0.88 | 26 | 23 | 26 | 23 | 23 | 1.030 | 66.1 | 25325 |
| S63 | 093 | 168 | G2 | 81K | -50° to +50° | 5° | 0.88 | 25 | 22 | 25 | 21 | 23 | 1.000 | 79.4 | 25326 |
| S64 | 094 | 168 | G2 | 81K | -50° to +50° | 5° | 0.88 | 26 | 23 | 29 | 25 | 24 | 1.000 | 37.4 | 25327 |
| S65 | 095 | 168 | G2 | 81K | -50° to +50° | 5° | 0.88 | 27 | 23 | 26 | 21 | 25 | 1.000 | 50.6 | 25328 |

| | | | | | | | | | | | | | | | |
|---|---|---|---|---|---|---|---|---|---|---|---|---|---|---|---|
| S66 | 096 | 168 | G2 | 81K | -50° to +50° | 5° | 0.88 | 25 | 22 | 26 | 22 | 24 | 1.000 | 53.3 | 25329 |
| S67 | 097 | 168 | G2 | 81K | -50° to +50° | 5° | 0.88 | 26 | 23 | 26 | 23 | 22 | 1.000 | 82.4 | 25330 |
| S68 | 098 | 168 | G2 | 81K | -50° to +50° | 5° | 0.88 | 25 | 16 | 26 | 22 | 23 | 1.000 | 67.3 | 25331 |
| S69 | 099 | 168 | G2 | 81K | -50° to +50° | 5° | 0.88 | 25 | 22 | 26 | 24 | 24 | 1.000 | 81.7 | 25332 |
| S70 | 100 | 168 | G2 | 81K | -50° to +50° | 5° | 0.88 | 27 | 24 | 26 | 23 | 22 | 1.000 | 60.0 | 25333 |
| S71 | 101 | 168 | G2 | 81K | -50° to +50° | 5° | 0.88 | 25 | 21 | 26 | 22 | 23 | 1.000 | 86.6 | 25334 |
| S72 | 102 | 168 | G2 | 81K | -50° to +50° | 5° | 0.88 | 25 | 23 | 25 | 23 | 24 | 1.000 | 73.6 | 25335 |
| S73 | 103 | 168 | G2 | 81K | -50° to +50° | 5° | 0.88 | 27 | 22 | 26 | 22 | 24 | 1.000 | 70.7 | 25336 |
| S74 | 104 | 168 | G2 | 81K | -50° to +50° | 5° | 0.88 | 26 | 18 | 27 | 24 | 24 | 1.000 | 38.3 | 25337 |
| S75 | 105 | 168 | G2 | 81K | -50° to +50° | 5° | 0.88 | 26 | 22 | 26 | 23 | 23 | 1.000 | 66.3 | 25338 |
| S76 | 106 | 168 | G2 | 81K | -50° to +50° | 5° | 0.88 | 25 | 22 | 25 | 23 | 22 | 1.000 | 67.7 | 25339 |
| S77 | 107 | 168 | G2 | 81K | -50° to +50° | 5° | 0.88 | 24 | 22 | 25 | 22 | 23 | 1.000 | 76.3 | 25340 |
| S78 | 108 | 168 | G2 | 81K | -50° to +50° | 5° | 0.88 | 23 | 18 | 24 | 22 | 22 | 1.000 | 73.8 | 25341 |
| S79 | 109 | 168 | G2 | 81K | -50° to +50° | 5° | 0.88 | 24 | 20 | 26 | 22 | 26 | 1.050 | 43.1 | 25342 |
| S80 | 110 | 168 | G2 | 81K | -50° to +50° | 5° | 0.88 | 24 | 21 | 25 | 22 | 23 | 1.000 | 78.8 | 25343 |
| S81 | 111 | 168 | G2 | 81K | -50° to +50° | 5° | 0.88 | 25 | 16 | 26 | 22 | 24 | 1.000 | 66.6 | 25344 |
| S82 | 112 | 168 | G2 | 81K | -50° to +50° | 5° | 0.88 | 25 | 23 | 25 | 23 | 22 | 1.000 | 66.1 | 25345 |
| S83 | 113 | 168 | G2 | 81K | -50° to +50° | 5° | 0.88 | 25 | 22 | 25 | 22 | 23 | 0.900 | 64.0 | 25346 |
| S84 | 114 | 168 | G2 | 81K | -50° to +50° | 5° | 0.88 | 24 | 17 | 24 | 21 | 22 | 1.000 | 67.3 | 25347 |
| S85 | 115 | 168 | G2 | 81K | -50° to +50° | 5° | 0.88 | 26 | 20 | 27 | 22 | 27 | 0.900 | 41.9 | 25348 |
| S86 | 116 | 168 | G2 | 81K | -50° to +50° | 5° | 0.88 | 26 | 22 | 27 | 23 | 25 | 1.000 | 51.4 | 25349 |
| S87 | 117 | 168 | G2 | 81K | -50° to +50° | 5° | 0.88 | 26 | 22 | 27 | 23 | 24 | 0.850 | 45.9 | 25350 |
| S88 | 118 | 168 | G2 | 81K | -50° to +50° | 5° | 0.88 | 26 | 22 | 26 | 23 | 24 | 1.000 | 70.6 | 25351 |
| S89 | 119 | 168 | G2 | 81K | -50° to +50° | 5° | 0.88 | 24 | 17 | 26 | 22 | 23 | 1.000 | 64.5 | 25352 |
| S90 | 120 | 168 | G2 | 81K | -50° to +50° | 5° | 0.88 | 25 | 16 | 26 | 23 | 23 | 0.950 | 72.5 | 25353 |
| S91 | 017 | 120 | G3i | 53K | -49° to +49° | 7° | 2.61 | 29 | 20 | 29 | 25 | 24 | 1.000 | 79.6 | 40371 |
| S92 | 018 | 120 | G3i | 53K | -49° to +49° | 7° | 2.61 | 29 | 23 | 29 | 25 | 25 | 0.930 | 60.5 | 40372 |
| S93 | 019 | 120 | G3i | 53K | -49° to +49° | 7° | 2.61 | 27 | 24 | 32 | 24 | 24 | 0.950 | 70.7 | 40373 |
| S94 | 020 | 120 | G3i | 53K | -49° to +49° | 7° | 2.61 | 30 | 25 | 31 | 26 | 26 | 0.950 | 69.9 | 40374 |
| S95 | 021 | 120 | G3i | 53K | -49° to +49° | 7° | 2.61 | 27 | 21 | 32 | 26 | 24 | 0.935 | 39.4 | 40375 |
| S96 | 022 | 120 | G3i | 53K | -49° to +49° | 7° | 2.61 | 29 | 24 | 29 | 25 | 24 | 0.942 | 68.6 | 40377 |
| S97 | 023 | 120 | G3i | 53K | -49° to +49° | 7° | 2.61 | 27 | 25 | 28 | 24 | 23 | 1.000 | 79.8 | 40378 |
| S98 | 024 | 120 | G3i | 53K | -49° to +49° | 7° | 2.61 | 28 | 24 | 29 | 24 | 27 | 1.000 | 63.0 | 40379 |
| S99 | 025 | 120 | G3i | 53K | -49° to +49° | 7° | 2.61 | 35 | 23 | 42 | 27 | 29 | 0.988 | 62.2 | 40380 |
| S100 | 026 | 120 | G3i | 53K | -49° to +49° | 7° | 2.61 | 27 | 24 | 29 | 25 | 30 | 1.000 | 45.8 | 40381 |
| S101 | 027 | 120 | G3i | 53K | -49° to +49° | 7° | 2.61 | 29 | 14 | 30 | 25 | 24 | 0.920 | 69.9 | 40382 |
| S102 | 028 | 120 | G3i | 53K | -49° to +49° | 7° | 2.61 | 29 | 23 | 31 | 19 | 23 | 0.950 | 72.5 | 40383 |
| S103 | 029 | 120 | G3i | 53K | -49° to +49° | 7° | 2.61 | 33 | 24 | 35 | 25 | 26 | 1.000 | 68.9 | 40384 |
| S104 | 030 | 120 | G3i | 53K | -49° to +49° | 7° | 2.61 | 30 | 24 | 32 | 25 | 27 | 0.900 | 92.7 | 40385 |
| S105 | 031 | 120 | G3i | 53K | -49° to +49° | 7° | 2.61 | 28 | 22 | 31 | 25 | 25 | 1.000 | 67.7 | 40386 |
| S106 | 032 | 120 | G3i | 53K | -49° to +49° | 7° | 2.61 | 27 | 19 | 32 | 27 | 26 | 1.000 | 79.5 | 40387 |

| | | | | | | | | | | | | | | | |
|---|---|---|---|---|---|---|---|---|---|---|---|---|---|---|---|
| S107 | 033 | 120 | G3i | 53K | -49° to +49° | 7° | 2.61 | 31 | 25 | 34 | 24 | 24 | 0.880 | 48.1 | 40388 |
| S108 | 034 | 120 | G3i | 53K | -49° to +49° | 7° | 2.61 | 27 | 25 | 29 | 25 | 24 | 0.935 | 77.3 | 40389 |
| S109 | 035 | 120 | G3i | 53K | -49° to +49° | 7° | 2.61 | 31 | 21 | 35 | 23 | 28 | 0.930 | 43.0 | 40390 |
| S110 | 012 | 107 | G3i | 53K | -50° to +50° | 5° | 1.75 | 32 | 21 | 31 | 25 | 35 | 0.900 | 78.5 | 40366 |
| S111 | 013 | 107 | G3i | 53K | -50° to +50° | 5° | 1.75 | 35 | 25 | 33 | 25 | 24 | 0.861 | 71.6 | 40367 |
| S112 | 014 | 107 | G3i | 53K | -50° to +50° | 5° | 1.75 | 33 | 22 | 35 | 23 | 25 | 0.816 | 76.2 | 40368 |
| S113 | 015 | 107 | G3i | 53K | -50° to +50° | 5° | 1.75 | 30 | 23 | 30 | 25 | 30 | 0.900 | 92.6 | 40369 |
| S114 | 016 | 107 | G3i | 53K | -50° to +50° | 5° | 1.75 | 32 | 27 | 33 | 29 | 29 | 0.910 | 39.6 | 40370 |
| S115 | 001 | 54 | G3i | 53K | -50° to +50° | 5° | 0.88 | 27 | 25 | 29 | 24 | 23 | 1.000 | 74.7 | 40353 |
| S116 | 002 | 54 | G3i | 53K | -50° to +50° | 5° | 0.88 | 32 | 24 | 30 | 24 | 27 | 1.000 | 84.9 | 40356 |
| S117 | 003 | 54 | G3i | 53K | -50° to +50° | 5° | 0.88 | 28 | 24 | 29 | 24 | 24 | 0.900 | 59.4 | 40357 |
| S118 | 004 | 54 | G3i | 53K | -50° to +50° | 5° | 0.88 | 33 | 28 | 29 | 25 | 26 | 0.876 | 76.3 | 40358 |
| S119 | 005 | 54 | G3i | 53K | -50° to +50° | 5° | 0.88 | 29 | 23 | 28 | 20 | 24 | 0.937 | 43.5 | 40359 |
| S120 | 006 | 54 | G3i | 53K | -50° to +50° | 5° | 0.88 | 26 | 23 | 26 | 23 | 28 | 0.942 | 48.3 | 40360 |
| S121 | 007 | 54 | G3i | 53K | -50° to +50° | 5° | 0.88 | 29 | 25 | 31 | 25 | 30 | 1.030 | 38.4 | 40361 |
| S122 | 008 | 54 | G3i | 53K | -50° to +50° | 5° | 0.88 | 29 | 24 | 29 | 24 | 23 | 1.000 | 64.0 | 40362 |
| S123 | 009 | 54 | G3i | 53K | -50° to +50° | 5° | 0.88 | 27 | 18 | 28 | 24 | 25 | 0.927 | 68.6 | 40363 |
| S124 | 010 | 54 | G3i | 53K | -50° to +50° | 5° | 0.88 | 32 | 23 | 34 | 26 | 25 | 0.878 | 69.9 | 40364 |
| S125 | 011 | 54 | G3i | 53K | -50° to +50° | 5° | 0.88 | 28 | 23 | 30 | 25 | 23 | 0.900 | 70.4 | 40365 |
| S126 | 121 | 200 | G2 | 81K | -60° to +60° | 5° | 0.88 | 25 | 20 | 27 | 23 | 22 | 1.010 | - | 25230 |
| S127 | 122 | 200 | G2 | 81K | -60° to +60° | 5° | 0.88 | 25 | 20 | 28 | 22 | 24 | 0.918 | - | 25231 |
| S128 | 123 | 200 | G2 | 81K | -60° to +60° | 5° | 0.88 | 26 | 22 | 26 | 23 | 24 | 0.934 | - | 25232 |
| S129 | 124 | 200 | G2 | 81K | -60° to +60° | 5° | 0.88 | 28 | 23 | 28 | 23 | 24 | 1.030 | - | 25233 |
| S130 | 125 | 200 | G2 | 81K | -60° to +60° | 5° | 0.88 | 26 | 22 | 31 | 22 | 24 | 1.000 | - | 25234 |
| S131 | 126 | 200 | G2 | 81K | -60° to +60° | 5° | 0.88 | 26 | 21 | 27 | 21 | 23 | 1.070 | - | 25235 |
| S132 | 127 | 200 | G2 | 81K | -60° to +60° | 5° | 0.88 | 23 | 21 | 24 | 22 | 22 | 1.120 | - | 25236 |
| S133 | 128 | 200 | G2 | 81K | -60° to +60° | 5° | 0.88 | 25 | 22 | 26 | 23 | 23 | 1.120 | - | 25237 |
| S134 | 129 | 200 | G2 | 81K | -60° to +60° | 5° | 0.88 | 25 | 18 | 26 | 22 | 22 | 1.000 | - | 25238 |
| S135 | 130 | 200 | G2 | 81K | -60° to +60° | 5° | 0.88 | 26 | 18 | 28 | 22 | 24 | 1.000 | - | 25239 |
| S136 | 131 | 200 | G2 | 81K | -60° to +60° | 5° | 0.88 | 24 | 21 | 26 | 22 | 22 | 1.040 | - | 25240 |
| S137 | 132 | 221 | G2 | 81K | -56° to +56° | 7° | 1.43 | 26 | 21 | 36 | 24 | 23 | 1.030 | - | 25250 |
| S138 | 133 | 221 | G2 | 81K | -56° to +56° | 7° | 1.43 | 26 | 23 | 28 | 24 | 23 | 1.000 | - | 25251 |
| S139 | 134 | 221 | G2 | 81K | -56° to +56° | 7° | 1.43 | 25 | 22 | 28 | 23 | 22 | 1.100 | - | 25252 |
| S140 | 135 | 221 | G2 | 81K | -56° to +56° | 7° | 1.43 | 27 | 16 | 29 | 25 | 22 | 1.000 | - | 25253 |
| S141 | 136 | 221 | G2 | 81K | -56° to +56° | 7° | 1.43 | 26 | 22 | 31 | 24 | 25 | 1.000 | - | 25254 |
| S142 | 137 | 221 | G2 | 81K | -56° to +56° | 7° | 1.43 | 28 | 16 | 32 | 25 | 23 | 1.050 | - | 25255 |
| S143 | 138 | 221 | G2 | 81K | -56° to +56° | 7° | 1.43 | 26 | 22 | 28 | 23 | 23 | 1.000 | - | 25256 |
| S144 | 139 | 221 | G2 | 81K | -56° to +56° | 7° | 1.43 | 27 | 23 | 31 | 24 | 23 | 1.000 | - | 25257 |
| S145 | 140 | 221 | G2 | 81K | -56° to +56° | 7° | 1.43 | 26 | 24 | 28 | 24 | 24 | 1.080 | - | 25258 |
| S146 | 141 | 221 | G2 | 81K | -56° to +56° | 7° | 1.43 | 26 | 21 | 30 | 22 | 25 | 1.120 | - | 25259 |
| S147 | 142 | 280 | G2 | 81K | -51° to +51° | 3° | 0.88 | 28 | 22 | 26 | 22 | 23 | 1.000 | - | 25260 |

| | | | | | | | | | | | | | | | |
|---|---|---|---|---|---|---|---|---|---|---|---|---|---|---|---|
| S148 | 143 | 280 | G2 | 81K | -51° to +51° | 3° | 0.88 | 26 | 23 | 28 | 22 | 24 | 1.000 | - | 25261 |
| S149 | 144 | 280 | G2 | 81K | -51° to +51° | 3° | 0.88 | 25 | 22 | 26 | 21 | 24 | 1.000 | - | 25270 |
| S150 | 145 | 280 | G2 | 81K | -51° to +51° | 3° | 0.88 | 26 | 23 | 25 | 22 | 25 | 1.000 | - | 25271 |
| S151 | 146 | 280 | G2 | 81K | -51° to +51° | 3° | 0.88 | 25 | 14 | 25 | 22 | 23 | 1.000 | - | 25272 |
| S152 | 147 | 325 | G2 | 81K | -60° to +60° | 5° | 1.43 | 26 | 24 | 27 | 24 | 24 | 1.000 | - | 25241 |
| S153 | 148 | 325 | G2 | 81K | -60° to +60° | 5° | 1.43 | 26 | 22 | 26 | 23 | 24 | 1.120 | - | 25242 |
| S154 | 149 | 325 | G2 | 81K | -60° to +60° | 5° | 1.43 | 26 | 21 | 28 | 23 | 25 | 1.000 | - | 25243 |
| S155 | 150 | 325 | G2 | 81K | -60° to +60° | 5° | 1.43 | 26 | 22 | 28 | 25 | 25 | 1.000 | - | 25244 |
| S156 | 151 | 325 | G2 | 81K | -60° to +60° | 5° | 1.43 | 26 | 23 | 27 | 24 | 24 | 1.000 | - | 25245 |
| S157 | 152 | 325 | G2 | 81K | -60° to +60° | 5° | 1.43 | 26 | 20 | 28 | 23 | 24 | 1.000 | - | 25246 |
| S158 | 153 | 325 | G2 | 81K | -60° to +60° | 5° | 1.43 | 26 | 23 | 27 | 24 | 24 | 1.000 | - | 25247 |
| S159 | 154 | 325 | G2 | 81K | -60° to +60° | 5° | 1.43 | 26 | 23 | 27 | 24 | 23 | 1.000 | - | 25248 |
| S160 | 155 | 325 | G2 | 81K | -60° to +60° | 5° | 1.43 | 26 | 16 | 27 | 24 | 24 | 1.110 | - | 25249 |
| S161 | 156 | 451 | G3i | 53K | -50° to +50° | 5° | 7.39 | 27 | 16 | 28 | 25 | 26 | 0.900 | - | 40391 |
| S162 | 157 | 451 | G3i | 53K | -50° to +50° | 5° | 7.39 | 26 | 15 | 26 | 23 | 24 | 0.900 | - | 40392 |
| S163 | 158 | 451 | G3i | 53K | -50° to +50° | 5° | 7.39 | 26 | 23 | 25 | 20 | 24 | 0.900 | - | 40404 |
| S164 | 159 | 451 | G3i | 53K | -50° to +50° | 5° | 7.39 | 25 | 18 | 28 | 24 | 26 | 0.900 | - | 40393 |
| S165 | 160 | 451 | G3i | 53K | -50° to +50° | 5° | 7.39 | 29 | 13 | 27 | 25 | 25 | 0.900 | - | 40394 |
| S166 | 161 | 451 | G3i | 53K | -50° to +50° | 5° | 7.39 | 26 | 24 | 27 | 25 | 26 | 0.900 | - | 40395 |
| S167 | 162 | 451 | G3i | 53K | -50° to +50° | 5° | 7.39 | 25 | 17 | 26 | 24 | 29 | 0.900 | - | 40396 |
| S168 | 163 | 451 | G3i | 53K | -50° to +50° | 5° | 7.39 | 27 | 19 | 30 | 21 | 35 | 0.900 | - | 40397 |
| S169 | 164 | 597 | G3i | 53K | -50° to +50° | 5° | 9.76 | 28 | 18 | 32 | 23 | 27 | 0.900 | - | 40398 |
| S170 | 165 | 597 | G3i | 53K | -50° to +50° | 5° | 9.76 | 27 | 18 | 33 | 23 | 26 | 0.900 | - | 40399 |
| S171 | 166 | 597 | G3i | 53K | -50° to +50° | 5° | 9.76 | 27 | 18 | 31 | 19 | 33 | 0.900 | - | 40400 |
| S172 | 167 | 597 | G3i | 53K | -50° to +50° | 5° | 9.76 | 26 | 15 | 30 | 19 | 24 | 0.900 | - | 40401 |
| S173 | 168 | 597 | G3i | 53K | -50° to +50° | 5° | 9.76 | 28 | 22 | 35 | 25 | 28 | 0.900 | - | 40402 |
| S174 | 169 | 597 | G3i | 53K | -50° to +50° | 5° | 9.76 | 26 | 19 | 26 | 22 | 26 | 0.900 | - | 40403 |
| S175 | 170 | 597 | G3i | 53K | -50° to +50° | 5° | 9.76 | 26 | 16 | 29 | 24 | 29 | 0.900 | - | 40376 |
| **Figure** | **Part. ID** | **Total dose ($e^-/Å^2$)** | **TEM** | **Magnification** | **Angle range** | **Tilt step** | **Exposure time (s)** | **Meth. 1[a]** | **Meth. 2[b]** | **Meth. 3[c]** | **Meth. 4[d]** | **Meth. 5[e]** | **Contour level** | **MP (%)** | **EMDB ID** |
| | | | | | | | | **Resolution (Å)** | | | | | | | |

**Supplementary Video:**

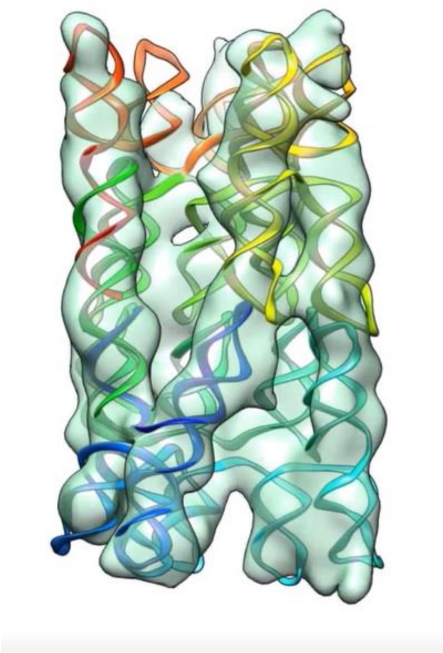

**Supplementary Video 1: The process for 3D reconstruction of individual particles, model fitting, and analysis of structural variability.**

**Supplementary Figures:**

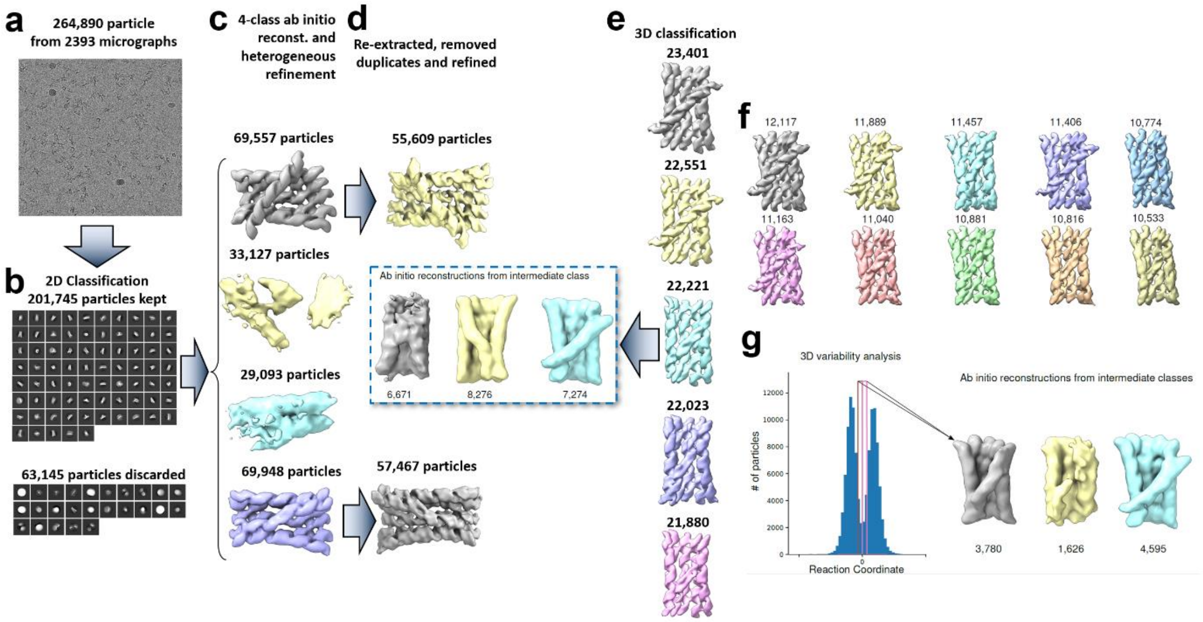

**Supplementary Fig. 1: The analyses of structural heterogeneity from a cryo-EM data set by SPA.** Single particle workflow from micrographs and particle picking (a) to 2D classification to remove junk particles (b) , multi-class *ab initio* reconstruction (c), re-extraction and refinement (d). Particles from the two major classes were re-combined and used for 3D classification without alignment (e,f), multi-class *ab initio* reconstruction using particles from an intermediate class (e, cyan) from the 5 class 3D classification resulted in a low-detail map, and the two major conformations. 3D variability analysis shows two populations with minimal overlap(g), multi-class *ab initio* reconstruction using particles from the overlap region also reproduced the two major conformations and one low detail map.

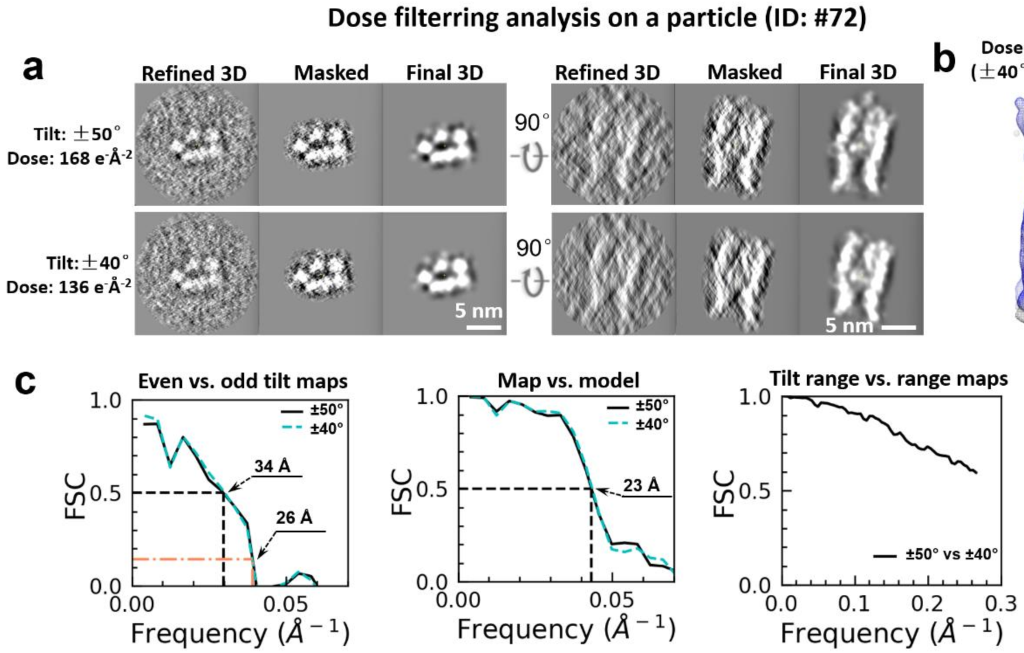

**Supplementary Fig. 2: Analysis of dose-filtering on an individual-particle 3D reconstruction. a,** Comparison of central slices of a 3D map reconstructed from a tilt series acquired at a dose of 168 $e^{-}Å^{-2}$, to their corresponding slices after dose-filtering to 136 $e^{-}Å^{-2}$. The filtering was conducted by removing the last acquired images corresponding to the high-tilt and high-dose images. The central slices of the reconstructions before and after applying the soft-mask and reducing-noise filters are shown in two perpendicular viewing directions. **b,** Overlay of the dose-filtered 3D map on the original map. The original map is shown in gray; the dose of 136 $e^{-}Å^{-2}$ are shown in blue. **c,** Resolution analysis of the dose-filtered map using the two-half maps generated from the even and odd tilt images, respectively. The FSC curve between the two-half maps reconstructed by the even and odd tilt images is calculated at each dose condition. **d,** Resolution analysis of the dose-filtered map using the map vs. fitting model method. The FSC curve between the IEPT reconstruction and the fitting model-generated density map is computed. **e,** Quantitative analysis of map similarity by FSC curves. The FSC curves between each two maps are computed.

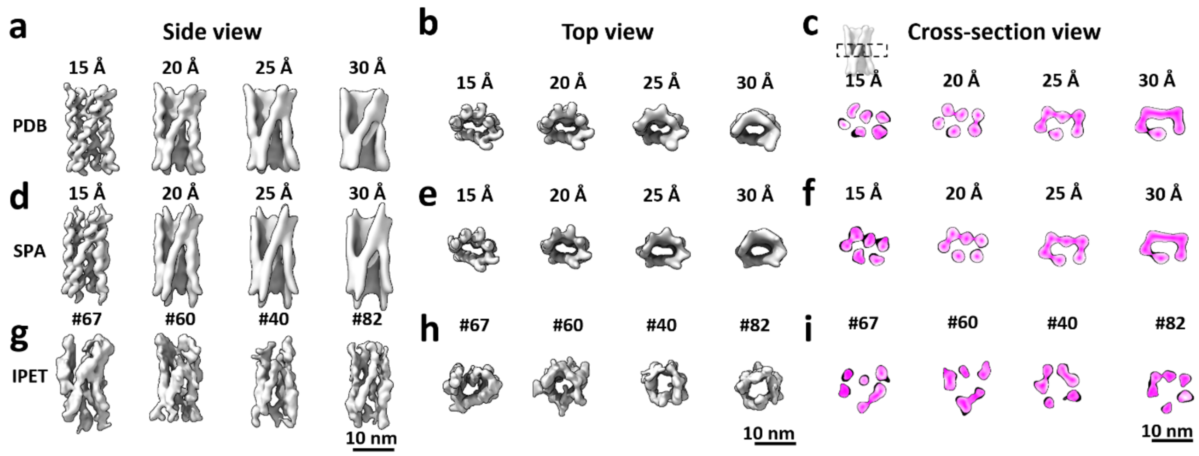

**Supplementary Fig. 3: Evaluating resolution using structural features of RNA helix. a.** Side view of the density map generated from the PDB model of the "mature" conformation, determined by SPA[11] at a dose of 60 e$^{-}$Å$^{-2}$. The map is displayed with low-pass filters at resolutions of 15, 20, 25, and 30 Å. **b,** Top views of the corresponding filtered maps. **c,** Central cross-section views of the filtered maps. **d,** Experimental 3D density map of the "mature" conformation determined by SPA[11], displayed with low-pass filters at resolutions of 15, 20, 25, and 30 Å. **e,** Top views of the corresponding filtered maps. **f,** Central cross-section view of the SPA[11] map. **g,** Four IPET 3D reconstructions corresponding to particles #67, #60, #40, and #82. The maps are displayed from side view, **h,** top view, and **i,** central cross-section view. The cross-section surface is colored according to the density gradient from high to low (purple to white).

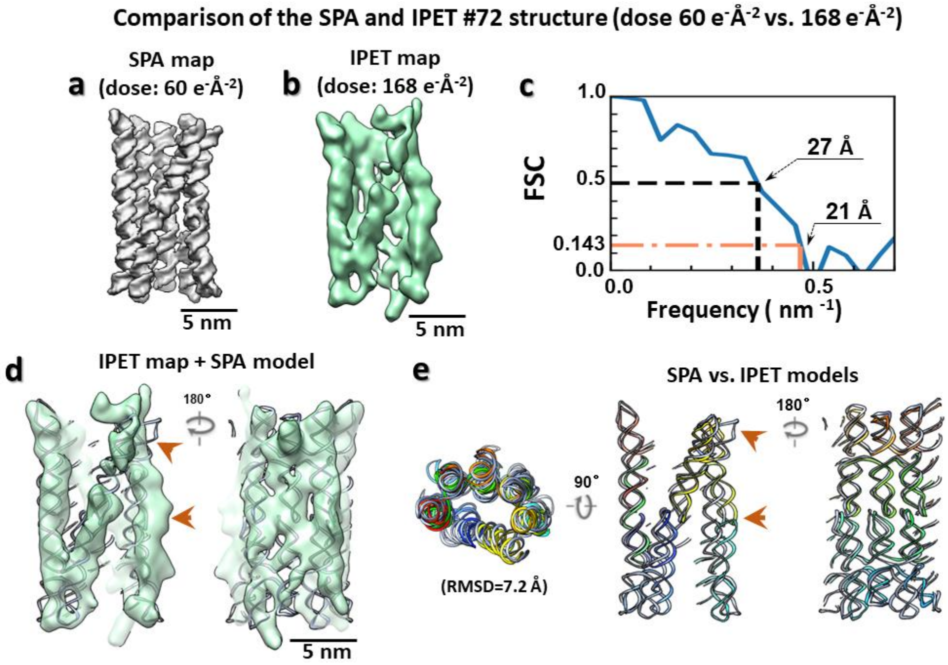

**Supplementary Fig. 4: Comparison of the structure of the SPA mature and IPET obtained at a dose of 60 and 168 e⁻Å$^{-2}$, respectively. a,** The "mature" conformation obtained by conventional SPA[11] at a dose of 60 e⁻Å$^{-2}$ and a resolution of 4.9 Å. **b,** IPET 3D reconstruction of an individual particle (particle # 072), achieved at a dose of 168 e⁻Å$^{-2}$. **c,** FSC analysis between the SPA map and IPET map. **d,** Rigid-body docking of the "mature" structure into the IPET map, displayed from opposite views. **e,** Overlay of the IPET model on the "mature" structure, shown in three views. The major mismatching positions are indicated by orange arrows.

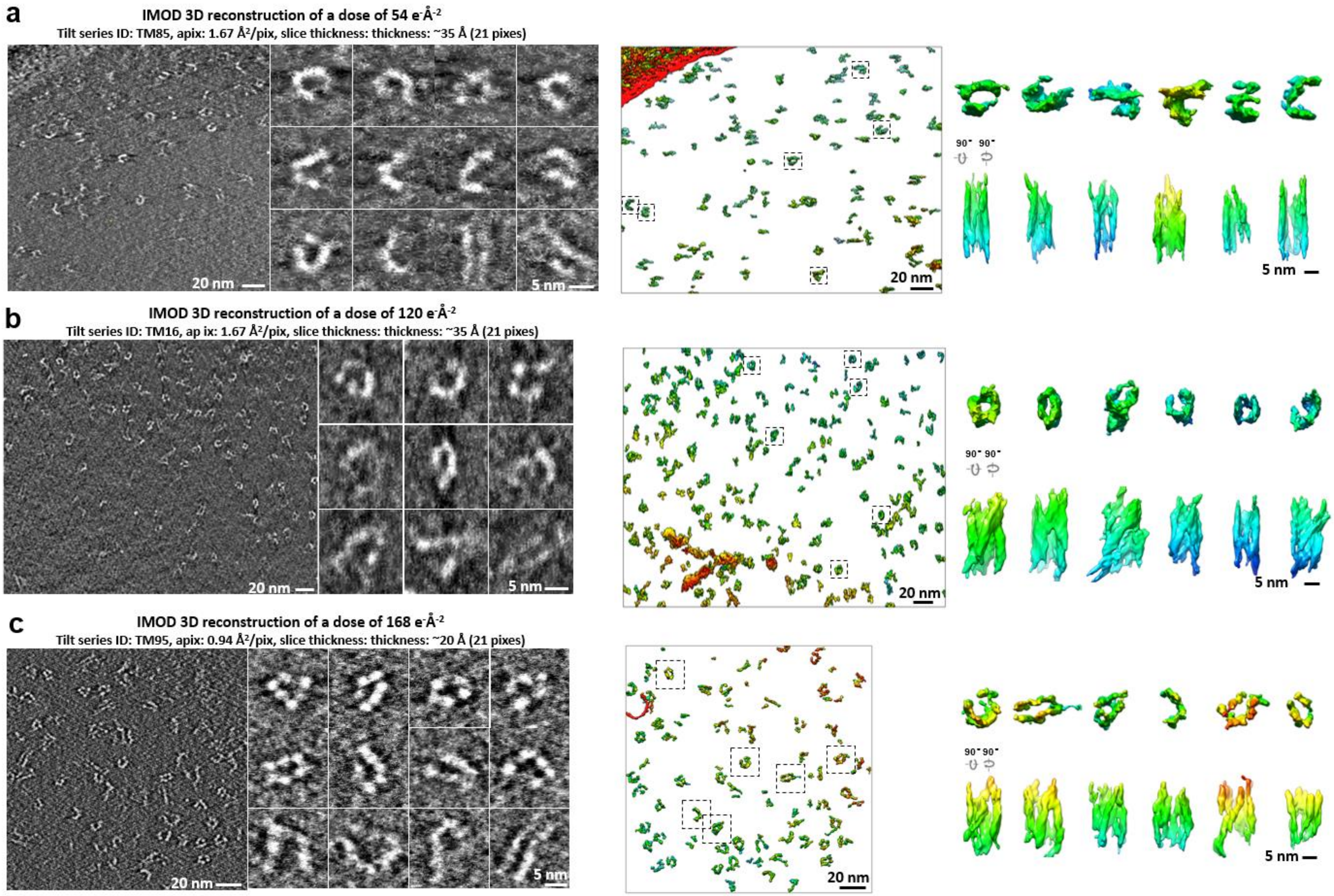
a
IMOD 3D reconstruction of a dose of 54 $e^{-}Å^{-2}$
Tilt series ID: TM85, apix: 1.67 $Å^2$/pix, slice thickness: thickness: ~35 Å (21 pixes)
20 nm
5 nm
20 nm
90° 90°
5 nm
b
IMOD 3D reconstruction of a dose of 120 $e^{-}Å^{-2}$
Tilt series ID: TM16, ap ix: 1.67 $Å^2$/pix, slice thickness: thickness: ~35 Å (21 pixes)
20 nm
5 nm
20 nm
90° 90°
5 nm
c
IMOD 3D reconstruction of a dose of 168 $e^{-}Å^{-2}$
Tilt series ID: TM95, apix: 0.94 $Å^2$/pix, slice thickness: thickness: ~20 Å (21 pixes)
20 nm
5 nm
20 nm
90° 90°
5 nm

**Supplementary Fig. 5: Validation of the 3D reconstruction using IMOD software. a**, The central slice (left top panel) and its 3D reconstruction (left bottom panel) by IMOD software from the cryo-ET tilt series acquired at a dose of 54 $e^{-}Å^{-2}$ (left column). The zoomed-in central slice images of twelve representative particles (right top panel) and 3D sub-volume of six representative particles that were viewed from two perpendicular directions (right bottom panel). **b**, The central slices and 3D reconstructions obtained by IMOD software at a dose of 120 $e^{-}Å^{-2}$. **c**, The central slices and 3D reconstructions achieved by IMOD software at a dose of 168 $e^{-}Å^{-2}$.

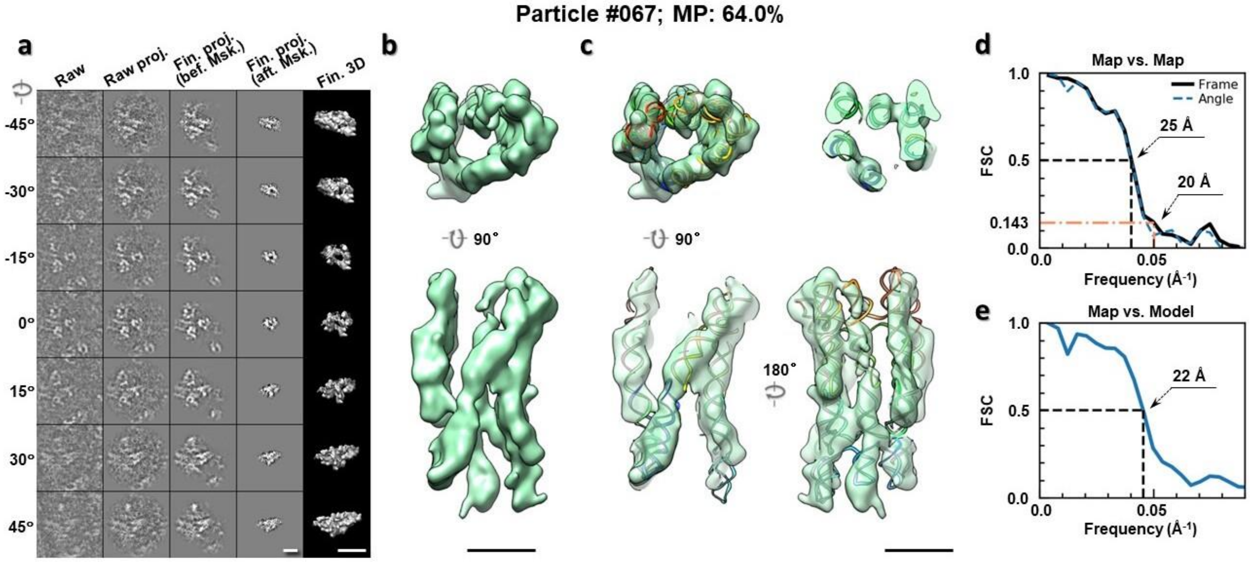

**Supplementary Fig. 6: IPET 3D reconstruction of an individual 6HBC RNA particle #67.** **a,** Seven representative tilt images of a single particle are shown in the first column from the left. The tilt images are aligned to a common center using IPET through iterative refinements. The projections of the raw, intermediate, and final 3D reconstructions at the corresponding angles are displayed in the next four columns. **b,** Two orthogonal views of the final 3D map; **c,** Overlap of the map with the fitted model; **d,** FSC analyses of the final map resolution using two methods, "map-map FSC", in which each map is reconstructed from one half of the images, based on the even vs. odd indices of frames or tilt angles. respectively, **e,** FSC analysis of map-model, where the model map is generated by the fitted model after being low-pass filtered to 8 Å. The resolutions are assessed based on the frequencies of the FSC curve falls at 0.5. Scale bars represent 10 nm in a, and 5 nm in b and c.

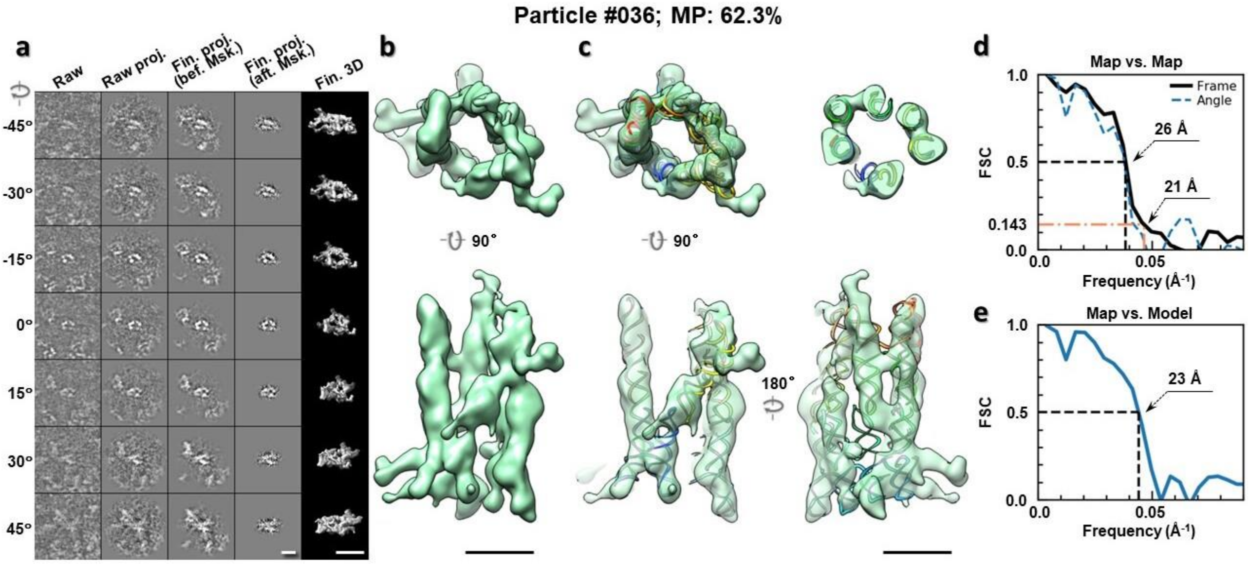

**Supplementary Fig. 7: IPET 3D reconstruction of an individual 6HBC RNA particle #36. a,** Seven representative tilt images of a single particle are shown in the first column from the left. The tilt images are aligned to a common center using IPET through iterative refinements. The projections of the raw, intermediate, and final 3D reconstructions at the corresponding angles are displayed in the next four columns. **b,** Two orthogonal views of the final 3D map; **c,** Overlap of the map with the fitted model; **d,** FSC analyses of the final map resolution using two methods, "map-map FSC", in which each map is reconstructed from one half of the images, based on the even vs. odd indices of frames or tilt angles. respectively, **e,** FSC analysis of map-model, where the model map is generated by the fitted model after being low-pass filtered to 8 Å. The resolutions are assessed based on the frequencies of the FSC curve falls at 0.5. Scale bars represent 10 nm in a, and 5 nm in b and c.

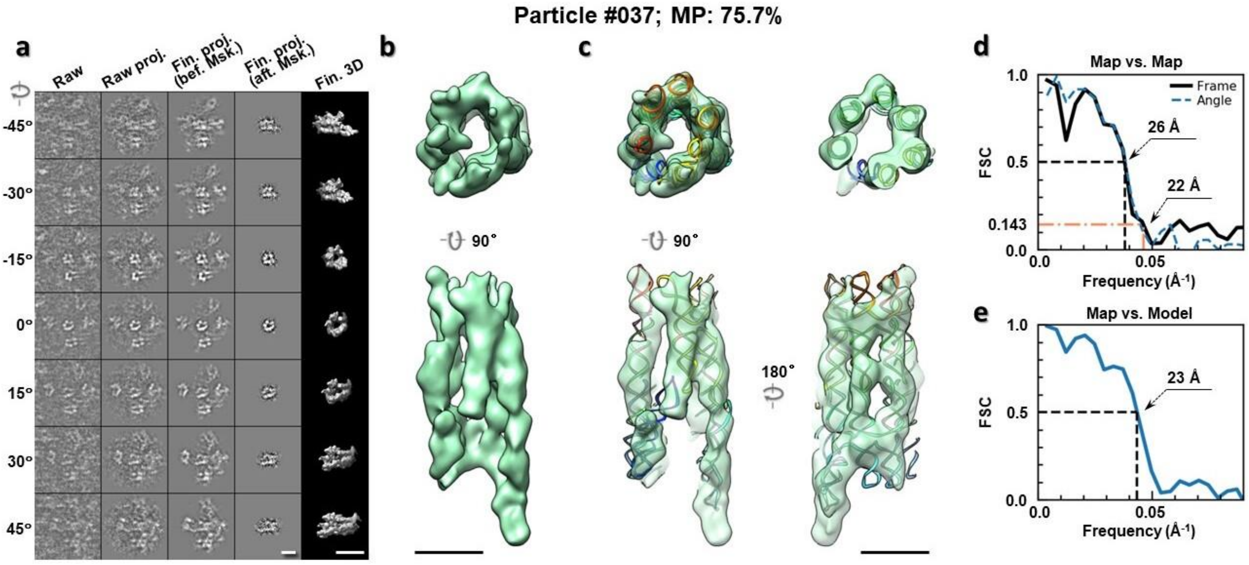

**Supplementary Fig. 8: IPET 3D reconstruction of an individual 6HBC RNA particle #37. a,** Seven representative tilt images of a single particle are shown in the first column from the left. The tilt images are aligned to a common center using IPET through iterative refinements. The projections of the raw, intermediate, and final 3D reconstructions at the corresponding angles are displayed in the next four

columns. **b,** Two orthogonal views of the final 3D map; **c,** Overlap of the map with the fitted model; **d,** FSC analyses of the final map resolution using two methods, "map-map FSC", in which each map is reconstructed from one half of the images, based on the even vs. odd indices of frames or tilt angles. respectively, **e,** FSC analysis of map-model, where the model map is generated by the fitted model after being low-pass filtered to 8 Å. The resolutions are assessed based on the frequencies of the FSC curve falls at 0.5. Scale bars represent 10 nm in a, and 5 nm in b and c.

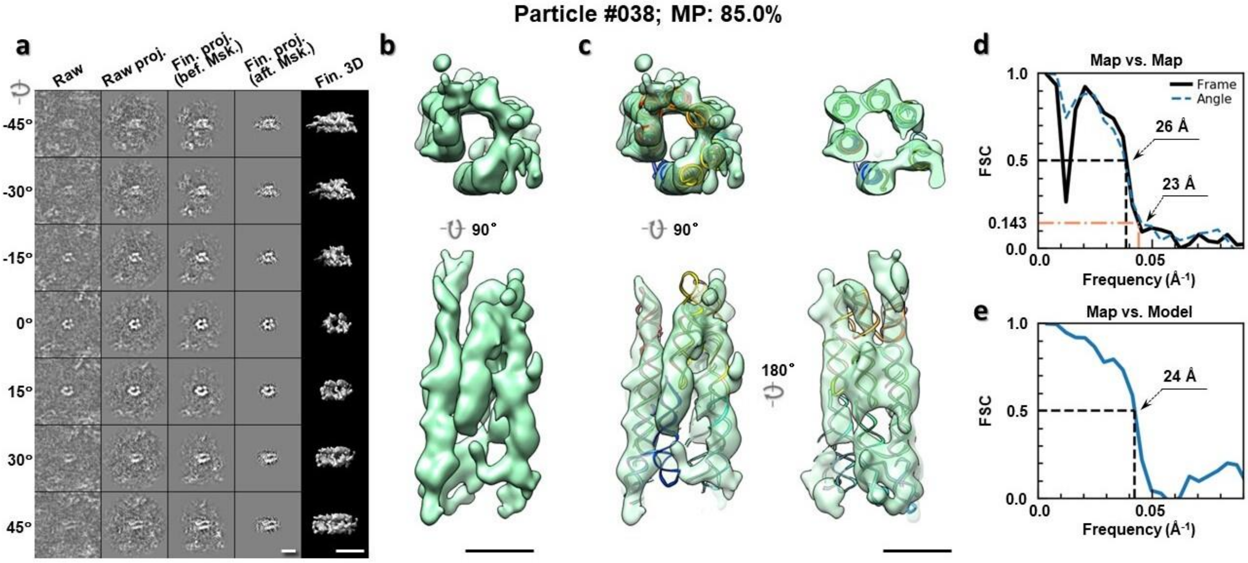

**Supplementary Fig. 9: IPET 3D reconstruction of an individual 6HBC RNA particle #38. a,** Seven representative tilt images of a single particle are shown in the first column from the left. The tilt images are aligned to a common center using IPET through iterative refinements. The projections of the raw, intermediate, and final 3D reconstructions at the corresponding angles are displayed in the next four columns. **b,** Two orthogonal views of the final 3D map; **c,** Overlap of the map with the fitted model; **d,** FSC analyses of the final map resolution using two methods, "map-map FSC", in which each map is reconstructed from one half of the images, based on the even vs. odd indices of frames or tilt angles. respectively, **e,** FSC analysis of map-model, where the model map is generated by the fitted model after being low-pass filtered to 8 Å. The resolutions are assessed based on the frequencies of the FSC curve falls at 0.5. Scale bars represent 10 nm in a, and 5 nm in b and c.

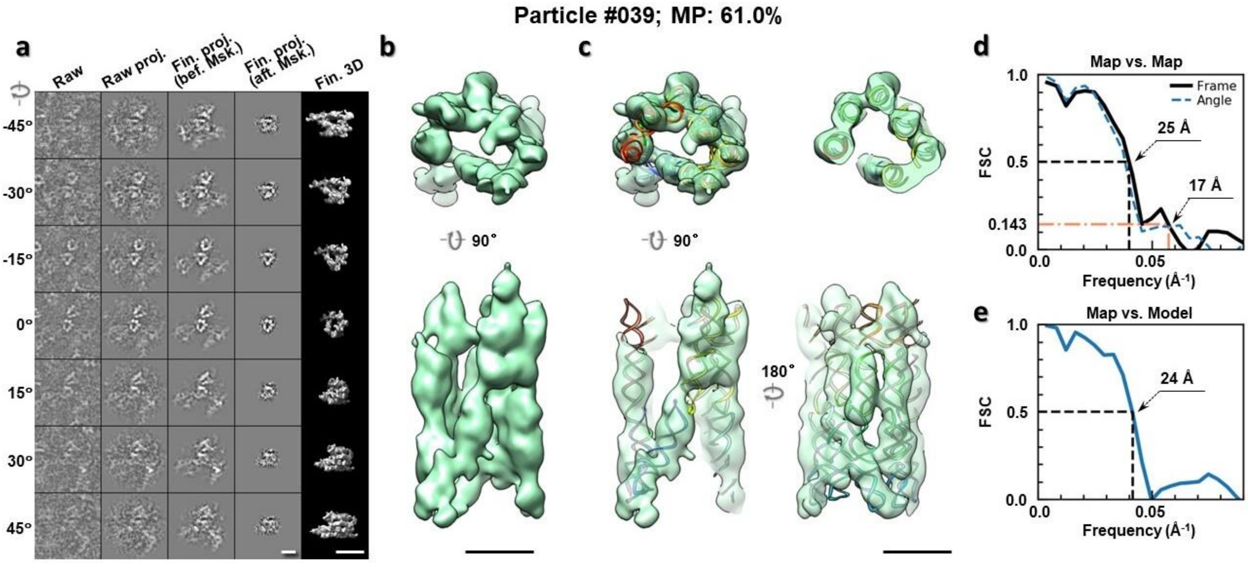

**Supplementary Fig. 10: IPET 3D reconstruction of an individual 6HBC RNA particle #39.** **a,** Seven representative tilt images of a single particle are shown in the first column from the left. The tilt images are aligned to a common center using IPET through iterative refinements. The projections of the raw, intermediate, and final 3D reconstructions at the corresponding angles are displayed in the next four columns. **b,** Two orthogonal views of the final 3D map; **c,** Overlap of the map with the fitted model; **d,** FSC analyses of the final map resolution using two methods, "map-map FSC", in which each map is reconstructed from one half of the images, based on the even vs. odd indices of frames or tilt angles. respectively, **e,** FSC analysis of map-model, where the model map is generated by the fitted model after being low-pass filtered to 8 Å. The resolutions are assessed based on the frequencies of the FSC curve falls at 0.5. Scale bars represent 10 nm in a, and 5 nm in b and c.

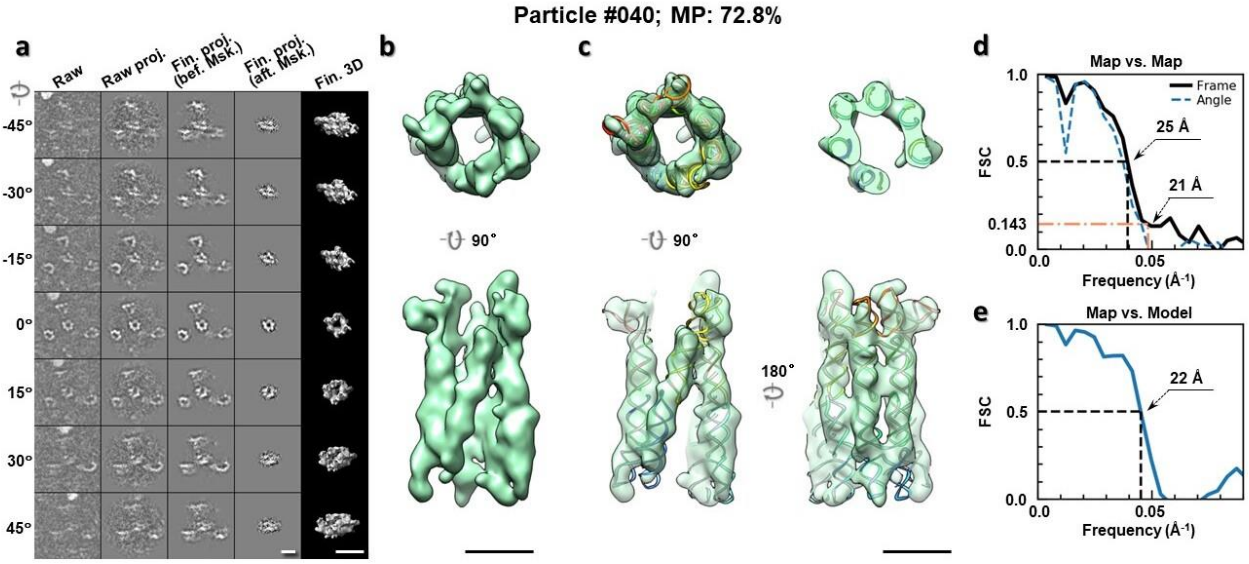

**Supplementary Fig. 11: IPET 3D reconstruction of an individual 6HBC RNA particle #40.** **a,** Seven representative tilt images of a single particle are shown in the first column from the left. The tilt images are aligned to a common center using IPET through iterative refinements. The projections of the raw, intermediate, and final 3D reconstructions at the corresponding angles are displayed in the next four

columns. **b,** Two orthogonal views of the final 3D map; **c,** Overlap of the map with the fitted model; **d,** FSC analyses of the final map resolution using two methods, "map-map FSC", in which each map is reconstructed from one half of the images, based on the even vs. odd indices of frames or tilt angles. respectively, **e,** FSC analysis of map-model, where the model map is generated by the fitted model after being low-pass filtered to 8 Å. The resolutions are assessed based on the frequencies of the FSC curve falls at 0.5. Scale bars represent 10 nm in a, and 5 nm in b and c.

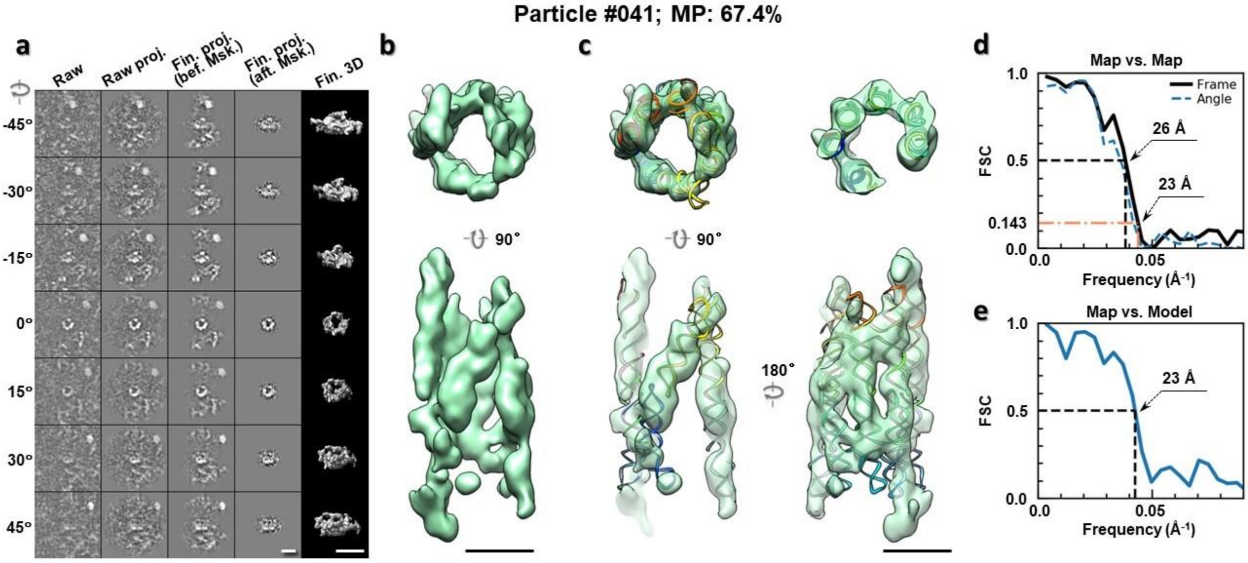

**Supplementary Fig. 12: IPET 3D reconstruction of an individual 6HBC RNA particle #41.** **a,** Seven representative tilt images of a single particle are shown in the first column from the left. The tilt images are aligned to a common center using IPET through iterative refinements. The projections of the raw, intermediate, and final 3D reconstructions at the corresponding angles are displayed in the next four columns. **b,** Two orthogonal views of the final 3D map; **c,** Overlap of the map with the fitted model; **d,** FSC analyses of the final map resolution using two methods, "map-map FSC", in which each map is reconstructed from one half of the images, based on the even vs. odd indices of frames or tilt angles. respectively, **e,** FSC analysis of map-model, where the model map is generated by the fitted model after being low-pass filtered to 8 Å. The resolutions are assessed based on the frequencies of the FSC curve falls at 0.5. Scale bars represent 10 nm in a, and 5 nm in b and c.

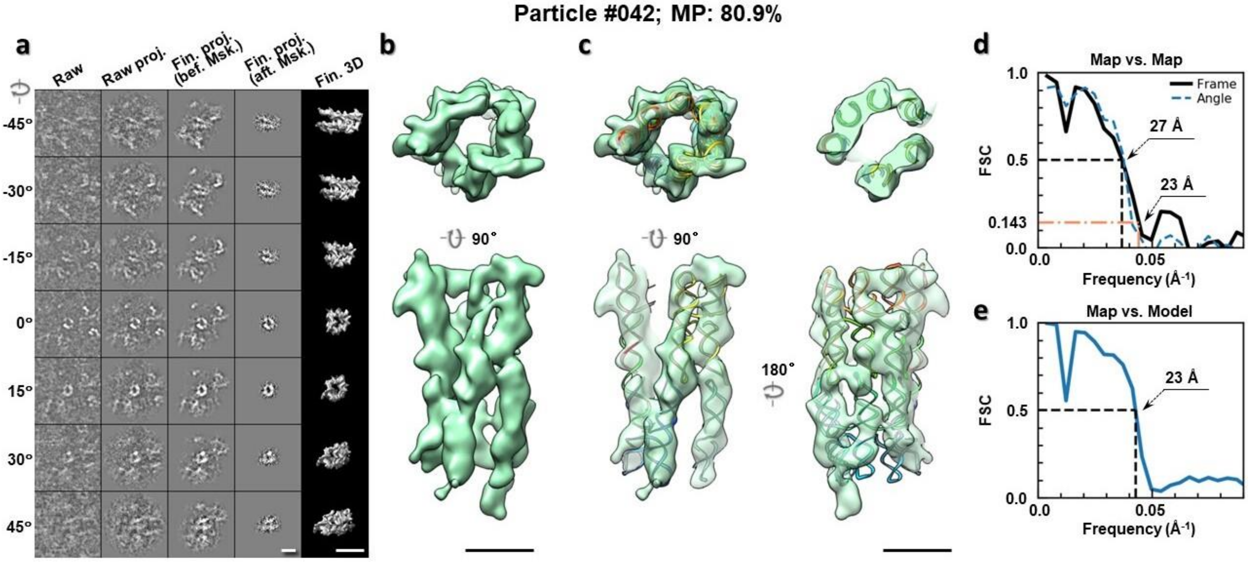

**Supplementary Fig. 13: IPET 3D reconstruction of an individual 6HBC RNA particle #42.** **a,** Seven representative tilt images of a single particle are shown in the first column from the left. The tilt images are aligned to a common center using IPET through iterative refinements. The projections of the raw, intermediate, and final 3D reconstructions at the corresponding angles are displayed in the next four columns. **b,** Two orthogonal views of the final 3D map; **c,** Overlap of the map with the fitted model; **d,** FSC analyses of the final map resolution using two methods, "map-map FSC", in which each map is reconstructed from one half of the images, based on the even vs. odd indices of frames or tilt angles. respectively, **e,** FSC analysis of map-model, where the model map is generated by the fitted model after being low-pass filtered to 8 Å. The resolutions are assessed based on the frequencies of the FSC curve falls at 0.5. Scale bars represent 10 nm in a, and 5 nm in b and c.

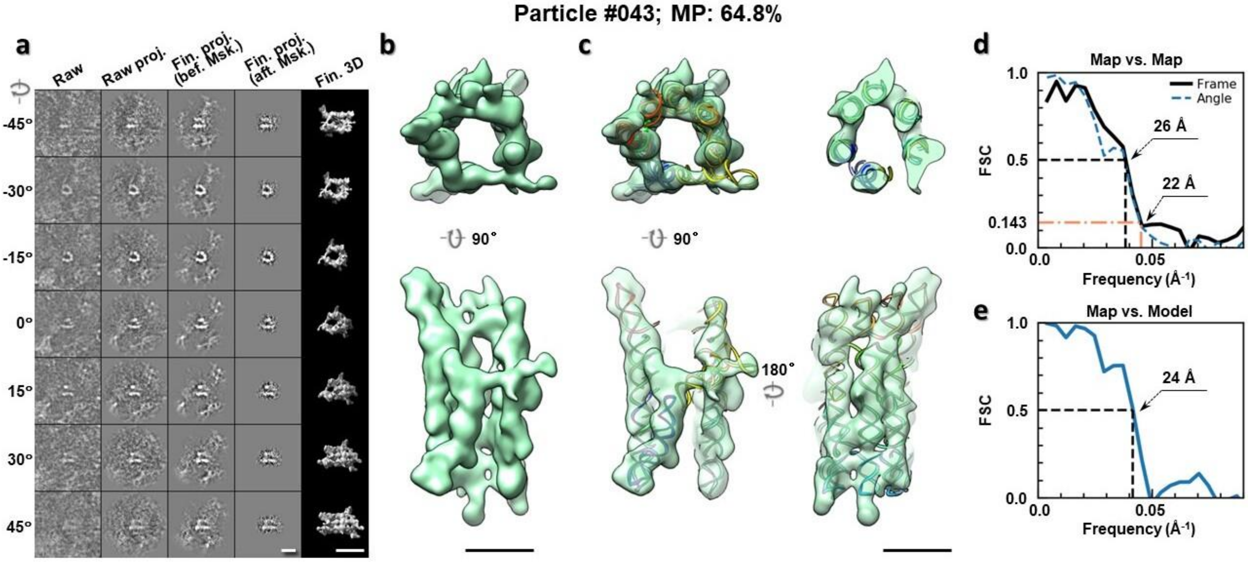

**Supplementary Fig. 14: IPET 3D reconstruction of an individual 6HBC RNA particle #43.** **a,** Seven representative tilt images of a single particle are shown in the first column from the left. The tilt images are aligned to a common center using IPET through iterative refinements. The projections of the raw, intermediate, and final 3D reconstructions at the corresponding angles are displayed in the next four

columns. **b,** Two orthogonal views of the final 3D map; **c,** Overlap of the map with the fitted model; **d,** FSC analyses of the final map resolution using two methods, "map-map FSC", in which each map is reconstructed from one half of the images, based on the even vs. odd indices of frames or tilt angles. respectively, **e,** FSC analysis of map-model, where the model map is generated by the fitted model after being low-pass filtered to 8 Å. The resolutions are assessed based on the frequencies of the FSC curve falls at 0.5. Scale bars represent 10 nm in a, and 5 nm in b and c.

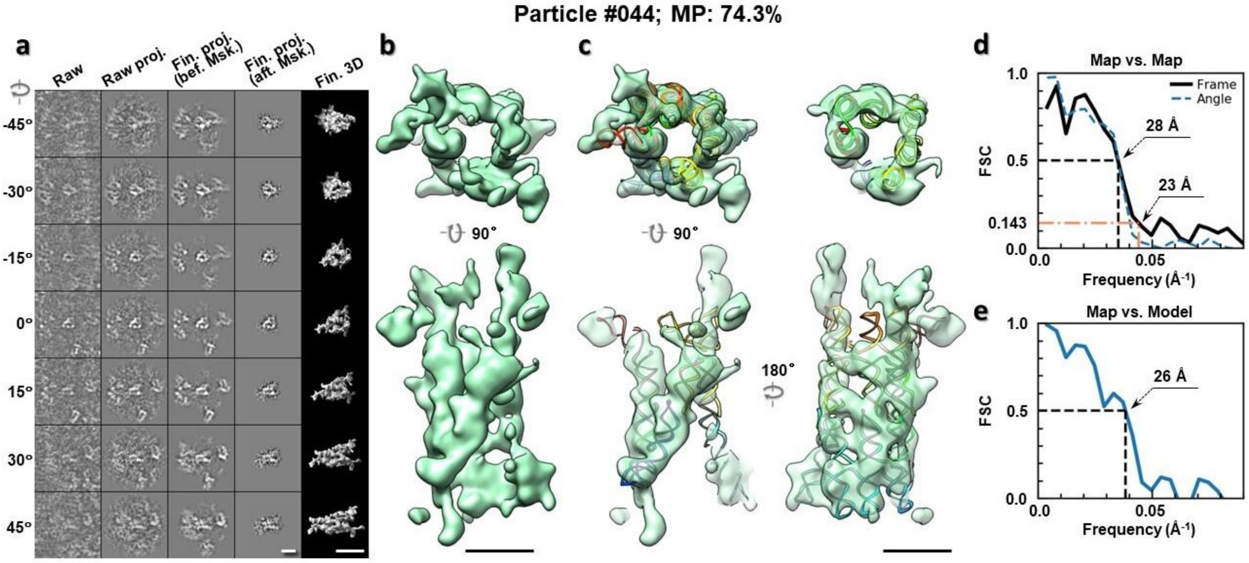

**Supplementary Fig. 15: IPET 3D reconstruction of an individual 6HBC RNA particle #44. a,** Seven representative tilt images of a single particle are shown in the first column from the left. The tilt images are aligned to a common center using IPET through iterative refinements. The projections of the raw, intermediate, and final 3D reconstructions at the corresponding angles are displayed in the next four columns. **b,** Two orthogonal views of the final 3D map; **c,** Overlap of the map with the fitted model; **d,** FSC analyses of the final map resolution using two methods, "map-map FSC", in which each map is reconstructed from one half of the images, based on the even vs. odd indices of frames or tilt angles. respectively, **e,** FSC analysis of map-model, where the model map is generated by the fitted model after being low-pass filtered to 8 Å. The resolutions are assessed based on the frequencies of the FSC curve falls at 0.5. Scale bars represent 10 nm in a, and 5 nm in b and c.

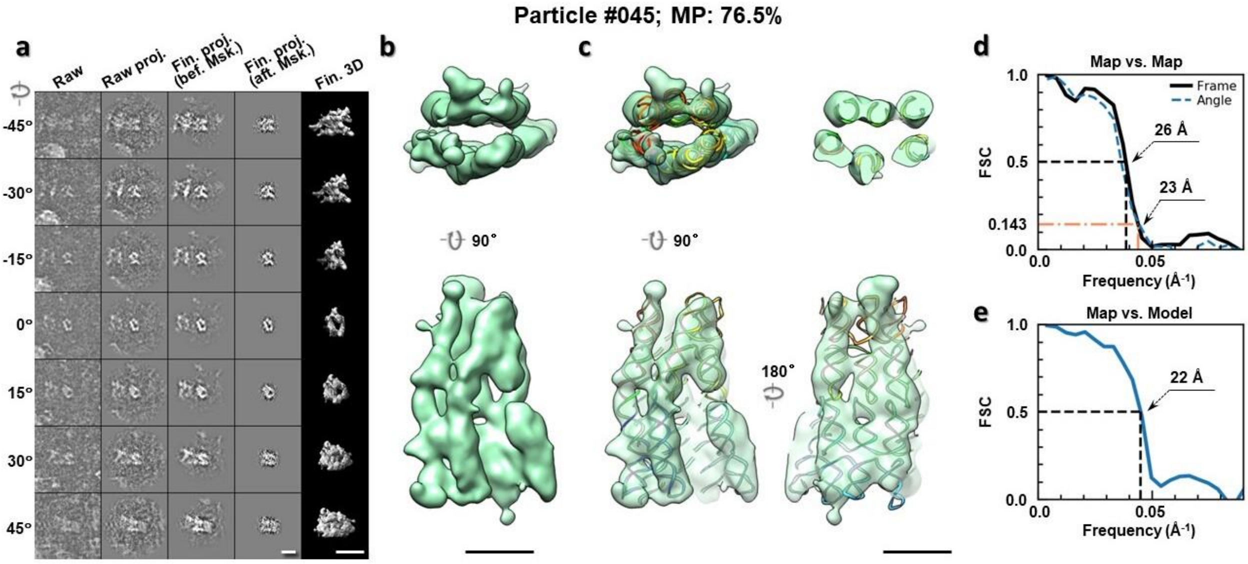

**Supplementary Fig. 16: IPET 3D reconstruction of an individual 6HBC RNA particle #45.** **a,** Seven representative tilt images of a single particle are shown in the first column from the left. The tilt images are aligned to a common center using IPET through iterative refinements. The projections of the raw, intermediate, and final 3D reconstructions at the corresponding angles are displayed in the next four columns. **b,** Two orthogonal views of the final 3D map; **c,** Overlap of the map with the fitted model; **d,** FSC analyses of the final map resolution using two methods, "map-map FSC", in which each map is reconstructed from one half of the images, based on the even vs. odd indices of frames or tilt angles. respectively, **e,** FSC analysis of map-model, where the model map is generated by the fitted model after being low-pass filtered to 8 Å. The resolutions are assessed based on the frequencies of the FSC curve falls at 0.5. Scale bars represent 10 nm in a, and 5 nm in b and c.

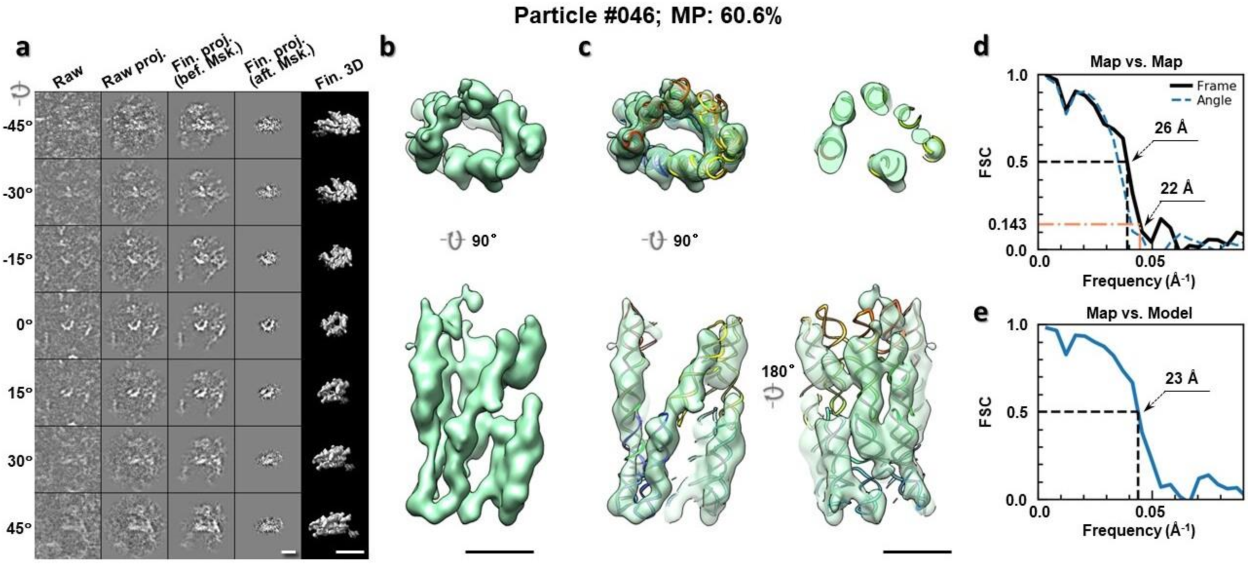

**Supplementary Fig. 17: IPET 3D reconstruction of an individual 6HBC RNA particle #46.** **a,** Seven representative tilt images of a single particle are shown in the first column from the left. The tilt images are aligned to a common center using IPET through iterative refinements. The projections of the raw, intermediate, and final 3D reconstructions at the corresponding angles are displayed in the next four

columns. **b,** Two orthogonal views of the final 3D map; **c,** Overlap of the map with the fitted model; **d,** FSC analyses of the final map resolution using two methods, "map-map FSC", in which each map is reconstructed from one half of the images, based on the even vs. odd indices of frames or tilt angles. respectively, **e,** FSC analysis of map-model, where the model map is generated by the fitted model after being low-pass filtered to 8 Å. The resolutions are assessed based on the frequencies of the FSC curve falls at 0.5. Scale bars represent 10 nm in a, and 5 nm in b and c.

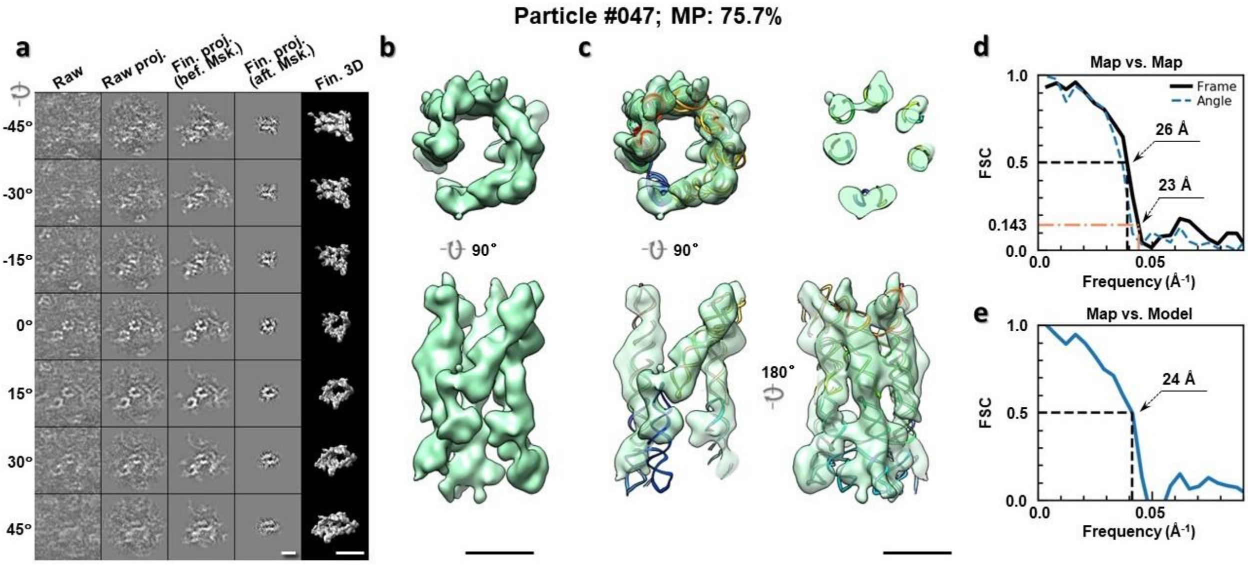

**Supplementary Fig. 18: IPET 3D reconstruction of an individual 6HBC RNA particle #47. a,** Seven representative tilt images of a single particle are shown in the first column from the left. The tilt images are aligned to a common center using IPET through iterative refinements. The projections of the raw, intermediate, and final 3D reconstructions at the corresponding angles are displayed in the next four columns. **b,** Two orthogonal views of the final 3D map; **c,** Overlap of the map with the fitted model; **d,** FSC analyses of the final map resolution using two methods, "map-map FSC", in which each map is reconstructed from one half of the images, based on the even vs. odd indices of frames or tilt angles. respectively, **e,** FSC analysis of map-model, where the model map is generated by the fitted model after being low-pass filtered to 8 Å. The resolutions are assessed based on the frequencies of the FSC curve falls at 0.5. Scale bars represent 10 nm in a, and 5 nm in b and c.

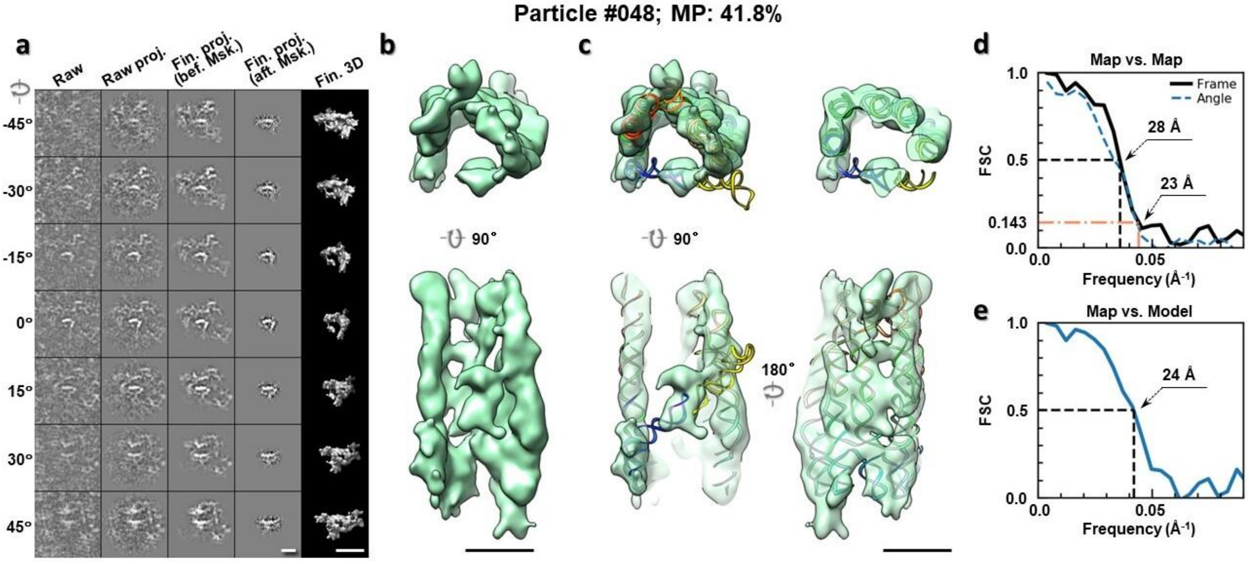

**Supplementary Fig. 19: IPET 3D reconstruction of an individual 6HBC RNA particle #48. a,** Seven representative tilt images of a single particle are shown in the first column from the left. The tilt images are aligned to a common center using IPET through iterative refinements. The projections of the raw, intermediate, and final 3D reconstructions at the corresponding angles are displayed in the next four columns. **b,** Two orthogonal views of the final 3D map; **c,** Overlap of the map with the fitted model; **d,** FSC analyses of the final map resolution using two methods, "map-map FSC", in which each map is reconstructed from one half of the images, based on the even vs. odd indices of frames or tilt angles. respectively, **e,** FSC analysis of map-model, where the model map is generated by the fitted model after being low-pass filtered to 8 Å. The resolutions are assessed based on the frequencies of the FSC curve falls at 0.5. Scale bars represent 10 nm in a, and 5 nm in b and c.

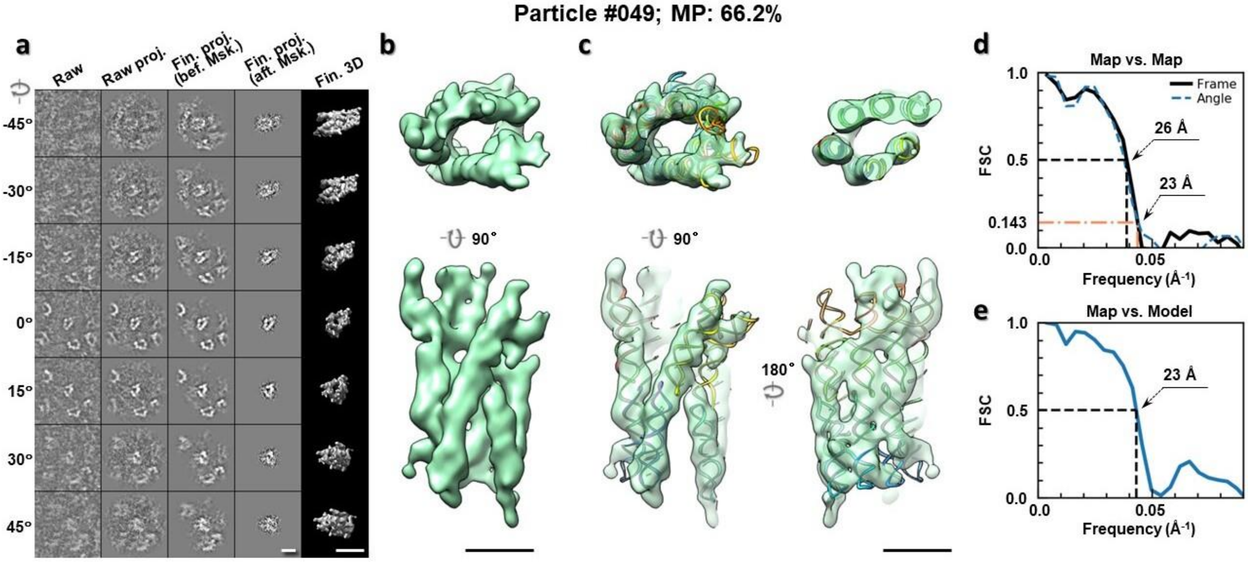

**Supplementary Fig. 20: IPET 3D reconstruction of an individual 6HBC RNA particle #49. a,** Seven representative tilt images of a single particle are shown in the first column from the left. The tilt images are aligned to a common center using IPET through iterative refinements. The projections of the raw, intermediate, and final 3D reconstructions at the corresponding angles are displayed in the next four

columns. **b,** Two orthogonal views of the final 3D map; **c,** Overlap of the map with the fitted model; **d,** FSC analyses of the final map resolution using two methods, "map-map FSC", in which each map is reconstructed from one half of the images, based on the even vs. odd indices of frames or tilt angles. respectively, **e,** FSC analysis of map-model, where the model map is generated by the fitted model after being low-pass filtered to 8 Å. The resolutions are assessed based on the frequencies of the FSC curve falls at 0.5. Scale bars represent 10 nm in a, and 5 nm in b and c.

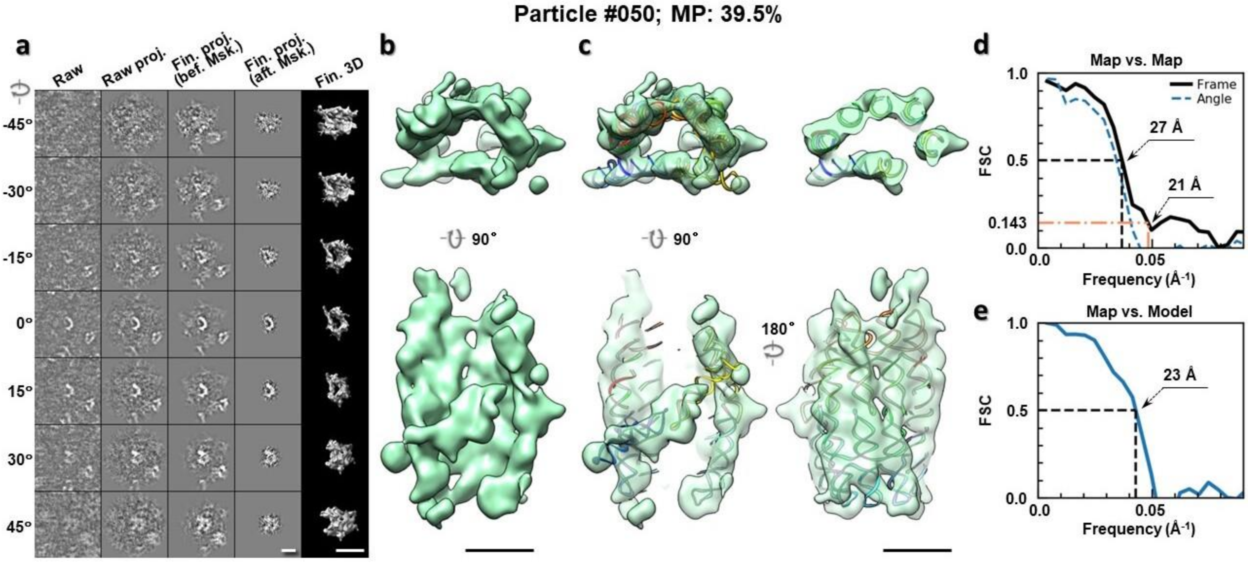

**Supplementary Fig. 21: IPET 3D reconstruction of an individual 6HBC RNA particle #50. a,** Seven representative tilt images of a single particle are shown in the first column from the left. The tilt images are aligned to a common center using IPET through iterative refinements. The projections of the raw, intermediate, and final 3D reconstructions at the corresponding angles are displayed in the next four columns. **b,** Two orthogonal views of the final 3D map; **c,** Overlap of the map with the fitted model; **d,** FSC analyses of the final map resolution using two methods, "map-map FSC", in which each map is reconstructed from one half of the images, based on the even vs. odd indices of frames or tilt angles. respectively, **e,** FSC analysis of map-model, where the model map is generated by the fitted model after being low-pass filtered to 8 Å. The resolutions are assessed based on the frequencies of the FSC curve falls at 0.5. Scale bars represent 10 nm in a, and 5 nm in b and c.

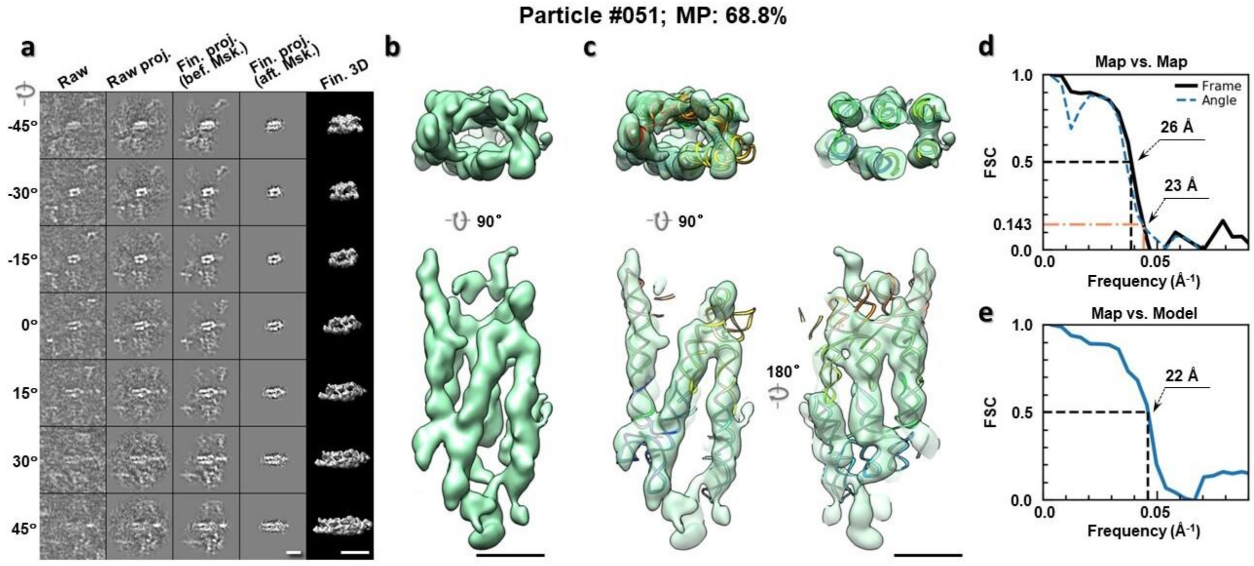

**Supplementary Fig. 22: IPET 3D reconstruction of an individual 6HBC RNA particle #51. a,** Seven representative tilt images of a single particle are shown in the first column from the left. The tilt images are aligned to a common center using IPET through iterative refinements. The projections of the raw, intermediate, and final 3D reconstructions at the corresponding angles are displayed in the next four columns. **b,** Two orthogonal views of the final 3D map; **c,** Overlap of the map with the fitted model; **d,** FSC analyses of the final map resolution using two methods, "map-map FSC", in which each map is reconstructed from one half of the images, based on the even vs. odd indices of frames or tilt angles. respectively, **e,** FSC analysis of map-model, where the model map is generated by the fitted model after being low-pass filtered to 8 Å. The resolutions are assessed based on the frequencies of the FSC curve falls at 0.5. Scale bars represent 10 nm in a, and 5 nm in b and c.

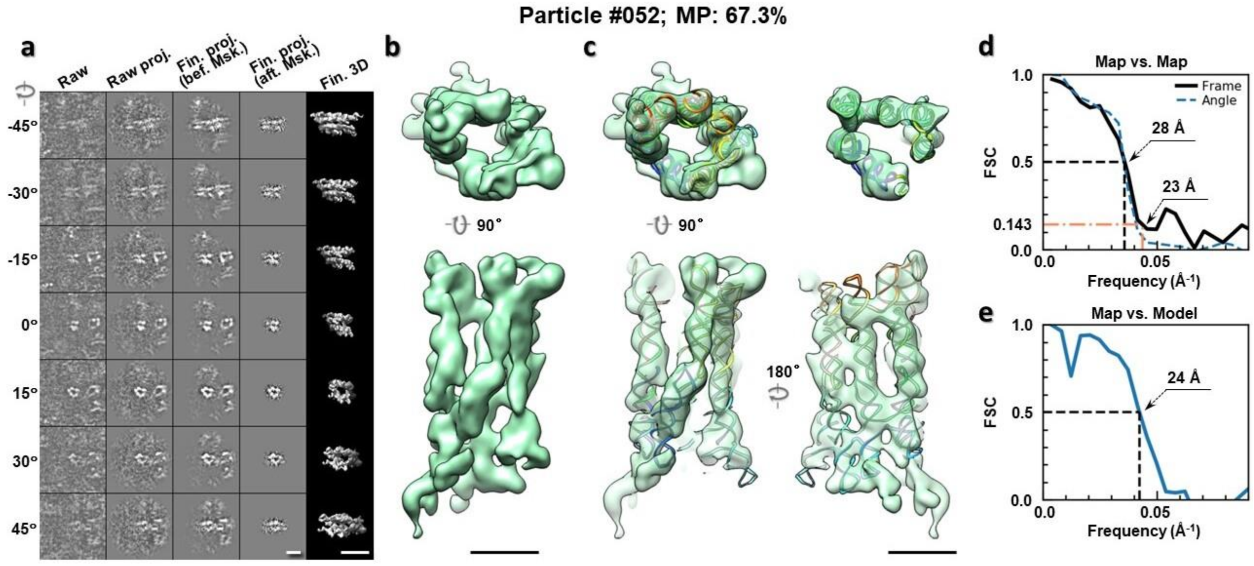

**Supplementary Fig. 23: IPET 3D reconstruction of an individual 6HBC RNA particle #52. a,** Seven representative tilt images of a single particle are shown in the first column from the left. The tilt images are aligned to a common center using IPET through iterative refinements. The projections of the raw, intermediate, and final 3D reconstructions at the corresponding angles are displayed in the next four

columns. **b,** Two orthogonal views of the final 3D map; **c,** Overlap of the map with the fitted model; **d,** FSC analyses of the final map resolution using two methods, "map-map FSC", in which each map is reconstructed from one half of the images, based on the even vs. odd indices of frames or tilt angles. respectively, **e,** FSC analysis of map-model, where the model map is generated by the fitted model after being low-pass filtered to 8 Å. The resolutions are assessed based on the frequencies of the FSC curve falls at 0.5. Scale bars represent 10 nm in a, and 5 nm in b and c.

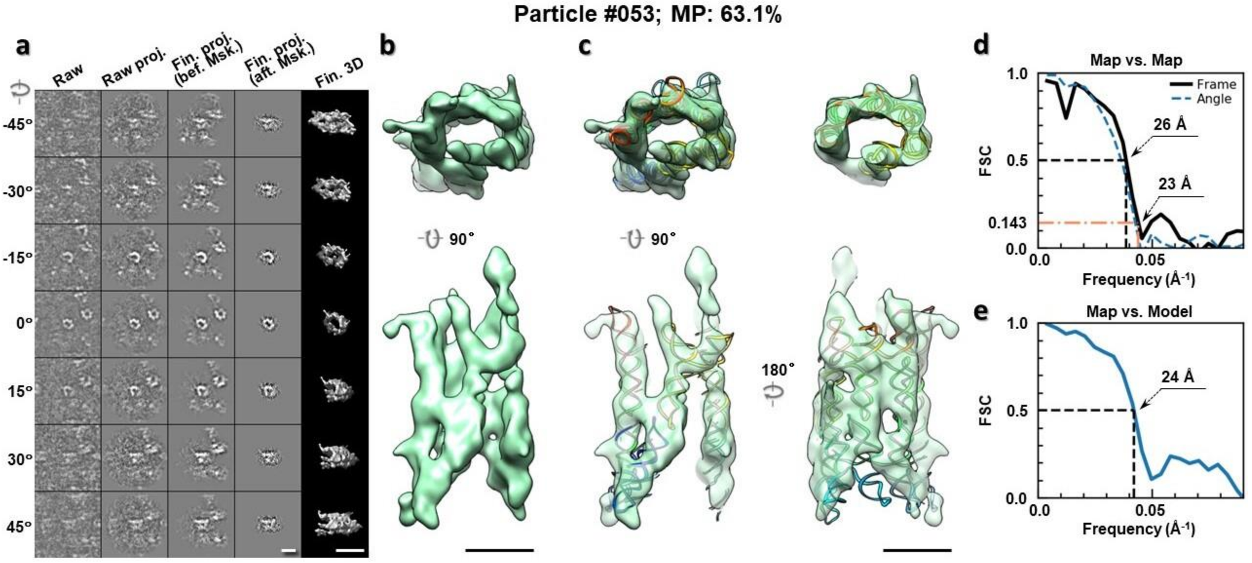

**Supplementary Fig. 24: IPET 3D reconstruction of an individual 6HBC RNA particle #53. a,** Seven representative tilt images of a single particle are shown in the first column from the left. The tilt images are aligned to a common center using IPET through iterative refinements. The projections of the raw, intermediate, and final 3D reconstructions at the corresponding angles are displayed in the next four columns. **b,** Two orthogonal views of the final 3D map; **c,** Overlap of the map with the fitted model; **d,** FSC analyses of the final map resolution using two methods, "map-map FSC", in which each map is reconstructed from one half of the images, based on the even vs. odd indices of frames or tilt angles. respectively, **e,** FSC analysis of map-model, where the model map is generated by the fitted model after being low-pass filtered to 8 Å. The resolutions are assessed based on the frequencies of the FSC curve falls at 0.5. Scale bars represent 10 nm in a, and 5 nm in b and c.

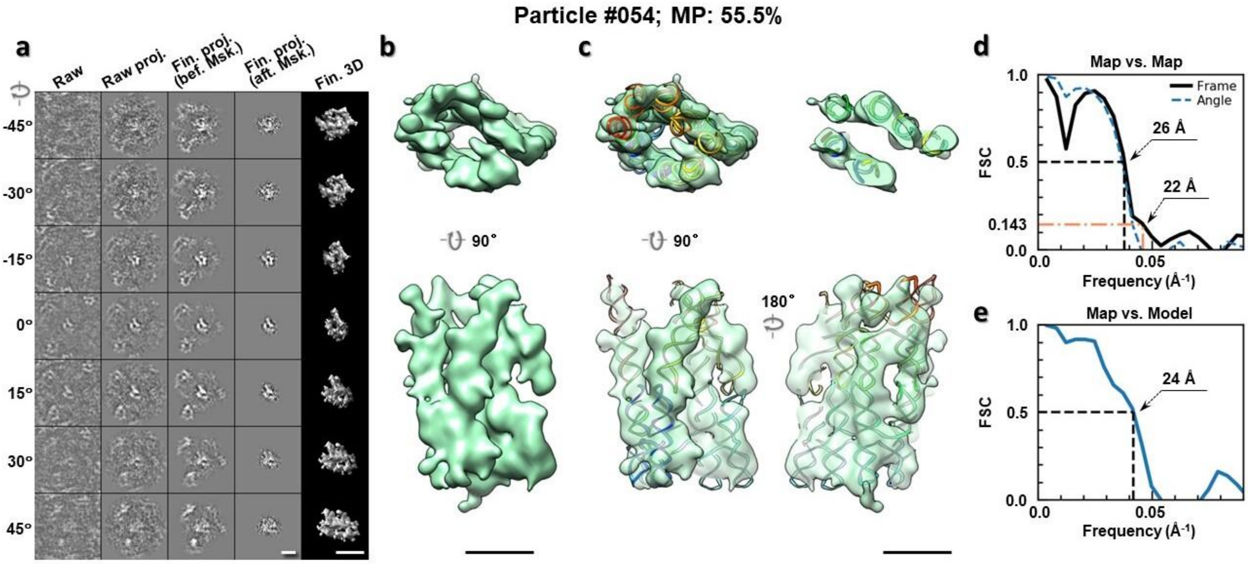

**Supplementary Fig. 25: IPET 3D reconstruction of an individual 6HBC RNA particle #54.** **a,** Seven representative tilt images of a single particle are shown in the first column from the left. The tilt images are aligned to a common center using IPET through iterative refinements. The projections of the raw, intermediate, and final 3D reconstructions at the corresponding angles are displayed in the next four columns. **b,** Two orthogonal views of the final 3D map; **c,** Overlap of the map with the fitted model; **d,** FSC analyses of the final map resolution using two methods, "map-map FSC", in which each map is reconstructed from one half of the images, based on the even vs. odd indices of frames or tilt angles. respectively, **e,** FSC analysis of map-model, where the model map is generated by the fitted model after being low-pass filtered to 8 Å. The resolutions are assessed based on the frequencies of the FSC curve falls at 0.5. Scale bars represent 10 nm in a, and 5 nm in b and c.

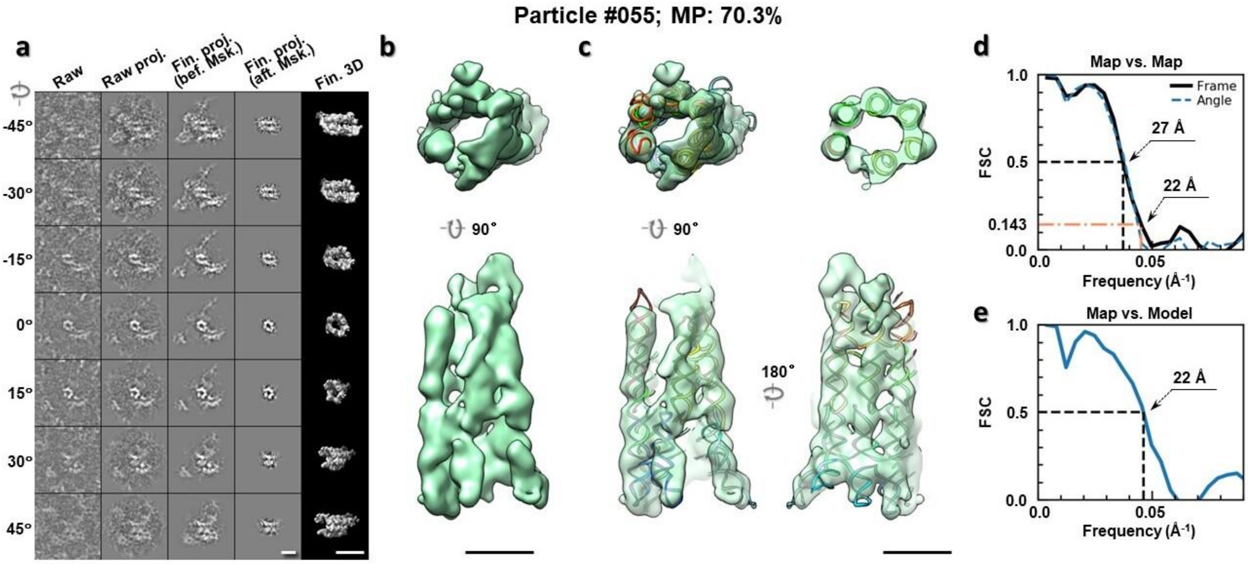

**Supplementary Fig. 26: IPET 3D reconstruction of an individual 6HBC RNA particle #55.** **a,** Seven representative tilt images of a single particle are shown in the first column from the left. The tilt images are aligned to a common center using IPET through iterative refinements. The projections of the raw, intermediate, and final 3D reconstructions at the corresponding angles are displayed in the next four

columns. **b,** Two orthogonal views of the final 3D map; **c,** Overlap of the map with the fitted model; **d,** FSC analyses of the final map resolution using two methods, "map-map FSC", in which each map is reconstructed from one half of the images, based on the even vs. odd indices of frames or tilt angles. respectively, **e,** FSC analysis of map-model, where the model map is generated by the fitted model after being low-pass filtered to 8 Å. The resolutions are assessed based on the frequencies of the FSC curve falls at 0.5. Scale bars represent 10 nm in a, and 5 nm in b and c.

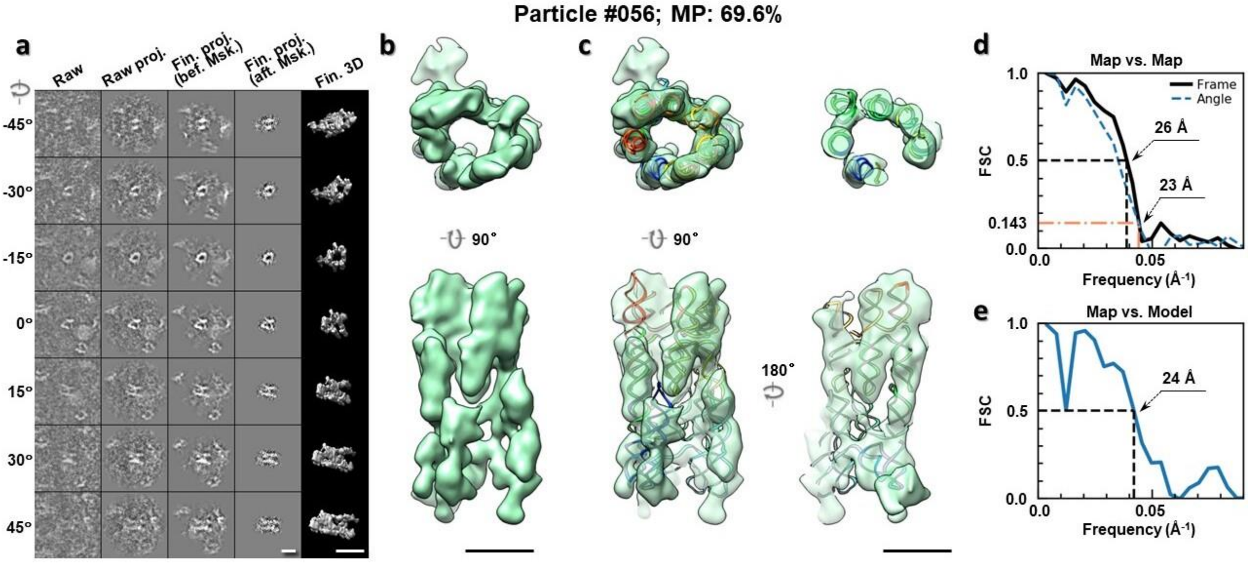

**Supplementary Fig. 27: IPET 3D reconstruction of an individual 6HBC RNA particle #56.** **a,** Seven representative tilt images of a single particle are shown in the first column from the left. The tilt images are aligned to a common center using IPET through iterative refinements. The projections of the raw, intermediate, and final 3D reconstructions at the corresponding angles are displayed in the next four columns. **b,** Two orthogonal views of the final 3D map; **c,** Overlap of the map with the fitted model; **d,** FSC analyses of the final map resolution using two methods, "map-map FSC", in which each map is reconstructed from one half of the images, based on the even vs. odd indices of frames or tilt angles. respectively, **e,** FSC analysis of map-model, where the model map is generated by the fitted model after being low-pass filtered to 8 Å. The resolutions are assessed based on the frequencies of the FSC curve falls at 0.5. Scale bars represent 10 nm in a, and 5 nm in b and c.

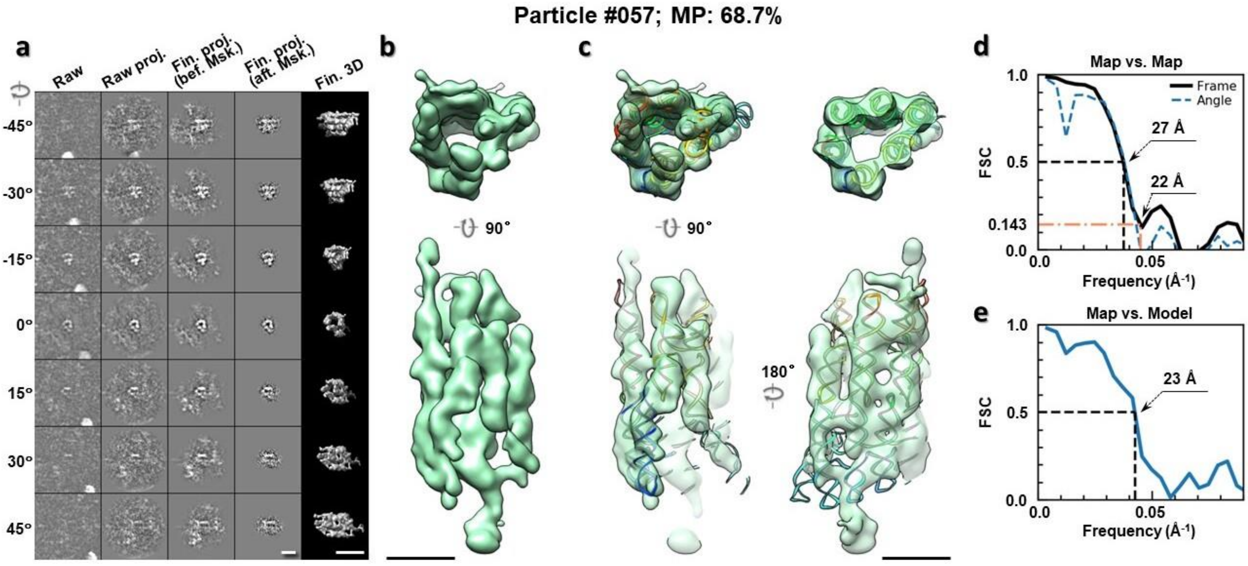

**Supplementary Fig. 28: IPET 3D reconstruction of an individual 6HBC RNA particle #57.** **a,** Seven representative tilt images of a single particle are shown in the first column from the left. The tilt images are aligned to a common center using IPET through iterative refinements. The projections of the raw, intermediate, and final 3D reconstructions at the corresponding angles are displayed in the next four columns. **b,** Two orthogonal views of the final 3D map; **c,** Overlap of the map with the fitted model; **d,** FSC analyses of the final map resolution using two methods, "map-map FSC", in which each map is reconstructed from one half of the images, based on the even vs. odd indices of frames or tilt angles. respectively, **e,** FSC analysis of map-model, where the model map is generated by the fitted model after being low-pass filtered to 8 Å. The resolutions are assessed based on the frequencies of the FSC curve falls at 0.5. Scale bars represent 10 nm in a, and 5 nm in b and c.

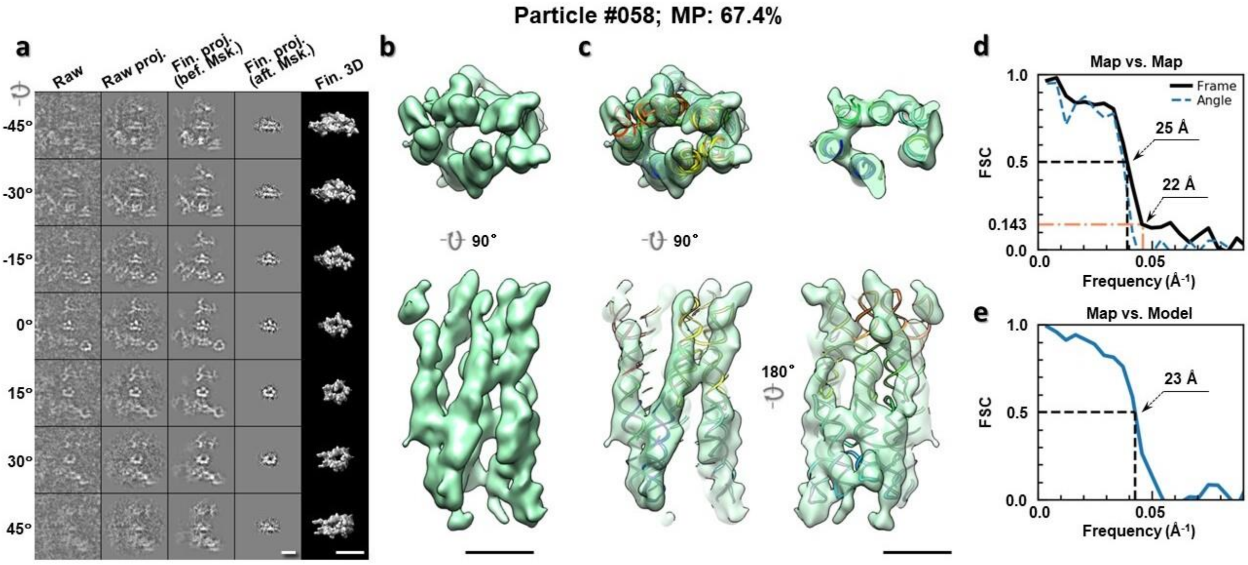

**Supplementary Fig. 28: IPET 3D reconstruction of an individual 6HBC RNA particle #58.** **a,** Seven representative tilt images of a single particle are shown in the first column from the left. The tilt images are aligned to a common center using IPET through iterative refinements. The projections of the raw, intermediate, and final 3D reconstructions at the corresponding angles are displayed in the next four

columns. **b,** Two orthogonal views of the final 3D map; **c,** Overlap of the map with the fitted model; **d,** FSC analyses of the final map resolution using two methods, "map-map FSC", in which each map is reconstructed from one half of the images, based on the even vs. odd indices of frames or tilt angles. respectively, **e,** FSC analysis of map-model, where the model map is generated by the fitted model after being low-pass filtered to 8 Å. The resolutions are assessed based on the frequencies of the FSC curve falls at 0.5. Scale bars represent 10 nm in a, and 5 nm in b and c.

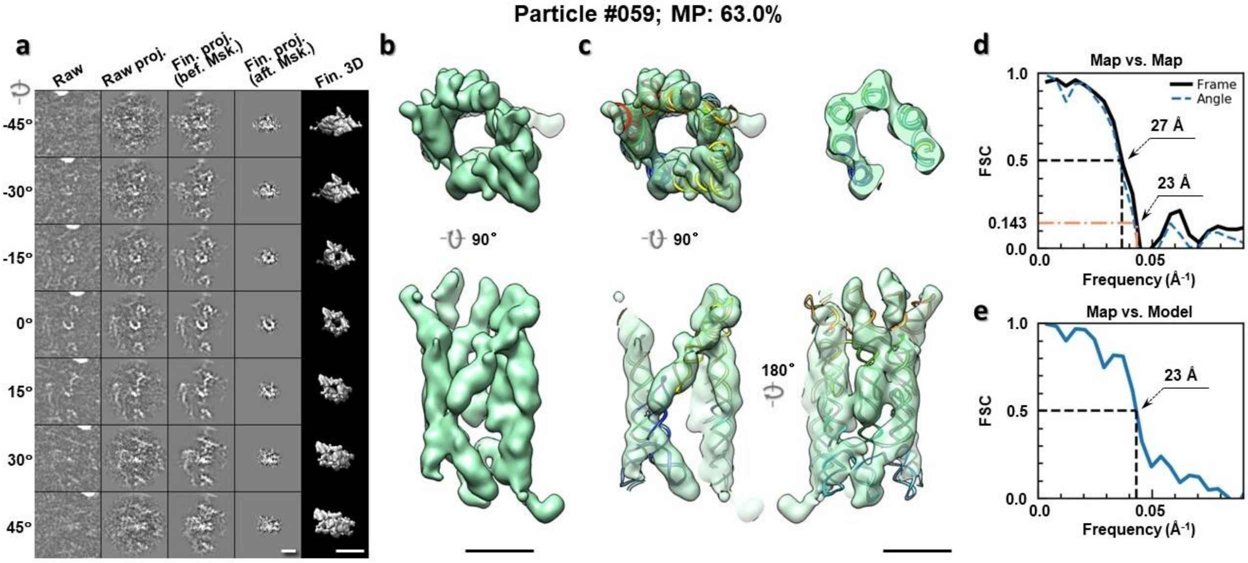

**Supplementary Fig. 30: IPET 3D reconstruction of an individual 6HBC RNA particle #59.** **a,** Seven representative tilt images of a single particle are shown in the first column from the left. The tilt images are aligned to a common center using IPET through iterative refinements. The projections of the raw, intermediate, and final 3D reconstructions at the corresponding angles are displayed in the next four columns. **b,** Two orthogonal views of the final 3D map; **c,** Overlap of the map with the fitted model; **d,** FSC analyses of the final map resolution using two methods, "map-map FSC", in which each map is reconstructed from one half of the images, based on the even vs. odd indices of frames or tilt angles. respectively, **e,** FSC analysis of map-model, where the model map is generated by the fitted model after being low-pass filtered to 8 Å. The resolutions are assessed based on the frequencies of the FSC curve falls at 0.5. Scale bars represent 10 nm in a, and 5 nm in b and c.

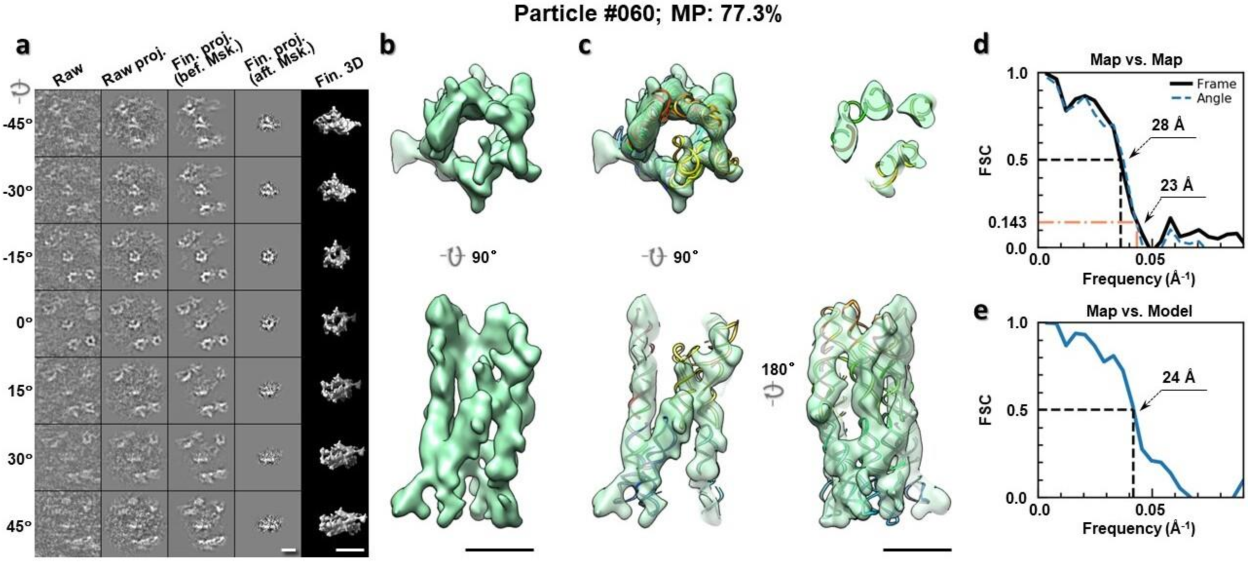

**Supplementary Fig. 31: IPET 3D reconstruction of an individual 6HBC RNA particle #60. a,** Seven representative tilt images of a single particle are shown in the first column from the left. The tilt images are aligned to a common center using IPET through iterative refinements. The projections of the raw, intermediate, and final 3D reconstructions at the corresponding angles are displayed in the next four columns. **b,** Two orthogonal views of the final 3D map; **c,** Overlap of the map with the fitted model; **d,** FSC analyses of the final map resolution using two methods, "map-map FSC", in which each map is reconstructed from one half of the images, based on the even vs. odd indices of frames or tilt angles. respectively, **e,** FSC analysis of map-model, where the model map is generated by the fitted model after being low-pass filtered to 8 Å. The resolutions are assessed based on the frequencies of the FSC curve falls at 0.5. Scale bars represent 10 nm in a, and 5 nm in b and c.

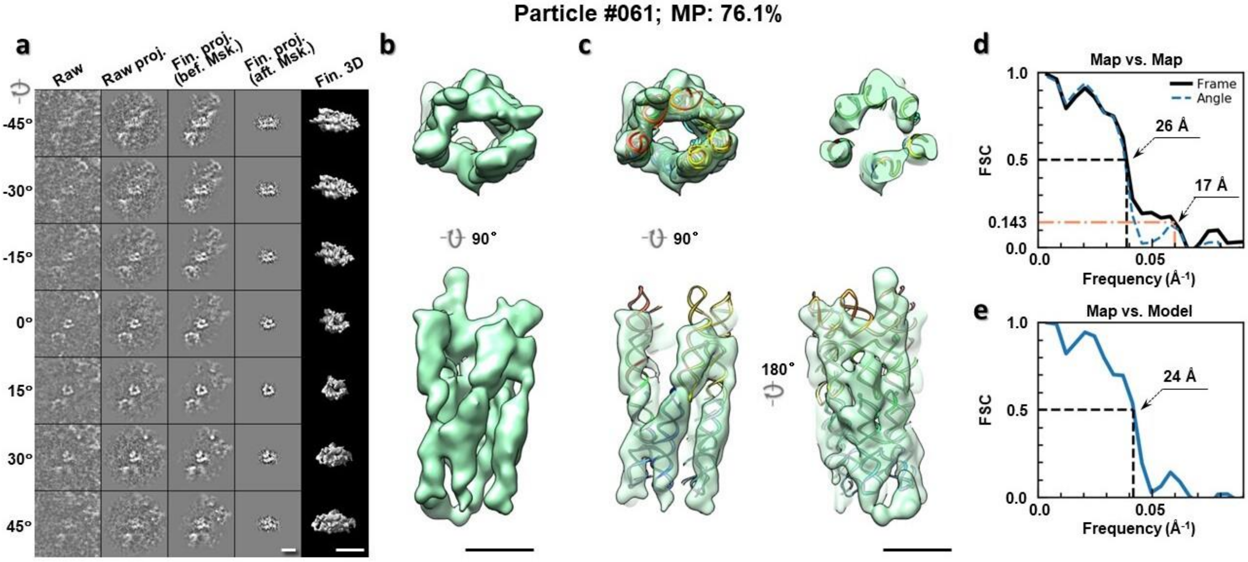

**Supplementary Fig. 32: IPET 3D reconstruction of an individual 6HBC RNA particle #61. a,** Seven representative tilt images of a single particle are shown in the first column from the left. The tilt images are aligned to a common center using IPET through iterative refinements. The projections of the raw, intermediate, and final 3D reconstructions at the corresponding angles are displayed in the next four

columns. **b,** Two orthogonal views of the final 3D map; **c,** Overlap of the map with the fitted model; **d,** FSC analyses of the final map resolution using two methods, "map-map FSC", in which each map is reconstructed from one half of the images, based on the even vs. odd indices of frames or tilt angles. respectively, **e,** FSC analysis of map-model, where the model map is generated by the fitted model after being low-pass filtered to 8 Å. The resolutions are assessed based on the frequencies of the FSC curve falls at 0.5. Scale bars represent 10 nm in a, and 5 nm in b and c.

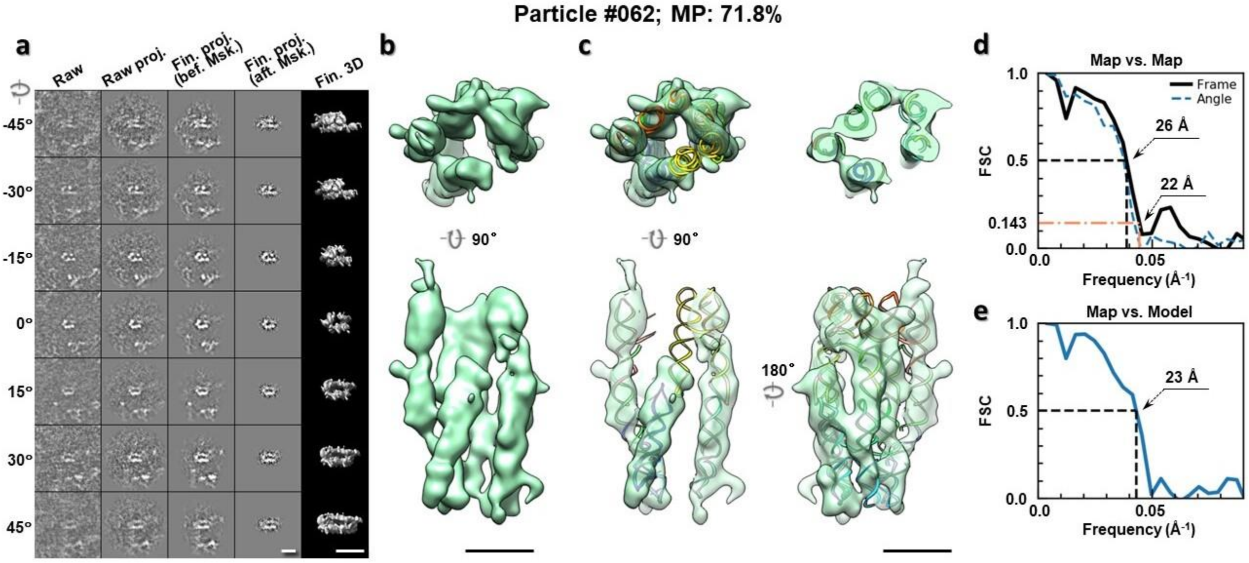

**Supplementary Fig. 33: IPET 3D reconstruction of an individual 6HBC RNA particle #62.** **a,** Seven representative tilt images of a single particle are shown in the first column from the left. The tilt images are aligned to a common center using IPET through iterative refinements. The projections of the raw, intermediate, and final 3D reconstructions at the corresponding angles are displayed in the next four columns. **b,** Two orthogonal views of the final 3D map; **c,** Overlap of the map with the fitted model; **d,** FSC analyses of the final map resolution using two methods, "map-map FSC", in which each map is reconstructed from one half of the images, based on the even vs. odd indices of frames or tilt angles. respectively, **e,** FSC analysis of map-model, where the model map is generated by the fitted model after being low-pass filtered to 8 Å. The resolutions are assessed based on the frequencies of the FSC curve falls at 0.5. Scale bars represent 10 nm in a, and 5 nm in b and c.

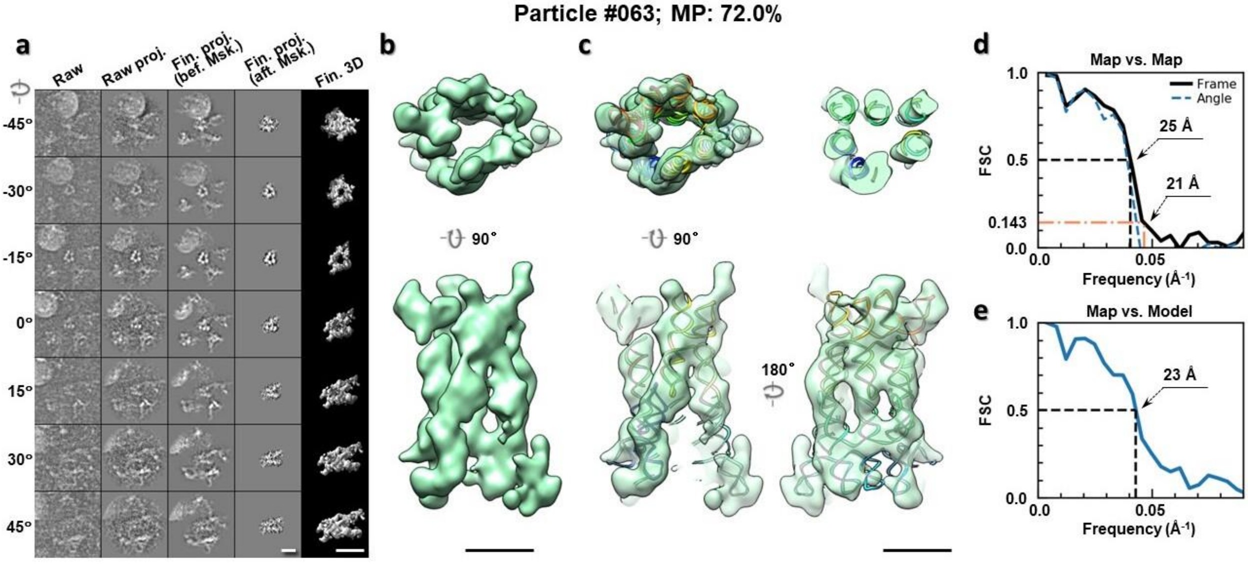

**Supplementary Fig. 34: IPET 3D reconstruction of an individual 6HBC RNA particle #63.** **a,** Seven representative tilt images of a single particle are shown in the first column from the left. The tilt images are aligned to a common center using IPET through iterative refinements. The projections of the raw, intermediate, and final 3D reconstructions at the corresponding angles are displayed in the next four columns. **b,** Two orthogonal views of the final 3D map; **c,** Overlap of the map with the fitted model; **d,** FSC analyses of the final map resolution using two methods, "map-map FSC", in which each map is reconstructed from one half of the images, based on the even vs. odd indices of frames or tilt angles. respectively, **e,** FSC analysis of map-model, where the model map is generated by the fitted model after being low-pass filtered to 8 Å. The resolutions are assessed based on the frequencies of the FSC curve falls at 0.5. Scale bars represent 10 nm in a, and 5 nm in b and c.

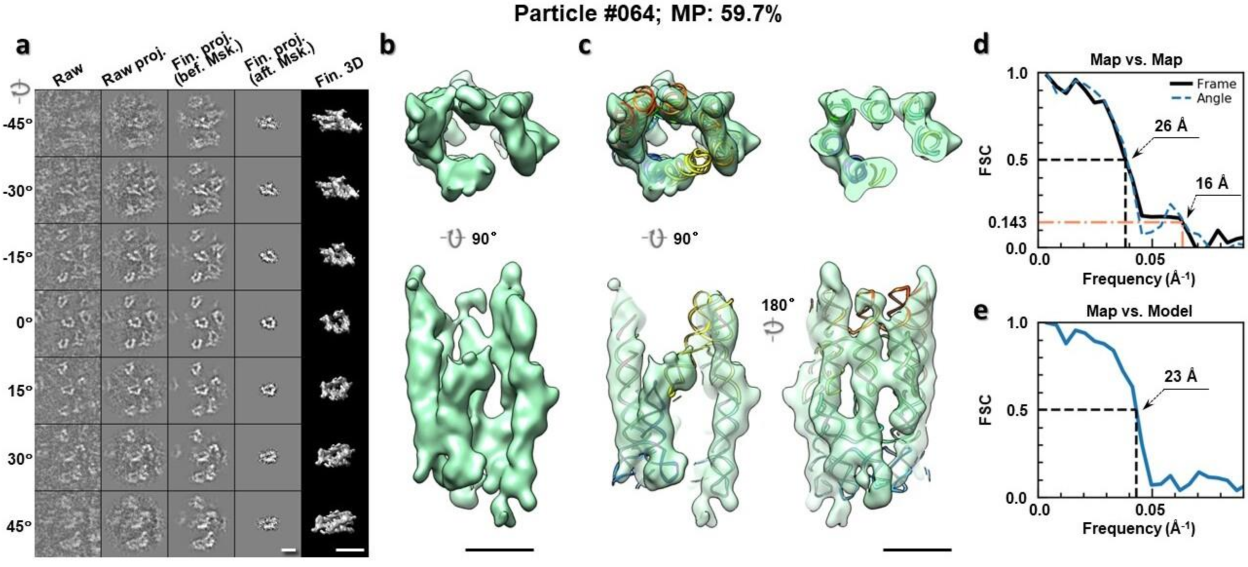

**Supplementary Fig. 35: IPET 3D reconstruction of an individual 6HBC RNA particle #64.** **a,** Seven representative tilt images of a single particle are shown in the first column from the left. The tilt images are aligned to a common center using IPET through iterative refinements. The projections of the raw, intermediate, and final 3D reconstructions at the corresponding angles are displayed in the next four

columns. **b,** Two orthogonal views of the final 3D map; **c,** Overlap of the map with the fitted model; **d,** FSC analyses of the final map resolution using two methods, "map-map FSC", in which each map is reconstructed from one half of the images, based on the even vs. odd indices of frames or tilt angles. respectively, **e,** FSC analysis of map-model, where the model map is generated by the fitted model after being low-pass filtered to 8 Å. The resolutions are assessed based on the frequencies of the FSC curve falls at 0.5. Scale bars represent 10 nm in a, and 5 nm in b and c.

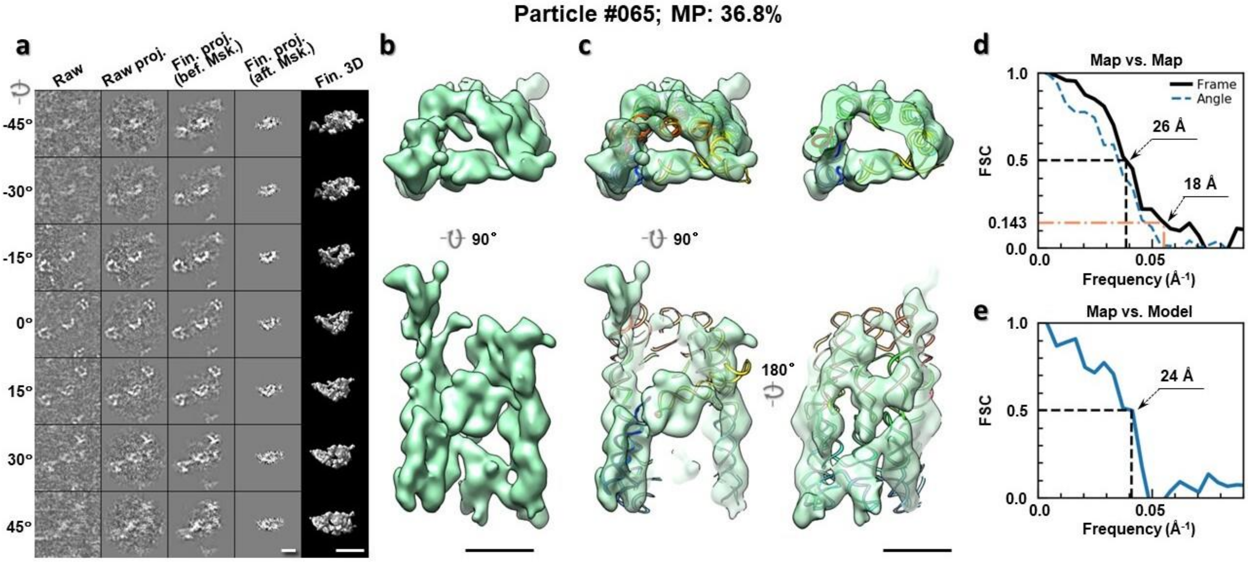

**Supplementary Fig. 36: IPET 3D reconstruction of an individual 6HBC RNA particle #65. a,** Seven representative tilt images of a single particle are shown in the first column from the left. The tilt images are aligned to a common center using IPET through iterative refinements. The projections of the raw, intermediate, and final 3D reconstructions at the corresponding angles are displayed in the next four columns. **b,** Two orthogonal views of the final 3D map; **c,** Overlap of the map with the fitted model; **d,** FSC analyses of the final map resolution using two methods, "map-map FSC", in which each map is reconstructed from one half of the images, based on the even vs. odd indices of frames or tilt angles. respectively, **e,** FSC analysis of map-model, where the model map is generated by the fitted model after being low-pass filtered to 8 Å. The resolutions are assessed based on the frequencies of the FSC curve falls at 0.5. Scale bars represent 10 nm in a, and 5 nm in b and c.

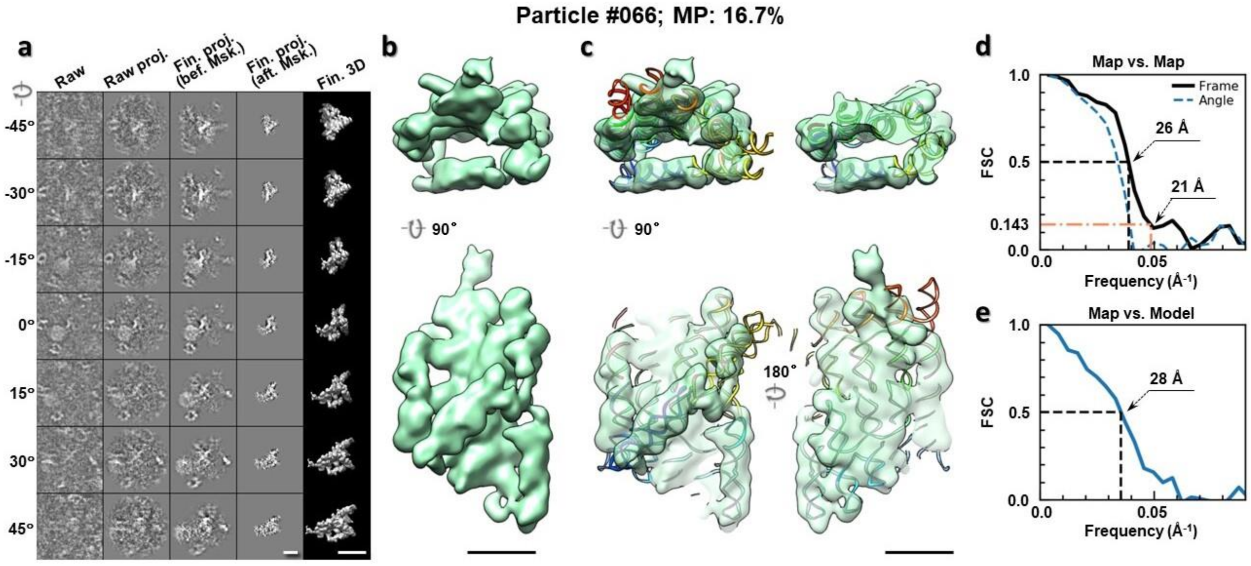

**Supplementary Fig. 37: IPET 3D reconstruction of an individual 6HBC RNA particle #66.** **a,** Seven representative tilt images of a single particle are shown in the first column from the left. The tilt images are aligned to a common center using IPET through iterative refinements. The projections of the raw, intermediate, and final 3D reconstructions at the corresponding angles are displayed in the next four columns. **b,** Two orthogonal views of the final 3D map; **c,** Overlap of the map with the fitted model; **d,** FSC analyses of the final map resolution using two methods, "map-map FSC", in which each map is reconstructed from one half of the images, based on the even vs. odd indices of frames or tilt angles. respectively, **e,** FSC analysis of map-model, where the model map is generated by the fitted model after being low-pass filtered to 8 Å. The resolutions are assessed based on the frequencies of the FSC curve falls at 0.5. Scale bars represent 10 nm in a, and 5 nm in b and c.

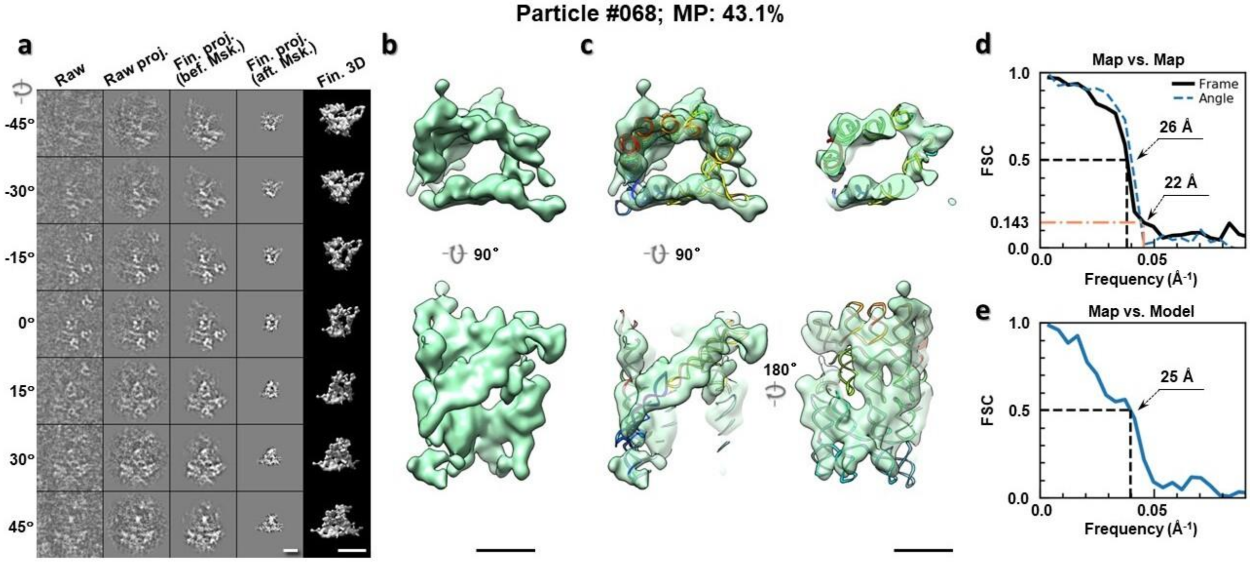

**Supplementary Fig. 38: IPET 3D reconstruction of an individual 6HBC RNA particle #68.** **a,** Seven representative tilt images of a single particle are shown in the first column from the left. The tilt images are aligned to a common center using IPET through iterative refinements. The projections of the raw, intermediate, and final 3D reconstructions at the corresponding angles are displayed in the next four

columns. **b,** Two orthogonal views of the final 3D map; **c,** Overlap of the map with the fitted model; **d,** FSC analyses of the final map resolution using two methods, "map-map FSC", in which each map is reconstructed from one half of the images, based on the even vs. odd indices of frames or tilt angles. respectively, **e,** FSC analysis of map-model, where the model map is generated by the fitted model after being low-pass filtered to 8 Å. The resolutions are assessed based on the frequencies of the FSC curve falls at 0.5. Scale bars represent 10 nm in a, and 5 nm in b and c.

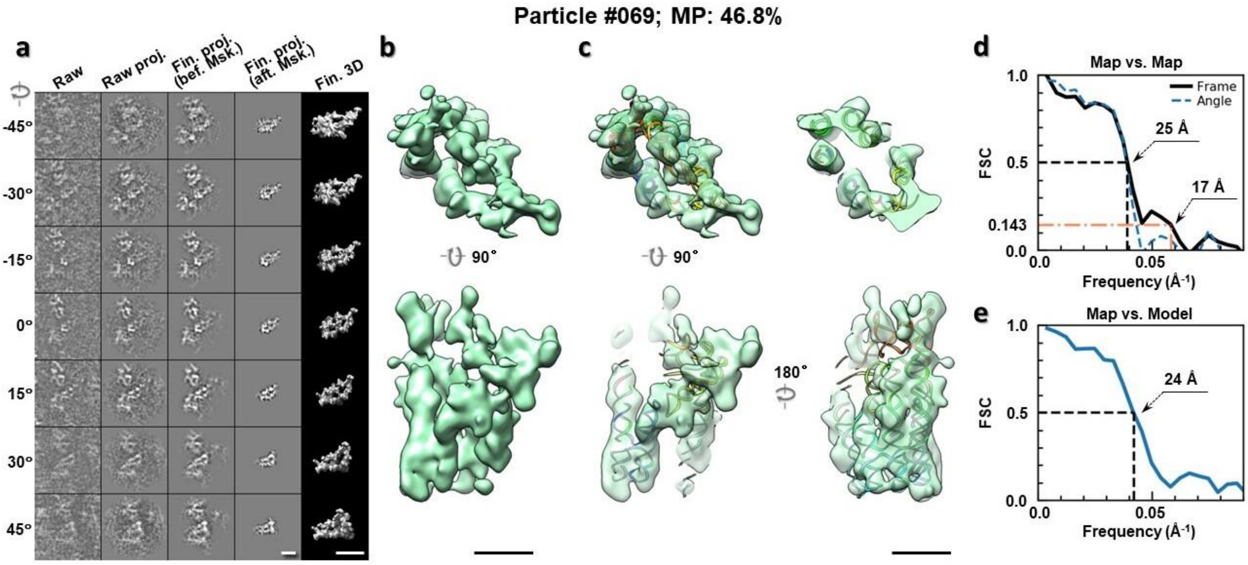

**Supplementary Fig. 39: IPET 3D reconstruction of an individual 6HBC RNA particle #69.** **a,** Seven representative tilt images of a single particle are shown in the first column from the left. The tilt images are aligned to a common center using IPET through iterative refinements. The projections of the raw, intermediate, and final 3D reconstructions at the corresponding angles are displayed in the next four columns. **b,** Two orthogonal views of the final 3D map; **c,** Overlap of the map with the fitted model; **d,** FSC analyses of the final map resolution using two methods, "map-map FSC", in which each map is reconstructed from one half of the images, based on the even vs. odd indices of frames or tilt angles. respectively, **e,** FSC analysis of map-model, where the model map is generated by the fitted model after being low-pass filtered to 8 Å. The resolutions are assessed based on the frequencies of the FSC curve falls at 0.5. Scale bars represent 10 nm in a, and 5 nm in b and c.

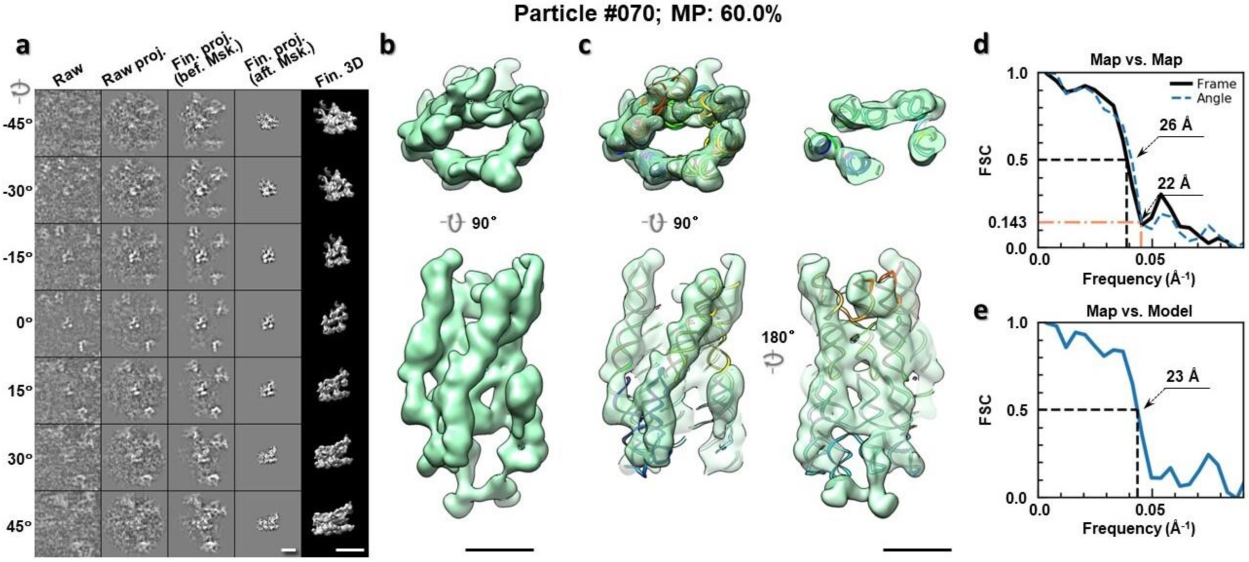

**Supplementary Fig. 40: IPET 3D reconstruction of an individual 6HBC RNA particle #70.** **a,** Seven representative tilt images of a single particle are shown in the first column from the left. The tilt images are aligned to a common center using IPET through iterative refinements. The projections of the raw, intermediate, and final 3D reconstructions at the corresponding angles are displayed in the next four columns. **b,** Two orthogonal views of the final 3D map; **c,** Overlap of the map with the fitted model; **d,** FSC analyses of the final map resolution using two methods, "map-map FSC", in which each map is reconstructed from one half of the images, based on the even vs. odd indices of frames or tilt angles. respectively, **e,** FSC analysis of map-model, where the model map is generated by the fitted model after being low-pass filtered to 8 Å. The resolutions are assessed based on the frequencies of the FSC curve falls at 0.5. Scale bars represent 10 nm in a, and 5 nm in b and c.

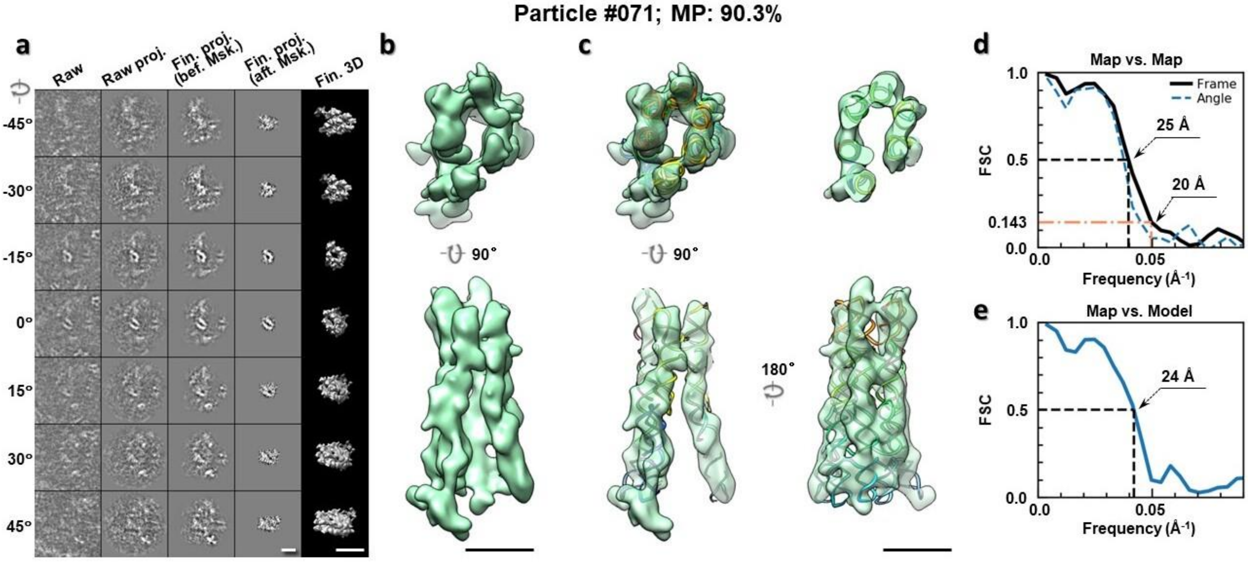

**Supplementary Fig. 41: IPET 3D reconstruction of an individual 6HBC RNA particle #71.** **a,** Seven representative tilt images of a single particle are shown in the first column from the left. The tilt images are aligned to a common center using IPET through iterative refinements. The projections of the raw, intermediate, and final 3D reconstructions at the corresponding angles are displayed in the next four

columns. **b,** Two orthogonal views of the final 3D map; **c,** Overlap of the map with the fitted model; **d,** FSC analyses of the final map resolution using two methods, "map-map FSC", in which each map is reconstructed from one half of the images, based on the even vs. odd indices of frames or tilt angles. respectively, **e,** FSC analysis of map-model, where the model map is generated by the fitted model after being low-pass filtered to 8 Å. The resolutions are assessed based on the frequencies of the FSC curve falls at 0.5. Scale bars represent 10 nm in a, and 5 nm in b and c.

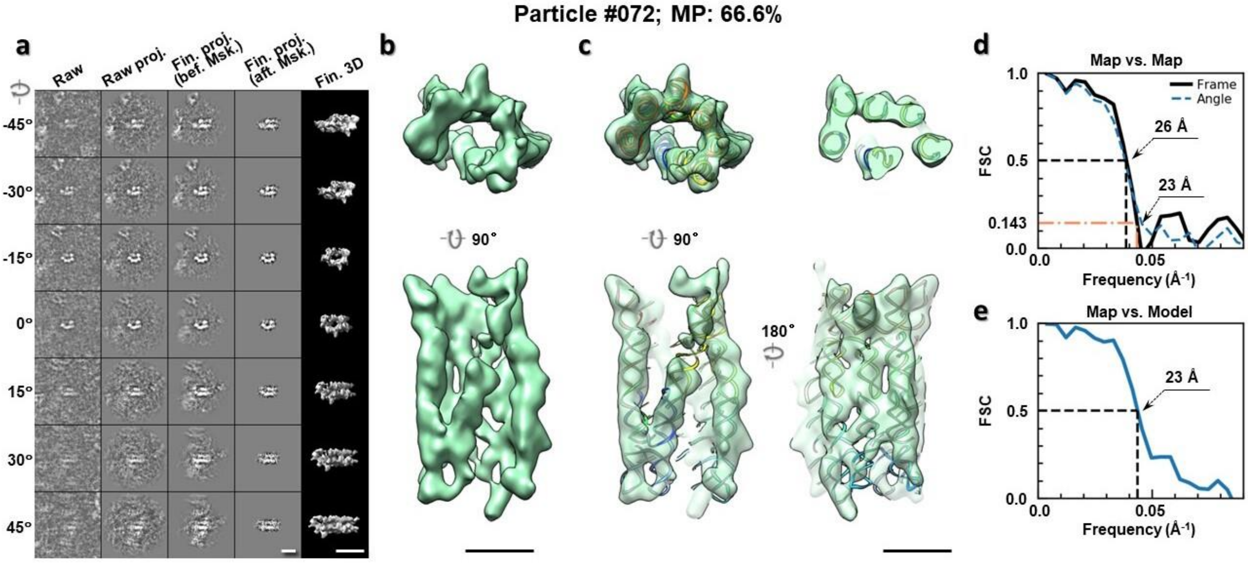

**Supplementary Fig. 42: IPET 3D reconstruction of an individual 6HBC RNA particle #72.** **a,** Seven representative tilt images of a single particle are shown in the first column from the left. The tilt images are aligned to a common center using IPET through iterative refinements. The projections of the raw, intermediate, and final 3D reconstructions at the corresponding angles are displayed in the next four columns. **b,** Two orthogonal views of the final 3D map; **c,** Overlap of the map with the fitted model; **d,** FSC analyses of the final map resolution using two methods, "map-map FSC", in which each map is reconstructed from one half of the images, based on the even vs. odd indices of frames or tilt angles. respectively, **e,** FSC analysis of map-model, where the model map is generated by the fitted model after being low-pass filtered to 8 Å. The resolutions are assessed based on the frequencies of the FSC curve falls at 0.5. Scale bars represent 10 nm in a, and 5 nm in b and c.

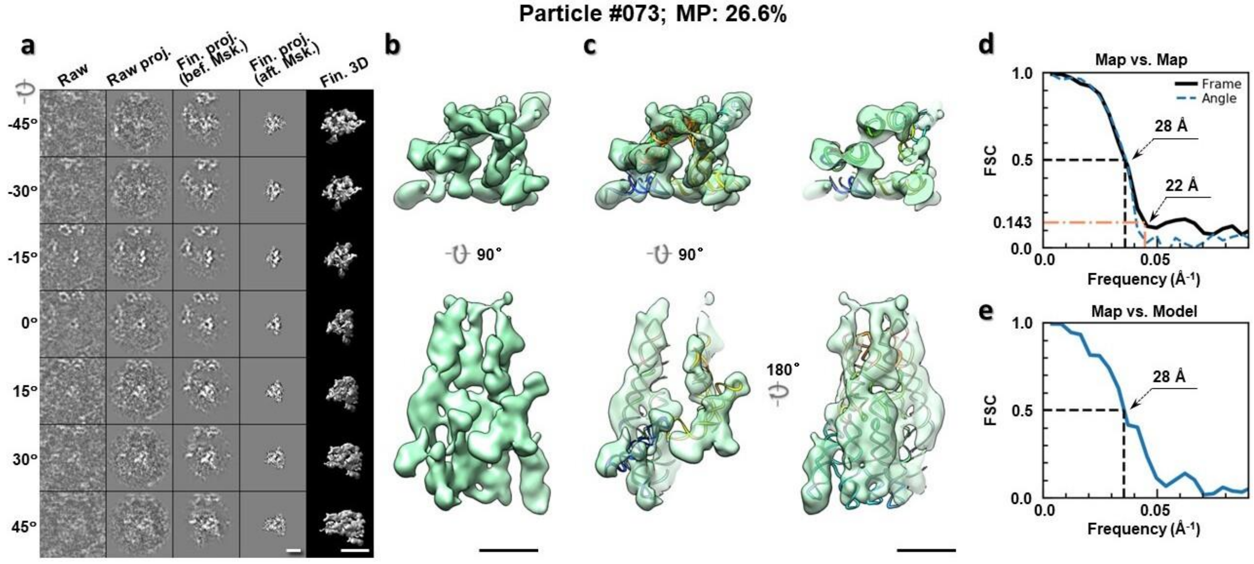

**Supplementary Fig. 43: IPET 3D reconstruction of an individual 6HBC RNA particle #73.** **a,** Seven representative tilt images of a single particle are shown in the first column from the left. The tilt images are aligned to a common center using IPET through iterative refinements. The projections of the raw, intermediate, and final 3D reconstructions at the corresponding angles are displayed in the next four columns. **b,** Two orthogonal views of the final 3D map; **c,** Overlap of the map with the fitted model; **d,** FSC analyses of the final map resolution using two methods, "map-map FSC", in which each map is reconstructed from one half of the images, based on the even vs. odd indices of frames or tilt angles. respectively, **e,** FSC analysis of map-model, where the model map is generated by the fitted model after being low-pass filtered to 8 Å. The resolutions are assessed based on the frequencies of the FSC curve falls at 0.5. Scale bars represent 10 nm in a, and 5 nm in b and c.

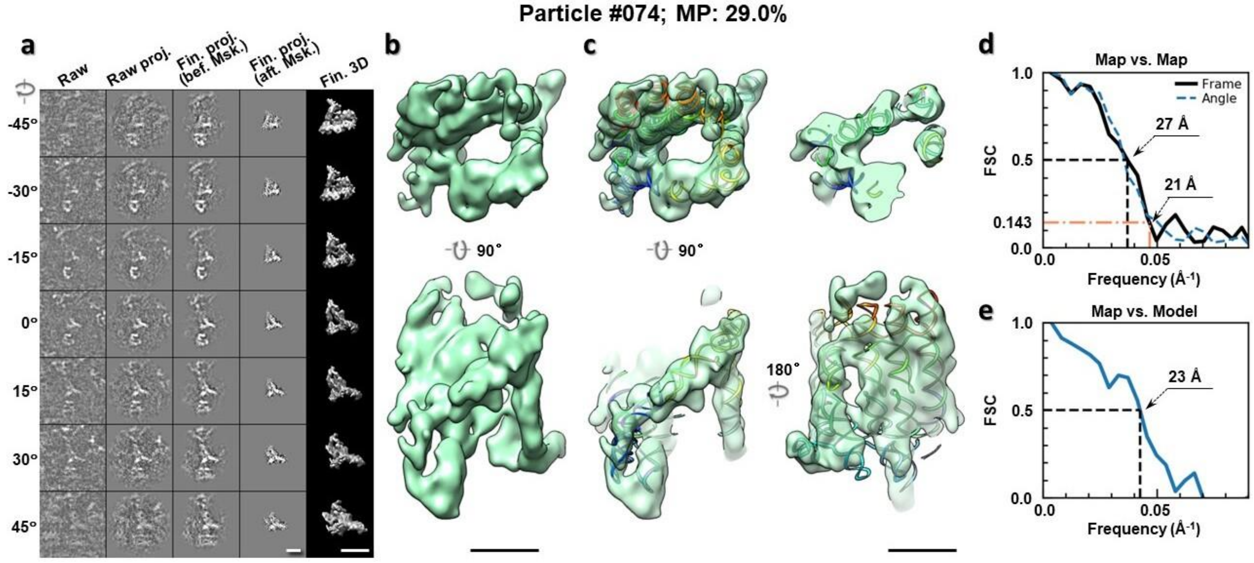

**Supplementary Fig. 44: IPET 3D reconstruction of an individual 6HBC RNA particle #74.** **a,** Seven representative tilt images of a single particle are shown in the first column from the left. The tilt images are aligned to a common center using IPET through iterative refinements. The projections of the raw, intermediate, and final 3D reconstructions at the corresponding angles are displayed in the next four

columns. **b,** Two orthogonal views of the final 3D map; **c,** Overlap of the map with the fitted model; **d,** FSC analyses of the final map resolution using two methods, "map-map FSC", in which each map is reconstructed from one half of the images, based on the even vs. odd indices of frames or tilt angles. respectively, **e,** FSC analysis of map-model, where the model map is generated by the fitted model after being low-pass filtered to 8 Å. The resolutions are assessed based on the frequencies of the FSC curve falls at 0.5. Scale bars represent 10 nm in a, and 5 nm in b and c.

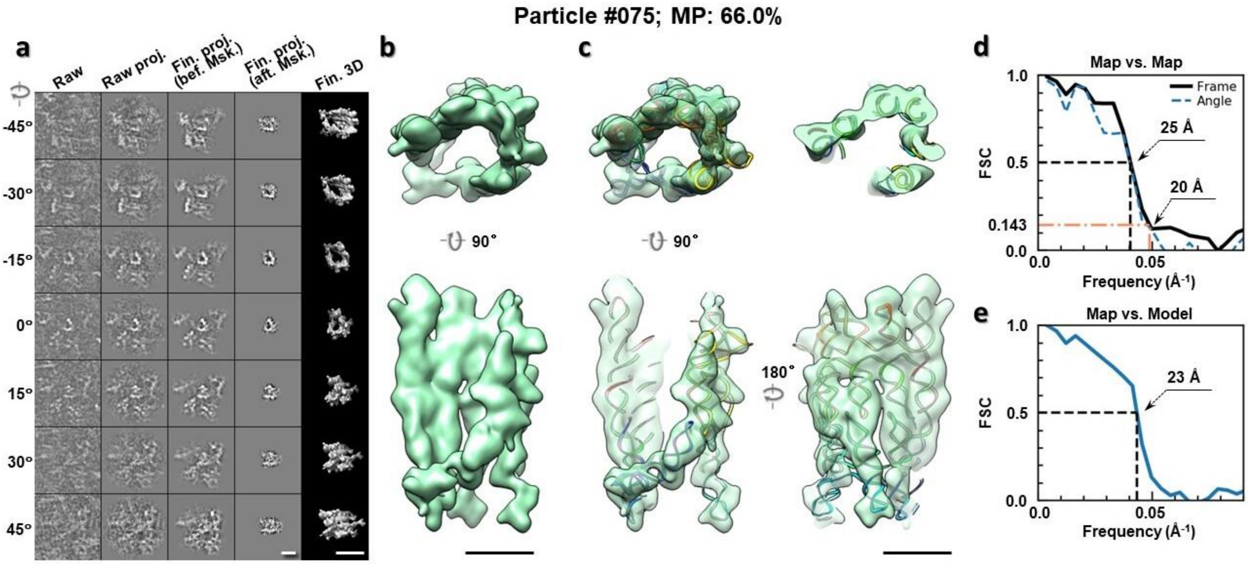

**Supplementary Fig. 45: IPET 3D reconstruction of an individual 6HBC RNA particle #75. a,** Seven representative tilt images of a single particle are shown in the first column from the left. The tilt images are aligned to a common center using IPET through iterative refinements. The projections of the raw, intermediate, and final 3D reconstructions at the corresponding angles are displayed in the next four columns. **b,** Two orthogonal views of the final 3D map; **c,** Overlap of the map with the fitted model; **d,** FSC analyses of the final map resolution using two methods, "map-map FSC", in which each map is reconstructed from one half of the images, based on the even vs. odd indices of frames or tilt angles. respectively, **e,** FSC analysis of map-model, where the model map is generated by the fitted model after being low-pass filtered to 8 Å. The resolutions are assessed based on the frequencies of the FSC curve falls at 0.5. Scale bars represent 10 nm in a, and 5 nm in b and c.

Particle #076; MP: 32.8%

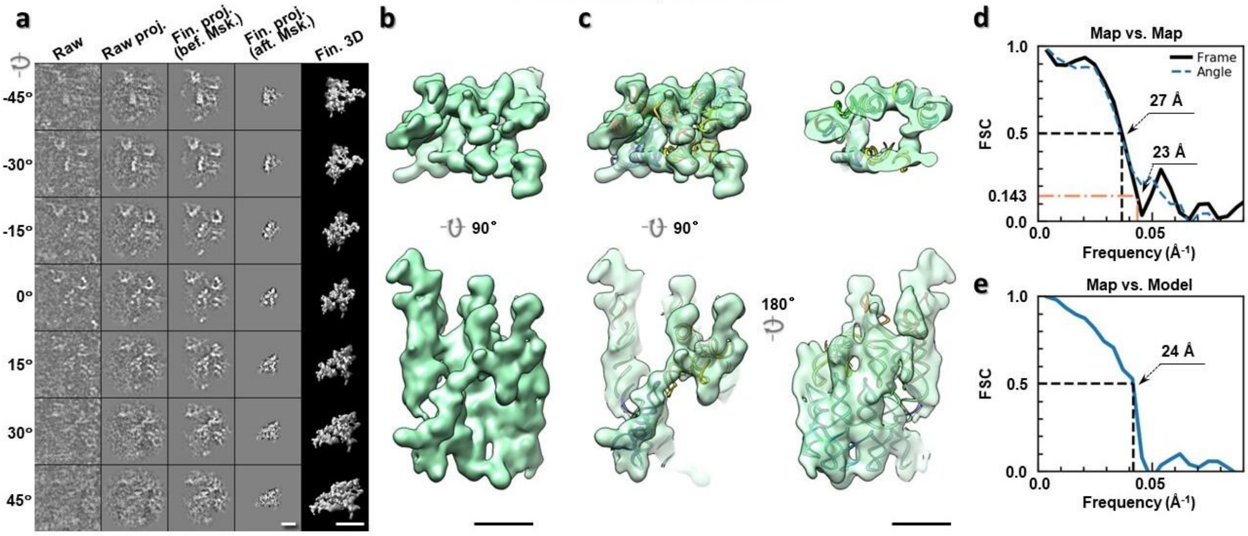

**Supplementary Fig. 46: IPET 3D reconstruction of an individual 6HBC RNA particle #76. a,** Seven representative tilt images of a single particle are shown in the first column from the left. The tilt images are aligned to a common center using IPET through iterative refinements. The projections of the raw, intermediate, and final 3D reconstructions at the corresponding angles are displayed in the next four columns. **b,** Two orthogonal views of the final 3D map; **c,** Overlap of the map with the fitted model; **d,** FSC analyses of the final map resolution using two methods, "map-map FSC", in which each map is reconstructed from one half of the images, based on the even vs. odd indices of frames or tilt angles. respectively, **e,** FSC analysis of map-model, where the model map is generated by the fitted model after being low-pass filtered to 8 Å. The resolutions are assessed based on the frequencies of the FSC curve falls at 0.5. Scale bars represent 10 nm in a, and 5 nm in b and c.

Particle #077; MP: 28.2%

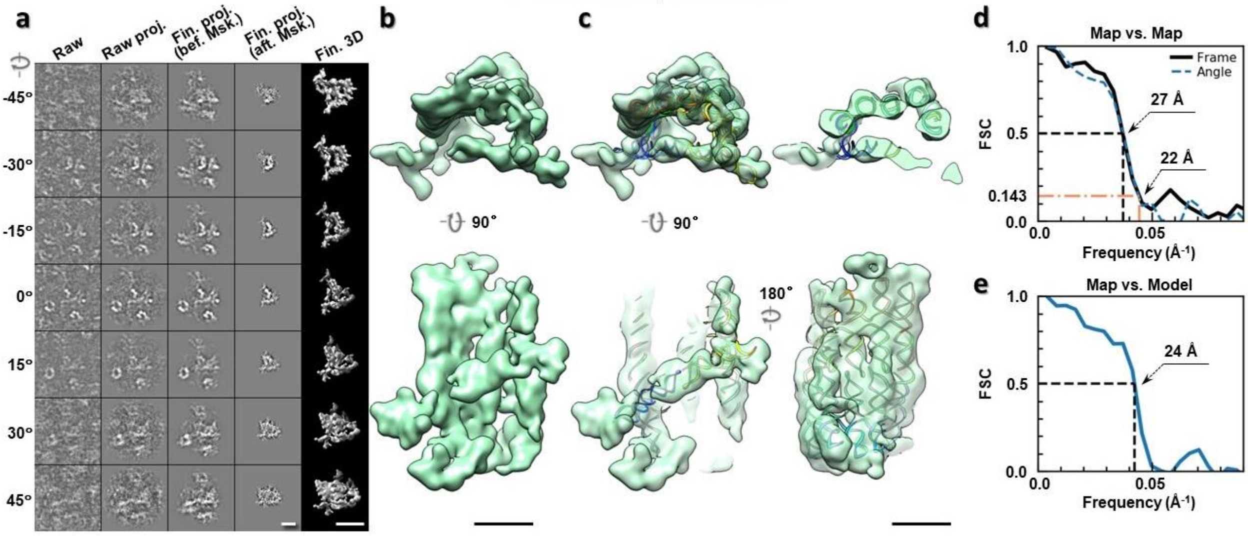

**Supplementary Fig. 47: IPET 3D reconstruction of an individual 6HBC RNA particle #77. a,** Seven representative tilt images of a single particle are shown in the first column from the left. The tilt images are aligned to a common center using IPET through iterative refinements. The projections of the raw, intermediate, and final 3D reconstructions at the corresponding angles are displayed in the next four

columns. **b,** Two orthogonal views of the final 3D map; **c,** Overlap of the map with the fitted model; **d,** FSC analyses of the final map resolution using two methods, "map-map FSC", in which each map is reconstructed from one half of the images, based on the even vs. odd indices of frames or tilt angles. respectively, **e,** FSC analysis of map-model, where the model map is generated by the fitted model after being low-pass filtered to 8 Å. The resolutions are assessed based on the frequencies of the FSC curve falls at 0.5. Scale bars represent 10 nm in a, and 5 nm in b and c.

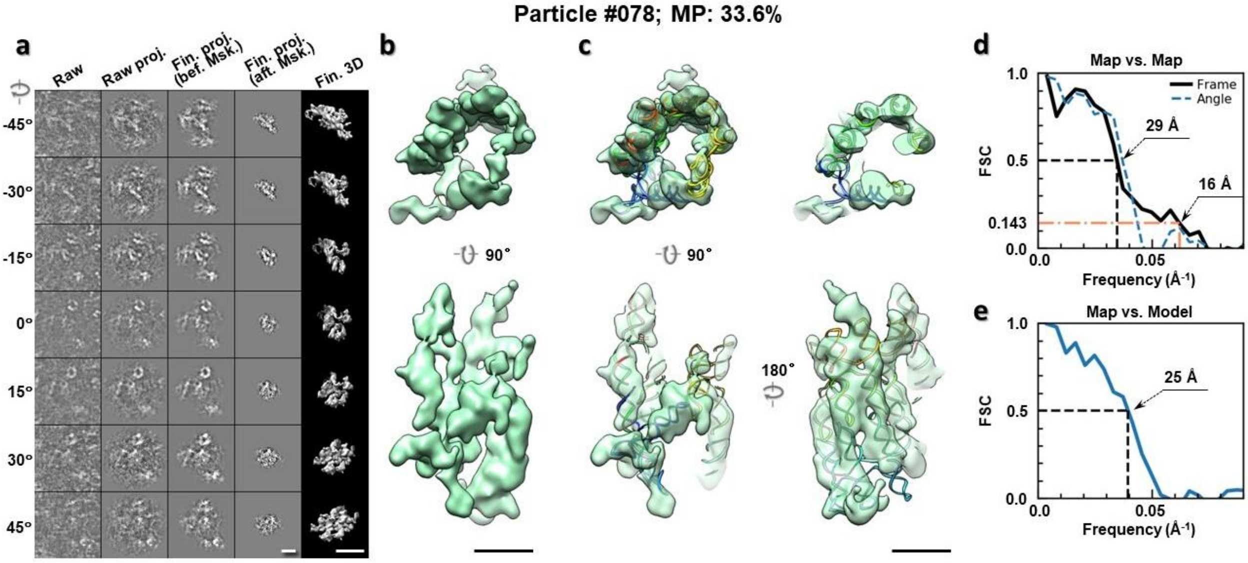

**Supplementary Fig. 48: IPET 3D reconstruction of an individual 6HBC RNA particle #78.** **a,** Seven representative tilt images of a single particle are shown in the first column from the left. The tilt images are aligned to a common center using IPET through iterative refinements. The projections of the raw, intermediate, and final 3D reconstructions at the corresponding angles are displayed in the next four columns. **b,** Two orthogonal views of the final 3D map; **c,** Overlap of the map with the fitted model; **d,** FSC analyses of the final map resolution using two methods, "map-map FSC", in which each map is reconstructed from one half of the images, based on the even vs. odd indices of frames or tilt angles. respectively, **e,** FSC analysis of map-model, where the model map is generated by the fitted model after being low-pass filtered to 8 Å. The resolutions are assessed based on the frequencies of the FSC curve falls at 0.5. Scale bars represent 10 nm in a, and 5 nm in b and c.

Particle #079; MP: 68.8%

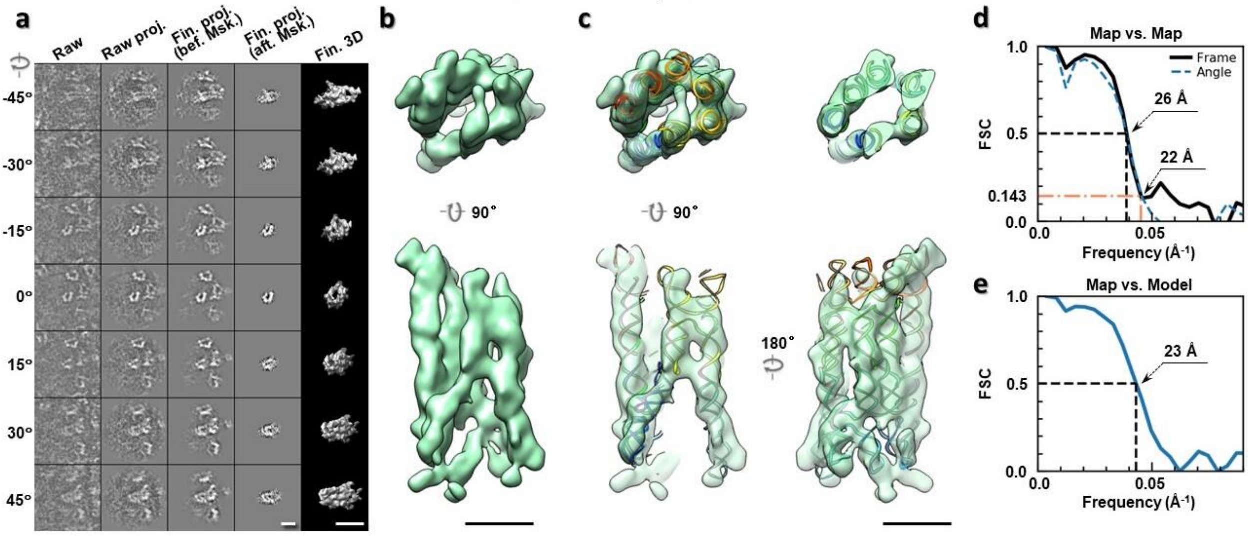

**Supplementary Fig. 49: IPET 3D reconstruction of an individual 6HBC RNA particle #79.** **a,** Seven representative tilt images of a single particle are shown in the first column from the left. The tilt images are aligned to a common center using IPET through iterative refinements. The projections of the raw, intermediate, and final 3D reconstructions at the corresponding angles are displayed in the next four columns. **b,** Two orthogonal views of the final 3D map; **c,** Overlap of the map with the fitted model; **d,** FSC analyses of the final map resolution using two methods, "map-map FSC", in which each map is reconstructed from one half of the images, based on the even vs. odd indices of frames or tilt angles. respectively, **e,** FSC analysis of map-model, where the model map is generated by the fitted model after being low-pass filtered to 8 Å. The resolutions are assessed based on the frequencies of the FSC curve falls at 0.5. Scale bars represent 10 nm in a, and 5 nm in b and c.

Particle #080; MP: 43.4%

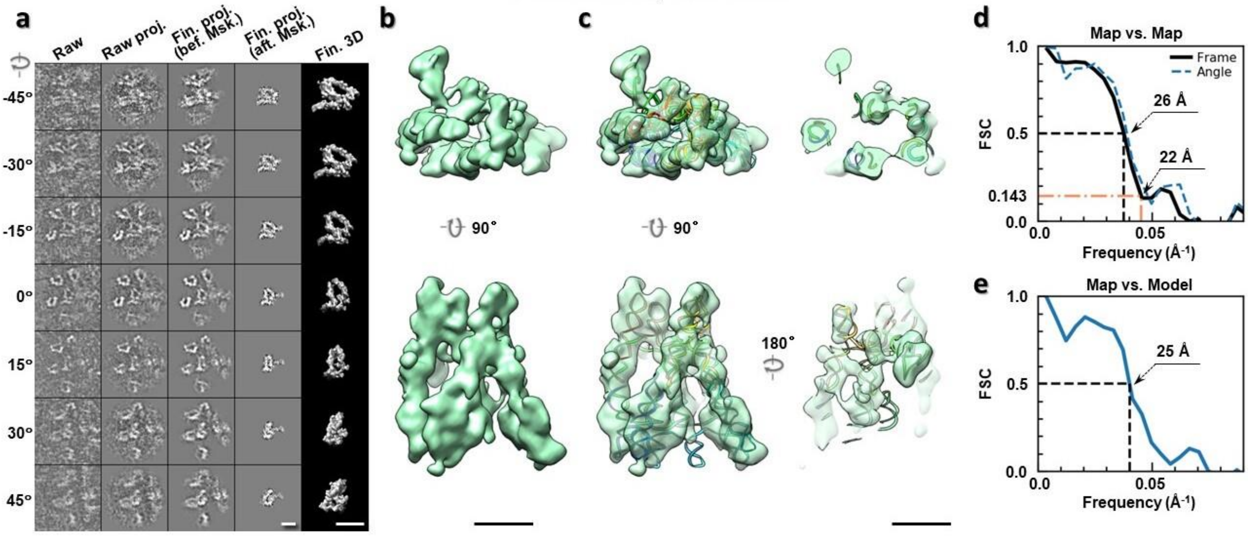

**Supplementary Fig. 50: IPET 3D reconstruction of an individual 6HBC RNA particle #80.** **a,** Seven representative tilt images of a single particle are shown in the first column from the left. The tilt images are aligned to a common center using IPET through iterative refinements. The projections of the raw, intermediate, and final 3D reconstructions at the corresponding angles are displayed in the next four

columns. **b,** Two orthogonal views of the final 3D map; **c,** Overlap of the map with the fitted model; **d,** FSC analyses of the final map resolution using two methods, "map-map FSC", in which each map is reconstructed from one half of the images, based on the even vs. odd indices of frames or tilt angles. respectively, **e,** FSC analysis of map-model, where the model map is generated by the fitted model after being low-pass filtered to 8 Å. The resolutions are assessed based on the frequencies of the FSC curve falls at 0.5. Scale bars represent 10 nm in a, and 5 nm in b and c.

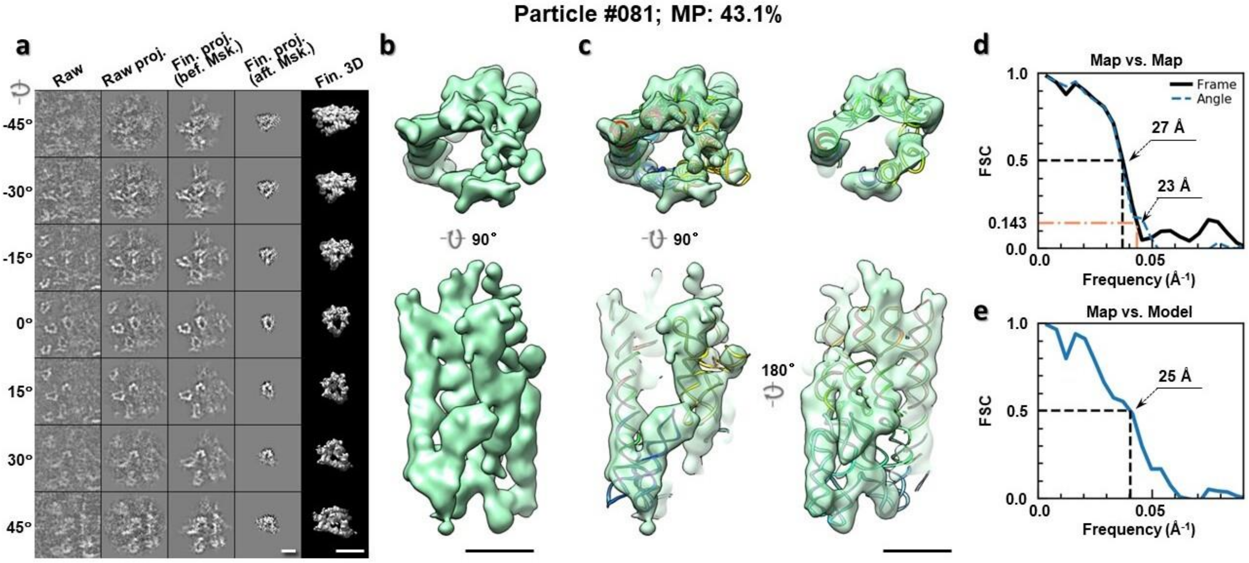

**Supplementary Fig. 51: IPET 3D reconstruction of an individual 6HBC RNA particle #81.** **a,** Seven representative tilt images of a single particle are shown in the first column from the left. The tilt images are aligned to a common center using IPET through iterative refinements. The projections of the raw, intermediate, and final 3D reconstructions at the corresponding angles are displayed in the next four columns. **b,** Two orthogonal views of the final 3D map; **c,** Overlap of the map with the fitted model; **d,** FSC analyses of the final map resolution using two methods, "map-map FSC", in which each map is reconstructed from one half of the images, based on the even vs. odd indices of frames or tilt angles. respectively, **e,** FSC analysis of map-model, where the model map is generated by the fitted model after being low-pass filtered to 8 Å. The resolutions are assessed based on the frequencies of the FSC curve falls at 0.5. Scale bars represent 10 nm in a, and 5 nm in b and c.

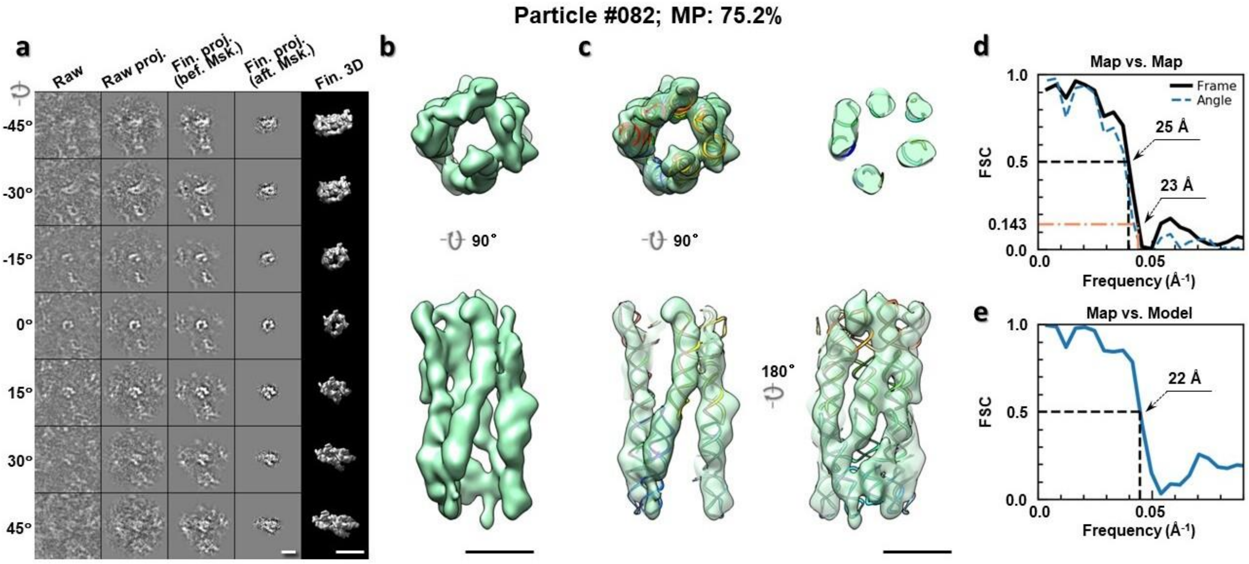

**Supplementary Fig. 52: IPET 3D reconstruction of an individual 6HBC RNA particle #82. a,** Seven representative tilt images of a single particle are shown in the first column from the left. The tilt images are aligned to a common center using IPET through iterative refinements. The projections of the raw, intermediate, and final 3D reconstructions at the corresponding angles are displayed in the next four columns. **b,** Two orthogonal views of the final 3D map; **c,** Overlap of the map with the fitted model; **d,** FSC analyses of the final map resolution using two methods, "map-map FSC", in which each map is reconstructed from one half of the images, based on the even vs. odd indices of frames or tilt angles. respectively, **e,** FSC analysis of map-model, where the model map is generated by the fitted model after being low-pass filtered to 8 Å. The resolutions are assessed based on the frequencies of the FSC curve falls at 0.5. Scale bars represent 10 nm in a, and 5 nm in b and c.

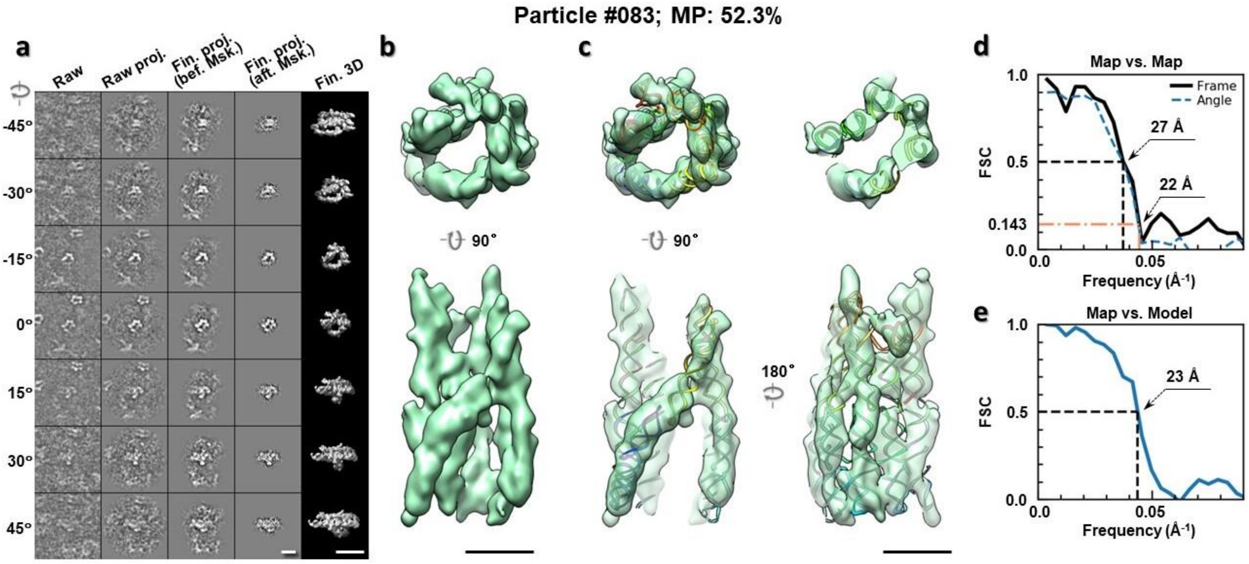

**Supplementary Fig. 53: IPET 3D reconstruction of an individual 6HBC RNA particle #83. a,** Seven representative tilt images of a single particle are shown in the first column from the left. The tilt images are aligned to a common center using IPET through iterative refinements. The projections of the raw, intermediate, and final 3D reconstructions at the corresponding angles are displayed in the next four

columns. **b,** Two orthogonal views of the final 3D map; **c,** Overlap of the map with the fitted model; **d,** FSC analyses of the final map resolution using two methods, "map-map FSC", in which each map is reconstructed from one half of the images, based on the even vs. odd indices of frames or tilt angles. respectively, **e,** FSC analysis of map-model, where the model map is generated by the fitted model after being low-pass filtered to 8 Å. The resolutions are assessed based on the frequencies of the FSC curve falls at 0.5. Scale bars represent 10 nm in a, and 5 nm in b and c.

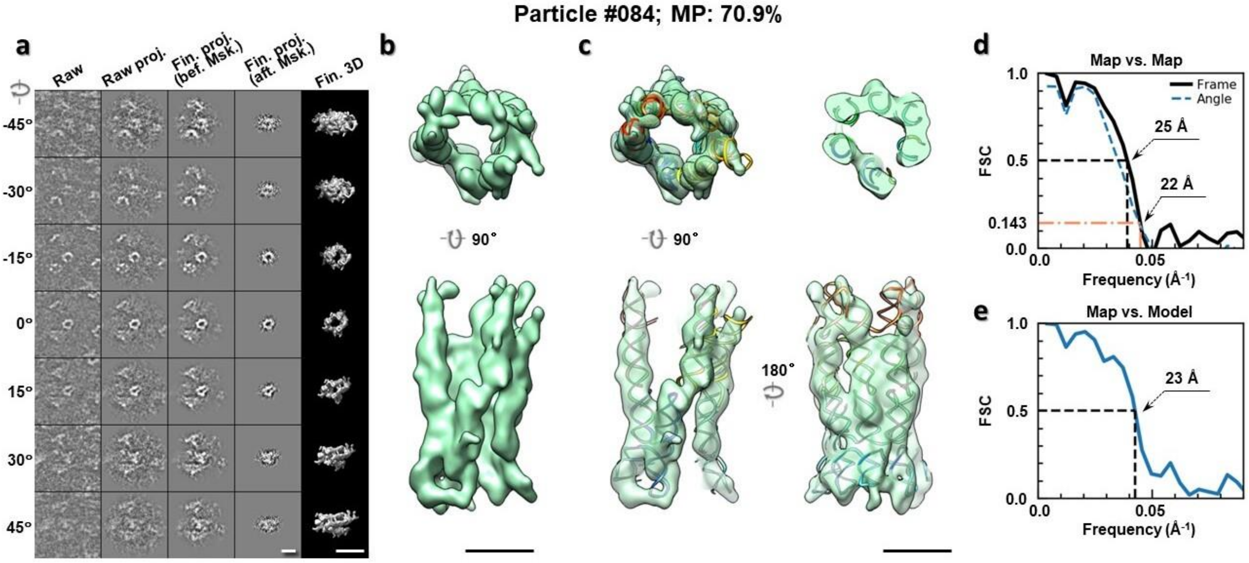

**Supplementary Fig. 54: IPET 3D reconstruction of an individual 6HBC RNA particle #84. a,** Seven representative tilt images of a single particle are shown in the first column from the left. The tilt images are aligned to a common center using IPET through iterative refinements. The projections of the raw, intermediate, and final 3D reconstructions at the corresponding angles are displayed in the next four columns. **b,** Two orthogonal views of the final 3D map; **c,** Overlap of the map with the fitted model; **d,** FSC analyses of the final map resolution using two methods, "map-map FSC", in which each map is reconstructed from one half of the images, based on the even vs. odd indices of frames or tilt angles. respectively, **e,** FSC analysis of map-model, where the model map is generated by the fitted model after being low-pass filtered to 8 Å. The resolutions are assessed based on the frequencies of the FSC curve falls at 0.5. Scale bars represent 10 nm in a, and 5 nm in b and c.

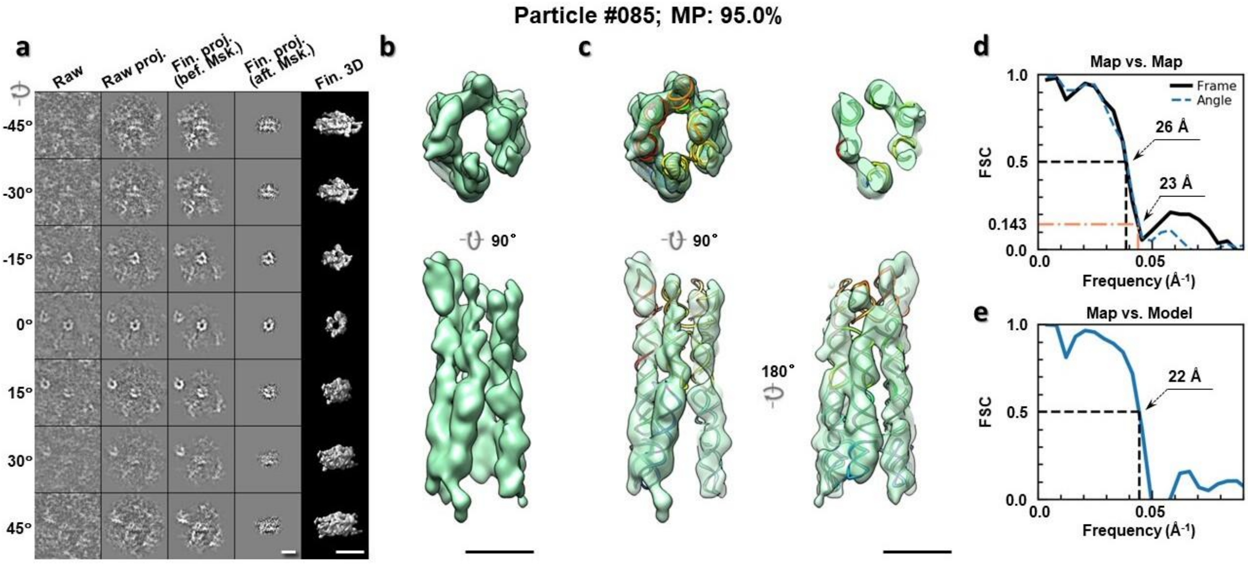

**Supplementary Fig. 55: IPET 3D reconstruction of an individual 6HBC RNA particle #85. a,** Seven representative tilt images of a single particle are shown in the first column from the left. The tilt images are aligned to a common center using IPET through iterative refinements. The projections of the raw, intermediate, and final 3D reconstructions at the corresponding angles are displayed in the next four columns. **b,** Two orthogonal views of the final 3D map; **c,** Overlap of the map with the fitted model; **d,** FSC analyses of the final map resolution using two methods, "map-map FSC", in which each map is reconstructed from one half of the images, based on the even vs. odd indices of frames or tilt angles. respectively, **e,** FSC analysis of map-model, where the model map is generated by the fitted model after being low-pass filtered to 8 Å. The resolutions are assessed based on the frequencies of the FSC curve falls at 0.5. Scale bars represent 10 nm in a, and 5 nm in b and c.

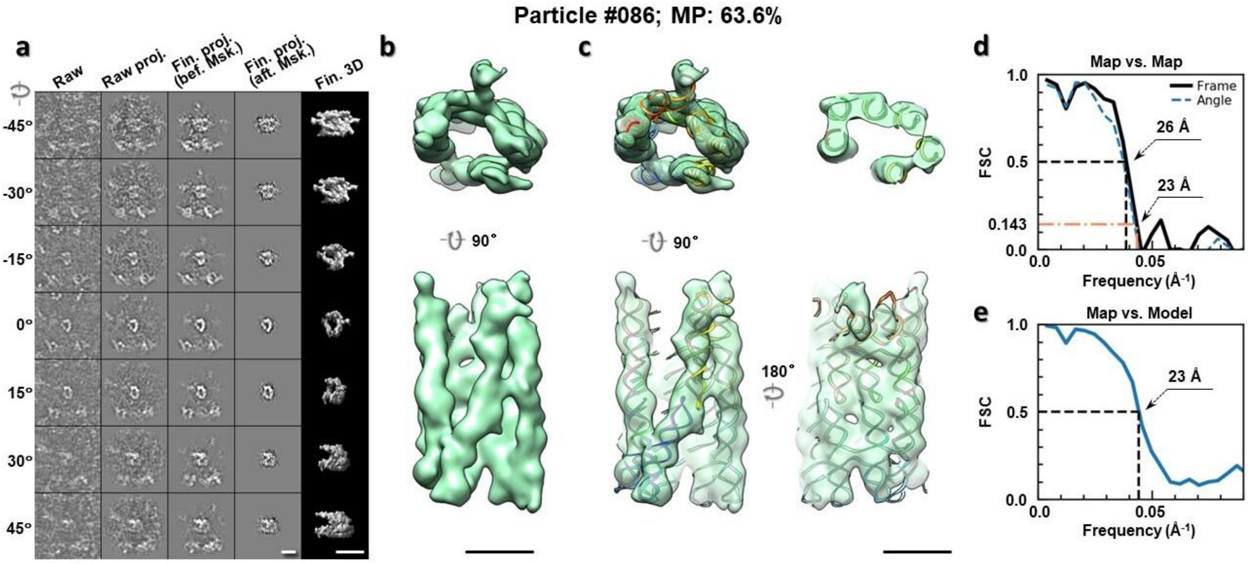

**Supplementary Fig. 56: IPET 3D reconstruction of an individual 6HBC RNA particle #86. a,** Seven representative tilt images of a single particle are shown in the first column from the left. The tilt images are aligned to a common center using IPET through iterative refinements. The projections of the raw, intermediate, and final 3D reconstructions at the corresponding angles are displayed in the next four

columns. **b,** Two orthogonal views of the final 3D map; **c,** Overlap of the map with the fitted model; **d,** FSC analyses of the final map resolution using two methods, "map-map FSC", in which each map is reconstructed from one half of the images, based on the even vs. odd indices of frames or tilt angles. respectively, **e,** FSC analysis of map-model, where the model map is generated by the fitted model after being low-pass filtered to 8 Å. The resolutions are assessed based on the frequencies of the FSC curve falls at 0.5. Scale bars represent 10 nm in a, and 5 nm in b and c.

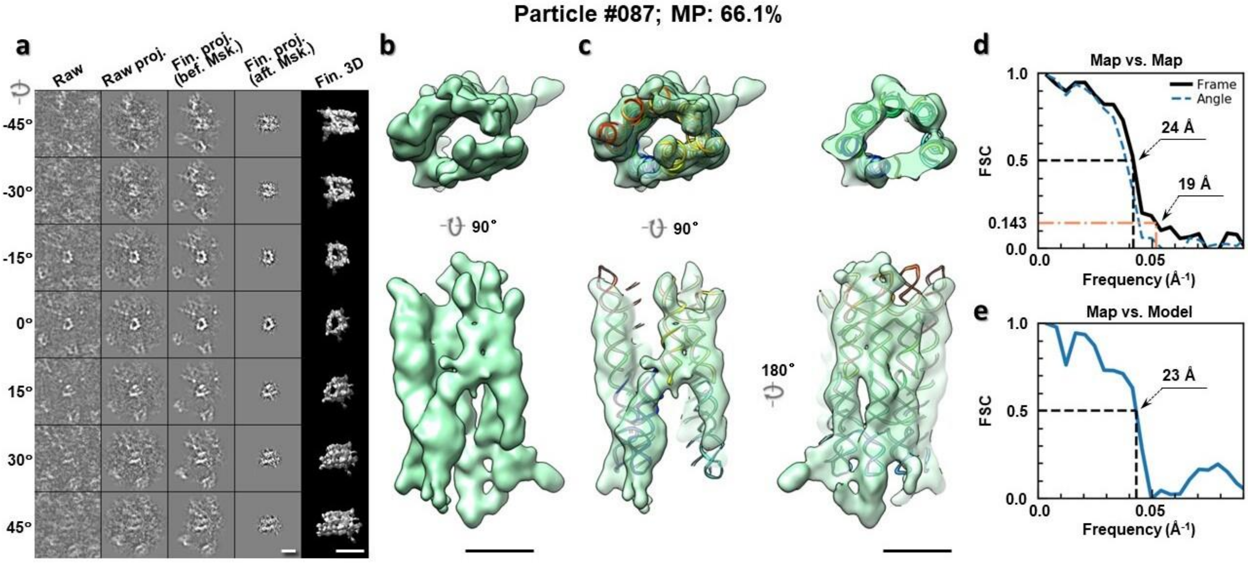

**Supplementary Fig. 57: IPET 3D reconstruction of an individual 6HBC RNA particle #87.** **a,** Seven representative tilt images of a single particle are shown in the first column from the left. The tilt images are aligned to a common center using IPET through iterative refinements. The projections of the raw, intermediate, and final 3D reconstructions at the corresponding angles are displayed in the next four columns. **b,** Two orthogonal views of the final 3D map; **c,** Overlap of the map with the fitted model; **d,** FSC analyses of the final map resolution using two methods, "map-map FSC", in which each map is reconstructed from one half of the images, based on the even vs. odd indices of frames or tilt angles. respectively, **e,** FSC analysis of map-model, where the model map is generated by the fitted model after being low-pass filtered to 8 Å. The resolutions are assessed based on the frequencies of the FSC curve falls at 0.5. Scale bars represent 10 nm in a, and 5 nm in b and c.

Particle #088; MP: 56.2%

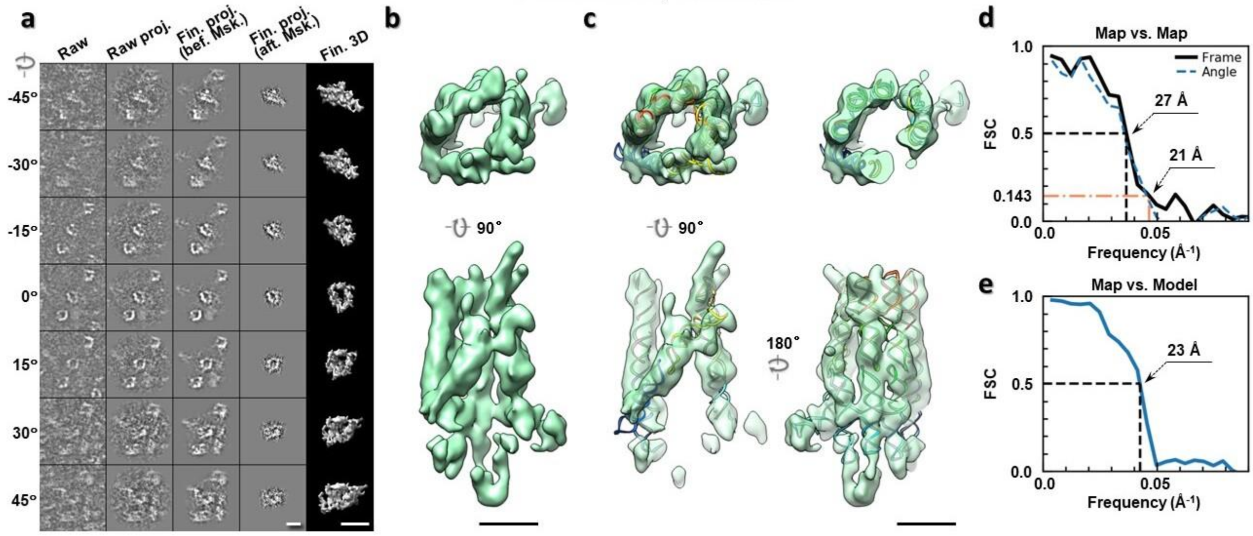

**Supplementary Fig. 58: IPET 3D reconstruction of an individual 6HBC RNA particle #88. a,** Seven representative tilt images of a single particle are shown in the first column from the left. The tilt images are aligned to a common center using IPET through iterative refinements. The projections of the raw, intermediate, and final 3D reconstructions at the corresponding angles are displayed in the next four columns. **b,** Two orthogonal views of the final 3D map; **c,** Overlap of the map with the fitted model; **d,** FSC analyses of the final map resolution using two methods, "map-map FSC", in which each map is reconstructed from one half of the images, based on the even vs. odd indices of frames or tilt angles. respectively, **e,** FSC analysis of map-model, where the model map is generated by the fitted model after being low-pass filtered to 8 Å. The resolutions are assessed based on the frequencies of the FSC curve falls at 0.5. Scale bars represent 10 nm in a, and 5 nm in b and c.

Particle #089; MP: 51.2%

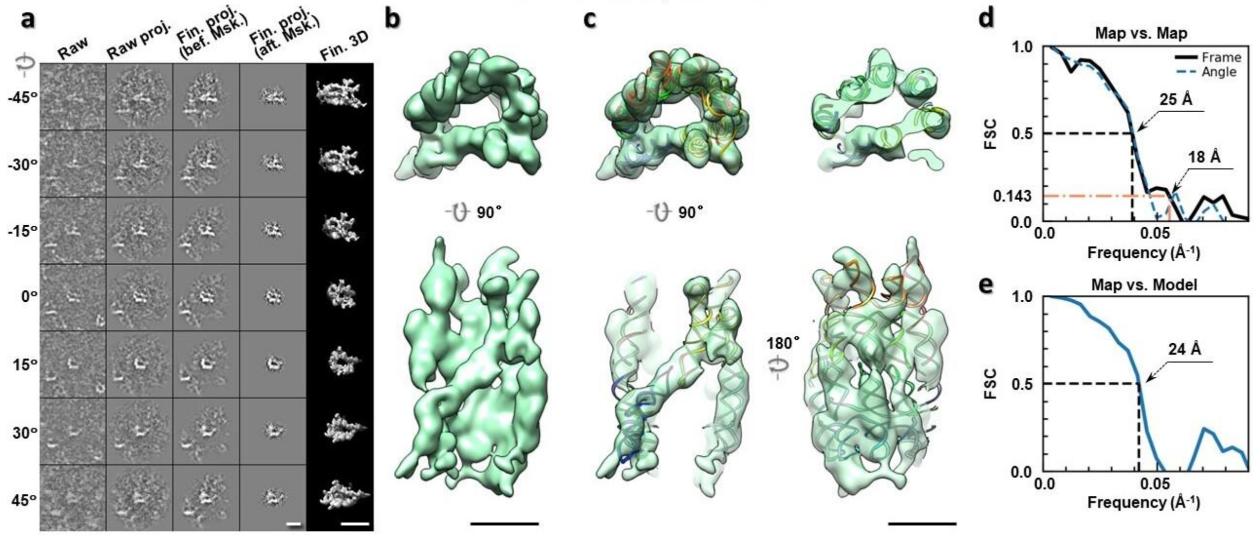

**Supplementary Fig. 59: IPET 3D reconstruction of an individual 6HBC RNA particle #89. a,** Seven representative tilt images of a single particle are shown in the first column from the left. The tilt images are aligned to a common center using IPET through iterative refinements. The projections of the raw, intermediate, and final 3D reconstructions at the corresponding angles are displayed in the next four

columns. **b,** Two orthogonal views of the final 3D map; **c,** Overlap of the map with the fitted model; **d,** FSC analyses of the final map resolution using two methods, "map-map FSC", in which each map is reconstructed from one half of the images, based on the even vs. odd indices of frames or tilt angles. respectively, **e,** FSC analysis of map-model, where the model map is generated by the fitted model after being low-pass filtered to 8 Å. The resolutions are assessed based on the frequencies of the FSC curve falls at 0.5. Scale bars represent 10 nm in a, and 5 nm in b and c.

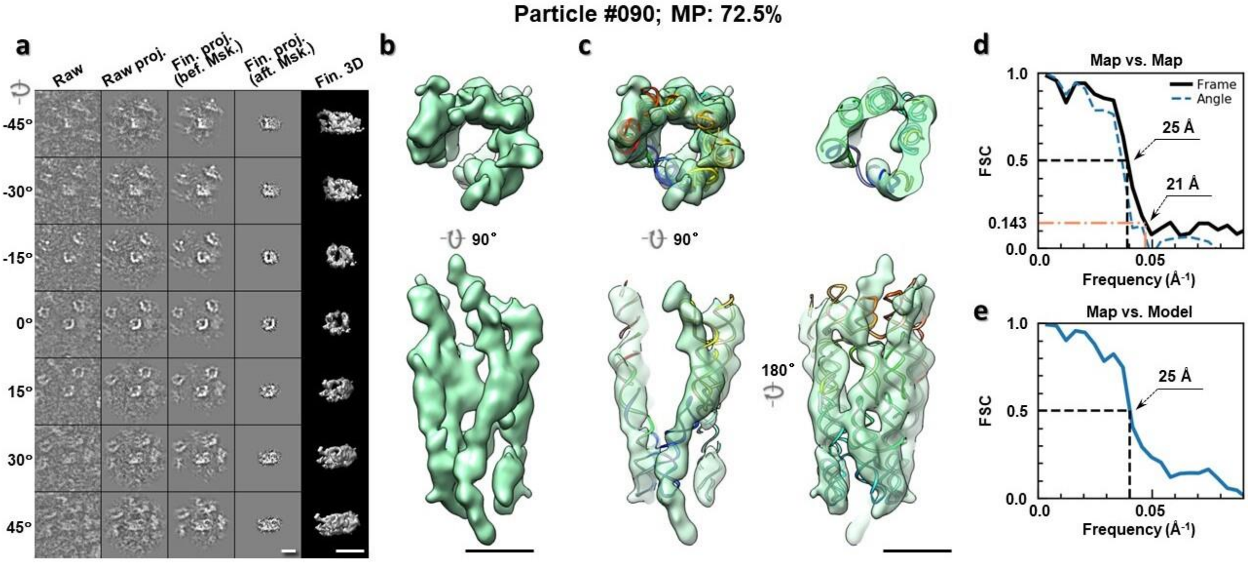

**Supplementary Fig. 60: IPET 3D reconstruction of an individual 6HBC RNA particle #90. a,** Seven representative tilt images of a single particle are shown in the first column from the left. The tilt images are aligned to a common center using IPET through iterative refinements. The projections of the raw, intermediate, and final 3D reconstructions at the corresponding angles are displayed in the next four columns. **b,** Two orthogonal views of the final 3D map; **c,** Overlap of the map with the fitted model; **d,** FSC analyses of the final map resolution using two methods, "map-map FSC", in which each map is reconstructed from one half of the images, based on the even vs. odd indices of frames or tilt angles. respectively, **e,** FSC analysis of map-model, where the model map is generated by the fitted model after being low-pass filtered to 8 Å. The resolutions are assessed based on the frequencies of the FSC curve falls at 0.5. Scale bars represent 10 nm in a, and 5 nm in b and c.

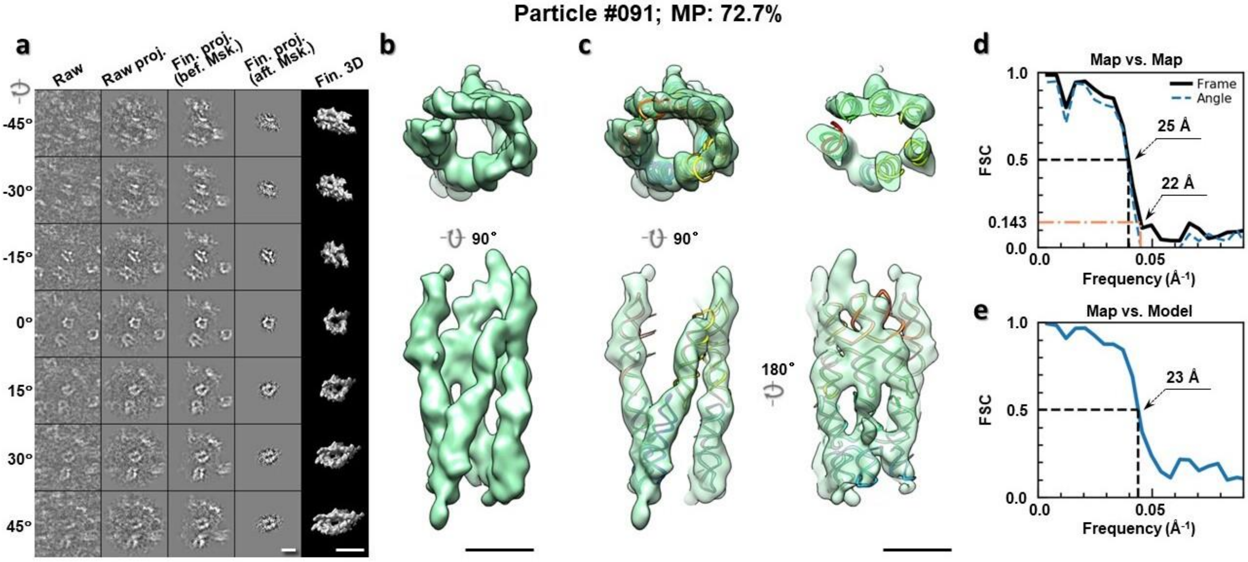

**Supplementary Fig. 61: IPET 3D reconstruction of an individual 6HBC RNA particle #91. a,** Seven representative tilt images of a single particle are shown in the first column from the left. The tilt images are aligned to a common center using IPET through iterative refinements. The projections of the raw, intermediate, and final 3D reconstructions at the corresponding angles are displayed in the next four columns. **b,** Two orthogonal views of the final 3D map; **c,** Overlap of the map with the fitted model; **d,** FSC analyses of the final map resolution using two methods, "map-map FSC", in which each map is reconstructed from one half of the images, based on the even vs. odd indices of frames or tilt angles. respectively, **e,** FSC analysis of map-model, where the model map is generated by the fitted model after being low-pass filtered to 8 Å. The resolutions are assessed based on the frequencies of the FSC curve falls at 0.5. Scale bars represent 10 nm in a, and 5 nm in b and c.

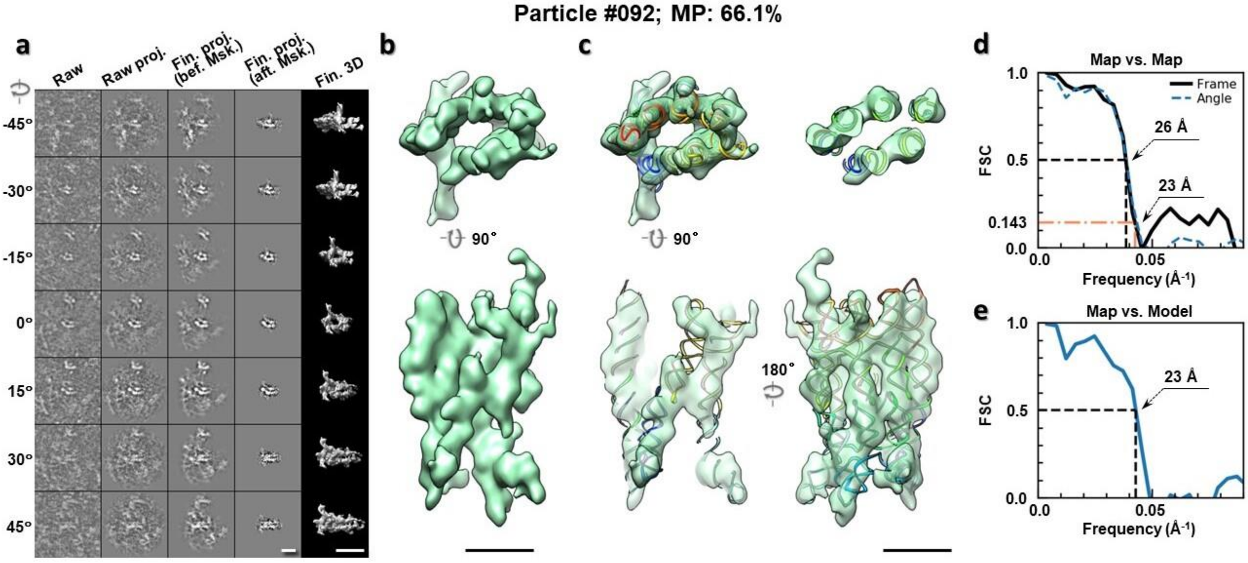

**Supplementary Fig. 62: IPET 3D reconstruction of an individual 6HBC RNA particle #92. a,** Seven representative tilt images of a single particle are shown in the first column from the left. The tilt images are aligned to a common center using IPET through iterative refinements. The projections of the raw, intermediate, and final 3D reconstructions at the corresponding angles are displayed in the next four

columns. **b,** Two orthogonal views of the final 3D map; **c,** Overlap of the map with the fitted model; **d,** FSC analyses of the final map resolution using two methods, "map-map FSC", in which each map is reconstructed from one half of the images, based on the even vs. odd indices of frames or tilt angles. respectively, **e,** FSC analysis of map-model, where the model map is generated by the fitted model after being low-pass filtered to 8 Å. The resolutions are assessed based on the frequencies of the FSC curve falls at 0.5. Scale bars represent 10 nm in a, and 5 nm in b and c.

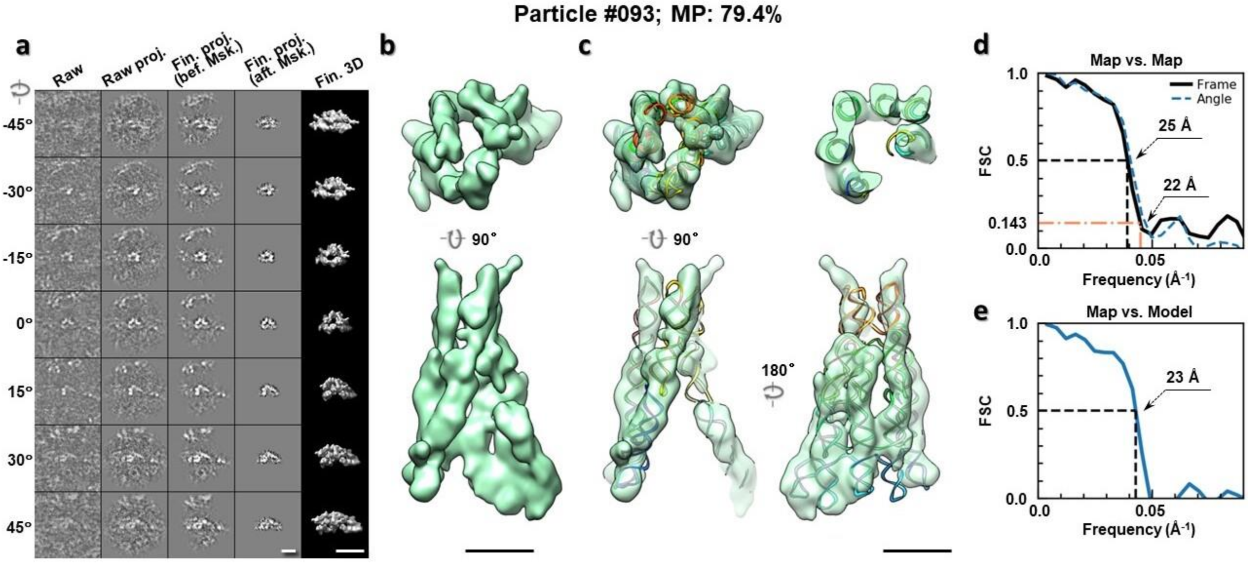

**Supplementary Fig. 63: IPET 3D reconstruction of an individual 6HBC RNA particle #93. a,** Seven representative tilt images of a single particle are shown in the first column from the left. The tilt images are aligned to a common center using IPET through iterative refinements. The projections of the raw, intermediate, and final 3D reconstructions at the corresponding angles are displayed in the next four columns. **b,** Two orthogonal views of the final 3D map; **c,** Overlap of the map with the fitted model; **d,** FSC analyses of the final map resolution using two methods, "map-map FSC", in which each map is reconstructed from one half of the images, based on the even vs. odd indices of frames or tilt angles. respectively, **e,** FSC analysis of map-model, where the model map is generated by the fitted model after being low-pass filtered to 8 Å. The resolutions are assessed based on the frequencies of the FSC curve falls at 0.5. Scale bars represent 10 nm in a, and 5 nm in b and c.

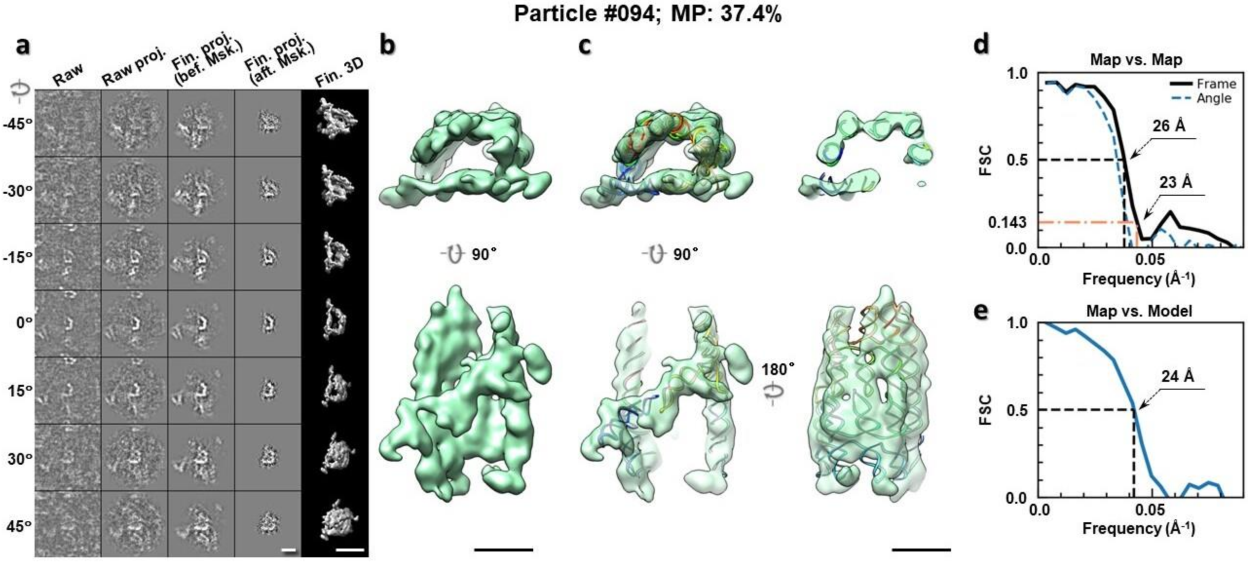

**Supplementary Fig. 64: IPET 3D reconstruction of an individual 6HBC RNA particle #94.** **a,** Seven representative tilt images of a single particle are shown in the first column from the left. The tilt images are aligned to a common center using IPET through iterative refinements. The projections of the raw, intermediate, and final 3D reconstructions at the corresponding angles are displayed in the next four columns. **b,** Two orthogonal views of the final 3D map; **c,** Overlap of the map with the fitted model; **d,** FSC analyses of the final map resolution using two methods, "map-map FSC", in which each map is reconstructed from one half of the images, based on the even vs. odd indices of frames or tilt angles. respectively, **e,** FSC analysis of map-model, where the model map is generated by the fitted model after being low-pass filtered to 8 Å. The resolutions are assessed based on the frequencies of the FSC curve falls at 0.5. Scale bars represent 10 nm in a, and 5 nm in b and c.

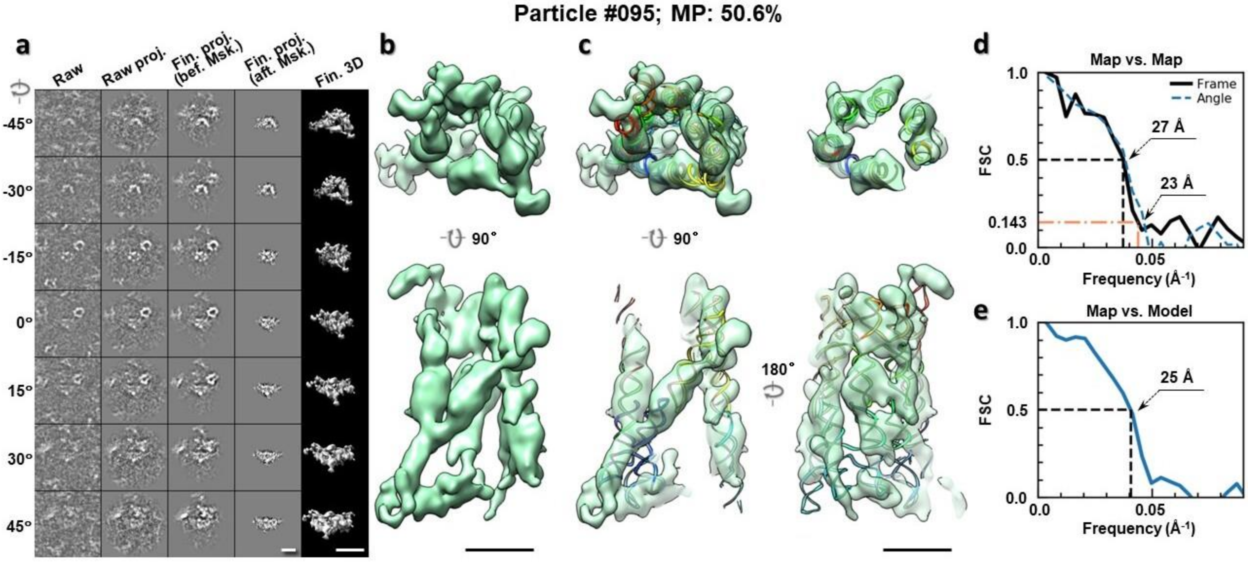

**Supplementary Fig. 65: IPET 3D reconstruction of an individual 6HBC RNA particle #95.** **a,** Seven representative tilt images of a single particle are shown in the first column from the left. The tilt images are aligned to a common center using IPET through iterative refinements. The projections of the raw, intermediate, and final 3D reconstructions at the corresponding angles are displayed in the next four

columns. **b,** Two orthogonal views of the final 3D map; **c,** Overlap of the map with the fitted model; **d,** FSC analyses of the final map resolution using two methods, "map-map FSC", in which each map is reconstructed from one half of the images, based on the even vs. odd indices of frames or tilt angles. respectively, **e,** FSC analysis of map-model, where the model map is generated by the fitted model after being low-pass filtered to 8 Å. The resolutions are assessed based on the frequencies of the FSC curve falls at 0.5. Scale bars represent 10 nm in a, and 5 nm in b and c.

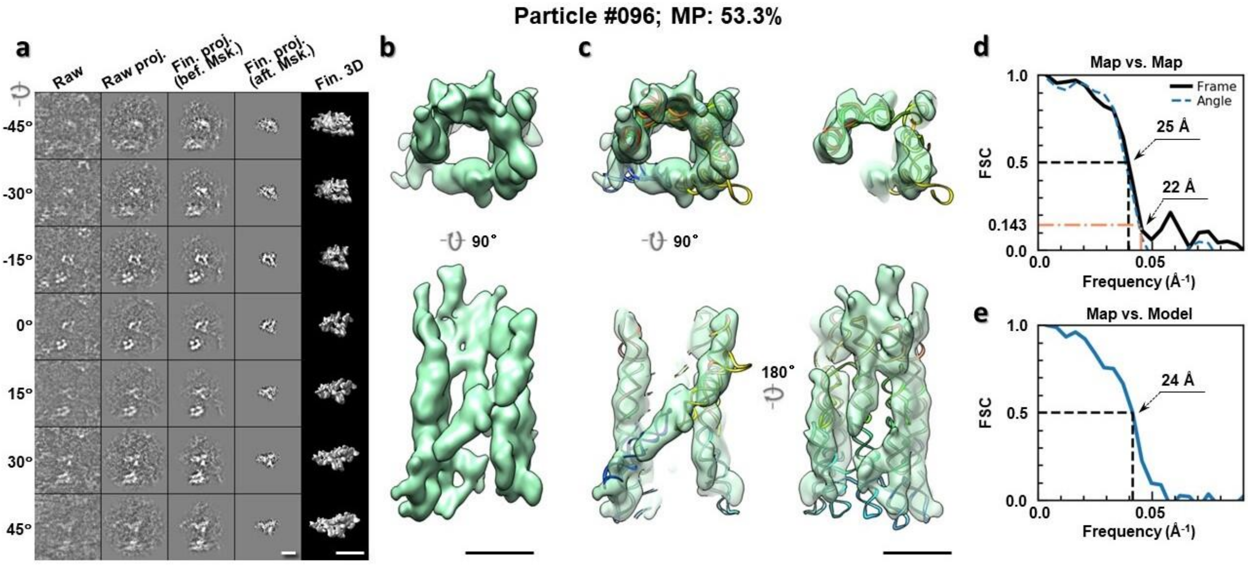

**Supplementary Fig. 66: IPET 3D reconstruction of an individual 6HBC RNA particle #96.** **a,** Seven representative tilt images of a single particle are shown in the first column from the left. The tilt images are aligned to a common center using IPET through iterative refinements. The projections of the raw, intermediate, and final 3D reconstructions at the corresponding angles are displayed in the next four columns. **b,** Two orthogonal views of the final 3D map; **c,** Overlap of the map with the fitted model; **d,** FSC analyses of the final map resolution using two methods, "map-map FSC", in which each map is reconstructed from one half of the images, based on the even vs. odd indices of frames or tilt angles. respectively, **e,** FSC analysis of map-model, where the model map is generated by the fitted model after being low-pass filtered to 8 Å. The resolutions are assessed based on the frequencies of the FSC curve falls at 0.5. Scale bars represent 10 nm in a, and 5 nm in b and c.

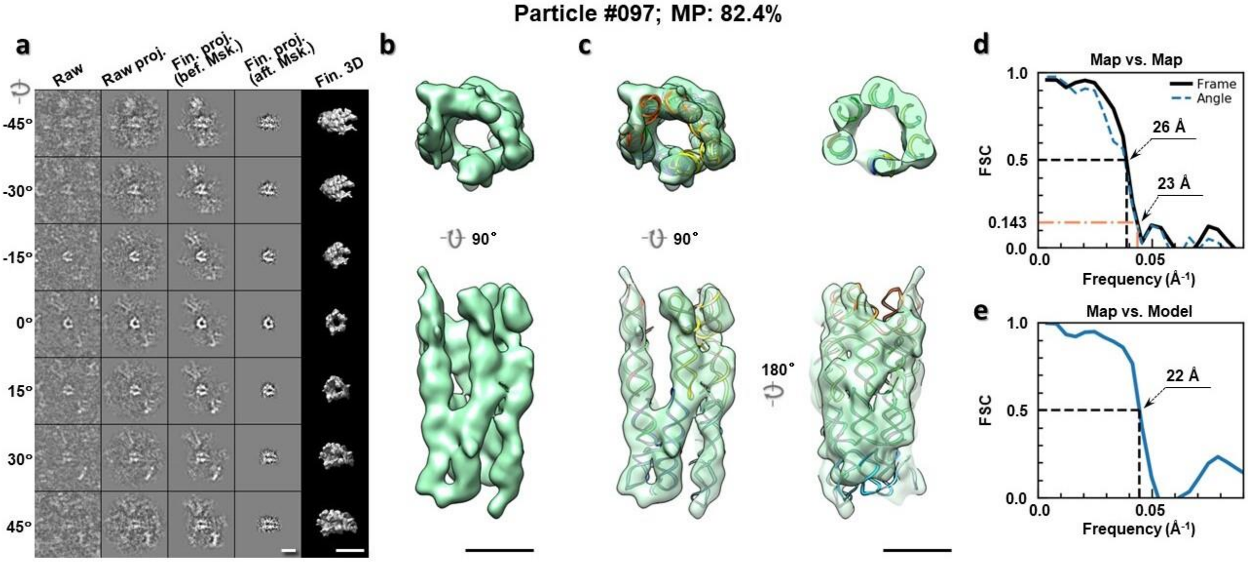

**Supplementary Fig. 67: IPET 3D reconstruction of an individual 6HBC RNA particle #97. a,** Seven representative tilt images of a single particle are shown in the first column from the left. The tilt images are aligned to a common center using IPET through iterative refinements. The projections of the raw, intermediate, and final 3D reconstructions at the corresponding angles are displayed in the next four columns. **b,** Two orthogonal views of the final 3D map; **c,** Overlap of the map with the fitted model; **d,** FSC analyses of the final map resolution using two methods, "map-map FSC", in which each map is reconstructed from one half of the images, based on the even vs. odd indices of frames or tilt angles. respectively, **e,** FSC analysis of map-model, where the model map is generated by the fitted model after being low-pass filtered to 8 Å. The resolutions are assessed based on the frequencies of the FSC curve falls at 0.5. Scale bars represent 10 nm in a, and 5 nm in b and c.

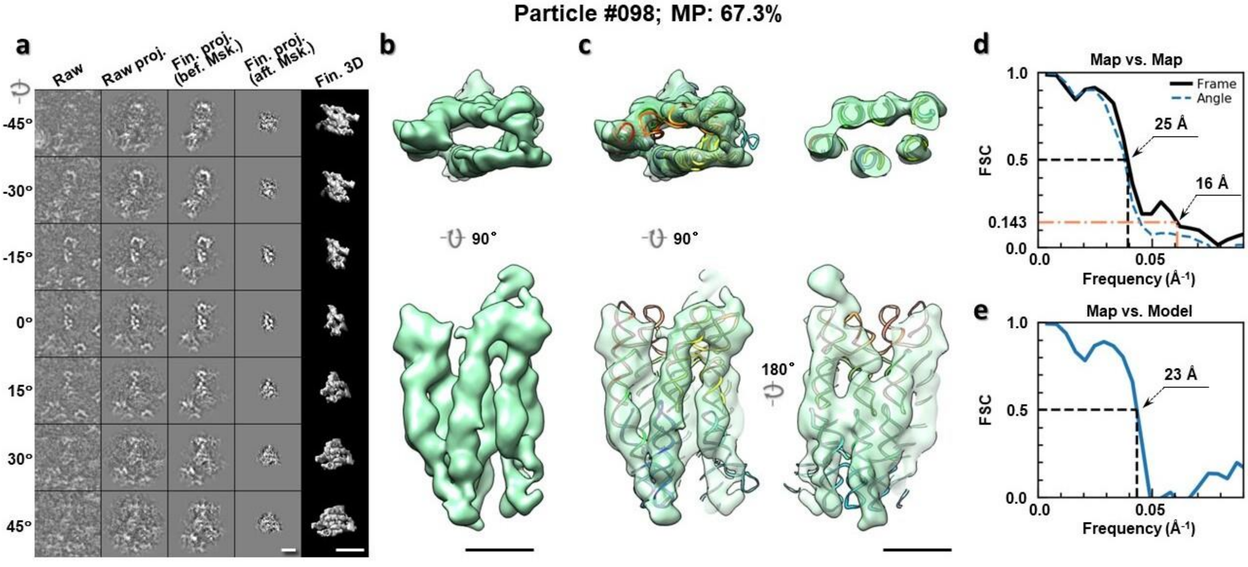

**Supplementary Fig. 68: IPET 3D reconstruction of an individual 6HBC RNA particle #98. a,** Seven representative tilt images of a single particle are shown in the first column from the left. The tilt images are aligned to a common center using IPET through iterative refinements. The projections of the raw, intermediate, and final 3D reconstructions at the corresponding angles are displayed in the next four

columns. **b,** Two orthogonal views of the final 3D map; **c,** Overlap of the map with the fitted model; **d,** FSC analyses of the final map resolution using two methods, "map-map FSC", in which each map is reconstructed from one half of the images, based on the even vs. odd indices of frames or tilt angles. respectively, **e,** FSC analysis of map-model, where the model map is generated by the fitted model after being low-pass filtered to 8 Å. The resolutions are assessed based on the frequencies of the FSC curve falls at 0.5. Scale bars represent 10 nm in a, and 5 nm in b and c.

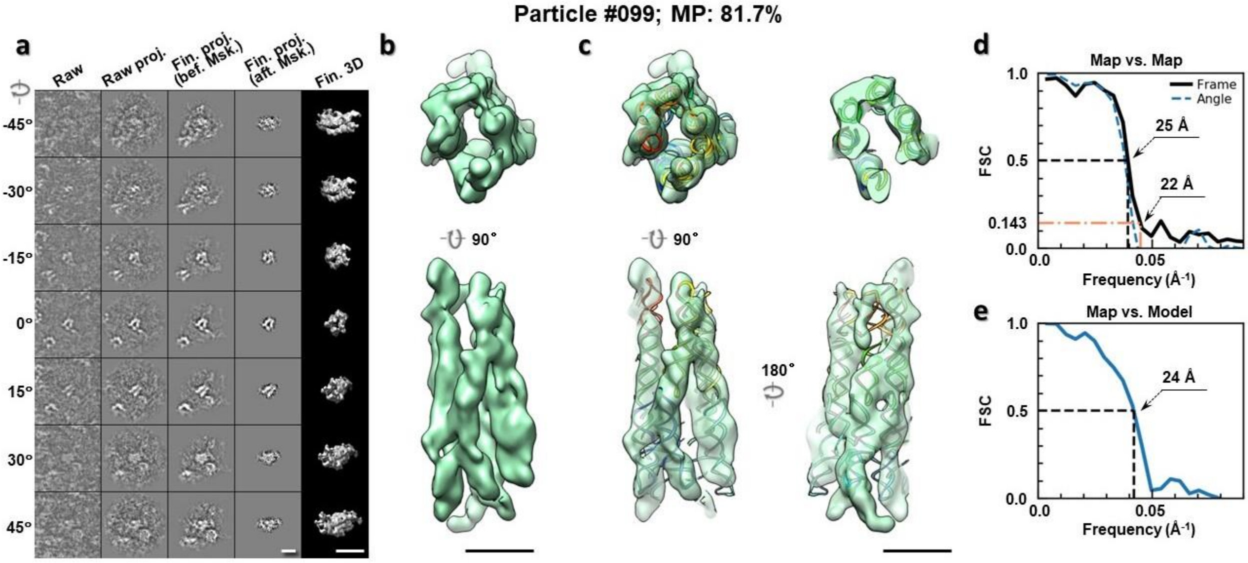

**Supplementary Fig. 69: IPET 3D reconstruction of an individual 6HBC RNA particle #99.** **a,** Seven representative tilt images of a single particle are shown in the first column from the left. The tilt images are aligned to a common center using IPET through iterative refinements. The projections of the raw, intermediate, and final 3D reconstructions at the corresponding angles are displayed in the next four columns. **b,** Two orthogonal views of the final 3D map; **c,** Overlap of the map with the fitted model; **d,** FSC analyses of the final map resolution using two methods, "map-map FSC", in which each map is reconstructed from one half of the images, based on the even vs. odd indices of frames or tilt angles. respectively, **e,** FSC analysis of map-model, where the model map is generated by the fitted model after being low-pass filtered to 8 Å. The resolutions are assessed based on the frequencies of the FSC curve falls at 0.5. Scale bars represent 10 nm in a, and 5 nm in b and c.

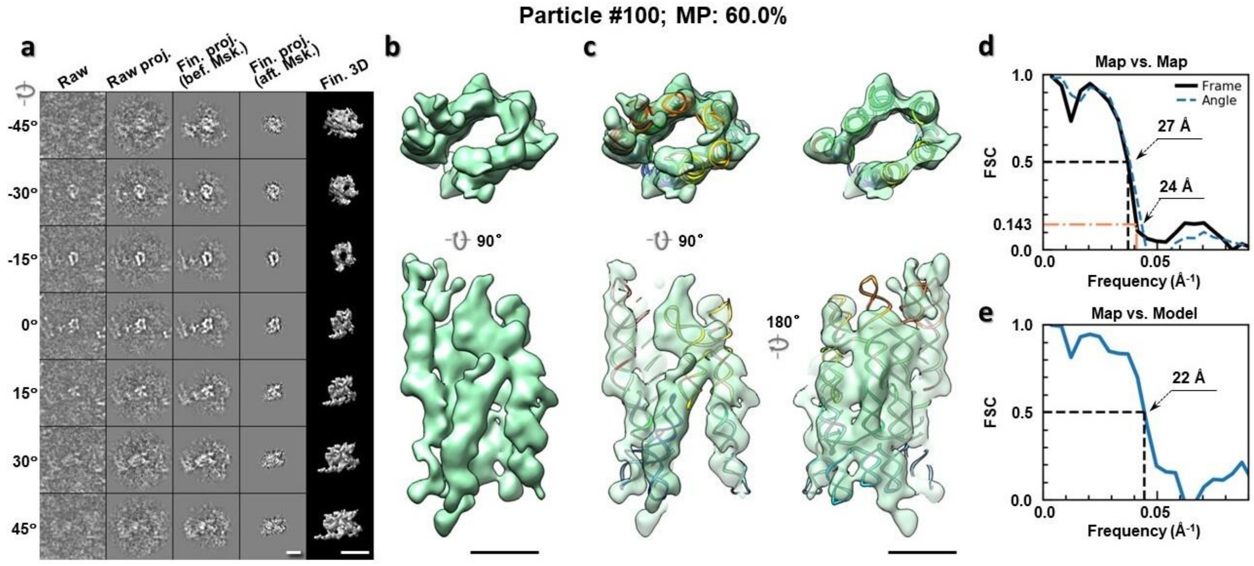

**Supplementary Fig. 70: IPET 3D reconstruction of an individual 6HBC RNA particle #100.** **a,** Seven representative tilt images of a single particle are shown in the first column from the left. The tilt images are aligned to a common center using IPET through iterative refinements. The projections of the raw, intermediate, and final 3D reconstructions at the corresponding angles are displayed in the next four columns. **b,** Two orthogonal views of the final 3D map; **c,** Overlap of the map with the fitted model; **d,** FSC analyses of the final map resolution using two methods, "map-map FSC", in which each map is reconstructed from one half of the images, based on the even vs. odd indices of frames or tilt angles. respectively, **e,** FSC analysis of map-model, where the model map is generated by the fitted model after being low-pass filtered to 8 Å. The resolutions are assessed based on the frequencies of the FSC curve falls at 0.5. Scale bars represent 10 nm in a, and 5 nm in b and c.

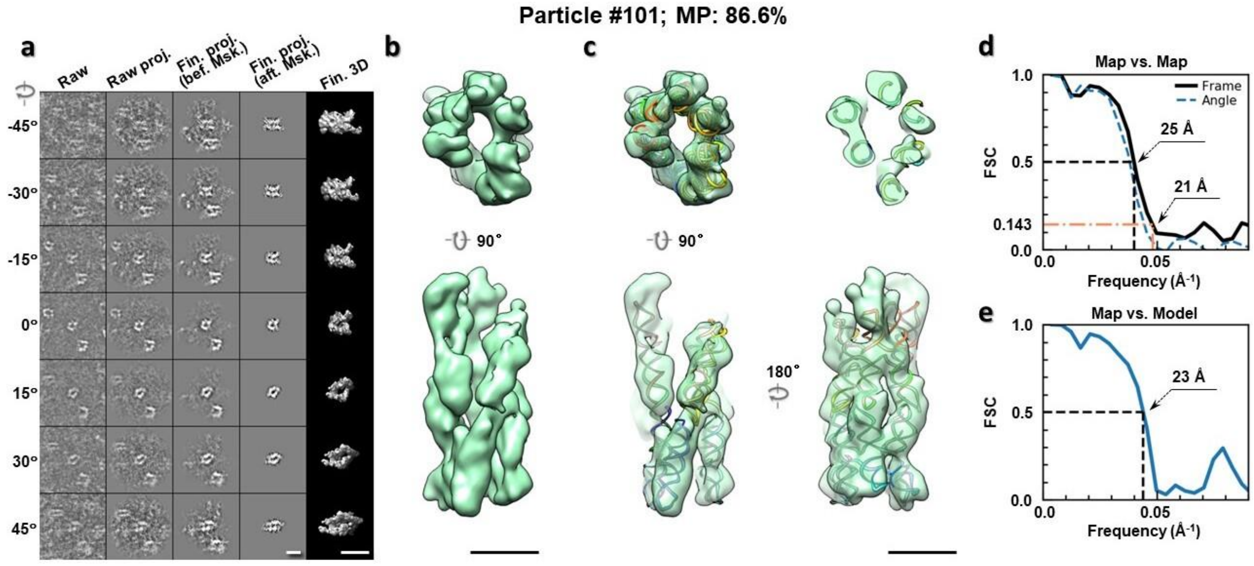

**Supplementary Fig. 71: IPET 3D reconstruction of an individual 6HBC RNA particle #101.** **a,** Seven representative tilt images of a single particle are shown in the first column from the left. The tilt images are aligned to a common center using IPET through iterative refinements. The projections of the raw, intermediate, and final 3D reconstructions at the corresponding angles are displayed in the next four

columns. **b,** Two orthogonal views of the final 3D map; **c,** Overlap of the map with the fitted model; **d,** FSC analyses of the final map resolution using two methods, "map-map FSC", in which each map is reconstructed from one half of the images, based on the even vs. odd indices of frames or tilt angles. respectively, **e,** FSC analysis of map-model, where the model map is generated by the fitted model after being low-pass filtered to 8 Å. The resolutions are assessed based on the frequencies of the FSC curve falls at 0.5. Scale bars represent 10 nm in a, and 5 nm in b and c.

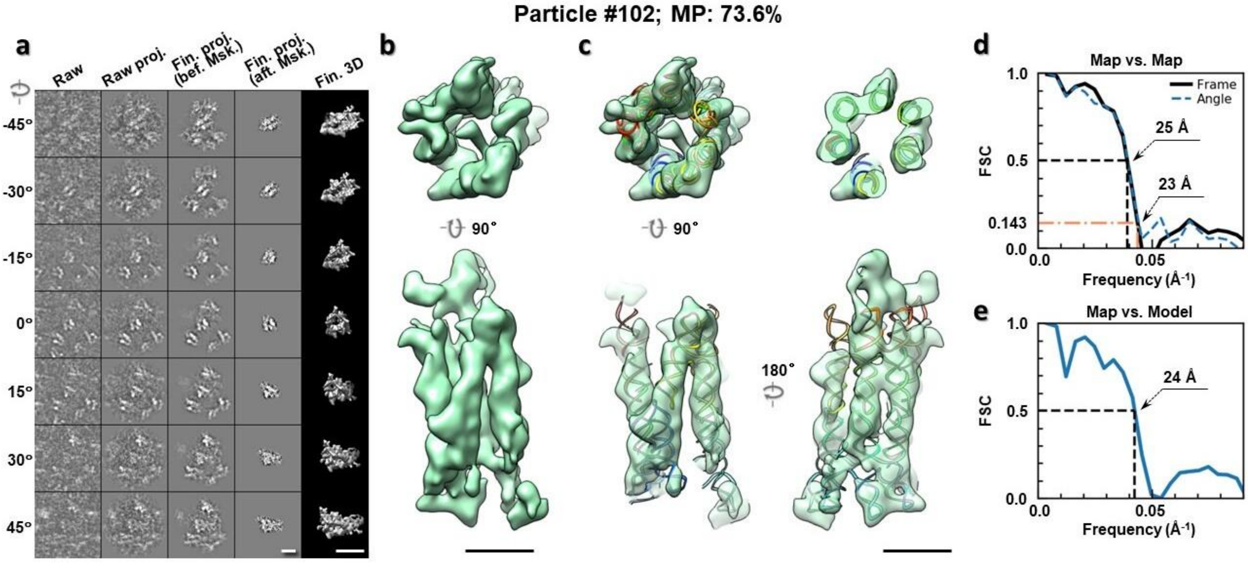

**Supplementary Fig. 72: IPET 3D reconstruction of an individual 6HBC RNA particle #102.** **a,** Seven representative tilt images of a single particle are shown in the first column from the left. The tilt images are aligned to a common center using IPET through iterative refinements. The projections of the raw, intermediate, and final 3D reconstructions at the corresponding angles are displayed in the next four columns. **b,** Two orthogonal views of the final 3D map; **c,** Overlap of the map with the fitted model; **d,** FSC analyses of the final map resolution using two methods, "map-map FSC", in which each map is reconstructed from one half of the images, based on the even vs. odd indices of frames or tilt angles. respectively, **e,** FSC analysis of map-model, where the model map is generated by the fitted model after being low-pass filtered to 8 Å. The resolutions are assessed based on the frequencies of the FSC curve falls at 0.5. Scale bars represent 10 nm in a, and 5 nm in b and c.

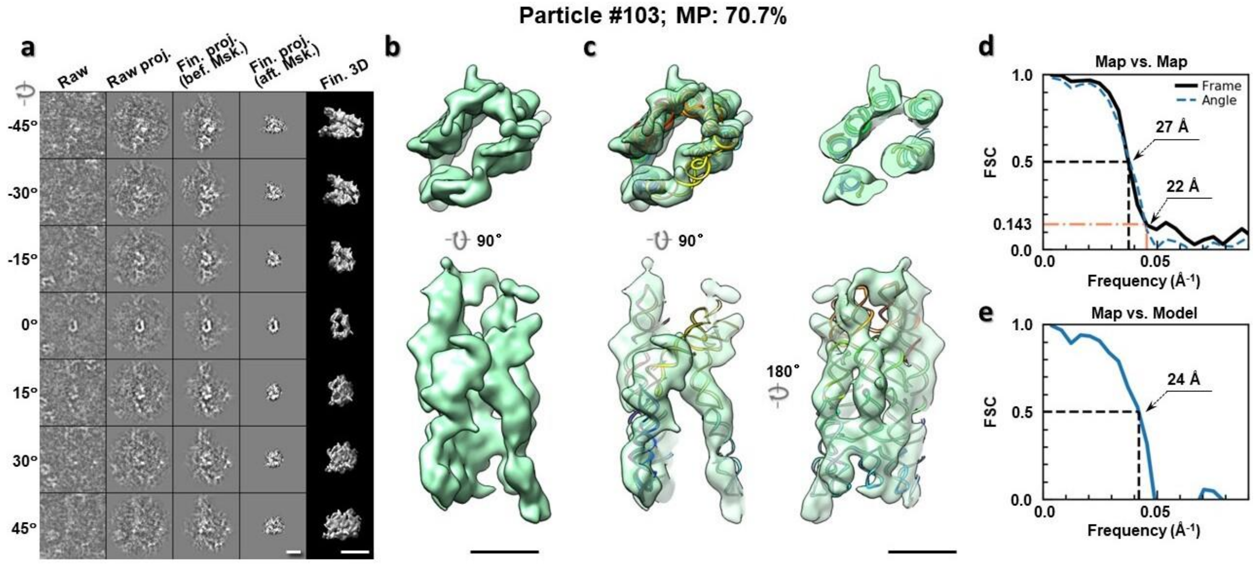

**Supplementary Fig. 73: IPET 3D reconstruction of an individual 6HBC RNA particle #103. a,** Seven representative tilt images of a single particle are shown in the first column from the left. The tilt images are aligned to a common center using IPET through iterative refinements. The projections of the raw, intermediate, and final 3D reconstructions at the corresponding angles are displayed in the next four columns. **b,** Two orthogonal views of the final 3D map; **c,** Overlap of the map with the fitted model; **d,** FSC analyses of the final map resolution using two methods, "map-map FSC", in which each map is reconstructed from one half of the images, based on the even vs. odd indices of frames or tilt angles. respectively, **e,** FSC analysis of map-model, where the model map is generated by the fitted model after being low-pass filtered to 8 Å. The resolutions are assessed based on the frequencies of the FSC curve falls at 0.5. Scale bars represent 10 nm in a, and 5 nm in b and c.

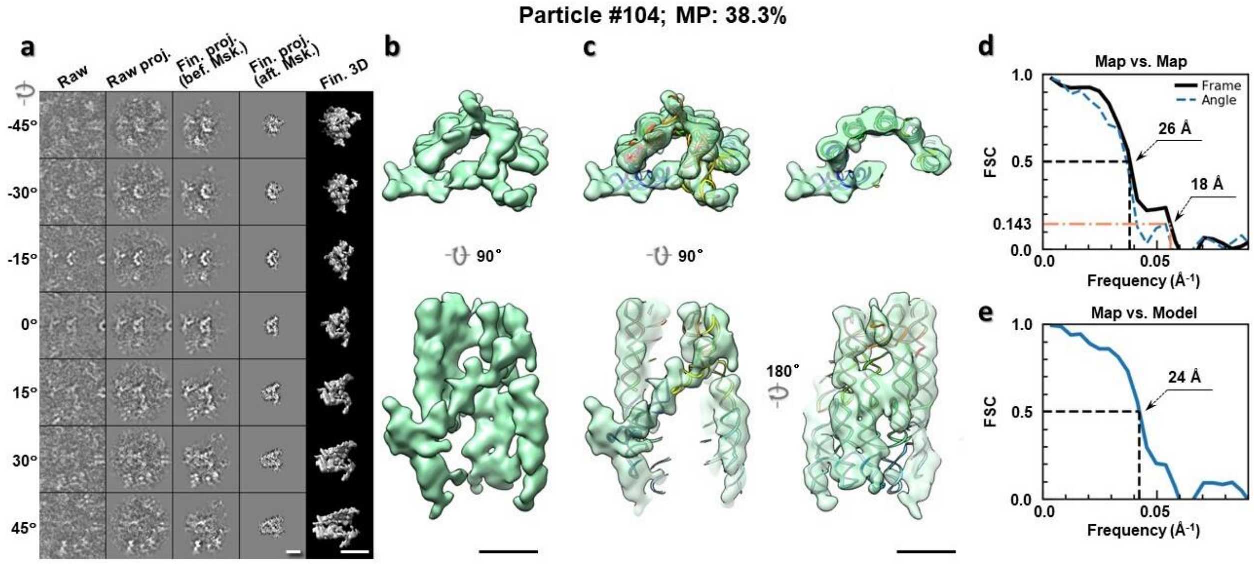

**Supplementary Fig. 74: IPET 3D reconstruction of an individual 6HBC RNA particle #104. a,** Seven representative tilt images of a single particle are shown in the first column from the left. The tilt images are aligned to a common center using IPET through iterative refinements. The projections of the raw, intermediate, and final 3D reconstructions at the corresponding angles are displayed in the next four

columns. **b,** Two orthogonal views of the final 3D map; **c,** Overlap of the map with the fitted model; **d,** FSC analyses of the final map resolution using two methods, "map-map FSC", in which each map is reconstructed from one half of the images, based on the even vs. odd indices of frames or tilt angles. respectively, **e,** FSC analysis of map-model, where the model map is generated by the fitted model after being low-pass filtered to 8 Å. The resolutions are assessed based on the frequencies of the FSC curve falls at 0.5. Scale bars represent 10 nm in a, and 5 nm in b and c.

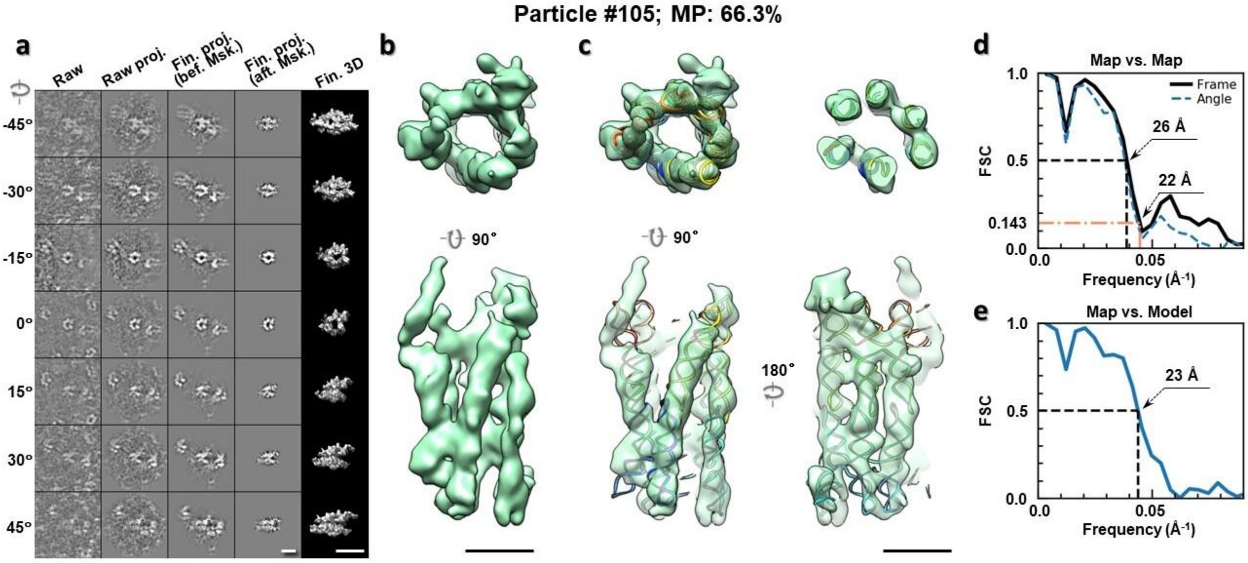

**Supplementary Fig. 75: IPET 3D reconstruction of an individual 6HBC RNA particle #105.** **a,** Seven representative tilt images of a single particle are shown in the first column from the left. The tilt images are aligned to a common center using IPET through iterative refinements. The projections of the raw, intermediate, and final 3D reconstructions at the corresponding angles are displayed in the next four columns. **b,** Two orthogonal views of the final 3D map; **c,** Overlap of the map with the fitted model; **d,** FSC analyses of the final map resolution using two methods, "map-map FSC", in which each map is reconstructed from one half of the images, based on the even vs. odd indices of frames or tilt angles. respectively, **e,** FSC analysis of map-model, where the model map is generated by the fitted model after being low-pass filtered to 8 Å. The resolutions are assessed based on the frequencies of the FSC curve falls at 0.5. Scale bars represent 10 nm in a, and 5 nm in b and c.

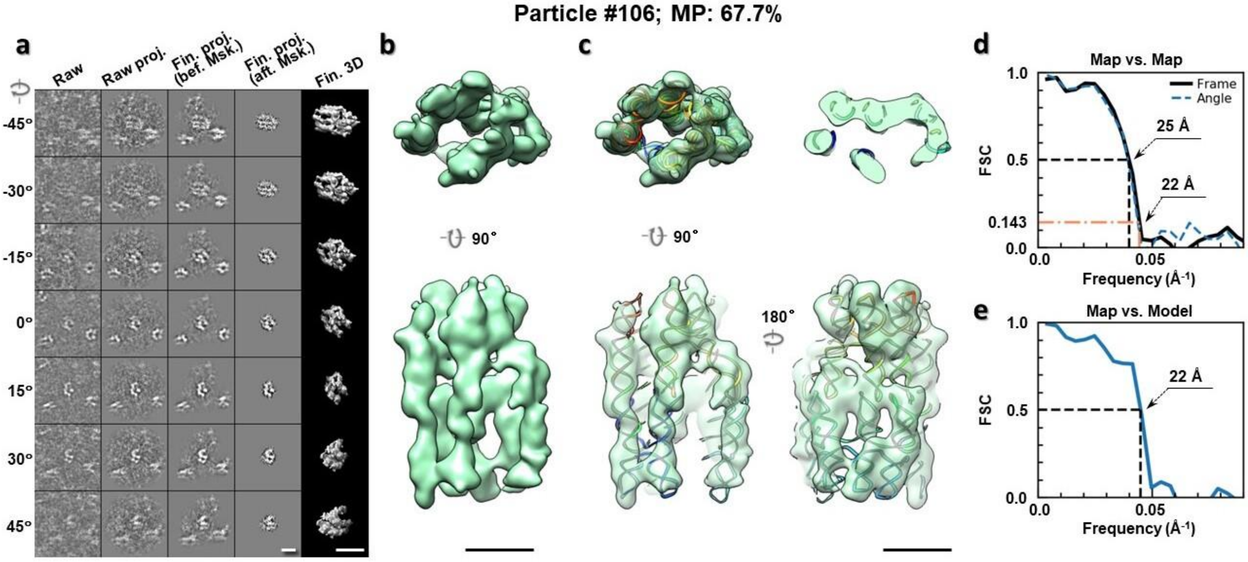

**Supplementary Fig. 76: IPET 3D reconstruction of an individual 6HBC RNA particle #106. a,** Seven representative tilt images of a single particle are shown in the first column from the left. The tilt images are aligned to a common center using IPET through iterative refinements. The projections of the raw, intermediate, and final 3D reconstructions at the corresponding angles are displayed in the next four columns. **b,** Two orthogonal views of the final 3D map; **c,** Overlap of the map with the fitted model; **d,** FSC analyses of the final map resolution using two methods, "map-map FSC", in which each map is reconstructed from one half of the images, based on the even vs. odd indices of frames or tilt angles. respectively, **e,** FSC analysis of map-model, where the model map is generated by the fitted model after being low-pass filtered to 8 Å. The resolutions are assessed based on the frequencies of the FSC curve falls at 0.5. Scale bars represent 10 nm in a, and 5 nm in b and c.

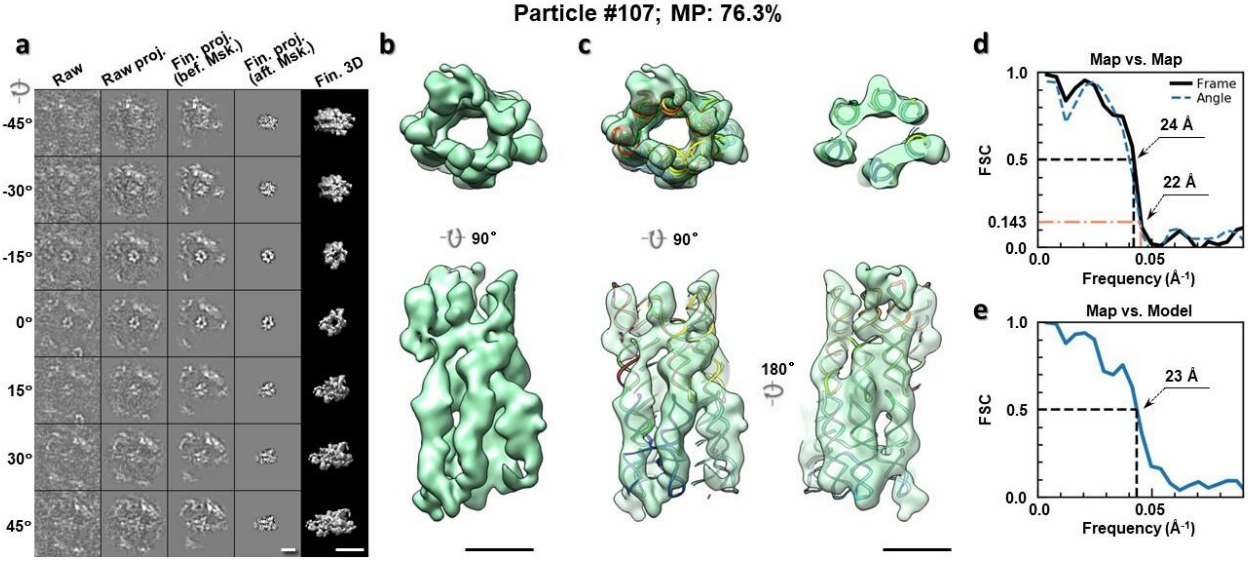

**Supplementary Fig. 77: IPET 3D reconstruction of an individual 6HBC RNA particle #107. a,** Seven representative tilt images of a single particle are shown in the first column from the left. The tilt images are aligned to a common center using IPET through iterative refinements. The projections of the raw, intermediate, and final 3D reconstructions at the corresponding angles are displayed in the next four

columns. **b,** Two orthogonal views of the final 3D map; **c,** Overlap of the map with the fitted model; **d,** FSC analyses of the final map resolution using two methods, "map-map FSC", in which each map is reconstructed from one half of the images, based on the even vs. odd indices of frames or tilt angles. respectively, **e,** FSC analysis of map-model, where the model map is generated by the fitted model after being low-pass filtered to 8 Å. The resolutions are assessed based on the frequencies of the FSC curve falls at 0.5. Scale bars represent 10 nm in a, and 5 nm in b and c.

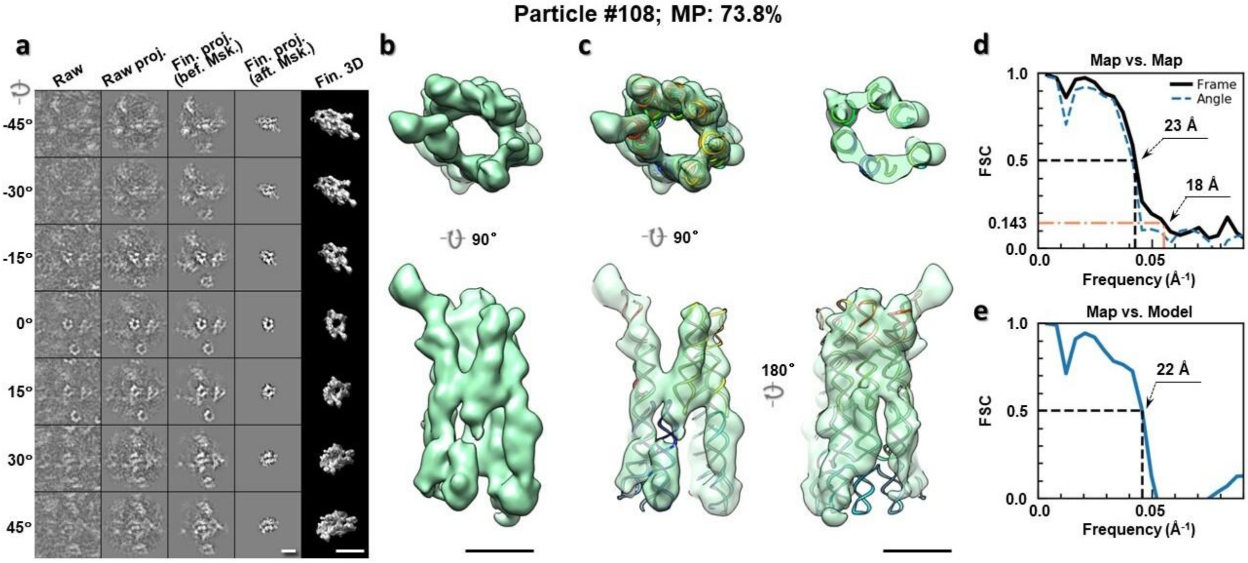

**Supplementary Fig. 78: IPET 3D reconstruction of an individual 6HBC RNA particle #108. a,** Seven representative tilt images of a single particle are shown in the first column from the left. The tilt images are aligned to a common center using IPET through iterative refinements. The projections of the raw, intermediate, and final 3D reconstructions at the corresponding angles are displayed in the next four columns. **b,** Two orthogonal views of the final 3D map; **c,** Overlap of the map with the fitted model; **d,** FSC analyses of the final map resolution using two methods, "map-map FSC", in which each map is reconstructed from one half of the images, based on the even vs. odd indices of frames or tilt angles. respectively, **e,** FSC analysis of map-model, where the model map is generated by the fitted model after being low-pass filtered to 8 Å. The resolutions are assessed based on the frequencies of the FSC curve falls at 0.5. Scale bars represent 10 nm in a, and 5 nm in b and c.

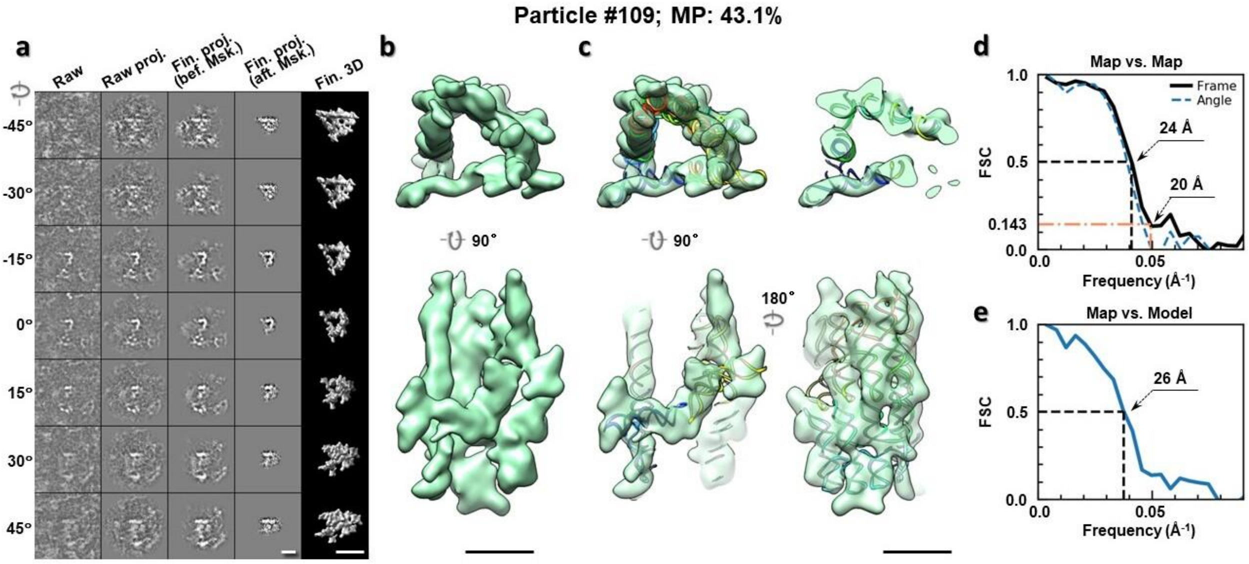

**Supplementary Fig. 79: IPET 3D reconstruction of an individual 6HBC RNA particle #109.** **a,** Seven representative tilt images of a single particle are shown in the first column from the left. The tilt images are aligned to a common center using IPET through iterative refinements. The projections of the raw, intermediate, and final 3D reconstructions at the corresponding angles are displayed in the next four columns. **b,** Two orthogonal views of the final 3D map; **c,** Overlap of the map with the fitted model; **d,** FSC analyses of the final map resolution using two methods, "map-map FSC", in which each map is reconstructed from one half of the images, based on the even vs. odd indices of frames or tilt angles. respectively, **e,** FSC analysis of map-model, where the model map is generated by the fitted model after being low-pass filtered to 8 Å. The resolutions are assessed based on the frequencies of the FSC curve falls at 0.5. Scale bars represent 10 nm in a, and 5 nm in b and c.

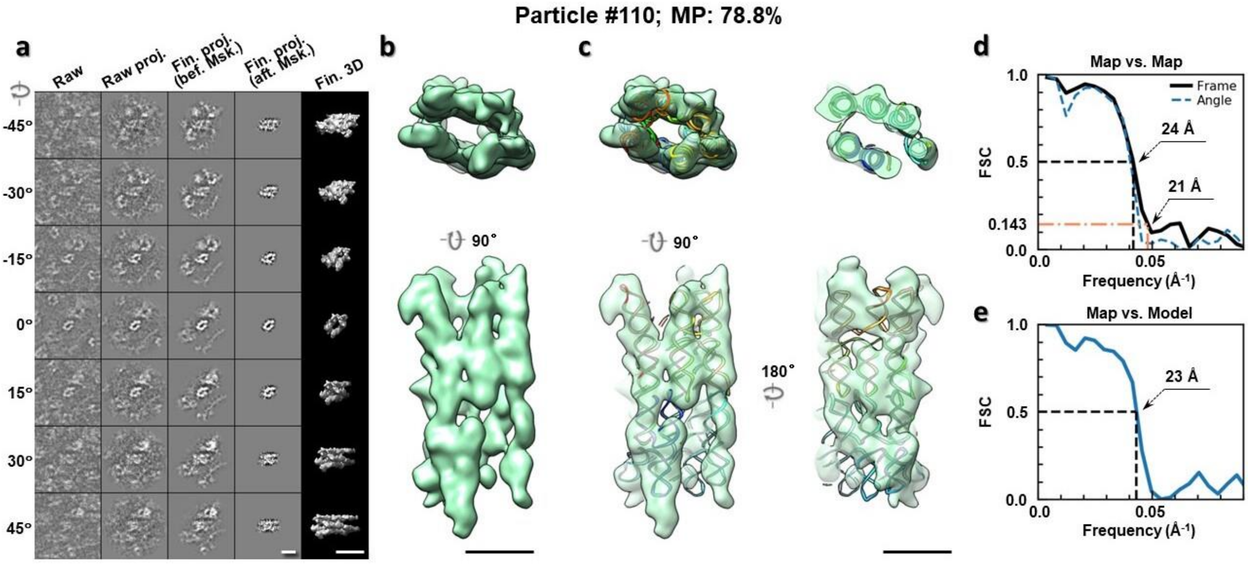

**Supplementary Fig. 80: IPET 3D reconstruction of an individual 6HBC RNA particle #110.** **a,** Seven representative tilt images of a single particle are shown in the first column from the left. The tilt images are aligned to a common center using IPET through iterative refinements. The projections of the raw, intermediate, and final 3D reconstructions at the corresponding angles are displayed in the next four

columns. **b,** Two orthogonal views of the final 3D map; **c,** Overlap of the map with the fitted model; **d,** FSC analyses of the final map resolution using two methods, "map-map FSC", in which each map is reconstructed from one half of the images, based on the even vs. odd indices of frames or tilt angles. respectively, **e,** FSC analysis of map-model, where the model map is generated by the fitted model after being low-pass filtered to 8 Å. The resolutions are assessed based on the frequencies of the FSC curve falls at 0.5. Scale bars represent 10 nm in a, and 5 nm in b and c.

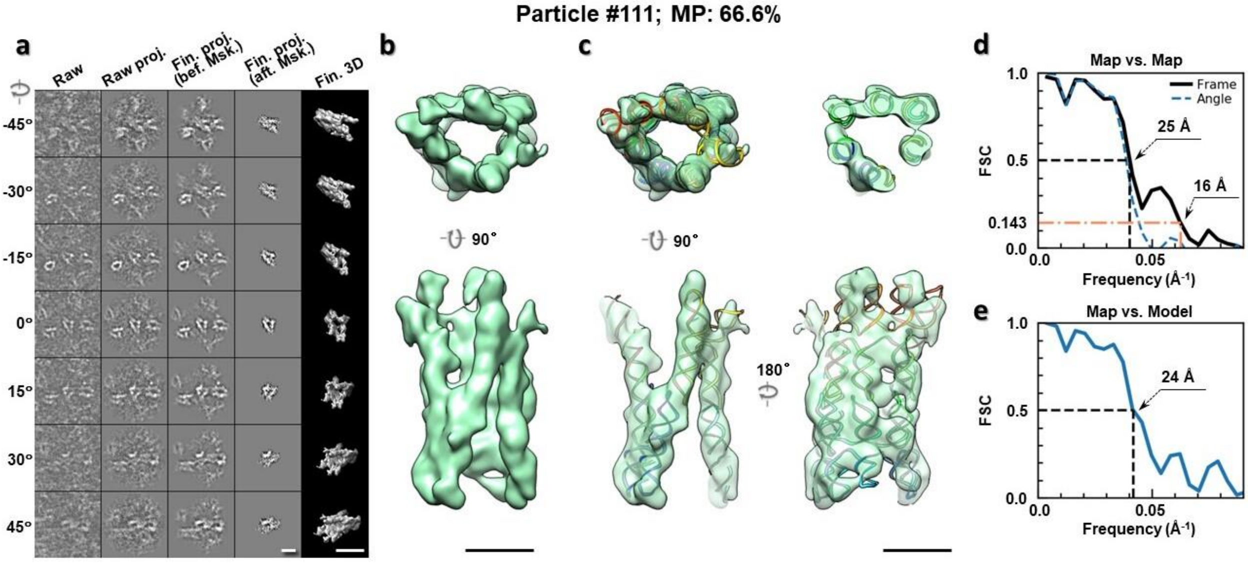

**Supplementary Fig. 81: IPET 3D reconstruction of an individual 6HBC RNA particle #111. a,** Seven representative tilt images of a single particle are shown in the first column from the left. The tilt images are aligned to a common center using IPET through iterative refinements. The projections of the raw, intermediate, and final 3D reconstructions at the corresponding angles are displayed in the next four columns. **b,** Two orthogonal views of the final 3D map; **c,** Overlap of the map with the fitted model; **d,** FSC analyses of the final map resolution using two methods, "map-map FSC", in which each map is reconstructed from one half of the images, based on the even vs. odd indices of frames or tilt angles. respectively, **e,** FSC analysis of map-model, where the model map is generated by the fitted model after being low-pass filtered to 8 Å. The resolutions are assessed based on the frequencies of the FSC curve falls at 0.5. Scale bars represent 10 nm in a, and 5 nm in b and c.

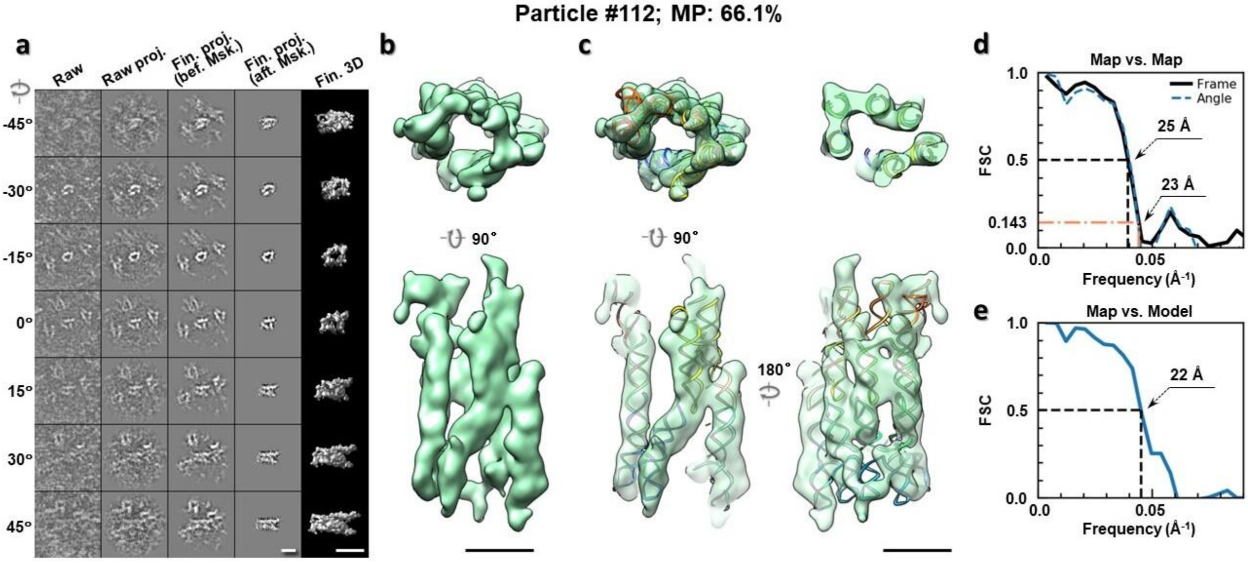

**Supplementary Fig. 82: IPET 3D reconstruction of an individual 6HBC RNA particle #112.** **a,** Seven representative tilt images of a single particle are shown in the first column from the left. The tilt images are aligned to a common center using IPET through iterative refinements. The projections of the raw, intermediate, and final 3D reconstructions at the corresponding angles are displayed in the next four columns. **b,** Two orthogonal views of the final 3D map; **c,** Overlap of the map with the fitted model; **d,** FSC analyses of the final map resolution using two methods, "map-map FSC", in which each map is reconstructed from one half of the images, based on the even vs. odd indices of frames or tilt angles. respectively, **e,** FSC analysis of map-model, where the model map is generated by the fitted model after being low-pass filtered to 8 Å. The resolutions are assessed based on the frequencies of the FSC curve falls at 0.5. Scale bars represent 10 nm in a, and 5 nm in b and c.

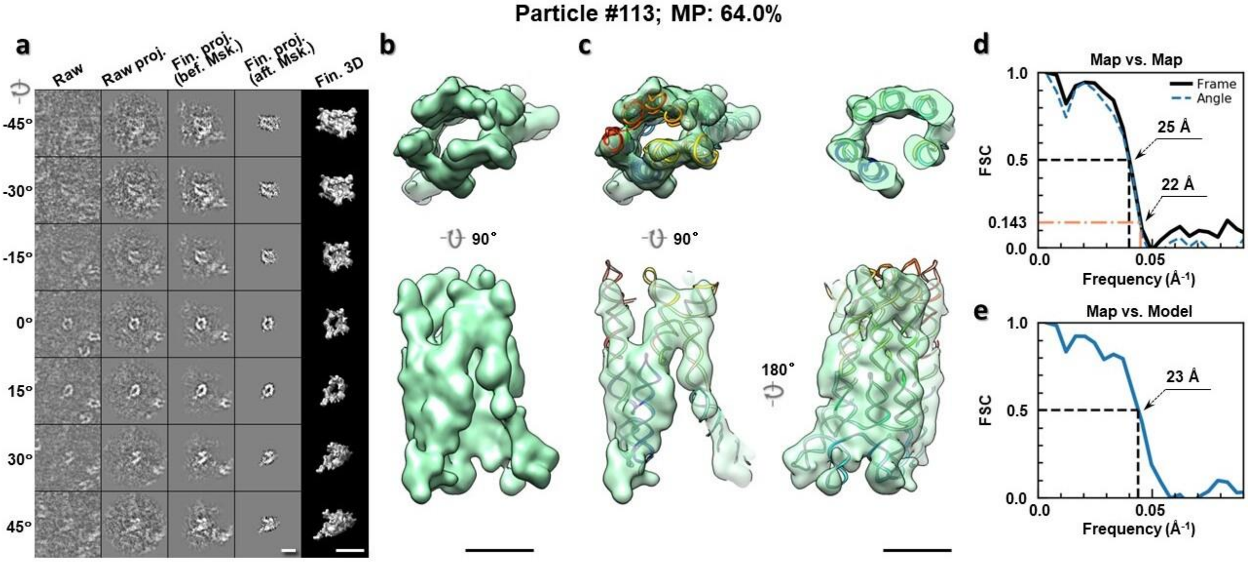

**Supplementary Fig. 83: IPET 3D reconstruction of an individual 6HBC RNA particle #113.** **a,** Seven representative tilt images of a single particle are shown in the first column from the left. The tilt images are aligned to a common center using IPET through iterative refinements. The projections of the raw, intermediate, and final 3D reconstructions at the corresponding angles are displayed in the next four

columns. **b,** Two orthogonal views of the final 3D map; **c,** Overlap of the map with the fitted model; **d,** FSC analyses of the final map resolution using two methods, "map-map FSC", in which each map is reconstructed from one half of the images, based on the even vs. odd indices of frames or tilt angles. respectively, **e,** FSC analysis of map-model, where the model map is generated by the fitted model after being low-pass filtered to 8 Å. The resolutions are assessed based on the frequencies of the FSC curve falls at 0.5. Scale bars represent 10 nm in a, and 5 nm in b and c.

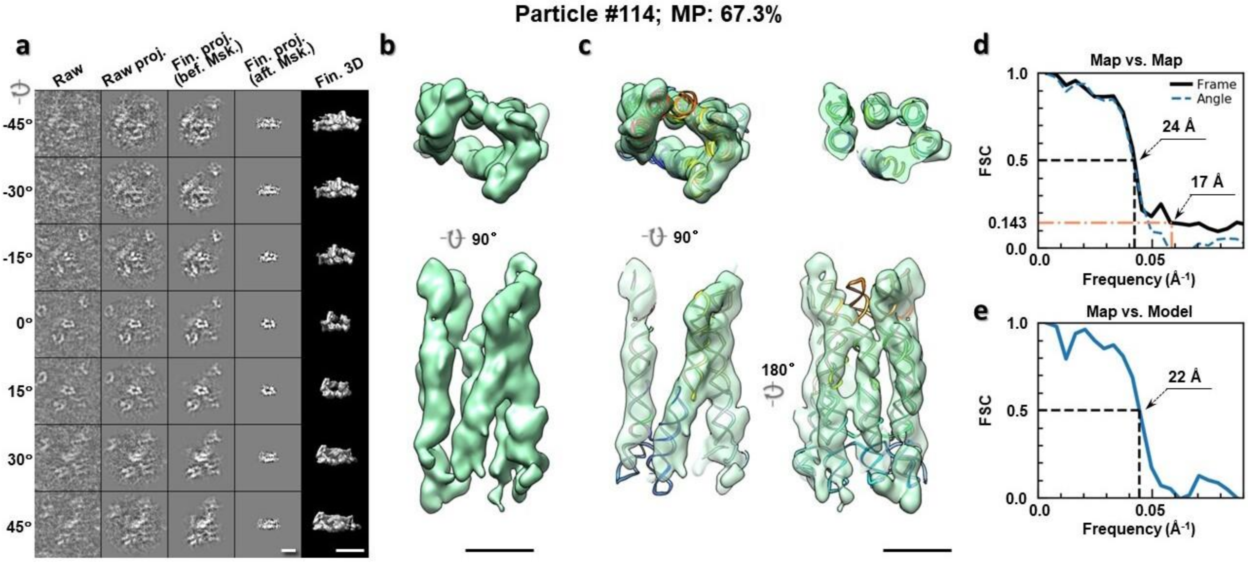

**Supplementary Fig. 84: IPET 3D reconstruction of an individual 6HBC RNA particle #114. a,** Seven representative tilt images of a single particle are shown in the first column from the left. The tilt images are aligned to a common center using IPET through iterative refinements. The projections of the raw, intermediate, and final 3D reconstructions at the corresponding angles are displayed in the next four columns. **b,** Two orthogonal views of the final 3D map; **c,** Overlap of the map with the fitted model; **d,** FSC analyses of the final map resolution using two methods, "map-map FSC", in which each map is reconstructed from one half of the images, based on the even vs. odd indices of frames or tilt angles. respectively, **e,** FSC analysis of map-model, where the model map is generated by the fitted model after being low-pass filtered to 8 Å. The resolutions are assessed based on the frequencies of the FSC curve falls at 0.5. Scale bars represent 10 nm in a, and 5 nm in b and c.

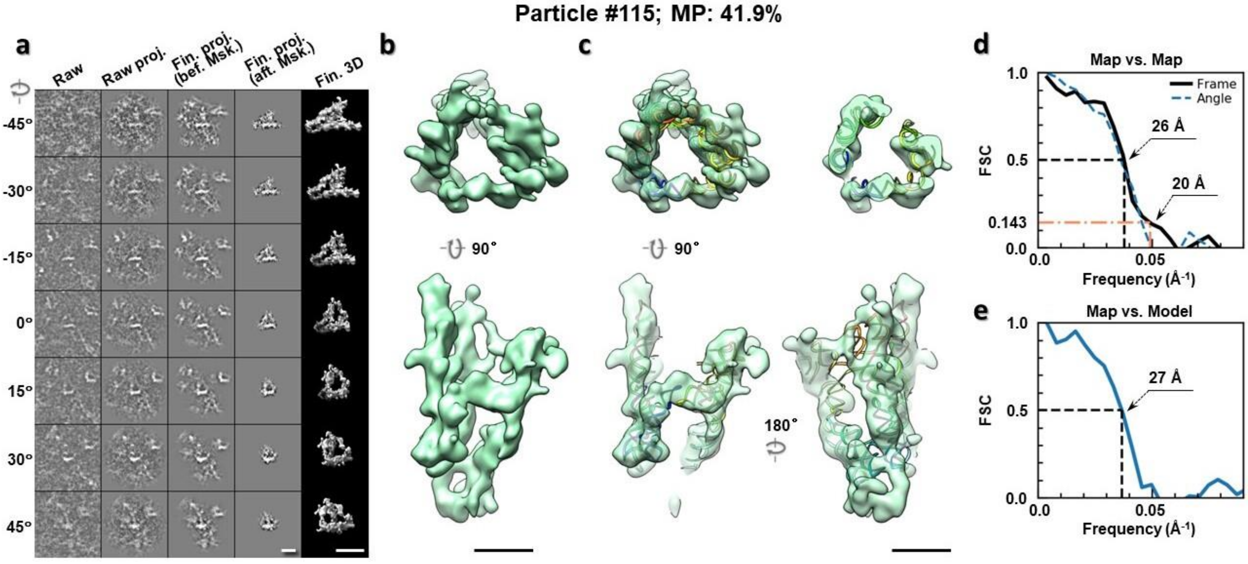

**Supplementary Fig. 85: IPET 3D reconstruction of an individual 6HBC RNA particle #115. a,** Seven representative tilt images of a single particle are shown in the first column from the left. The tilt images are aligned to a common center using IPET through iterative refinements. The projections of the raw, intermediate, and final 3D reconstructions at the corresponding angles are displayed in the next four columns. **b,** Two orthogonal views of the final 3D map; **c,** Overlap of the map with the fitted model; **d,** FSC analyses of the final map resolution using two methods, "map-map FSC", in which each map is reconstructed from one half of the images, based on the even vs. odd indices of frames or tilt angles. respectively, **e,** FSC analysis of map-model, where the model map is generated by the fitted model after being low-pass filtered to 8 Å. The resolutions are assessed based on the frequencies of the FSC curve falls at 0.5. Scale bars represent 10 nm in a, and 5 nm in b and c.

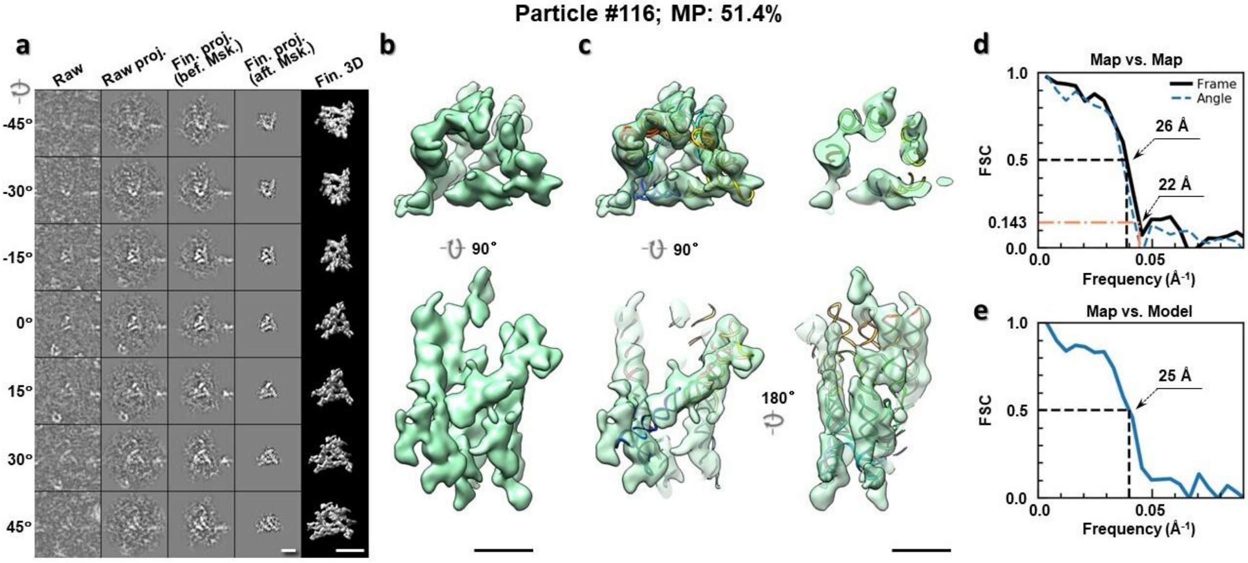

**Supplementary Fig. 86: IPET 3D reconstruction of an individual 6HBC RNA particle #116. a,** Seven representative tilt images of a single particle are shown in the first column from the left. The tilt images are aligned to a common center using IPET through iterative refinements. The projections of the raw, intermediate, and final 3D reconstructions at the corresponding angles are displayed in the next four

columns. **b,** Two orthogonal views of the final 3D map; **c,** Overlap of the map with the fitted model; **d,** FSC analyses of the final map resolution using two methods, "map-map FSC", in which each map is reconstructed from one half of the images, based on the even vs. odd indices of frames or tilt angles. respectively, **e,** FSC analysis of map-model, where the model map is generated by the fitted model after being low-pass filtered to 8 Å. The resolutions are assessed based on the frequencies of the FSC curve falls at 0.5. Scale bars represent 10 nm in a, and 5 nm in b and c.

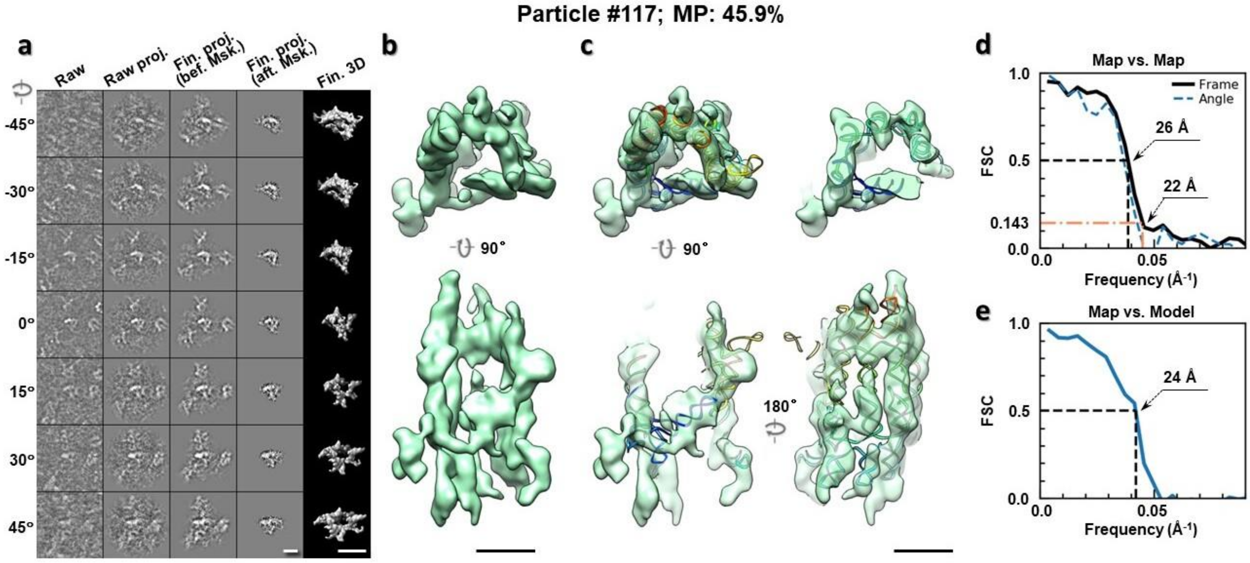

**Supplementary Fig. 87: IPET 3D reconstruction of an individual 6HBC RNA particle #117.** **a,** Seven representative tilt images of a single particle are shown in the first column from the left. The tilt images are aligned to a common center using IPET through iterative refinements. The projections of the raw, intermediate, and final 3D reconstructions at the corresponding angles are displayed in the next four columns. **b,** Two orthogonal views of the final 3D map; **c,** Overlap of the map with the fitted model; **d,** FSC analyses of the final map resolution using two methods, "map-map FSC", in which each map is reconstructed from one half of the images, based on the even vs. odd indices of frames or tilt angles. respectively, **e,** FSC analysis of map-model, where the model map is generated by the fitted model after being low-pass filtered to 8 Å. The resolutions are assessed based on the frequencies of the FSC curve falls at 0.5. Scale bars represent 10 nm in a, and 5 nm in b and c.

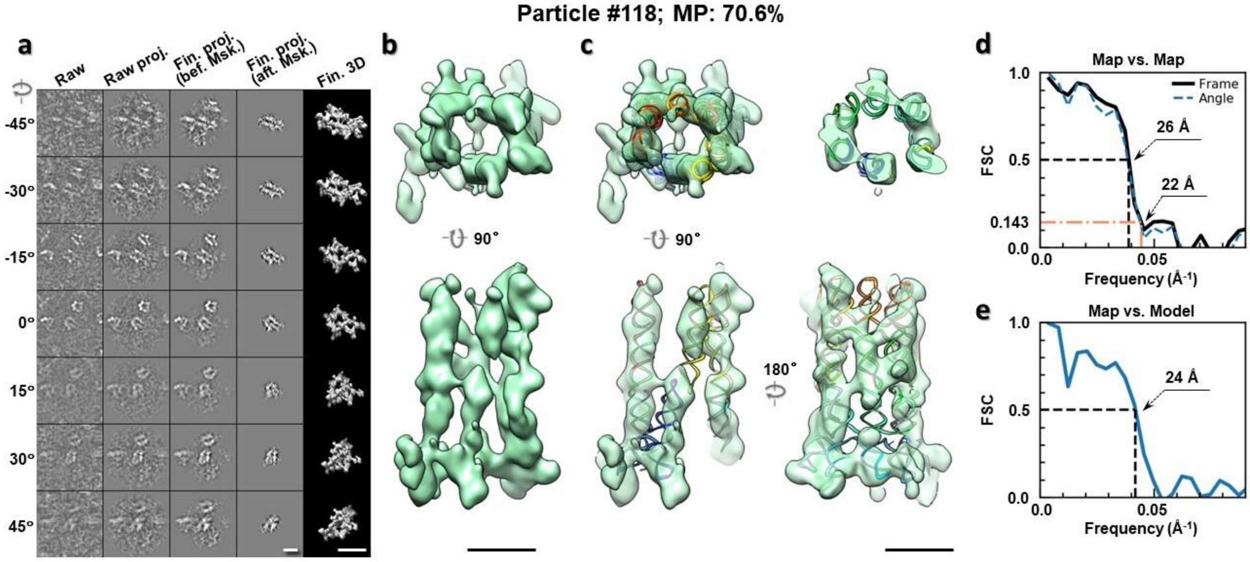

**Supplementary Fig. 88: IPET 3D reconstruction of an individual 6HBC RNA particle #118. a,** Seven representative tilt images of a single particle are shown in the first column from the left. The tilt images are aligned to a common center using IPET through iterative refinements. The projections of the raw, intermediate, and final 3D reconstructions at the corresponding angles are displayed in the next four columns. **b,** Two orthogonal views of the final 3D map; **c,** Overlap of the map with the fitted model; **d,** FSC analyses of the final map resolution using two methods, "map-map FSC", in which each map is reconstructed from one half of the images, based on the even vs. odd indices of frames or tilt angles. respectively, **e,** FSC analysis of map-model, where the model map is generated by the fitted model after being low-pass filtered to 8 Å. The resolutions are assessed based on the frequencies of the FSC curve falls at 0.5. Scale bars represent 10 nm in a, and 5 nm in b and c.

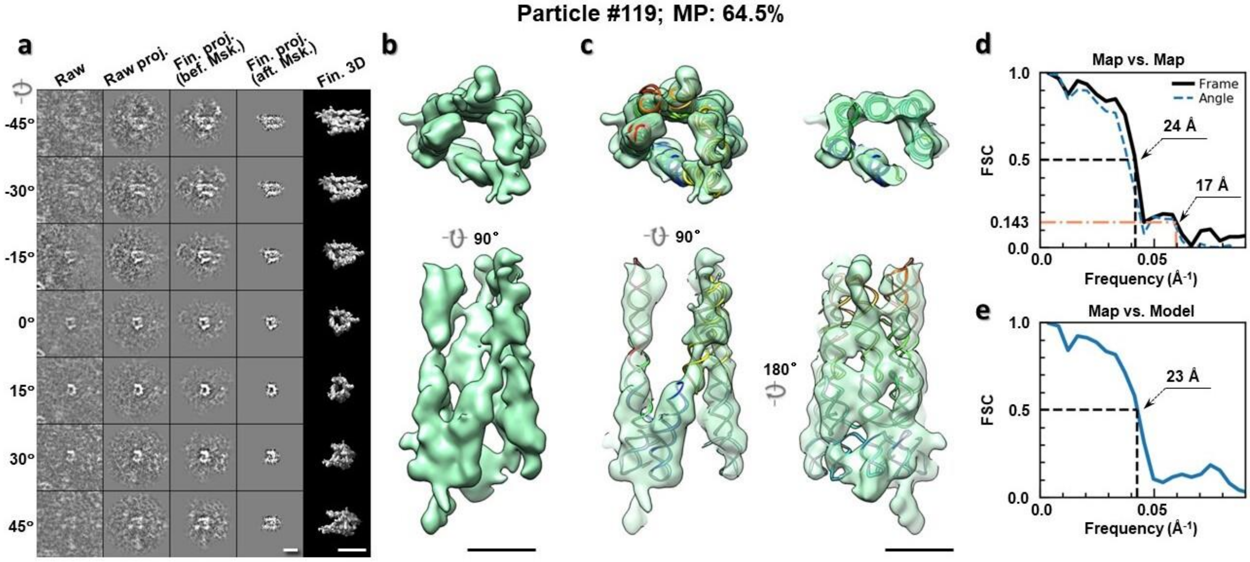

**Supplementary Fig. 89: IPET 3D reconstruction of an individual 6HBC RNA particle #119. a,** Seven representative tilt images of a single particle are shown in the first column from the left. The tilt images are aligned to a common center using IPET through iterative refinements. The projections of the raw, intermediate, and final 3D reconstructions at the corresponding angles are displayed in the next four

columns. **b,** Two orthogonal views of the final 3D map; **c,** Overlap of the map with the fitted model; **d,** FSC analyses of the final map resolution using two methods, "map-map FSC", in which each map is reconstructed from one half of the images, based on the even vs. odd indices of frames or tilt angles. respectively, **e,** FSC analysis of map-model, where the model map is generated by the fitted model after being low-pass filtered to 8 Å. The resolutions are assessed based on the frequencies of the FSC curve falls at 0.5. Scale bars represent 10 nm in a, and 5 nm in b and c.

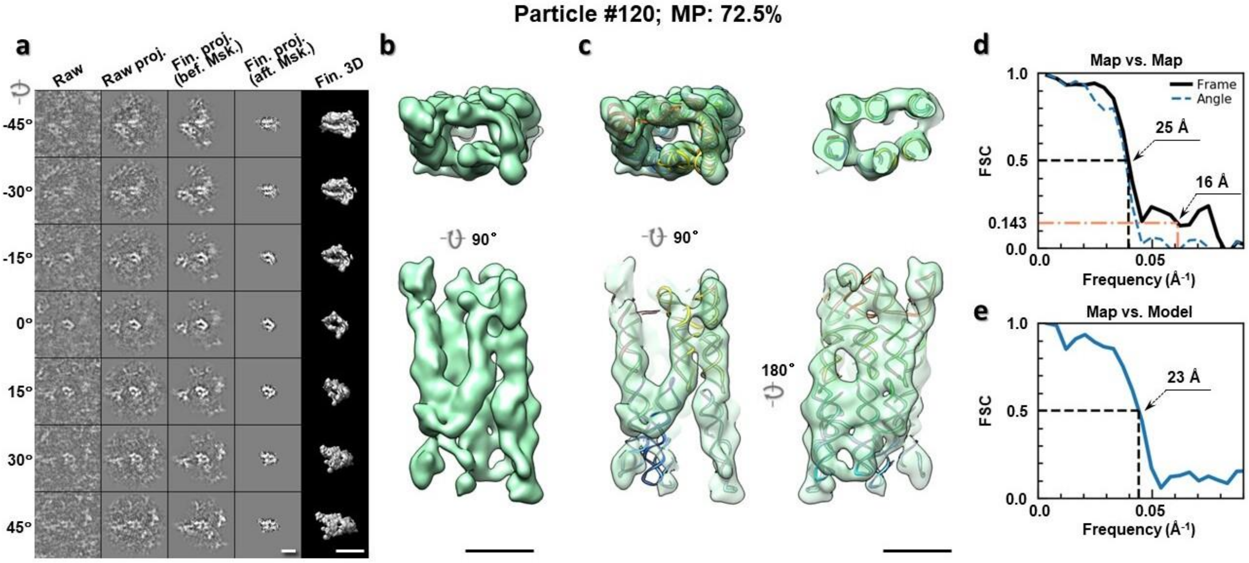

**Supplementary Fig. 90: IPET 3D reconstruction of an individual 6HBC RNA particle #120. a,** Seven representative tilt images of a single particle are shown in the first column from the left. The tilt images are aligned to a common center using IPET through iterative refinements. The projections of the raw, intermediate, and final 3D reconstructions at the corresponding angles are displayed in the next four columns. **b,** Two orthogonal views of the final 3D map; **c,** Overlap of the map with the fitted model; **d,** FSC analyses of the final map resolution using two methods, "map-map FSC", in which each map is reconstructed from one half of the images, based on the even vs. odd indices of frames or tilt angles. respectively, **e,** FSC analysis of map-model, where the model map is generated by the fitted model after being low-pass filtered to 8 Å. The resolutions are assessed based on the frequencies of the FSC curve falls at 0.5. Scale bars represent 10 nm in a, and 5 nm in b and c.

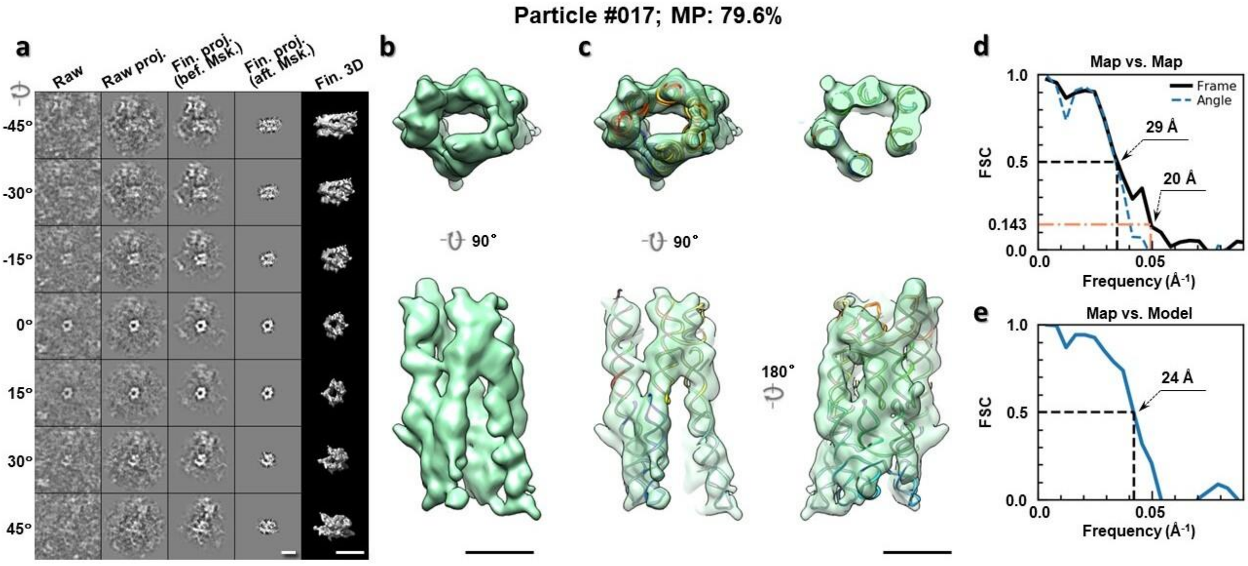

**Supplementary Fig. 91: IPET 3D reconstruction of an individual 6HBC RNA particle #017. a,** Seven representative tilt images of a single particle are shown in the first column from the left. The tilt images are aligned to a common center using IPET through iterative refinements. The projections of the raw, intermediate, and final 3D reconstructions at the corresponding angles are displayed in the next four columns. **b,** Two orthogonal views of the final 3D map; **c,** Overlap of the map with the fitted model; **d,** FSC analyses of the final map resolution using two methods, "map-map FSC", in which each map is reconstructed from one half of the images, based on the even vs. odd indices of frames or tilt angles. respectively, **e,** FSC analysis of map-model, where the model map is generated by the fitted model after being low-pass filtered to 8 Å. The resolutions are assessed based on the frequencies of the FSC curve falls at 0.5. Scale bars represent 10 nm in a, and 5 nm in b and c.

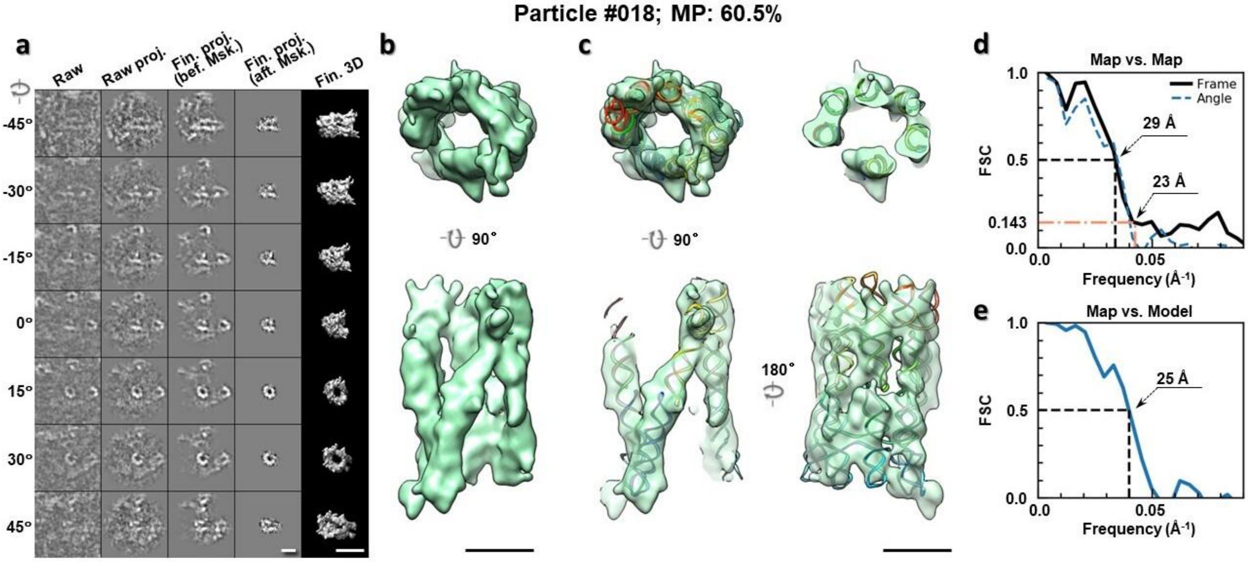

**Supplementary Fig. 92: IPET 3D reconstruction of an individual 6HBC RNA particle #018. a,** Seven representative tilt images of a single particle are shown in the first column from the left. The tilt images are aligned to a common center using IPET through iterative refinements. The projections of the raw, intermediate, and final 3D reconstructions at the corresponding angles are displayed in the next four

columns. **b,** Two orthogonal views of the final 3D map; **c,** Overlap of the map with the fitted model; **d,** FSC analyses of the final map resolution using two methods, "map-map FSC", in which each map is reconstructed from one half of the images, based on the even vs. odd indices of frames or tilt angles. respectively, **e,** FSC analysis of map-model, where the model map is generated by the fitted model after being low-pass filtered to 8 Å. The resolutions are assessed based on the frequencies of the FSC curve falls at 0.5. Scale bars represent 10 nm in a, and 5 nm in b and c.

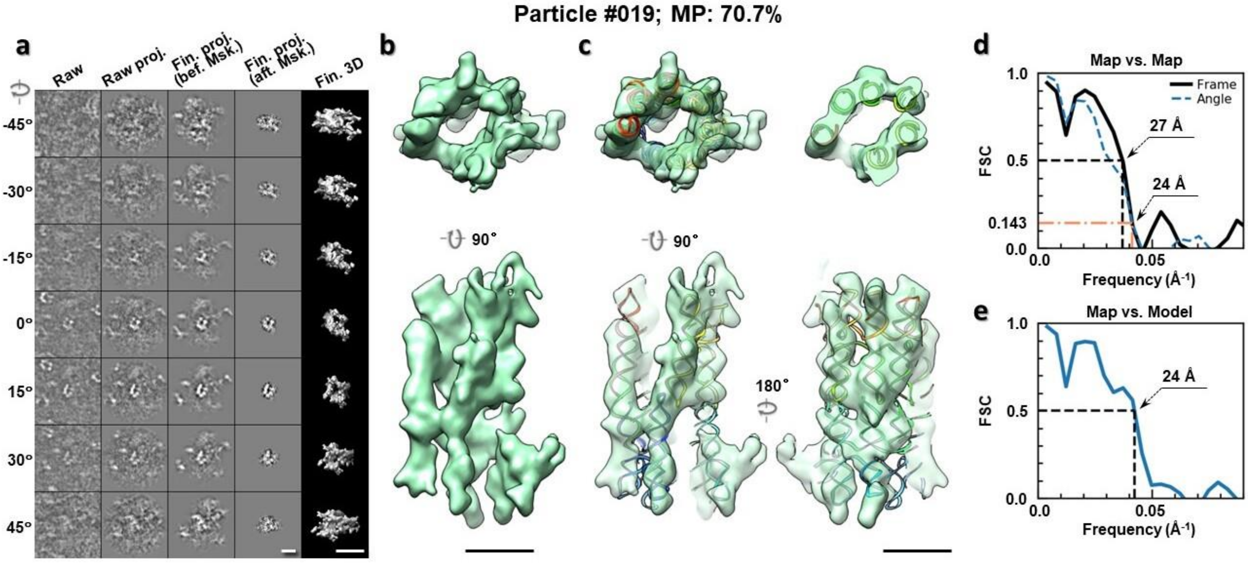

**Supplementary Fig. 93: IPET 3D reconstruction of an individual 6HBC RNA particle #019. a,** Seven representative tilt images of a single particle are shown in the first column from the left. The tilt images are aligned to a common center using IPET through iterative refinements. The projections of the raw, intermediate, and final 3D reconstructions at the corresponding angles are displayed in the next four columns. **b,** Two orthogonal views of the final 3D map; **c,** Overlap of the map with the fitted model; **d,** FSC analyses of the final map resolution using two methods, "map-map FSC", in which each map is reconstructed from one half of the images, based on the even vs. odd indices of frames or tilt angles. respectively, **e,** FSC analysis of map-model, where the model map is generated by the fitted model after being low-pass filtered to 8 Å. The resolutions are assessed based on the frequencies of the FSC curve falls at 0.5. Scale bars represent 10 nm in a, and 5 nm in b and c.

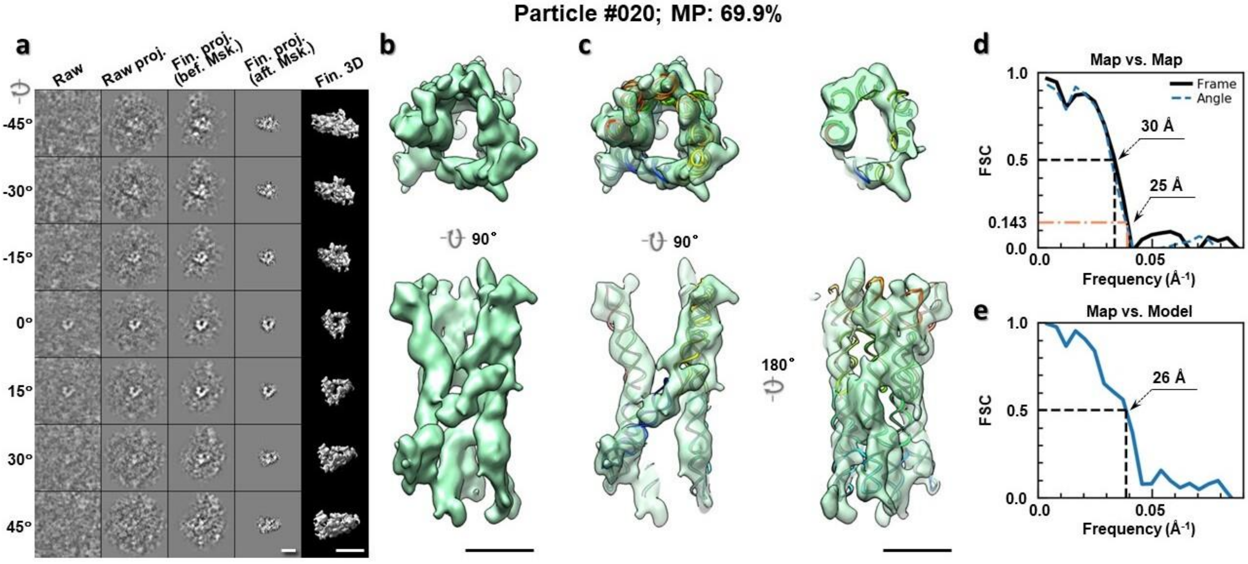

**Supplementary Fig. 94: IPET 3D reconstruction of an individual 6HBC RNA particle #020. a,** Seven representative tilt images of a single particle are shown in the first column from the left. The tilt images are aligned to a common center using IPET through iterative refinements. The projections of the raw, intermediate, and final 3D reconstructions at the corresponding angles are displayed in the next four columns. **b,** Two orthogonal views of the final 3D map; **c,** Overlap of the map with the fitted model; **d,** FSC analyses of the final map resolution using two methods, "map-map FSC", in which each map is reconstructed from one half of the images, based on the even vs. odd indices of frames or tilt angles. respectively, **e,** FSC analysis of map-model, where the model map is generated by the fitted model after being low-pass filtered to 8 Å. The resolutions are assessed based on the frequencies of the FSC curve falls at 0.5. Scale bars represent 10 nm in a, and 5 nm in b and c.

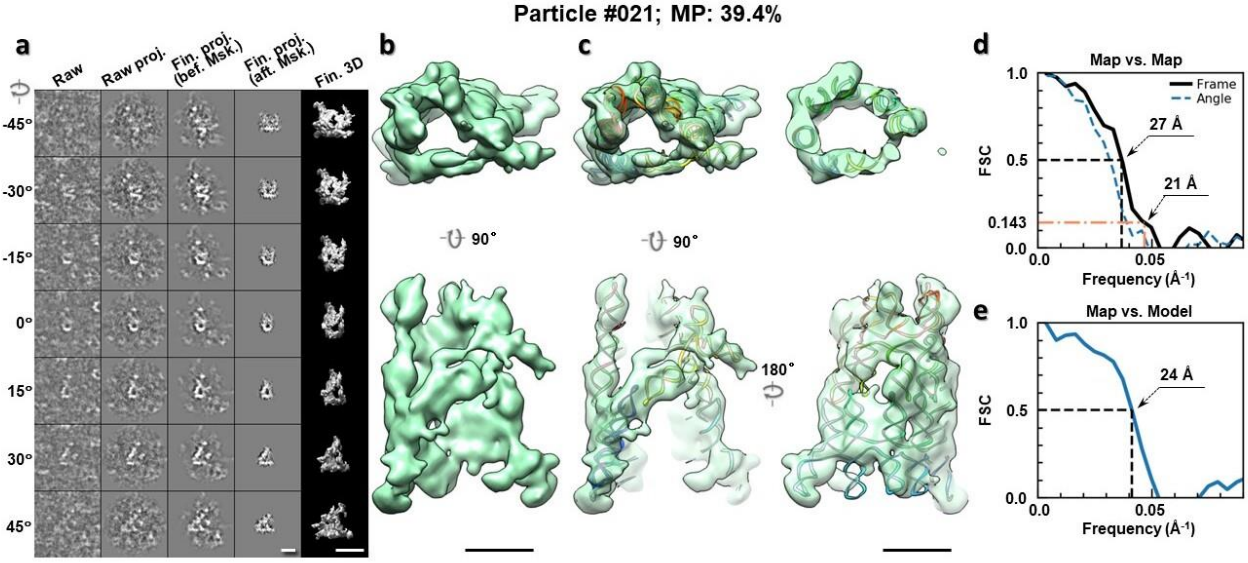

**Supplementary Fig. 95: IPET 3D reconstruction of an individual 6HBC RNA particle #021. a,** Seven representative tilt images of a single particle are shown in the first column from the left. The tilt images are aligned to a common center using IPET through iterative refinements. The projections of the raw, intermediate, and final 3D reconstructions at the corresponding angles are displayed in the next four

columns. **b,** Two orthogonal views of the final 3D map; **c,** Overlap of the map with the fitted model; **d,** FSC analyses of the final map resolution using two methods, "map-map FSC", in which each map is reconstructed from one half of the images, based on the even vs. odd indices of frames or tilt angles. respectively, **e,** FSC analysis of map-model, where the model map is generated by the fitted model after being low-pass filtered to 8 Å. The resolutions are assessed based on the frequencies of the FSC curve falls at 0.5. Scale bars represent 10 nm in a, and 5 nm in b and c.

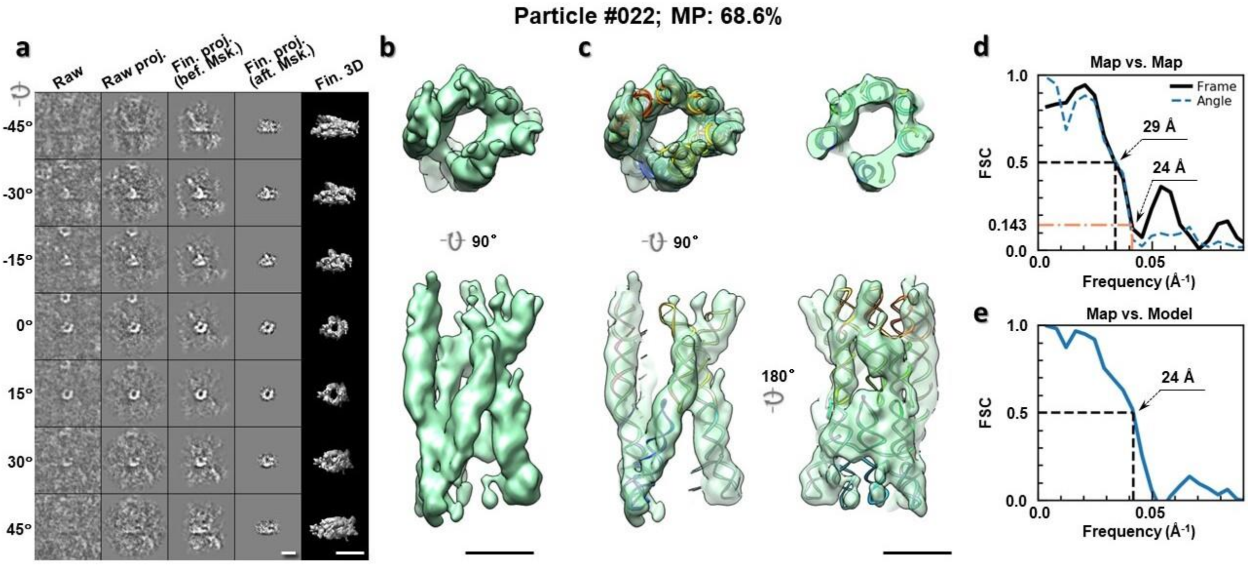

**Supplementary Fig. 96: IPET 3D reconstruction of an individual 6HBC RNA particle #022.** **a,** Seven representative tilt images of a single particle are shown in the first column from the left. The tilt images are aligned to a common center using IPET through iterative refinements. The projections of the raw, intermediate, and final 3D reconstructions at the corresponding angles are displayed in the next four columns. **b,** Two orthogonal views of the final 3D map; **c,** Overlap of the map with the fitted model; **d,** FSC analyses of the final map resolution using two methods, "map-map FSC", in which each map is reconstructed from one half of the images, based on the even vs. odd indices of frames or tilt angles. respectively, **e,** FSC analysis of map-model, where the model map is generated by the fitted model after being low-pass filtered to 8 Å. The resolutions are assessed based on the frequencies of the FSC curve falls at 0.5. Scale bars represent 10 nm in a, and 5 nm in b and c.

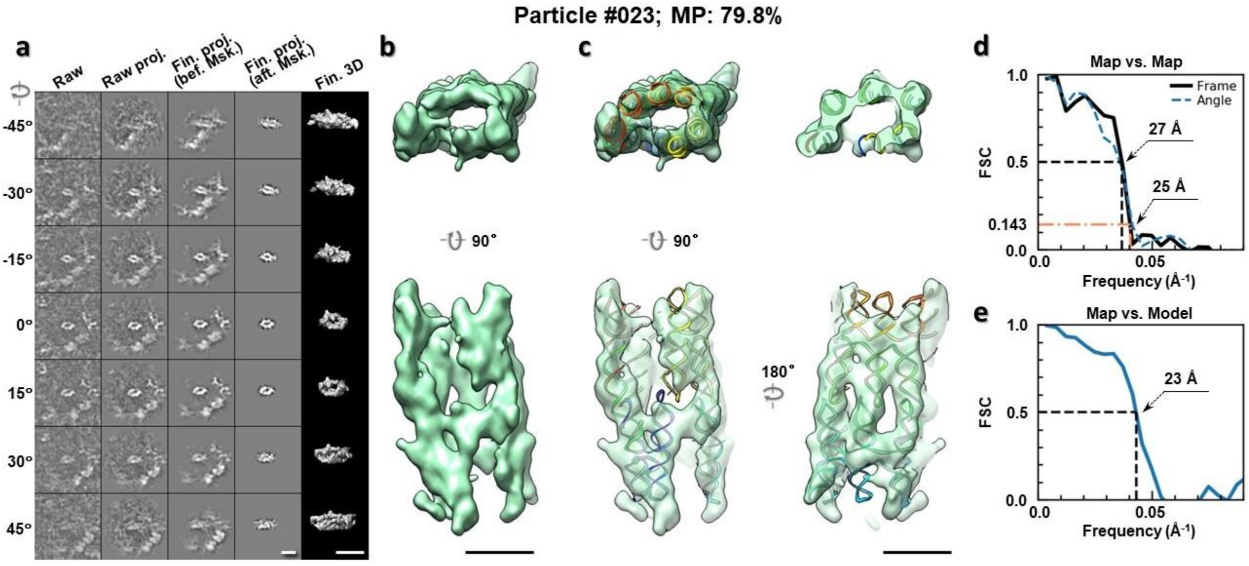

**Supplementary Fig. 97: IPET 3D reconstruction of an individual 6HBC RNA particle #023. a,** Seven representative tilt images of a single particle are shown in the first column from the left. The tilt images are aligned to a common center using IPET through iterative refinements. The projections of the raw, intermediate, and final 3D reconstructions at the corresponding angles are displayed in the next four columns. **b,** Two orthogonal views of the final 3D map; **c,** Overlap of the map with the fitted model; **d,** FSC analyses of the final map resolution using two methods, "map-map FSC", in which each map is reconstructed from one half of the images, based on the even vs. odd indices of frames or tilt angles. respectively, **e,** FSC analysis of map-model, where the model map is generated by the fitted model after being low-pass filtered to 8 Å. The resolutions are assessed based on the frequencies of the FSC curve falls at 0.5. Scale bars represent 10 nm in a, and 5 nm in b and c.

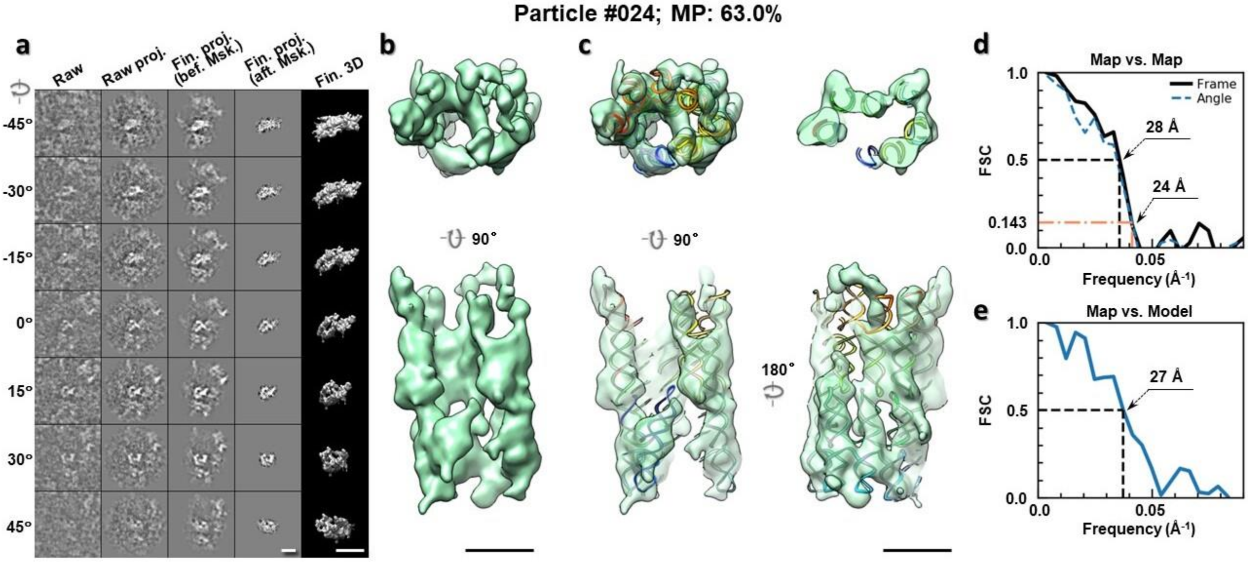

**Supplementary Fig. 98: IPET 3D reconstruction of an individual 6HBC RNA particle #024. a,** Seven representative tilt images of a single particle are shown in the first column from the left. The tilt images are aligned to a common center using IPET through iterative refinements. The projections of the raw, intermediate, and final 3D reconstructions at the corresponding angles are displayed in the next four

columns. **b,** Two orthogonal views of the final 3D map; **c,** Overlap of the map with the fitted model; **d,** FSC analyses of the final map resolution using two methods, "map-map FSC", in which each map is reconstructed from one half of the images, based on the even vs. odd indices of frames or tilt angles. respectively, **e,** FSC analysis of map-model, where the model map is generated by the fitted model after being low-pass filtered to 8 Å. The resolutions are assessed based on the frequencies of the FSC curve falls at 0.5. Scale bars represent 10 nm in a, and 5 nm in b and c.

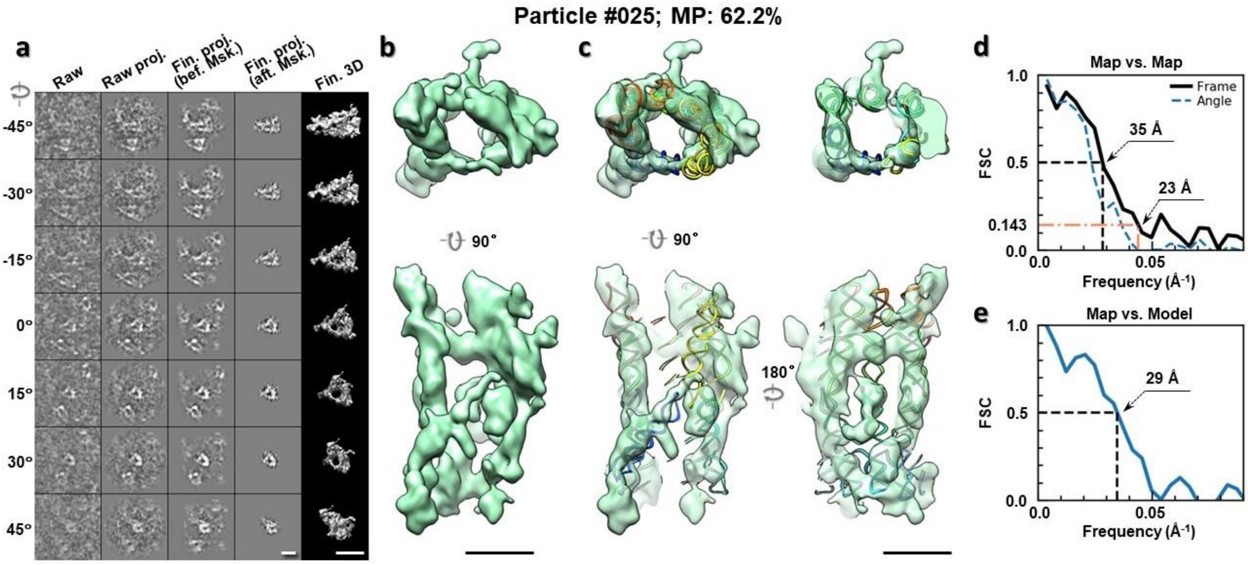

**Supplementary Fig. 99: IPET 3D reconstruction of an individual 6HBC RNA particle #025. a,** Seven representative tilt images of a single particle are shown in the first column from the left. The tilt images are aligned to a common center using IPET through iterative refinements. The projections of the raw, intermediate, and final 3D reconstructions at the corresponding angles are displayed in the next four columns. **b,** Two orthogonal views of the final 3D map; **c,** Overlap of the map with the fitted model; **d,** FSC analyses of the final map resolution using two methods, "map-map FSC", in which each map is reconstructed from one half of the images, based on the even vs. odd indices of frames or tilt angles. respectively, **e,** FSC analysis of map-model, where the model map is generated by the fitted model after being low-pass filtered to 8 Å. The resolutions are assessed based on the frequencies of the FSC curve falls at 0.5. Scale bars represent 10 nm in a, and 5 nm in b and c.

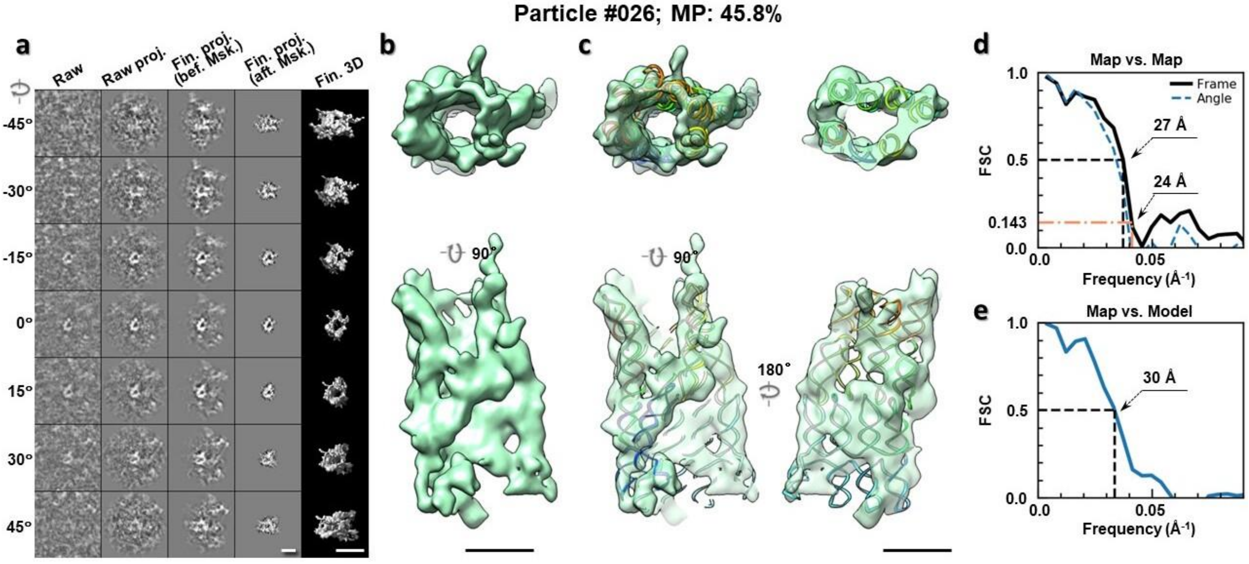

**Supplementary Fig. 100: IPET 3D reconstruction of an individual 6HBC RNA particle #026. a,** Seven representative tilt images of a single particle are shown in the first column from the left. The tilt images are aligned to a common center using IPET through iterative refinements. The projections of the raw, intermediate, and final 3D reconstructions at the corresponding angles are displayed in the next four columns. **b,** Two orthogonal views of the final 3D map; **c,** Overlap of the map with the fitted model; **d,** FSC analyses of the final map resolution using two methods, "map-map FSC", in which each map is reconstructed from one half of the images, based on the even vs. odd indices of frames or tilt angles. respectively, **e,** FSC analysis of map-model, where the model map is generated by the fitted model after being low-pass filtered to 8 Å. The resolutions are assessed based on the frequencies of the FSC curve falls at 0.5. Scale bars represent 10 nm in a, and 5 nm in b and c.

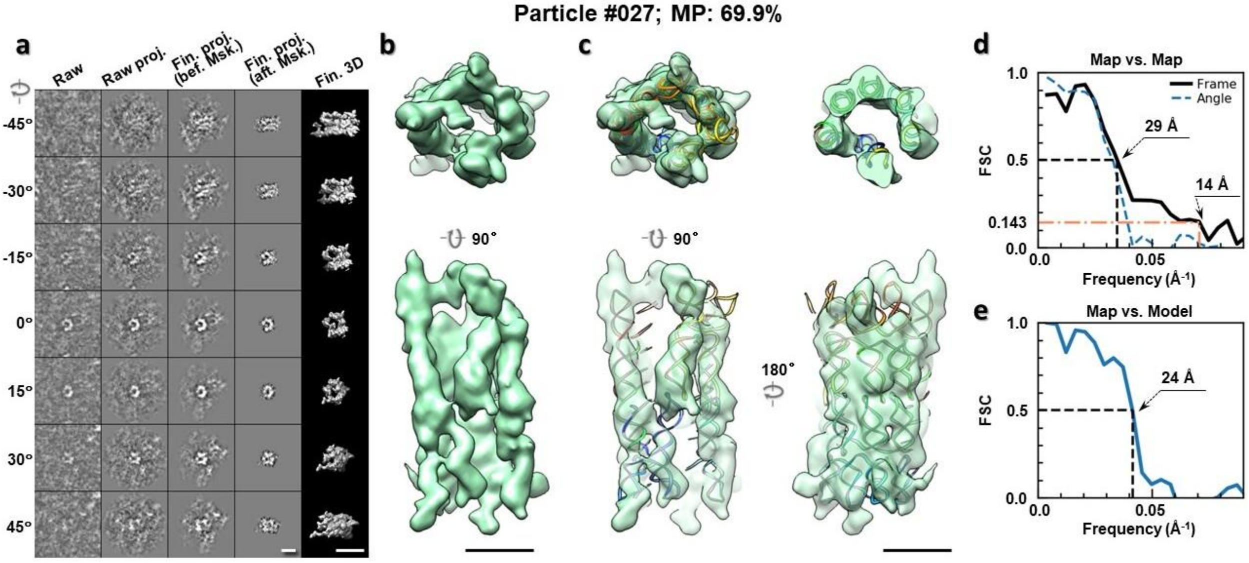

**Supplementary Fig. 101: IPET 3D reconstruction of an individual 6HBC RNA particle #027. a,** Seven representative tilt images of a single particle are shown in the first column from the left. The tilt images are aligned to a common center using IPET through iterative refinements. The projections of the raw, intermediate, and final 3D reconstructions at the corresponding angles are displayed in the next four

columns. **b,** Two orthogonal views of the final 3D map; **c,** Overlap of the map with the fitted model; **d,** FSC analyses of the final map resolution using two methods, "map-map FSC", in which each map is reconstructed from one half of the images, based on the even vs. odd indices of frames or tilt angles. respectively, **e,** FSC analysis of map-model, where the model map is generated by the fitted model after being low-pass filtered to 8 Å. The resolutions are assessed based on the frequencies of the FSC curve falls at 0.5. Scale bars represent 10 nm in a, and 5 nm in b and c.

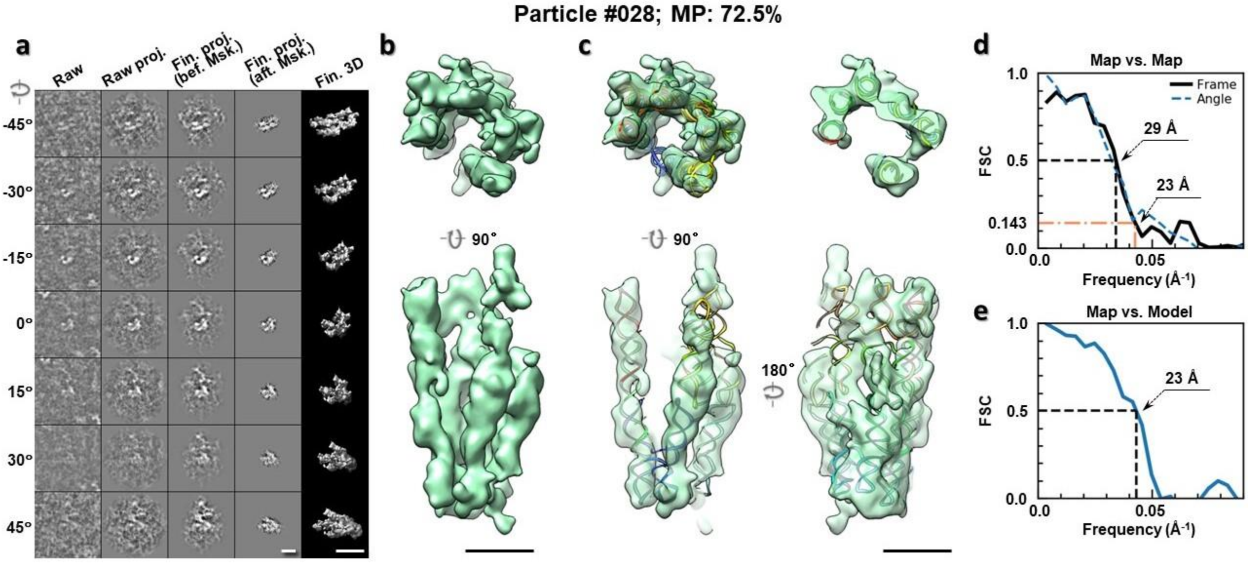

**Supplementary Fig. 102: IPET 3D reconstruction of an individual 6HBC RNA particle #028. a,** Seven representative tilt images of a single particle are shown in the first column from the left. The tilt images are aligned to a common center using IPET through iterative refinements. The projections of the raw, intermediate, and final 3D reconstructions at the corresponding angles are displayed in the next four columns. **b,** Two orthogonal views of the final 3D map; **c,** Overlap of the map with the fitted model; **d,** FSC analyses of the final map resolution using two methods, "map-map FSC", in which each map is reconstructed from one half of the images, based on the even vs. odd indices of frames or tilt angles. respectively, **e,** FSC analysis of map-model, where the model map is generated by the fitted model after being low-pass filtered to 8 Å. The resolutions are assessed based on the frequencies of the FSC curve falls at 0.5. Scale bars represent 10 nm in a, and 5 nm in b and c.

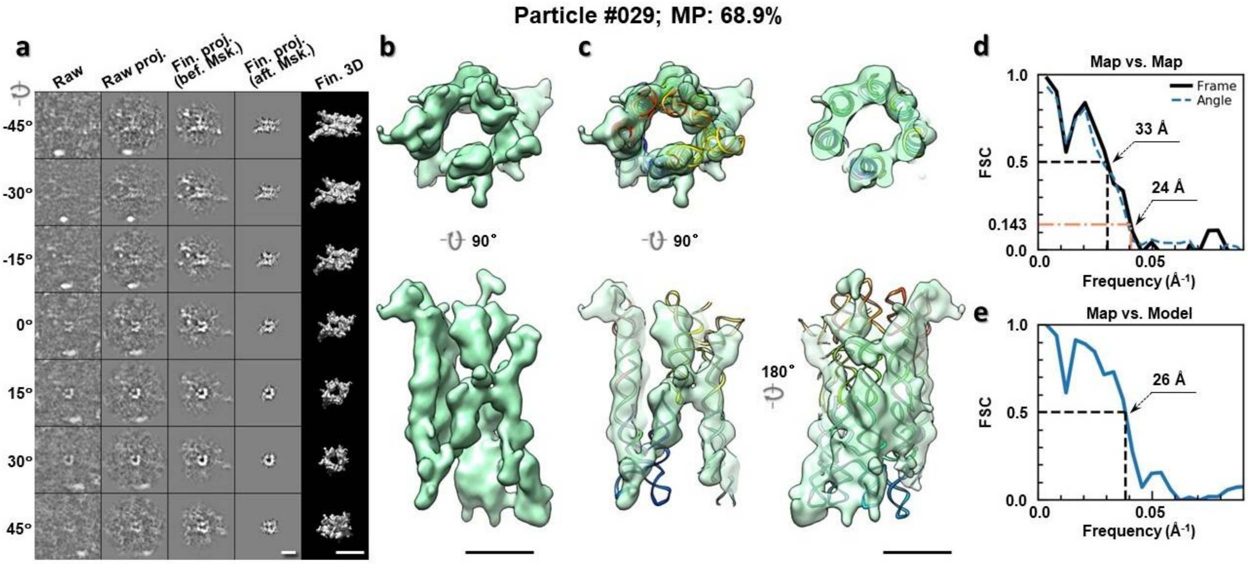

**Supplementary Fig. 103: IPET 3D reconstruction of an individual 6HBC RNA particle #029. a,** Seven representative tilt images of a single particle are shown in the first column from the left. The tilt images are aligned to a common center using IPET through iterative refinements. The projections of the raw, intermediate, and final 3D reconstructions at the corresponding angles are displayed in the next four columns. **b,** Two orthogonal views of the final 3D map; **c,** Overlap of the map with the fitted model; **d,** FSC analyses of the final map resolution using two methods, "map-map FSC", in which each map is reconstructed from one half of the images, based on the even vs. odd indices of frames or tilt angles. respectively, **e,** FSC analysis of map-model, where the model map is generated by the fitted model after being low-pass filtered to 8 Å. The resolutions are assessed based on the frequencies of the FSC curve falls at 0.5. Scale bars represent 10 nm in a, and 5 nm in b and c.

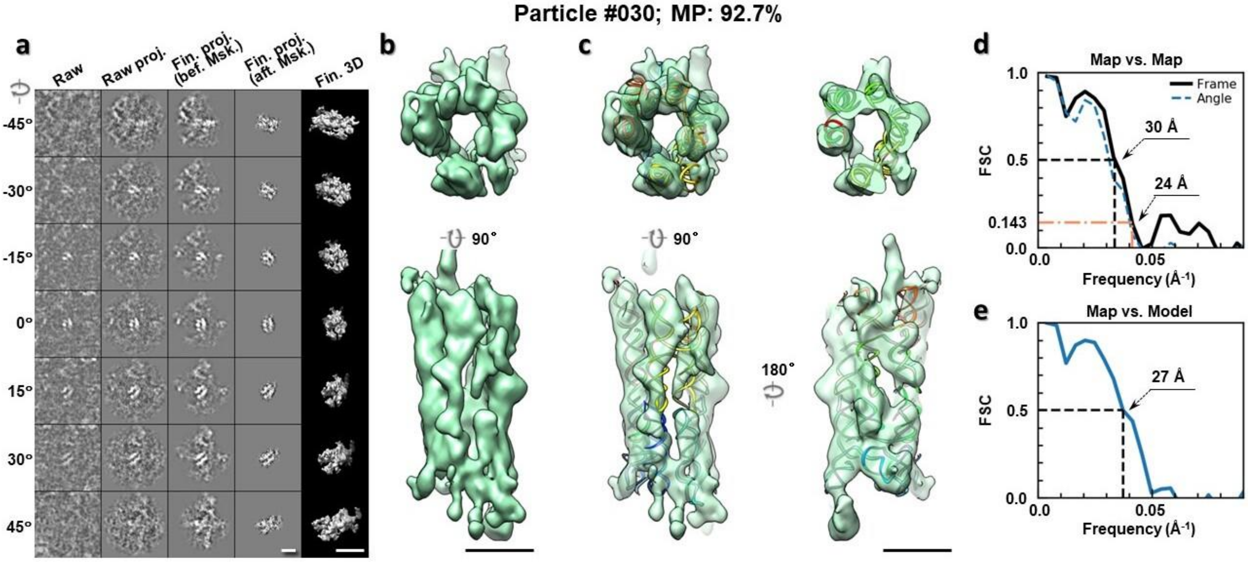

**Supplementary Fig. 104: IPET 3D reconstruction of an individual 6HBC RNA particle #030. a,** Seven representative tilt images of a single particle are shown in the first column from the left. The tilt images are aligned to a common center using IPET through iterative refinements. The projections of the raw, intermediate, and final 3D reconstructions at the corresponding angles are displayed in the next four

columns. **b,** Two orthogonal views of the final 3D map; **c,** Overlap of the map with the fitted model; **d,** FSC analyses of the final map resolution using two methods, "map-map FSC", in which each map is reconstructed from one half of the images, based on the even vs. odd indices of frames or tilt angles. respectively, **e,** FSC analysis of map-model, where the model map is generated by the fitted model after being low-pass filtered to 8 Å. The resolutions are assessed based on the frequencies of the FSC curve falls at 0.5. Scale bars represent 10 nm in a, and 5 nm in b and c.

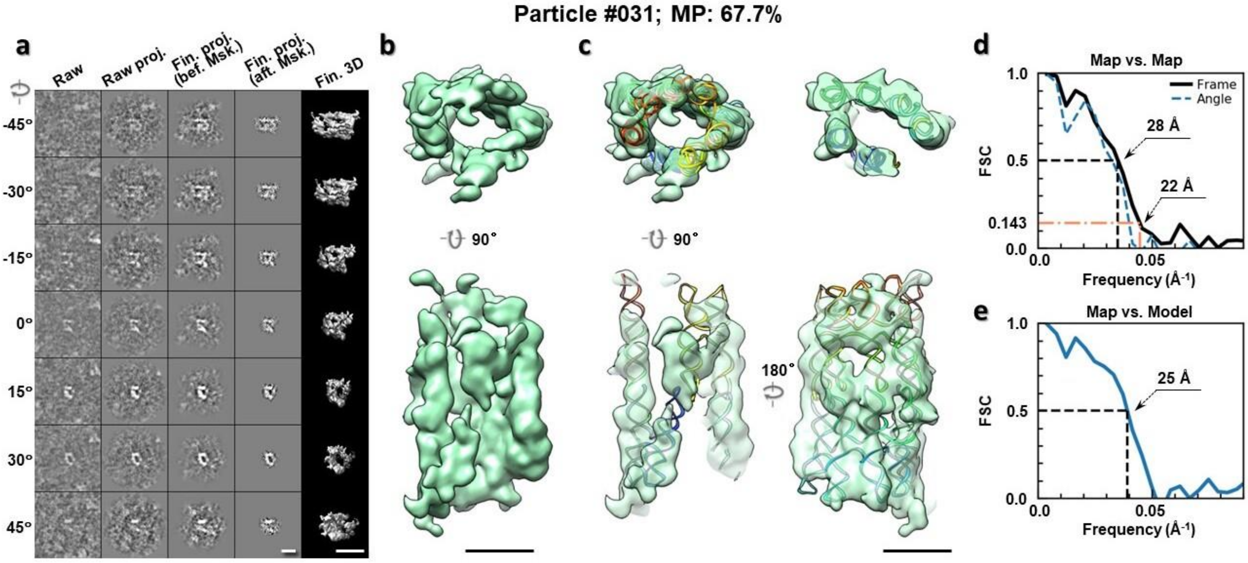

**Supplementary Fig. 105: IPET 3D reconstruction of an individual 6HBC RNA particle #031. a,** Seven representative tilt images of a single particle are shown in the first column from the left. The tilt images are aligned to a common center using IPET through iterative refinements. The projections of the raw, intermediate, and final 3D reconstructions at the corresponding angles are displayed in the next four columns. **b,** Two orthogonal views of the final 3D map; **c,** Overlap of the map with the fitted model; **d,** FSC analyses of the final map resolution using two methods, "map-map FSC", in which each map is reconstructed from one half of the images, based on the even vs. odd indices of frames or tilt angles. respectively, **e,** FSC analysis of map-model, where the model map is generated by the fitted model after being low-pass filtered to 8 Å. The resolutions are assessed based on the frequencies of the FSC curve falls at 0.5. Scale bars represent 10 nm in a, and 5 nm in b and c.

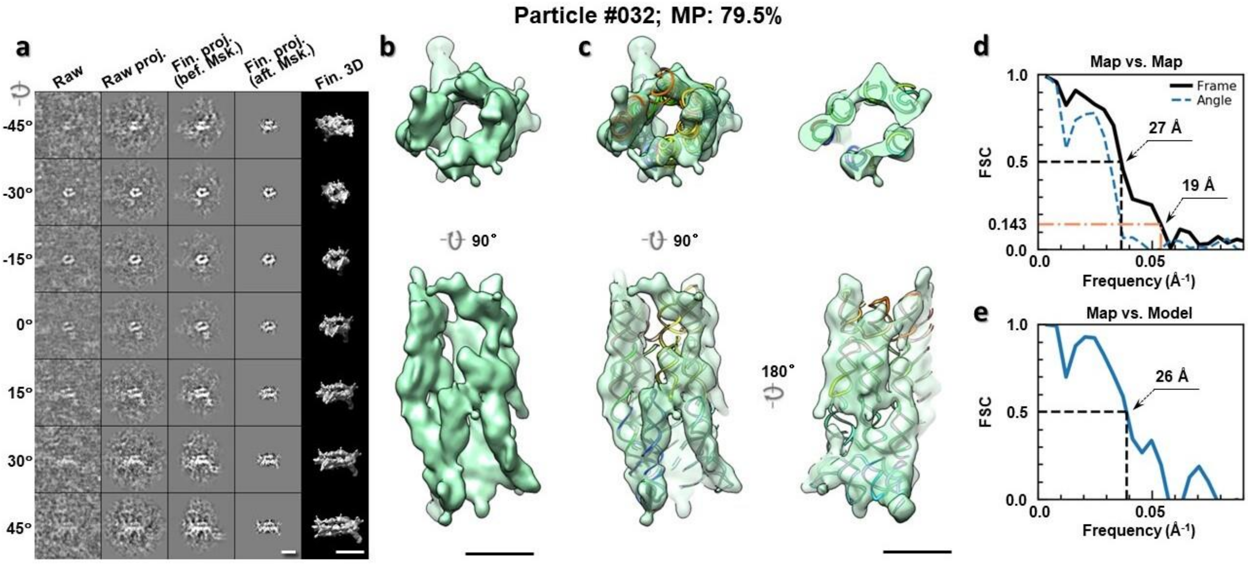

**Supplementary Fig. 106: IPET 3D reconstruction of an individual 6HBC RNA particle #032. a,** Seven representative tilt images of a single particle are shown in the first column from the left. The tilt images are aligned to a common center using IPET through iterative refinements. The projections of the raw, intermediate, and final 3D reconstructions at the corresponding angles are displayed in the next four columns. **b,** Two orthogonal views of the final 3D map; **c,** Overlap of the map with the fitted model; **d,** FSC analyses of the final map resolution using two methods, "map-map FSC", in which each map is reconstructed from one half of the images, based on the even vs. odd indices of frames or tilt angles. respectively, **e,** FSC analysis of map-model, where the model map is generated by the fitted model after being low-pass filtered to 8 Å. The resolutions are assessed based on the frequencies of the FSC curve falls at 0.5. Scale bars represent 10 nm in a, and 5 nm in b and c.

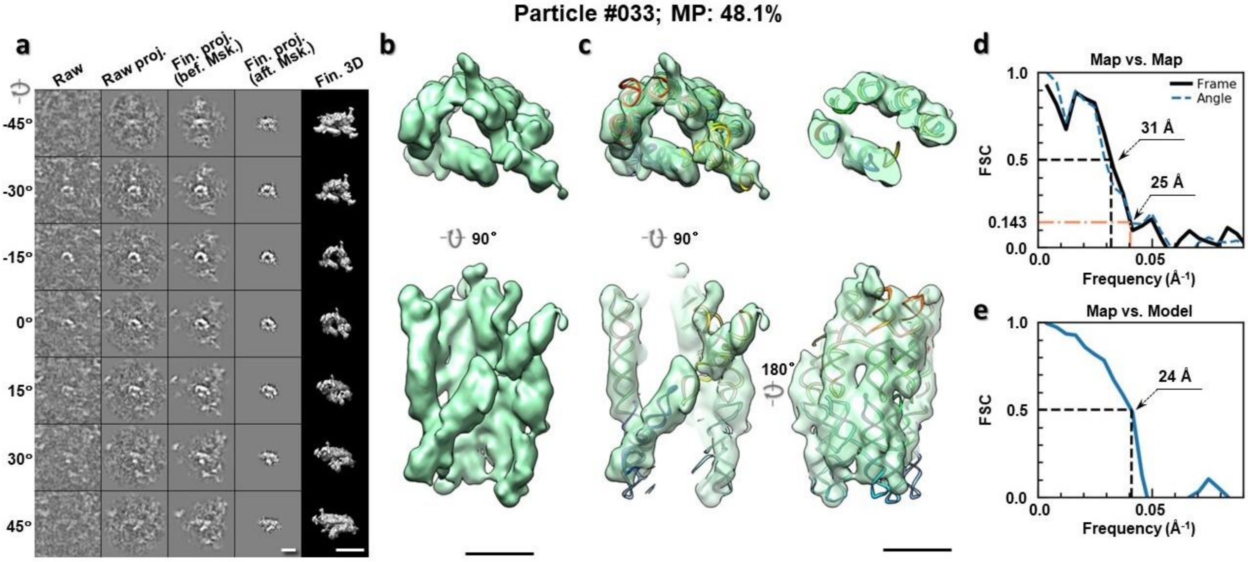

**Supplementary Fig. 107: IPET 3D reconstruction of an individual 6HBC RNA particle #033. a,** Seven representative tilt images of a single particle are shown in the first column from the left. The tilt images are aligned to a common center using IPET through iterative refinements. The projections of the raw, intermediate, and final 3D reconstructions at the corresponding angles are displayed in the next four

columns. **b,** Two orthogonal views of the final 3D map; **c,** Overlap of the map with the fitted model; **d,** FSC analyses of the final map resolution using two methods, "map-map FSC", in which each map is reconstructed from one half of the images, based on the even vs. odd indices of frames or tilt angles. respectively, **e,** FSC analysis of map-model, where the model map is generated by the fitted model after being low-pass filtered to 8 Å. The resolutions are assessed based on the frequencies of the FSC curve falls at 0.5. Scale bars represent 10 nm in a, and 5 nm in b and c.

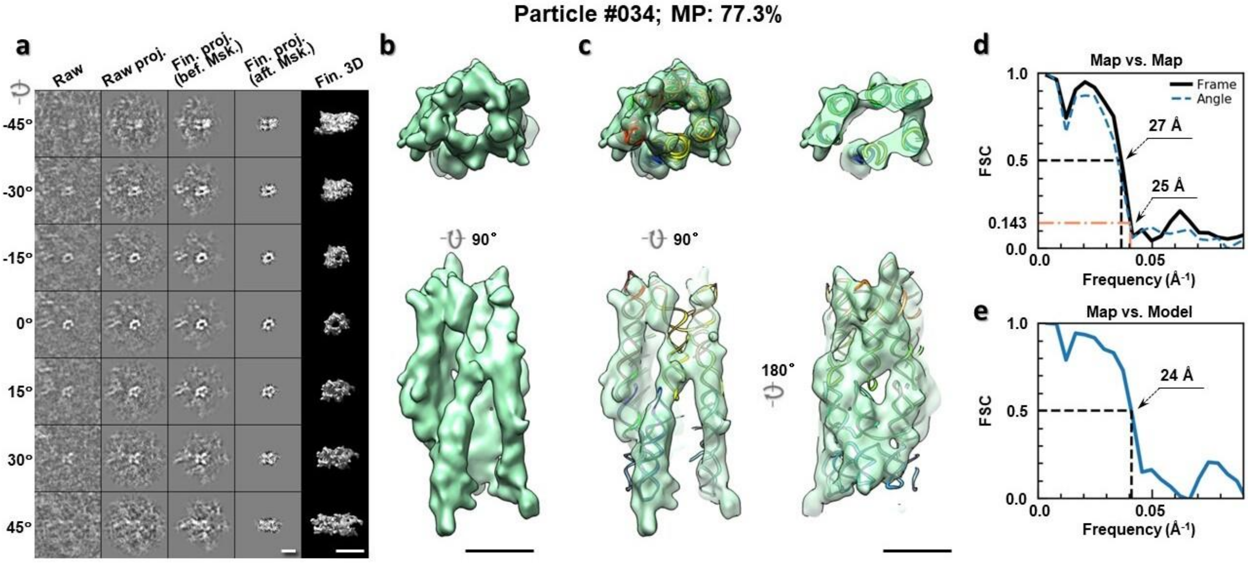

**Supplementary Fig. 108: IPET 3D reconstruction of an individual 6HBC RNA particle #034.** **a,** Seven representative tilt images of a single particle are shown in the first column from the left. The tilt images are aligned to a common center using IPET through iterative refinements. The projections of the raw, intermediate, and final 3D reconstructions at the corresponding angles are displayed in the next four columns. **b,** Two orthogonal views of the final 3D map; **c,** Overlap of the map with the fitted model; **d,** FSC analyses of the final map resolution using two methods, "map-map FSC", in which each map is reconstructed from one half of the images, based on the even vs. odd indices of frames or tilt angles. respectively, **e,** FSC analysis of map-model, where the model map is generated by the fitted model after being low-pass filtered to 8 Å. The resolutions are assessed based on the frequencies of the FSC curve falls at 0.5. Scale bars represent 10 nm in a, and 5 nm in b and c.

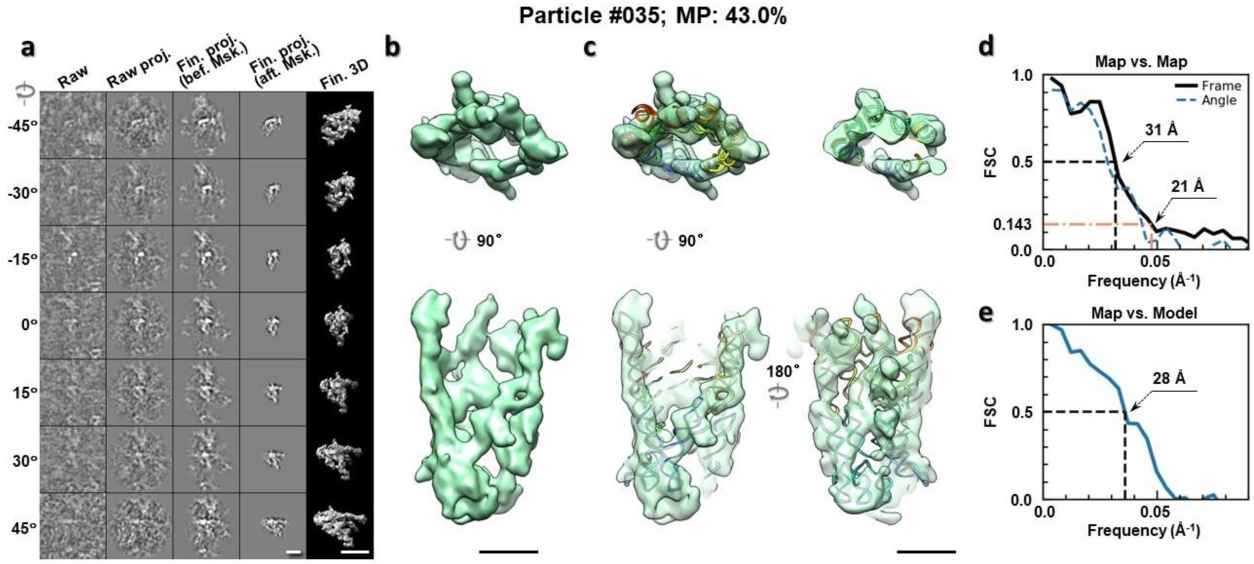

**Supplementary Fig. 109: IPET 3D reconstruction of an individual 6HBC RNA particle #035. a,** Seven representative tilt images of a single particle are shown in the first column from the left. The tilt images are aligned to a common center using IPET through iterative refinements. The projections of the raw, intermediate, and final 3D reconstructions at the corresponding angles are displayed in the next four columns. **b,** Two orthogonal views of the final 3D map; **c,** Overlap of the map with the fitted model; **d,** FSC analyses of the final map resolution using two methods, "map-map FSC", in which each map is reconstructed from one half of the images, based on the even vs. odd indices of frames or tilt angles. respectively, **e,** FSC analysis of map-model, where the model map is generated by the fitted model after being low-pass filtered to 8 Å. The resolutions are assessed based on the frequencies of the FSC curve falls at 0.5. Scale bars represent 10 nm in a, and 5 nm in b and c.

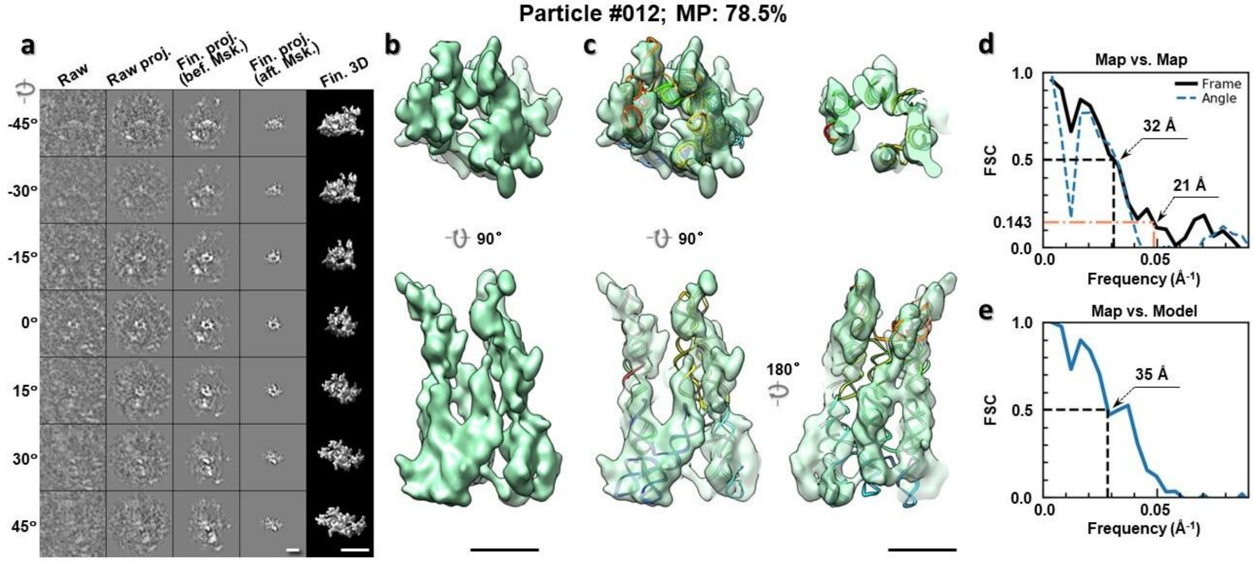

**Supplementary Fig. 110: IPET 3D reconstruction of an individual 6HBC RNA particle #012. a,** Seven representative tilt images of a single particle are shown in the first column from the left. The tilt images are aligned to a common center using IPET through iterative refinements. The projections of the raw, intermediate, and final 3D reconstructions at the corresponding angles are displayed in the next four

columns. **b,** Two orthogonal views of the final 3D map; **c,** Overlap of the map with the fitted model; **d,** FSC analyses of the final map resolution using two methods, "map-map FSC", in which each map is reconstructed from one half of the images, based on the even vs. odd indices of frames or tilt angles. respectively, **e,** FSC analysis of map-model, where the model map is generated by the fitted model after being low-pass filtered to 8 Å. The resolutions are assessed based on the frequencies of the FSC curve falls at 0.5. Scale bars represent 10 nm in a, and 5 nm in b and c.

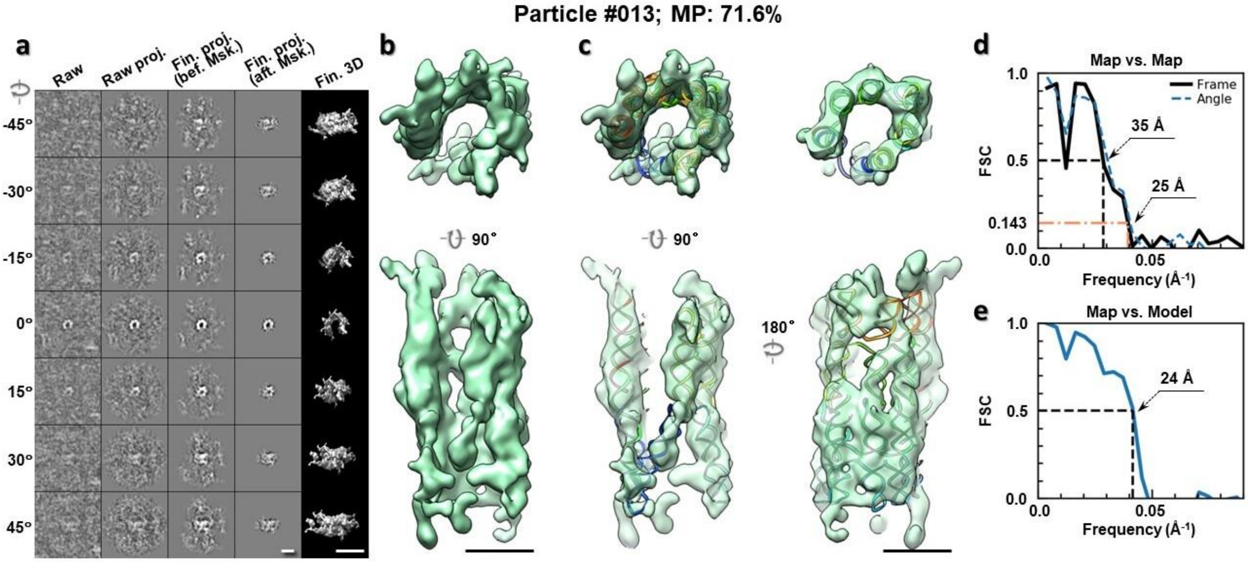

**Supplementary Fig. 111: IPET 3D reconstruction of an individual 6HBC RNA particle #013. a,** Seven representative tilt images of a single particle are shown in the first column from the left. The tilt images are aligned to a common center using IPET through iterative refinements. The projections of the raw, intermediate, and final 3D reconstructions at the corresponding angles are displayed in the next four columns. **b,** Two orthogonal views of the final 3D map; **c,** Overlap of the map with the fitted model; **d,** FSC analyses of the final map resolution using two methods, "map-map FSC", in which each map is reconstructed from one half of the images, based on the even vs. odd indices of frames or tilt angles. respectively, **e,** FSC analysis of map-model, where the model map is generated by the fitted model after being low-pass filtered to 8 Å. The resolutions are assessed based on the frequencies of the FSC curve falls at 0.5. Scale bars represent 10 nm in a, and 5 nm in b and c.

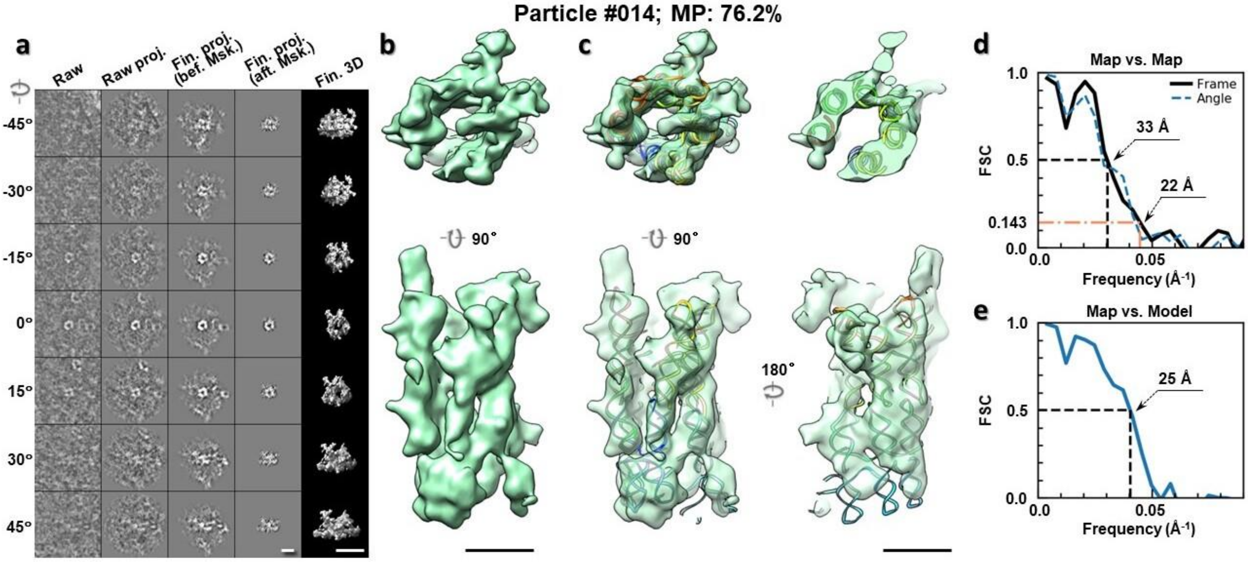

**Supplementary Fig. 112: IPET 3D reconstruction of an individual 6HBC RNA particle #014. a,** Seven representative tilt images of a single particle are shown in the first column from the left. The tilt images are aligned to a common center using IPET through iterative refinements. The projections of the raw, intermediate, and final 3D reconstructions at the corresponding angles are displayed in the next four columns. **b,** Two orthogonal views of the final 3D map; **c,** Overlap of the map with the fitted model; **d,** FSC analyses of the final map resolution using two methods, "map-map FSC", in which each map is reconstructed from one half of the images, based on the even vs. odd indices of frames or tilt angles. respectively, **e,** FSC analysis of map-model, where the model map is generated by the fitted model after being low-pass filtered to 8 Å. The resolutions are assessed based on the frequencies of the FSC curve falls at 0.5. Scale bars represent 10 nm in a, and 5 nm in b and c.

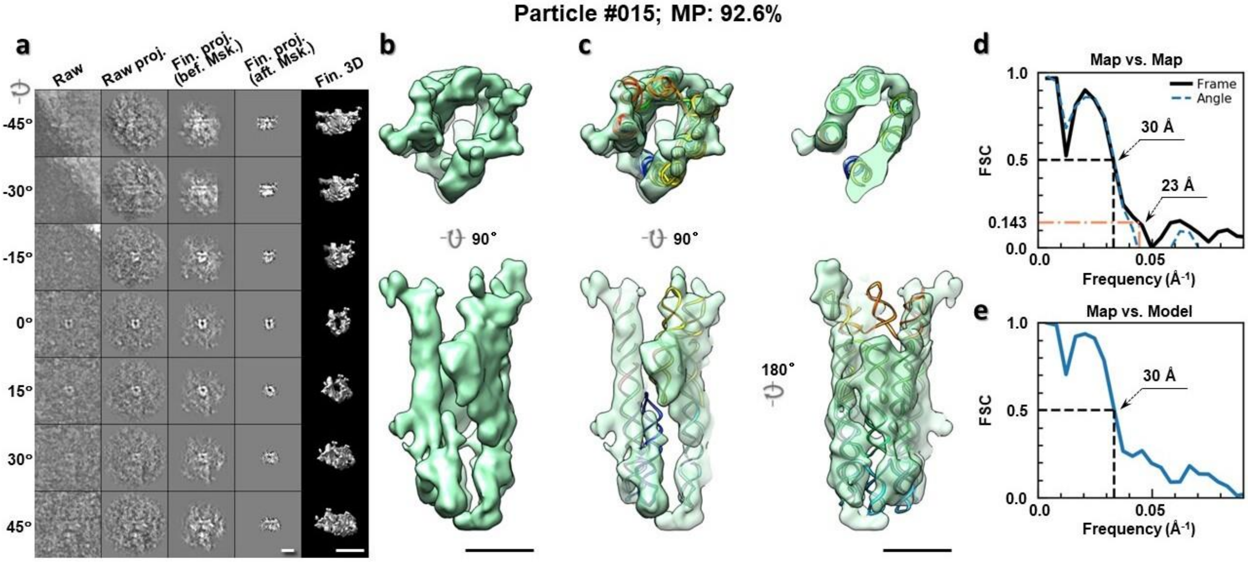

**Supplementary Fig. 113: IPET 3D reconstruction of an individual 6HBC RNA particle #015. a,** Seven representative tilt images of a single particle are shown in the first column from the left. The tilt images are aligned to a common center using IPET through iterative refinements. The projections of the raw, intermediate, and final 3D reconstructions at the corresponding angles are displayed in the next four

columns. **b,** Two orthogonal views of the final 3D map; **c,** Overlap of the map with the fitted model; **d,** FSC analyses of the final map resolution using two methods, "map-map FSC", in which each map is reconstructed from one half of the images, based on the even vs. odd indices of frames or tilt angles. respectively, **e,** FSC analysis of map-model, where the model map is generated by the fitted model after being low-pass filtered to 8 Å. The resolutions are assessed based on the frequencies of the FSC curve falls at 0.5. Scale bars represent 10 nm in a, and 5 nm in b and c.

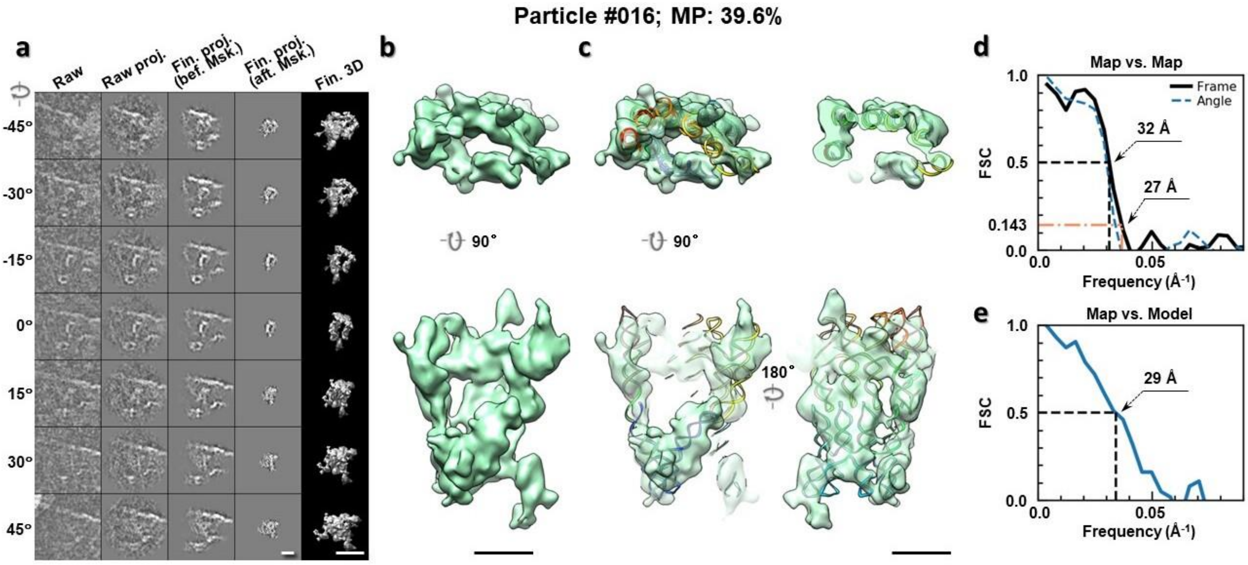

**Supplementary Fig. 114: IPET 3D reconstruction of an individual 6HBC RNA particle #016. a,** Seven representative tilt images of a single particle are shown in the first column from the left. The tilt images are aligned to a common center using IPET through iterative refinements. The projections of the raw, intermediate, and final 3D reconstructions at the corresponding angles are displayed in the next four columns. **b,** Two orthogonal views of the final 3D map; **c,** Overlap of the map with the fitted model; **d,** FSC analyses of the final map resolution using two methods, "map-map FSC", in which each map is reconstructed from one half of the images, based on the even vs. odd indices of frames or tilt angles. respectively, **e,** FSC analysis of map-model, where the model map is generated by the fitted model after being low-pass filtered to 8 Å. The resolutions are assessed based on the frequencies of the FSC curve falls at 0.5. Scale bars represent 10 nm in a, and 5 nm in b and c.

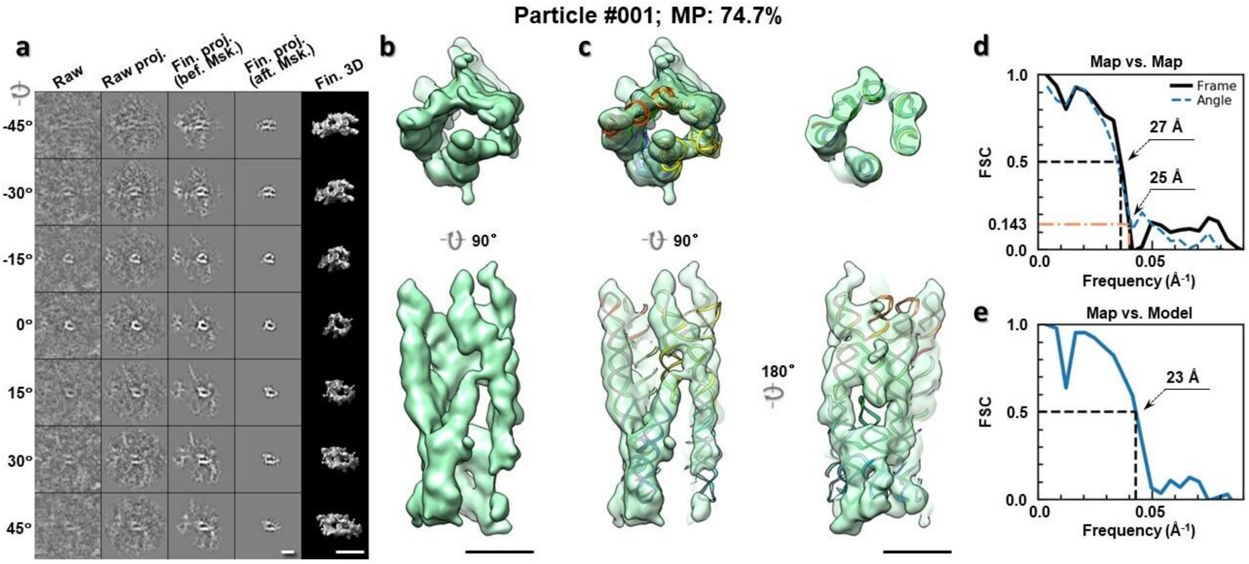

**Supplementary Fig. 115: IPET 3D reconstruction of an individual 6HBC RNA particle #001. a,** Seven representative tilt images of a single particle are shown in the first column from the left. The tilt images are aligned to a common center using IPET through iterative refinements. The projections of the raw, intermediate, and final 3D reconstructions at the corresponding angles are displayed in the next four columns. **b,** Two orthogonal views of the final 3D map; **c,** Overlap of the map with the fitted model; **d,** FSC analyses of the final map resolution using two methods, "map-map FSC", in which each map is reconstructed from one half of the images, based on the even vs. odd indices of frames or tilt angles. respectively, **e,** FSC analysis of map-model, where the model map is generated by the fitted model after being low-pass filtered to 8 Å. The resolutions are assessed based on the frequencies of the FSC curve falls at 0.5. Scale bars represent 10 nm in a, and 5 nm in b and c.

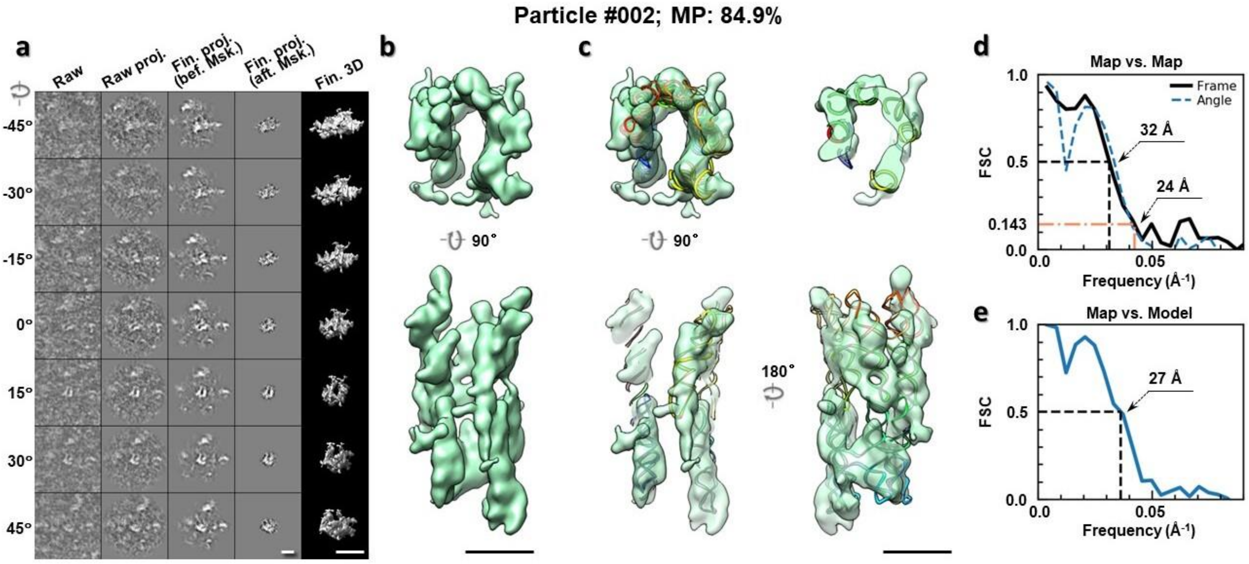

**Supplementary Fig. 116: IPET 3D reconstruction of an individual 6HBC RNA particle #002. a,** Seven representative tilt images of a single particle are shown in the first column from the left. The tilt images are aligned to a common center using IPET through iterative refinements. The projections of the raw, intermediate, and final 3D reconstructions at the corresponding angles are displayed in the next four

columns. **b,** Two orthogonal views of the final 3D map; **c,** Overlap of the map with the fitted model; **d,** FSC analyses of the final map resolution using two methods, "map-map FSC", in which each map is reconstructed from one half of the images, based on the even vs. odd indices of frames or tilt angles. respectively, **e,** FSC analysis of map-model, where the model map is generated by the fitted model after being low-pass filtered to 8 Å. The resolutions are assessed based on the frequencies of the FSC curve falls at 0.5. Scale bars represent 10 nm in a, and 5 nm in b and c.

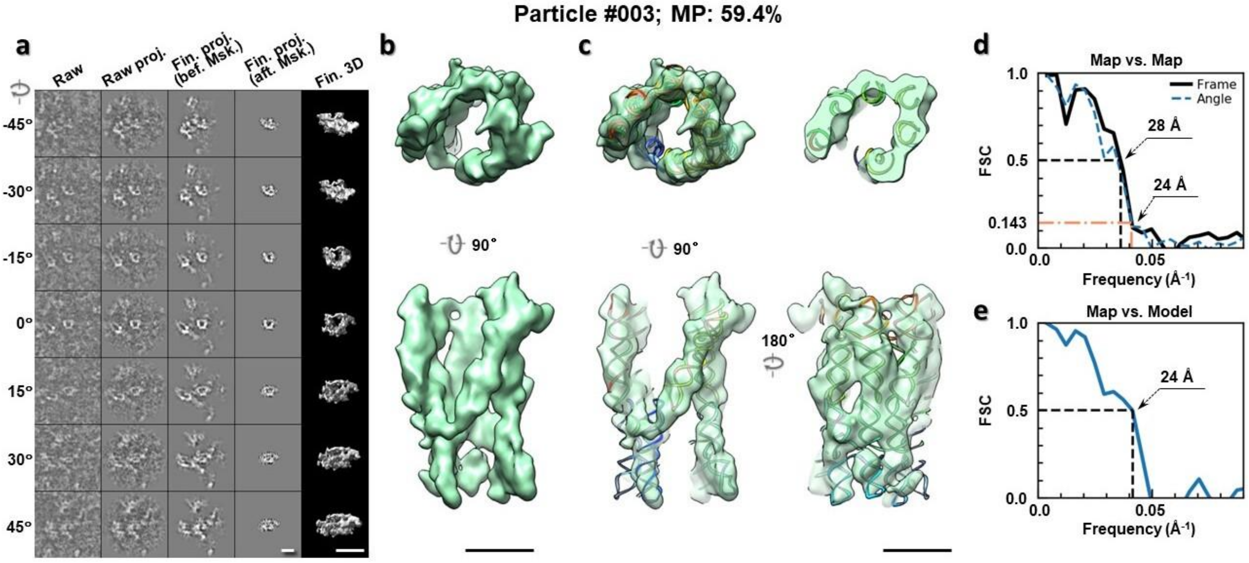

**Supplementary Fig. 117: IPET 3D reconstruction of an individual 6HBC RNA particle #003. a,** Seven representative tilt images of a single particle are shown in the first column from the left. The tilt images are aligned to a common center using IPET through iterative refinements. The projections of the raw, intermediate, and final 3D reconstructions at the corresponding angles are displayed in the next four columns. **b,** Two orthogonal views of the final 3D map; **c,** Overlap of the map with the fitted model; **d,** FSC analyses of the final map resolution using two methods, "map-map FSC", in which each map is reconstructed from one half of the images, based on the even vs. odd indices of frames or tilt angles. respectively, **e,** FSC analysis of map-model, where the model map is generated by the fitted model after being low-pass filtered to 8 Å. The resolutions are assessed based on the frequencies of the FSC curve falls at 0.5. Scale bars represent 10 nm in a, and 5 nm in b and c.

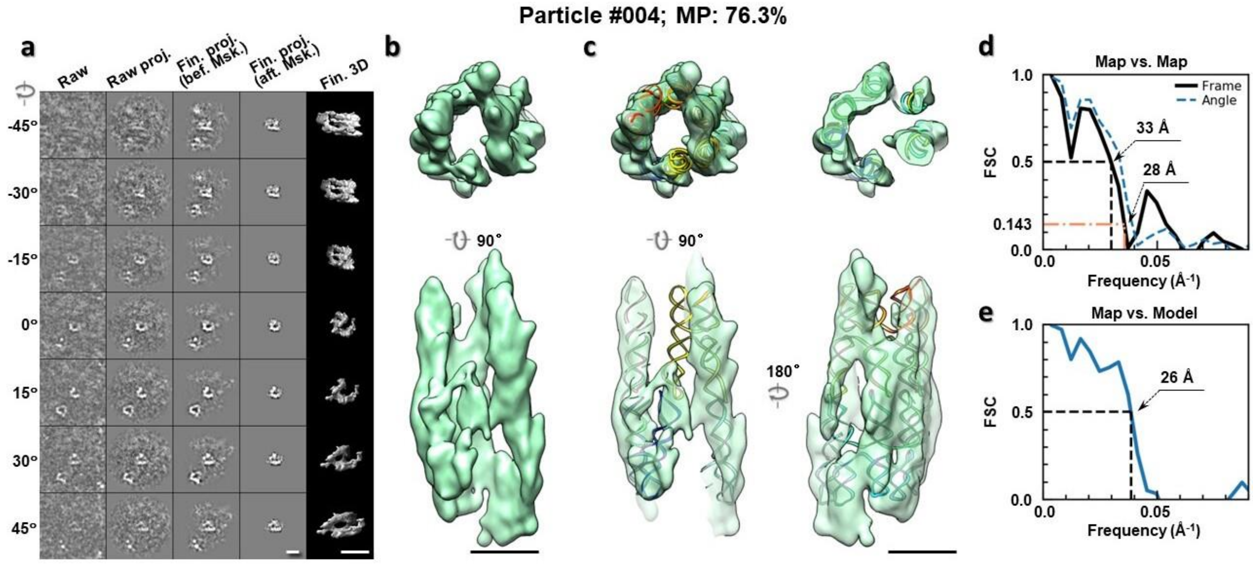

**Supplementary Fig. 118: IPET 3D reconstruction of an individual 6HBC RNA particle #004. a,** Seven representative tilt images of a single particle are shown in the first column from the left. The tilt images are aligned to a common center using IPET through iterative refinements. The projections of the raw, intermediate, and final 3D reconstructions at the corresponding angles are displayed in the next four columns. **b,** Two orthogonal views of the final 3D map; **c,** Overlap of the map with the fitted model; **d,** FSC analyses of the final map resolution using two methods, "map-map FSC", in which each map is reconstructed from one half of the images, based on the even vs. odd indices of frames or tilt angles. respectively, **e,** FSC analysis of map-model, where the model map is generated by the fitted model after being low-pass filtered to 8 Å. The resolutions are assessed based on the frequencies of the FSC curve falls at 0.5. Scale bars represent 10 nm in a, and 5 nm in b and c.

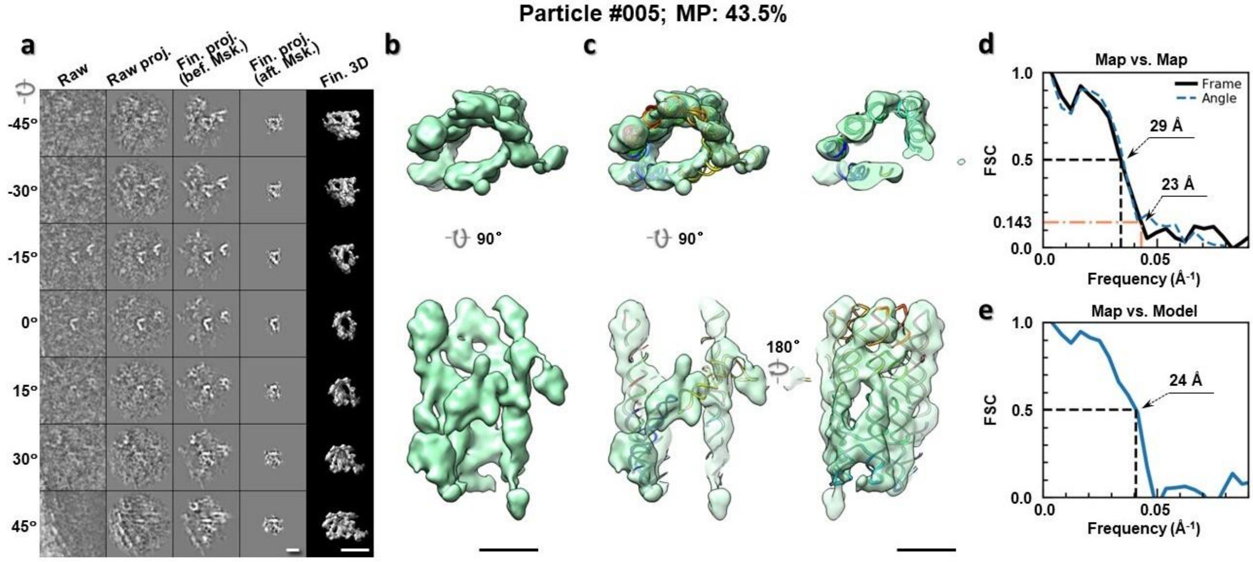

**Supplementary Fig. 119: IPET 3D reconstruction of an individual 6HBC RNA particle #005. a,** Seven representative tilt images of a single particle are shown in the first column from the left. The tilt images are aligned to a common center using IPET through iterative refinements. The projections of the raw, intermediate, and final 3D reconstructions at the corresponding angles are displayed in the next four

columns. **b,** Two orthogonal views of the final 3D map; **c,** Overlap of the map with the fitted model; **d,** FSC analyses of the final map resolution using two methods, "map-map FSC", in which each map is reconstructed from one half of the images, based on the even vs. odd indices of frames or tilt angles. respectively, **e,** FSC analysis of map-model, where the model map is generated by the fitted model after being low-pass filtered to 8 Å. The resolutions are assessed based on the frequencies of the FSC curve falls at 0.5. Scale bars represent 10 nm in a, and 5 nm in b and c.

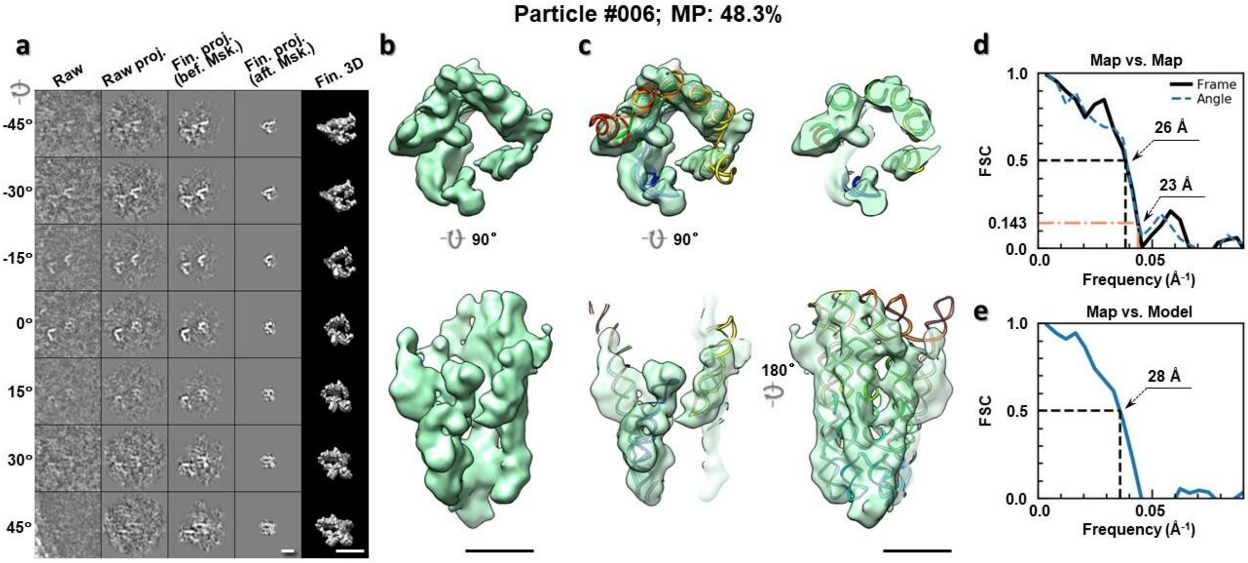

**Supplementary Fig. 120: IPET 3D reconstruction of an individual 6HBC RNA particle #006. a,** Seven representative tilt images of a single particle are shown in the first column from the left. The tilt images are aligned to a common center using IPET through iterative refinements. The projections of the raw, intermediate, and final 3D reconstructions at the corresponding angles are displayed in the next four columns. **b,** Two orthogonal views of the final 3D map; **c,** Overlap of the map with the fitted model; **d,** FSC analyses of the final map resolution using two methods, "map-map FSC", in which each map is reconstructed from one half of the images, based on the even vs. odd indices of frames or tilt angles. respectively, **e,** FSC analysis of map-model, where the model map is generated by the fitted model after being low-pass filtered to 8 Å. The resolutions are assessed based on the frequencies of the FSC curve falls at 0.5. Scale bars represent 10 nm in a, and 5 nm in b and c.

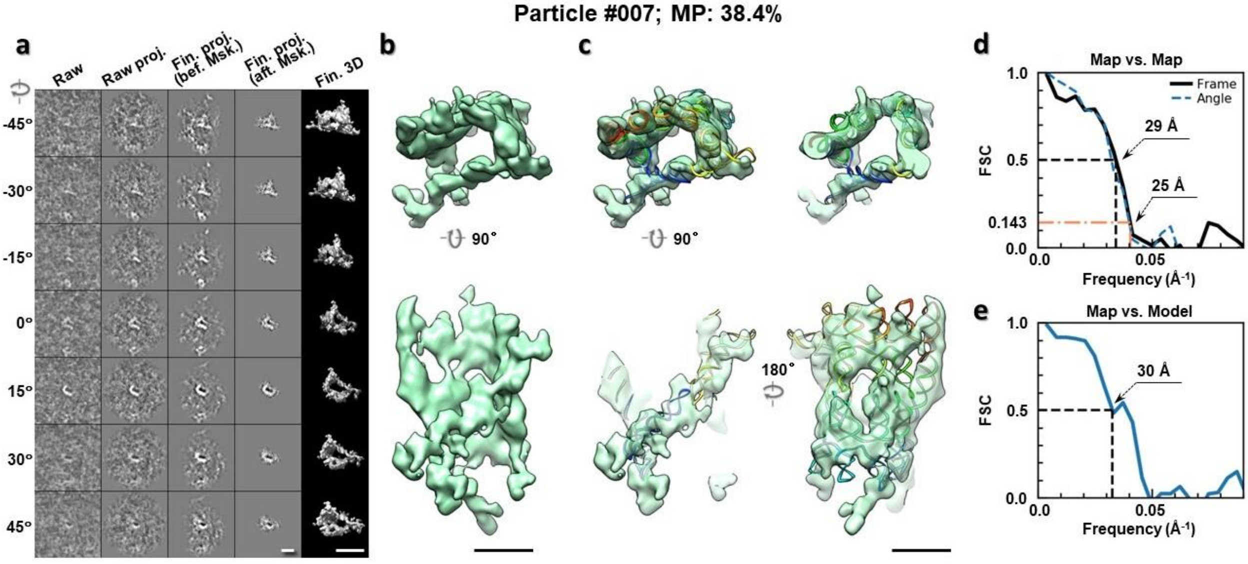

**Supplementary Fig. 121: IPET 3D reconstruction of an individual 6HBC RNA particle #007. a,** Seven representative tilt images of a single particle are shown in the first column from the left. The tilt images are aligned to a common center using IPET through iterative refinements. The projections of the raw, intermediate, and final 3D reconstructions at the corresponding angles are displayed in the next four columns. **b,** Two orthogonal views of the final 3D map; **c,** Overlap of the map with the fitted model; **d,** FSC analyses of the final map resolution using two methods, "map-map FSC", in which each map is reconstructed from one half of the images, based on the even vs. odd indices of frames or tilt angles. respectively, **e,** FSC analysis of map-model, where the model map is generated by the fitted model after being low-pass filtered to 8 Å. The resolutions are assessed based on the frequencies of the FSC curve falls at 0.5. Scale bars represent 10 nm in a, and 5 nm in b and c.

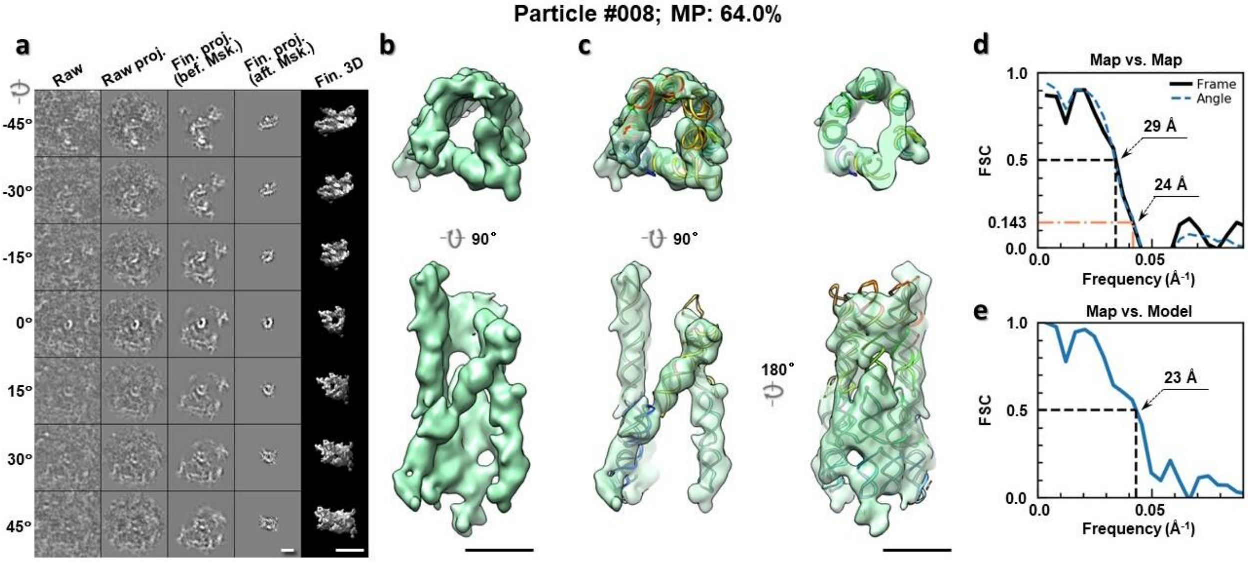

**Supplementary Fig. 122: IPET 3D reconstruction of an individual 6HBC RNA particle #008. a,** Seven representative tilt images of a single particle are shown in the first column from the left. The tilt images are aligned to a common center using IPET through iterative refinements. The projections of the raw, intermediate, and final 3D reconstructions at the corresponding angles are displayed in the next four

columns. **b,** Two orthogonal views of the final 3D map; **c,** Overlap of the map with the fitted model; **d,** FSC analyses of the final map resolution using two methods, "map-map FSC", in which each map is reconstructed from one half of the images, based on the even vs. odd indices of frames or tilt angles. respectively, **e,** FSC analysis of map-model, where the model map is generated by the fitted model after being low-pass filtered to 8 Å. The resolutions are assessed based on the frequencies of the FSC curve falls at 0.5. Scale bars represent 10 nm in a, and 5 nm in b and c.

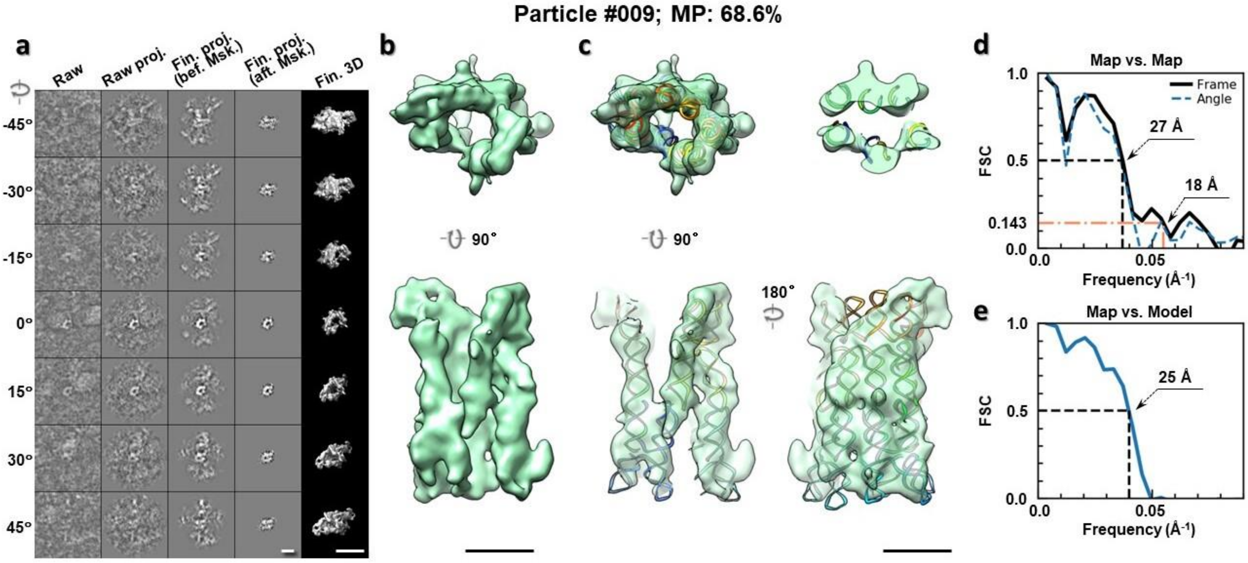

**Supplementary Fig. 123: IPET 3D reconstruction of an individual 6HBC RNA particle #009.** **a,** Seven representative tilt images of a single particle are shown in the first column from the left. The tilt images are aligned to a common center using IPET through iterative refinements. The projections of the raw, intermediate, and final 3D reconstructions at the corresponding angles are displayed in the next four columns. **b,** Two orthogonal views of the final 3D map; **c,** Overlap of the map with the fitted model; **d,** FSC analyses of the final map resolution using two methods, "map-map FSC", in which each map is reconstructed from one half of the images, based on the even vs. odd indices of frames or tilt angles. respectively, **e,** FSC analysis of map-model, where the model map is generated by the fitted model after being low-pass filtered to 8 Å. The resolutions are assessed based on the frequencies of the FSC curve falls at 0.5. Scale bars represent 10 nm in a, and 5 nm in b and c.

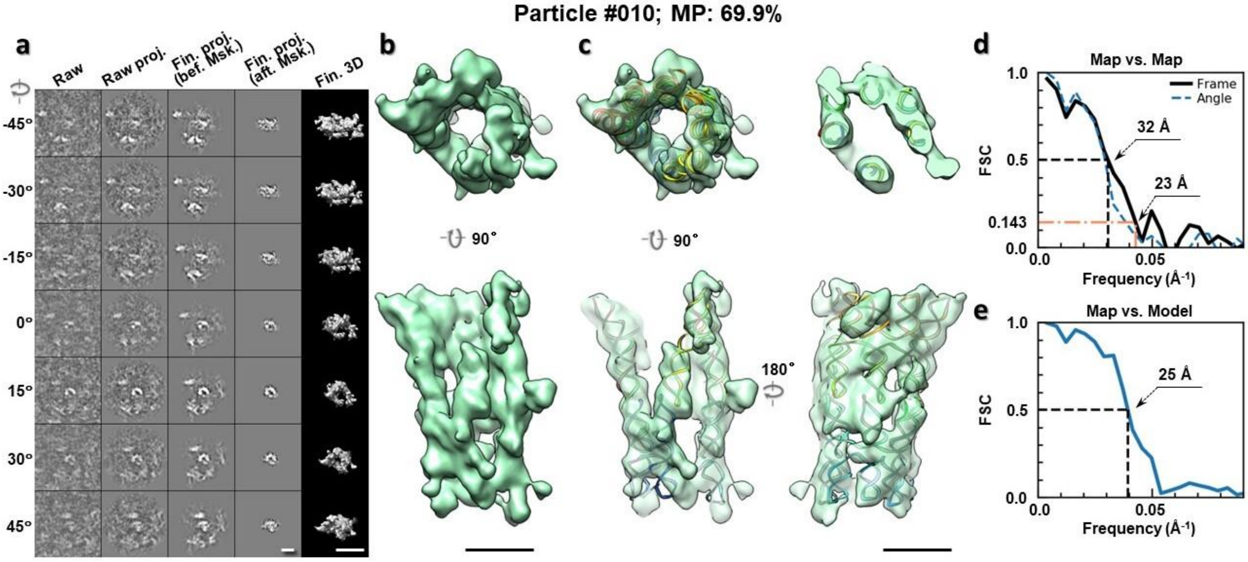

**Supplementary Fig. 124: IPET 3D reconstruction of an individual 6HBC RNA particle #010. a,** Seven representative tilt images of a single particle are shown in the first column from the left. The tilt images are aligned to a common center using IPET through iterative refinements. The projections of the raw, intermediate, and final 3D reconstructions at the corresponding angles are displayed in the next four columns. **b,** Two orthogonal views of the final 3D map; **c,** Overlap of the map with the fitted model; **d,** FSC analyses of the final map resolution using two methods, "map-map FSC", in which each map is reconstructed from one half of the images, based on the even vs. odd indices of frames or tilt angles. respectively, **e,** FSC analysis of map-model, where the model map is generated by the fitted model after being low-pass filtered to 8 Å. The resolutions are assessed based on the frequencies of the FSC curve falls at 0.5. Scale bars represent 10 nm in a, and 5 nm in b and c.

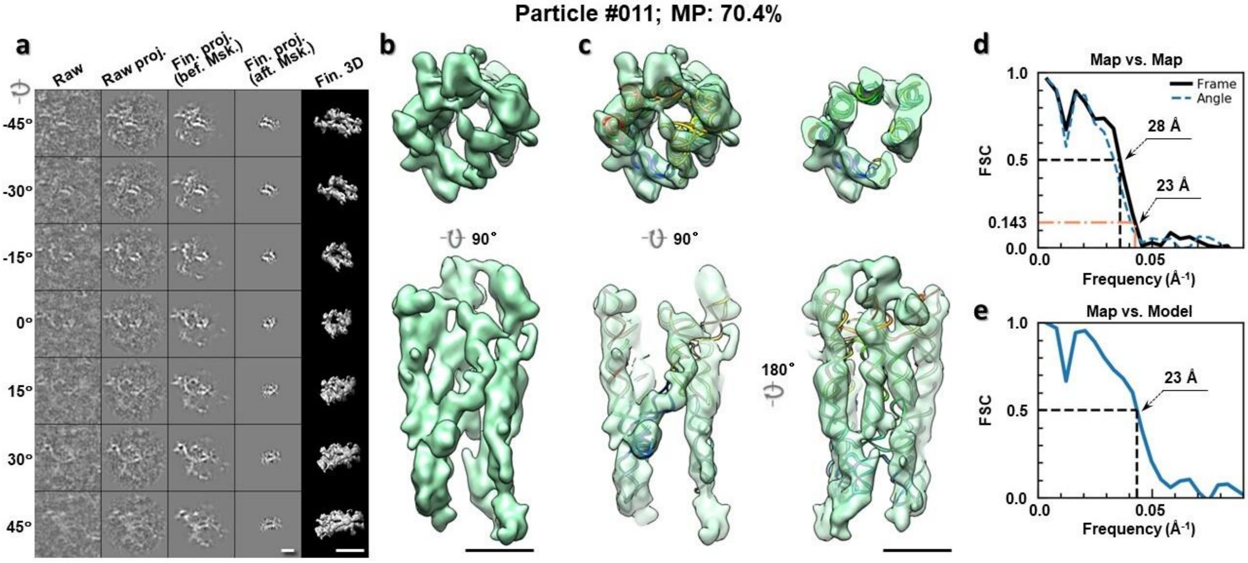

**Supplementary Fig. 125: IPET 3D reconstruction of an individual 6HBC RNA particle #011. a,** Seven representative tilt images of a single particle are shown in the first column from the left. The tilt images are aligned to a common center using IPET through iterative refinements. The projections of the raw, intermediate, and final 3D reconstructions at the corresponding angles are displayed in the next four

columns. **b,** Two orthogonal views of the final 3D map; **c,** Overlap of the map with the fitted model; **d,** FSC analyses of the final map resolution using two methods, "map-map FSC", in which each map is reconstructed from one half of the images, based on the even vs. odd indices of frames or tilt angles. respectively, **e,** FSC analysis of map-model, where the model map is generated by the fitted model after being low-pass filtered to 8 Å. The resolutions are assessed based on the frequencies of the FSC curve falls at 0.5. Scale bars represent 10 nm in a, and 5 nm in b and c.

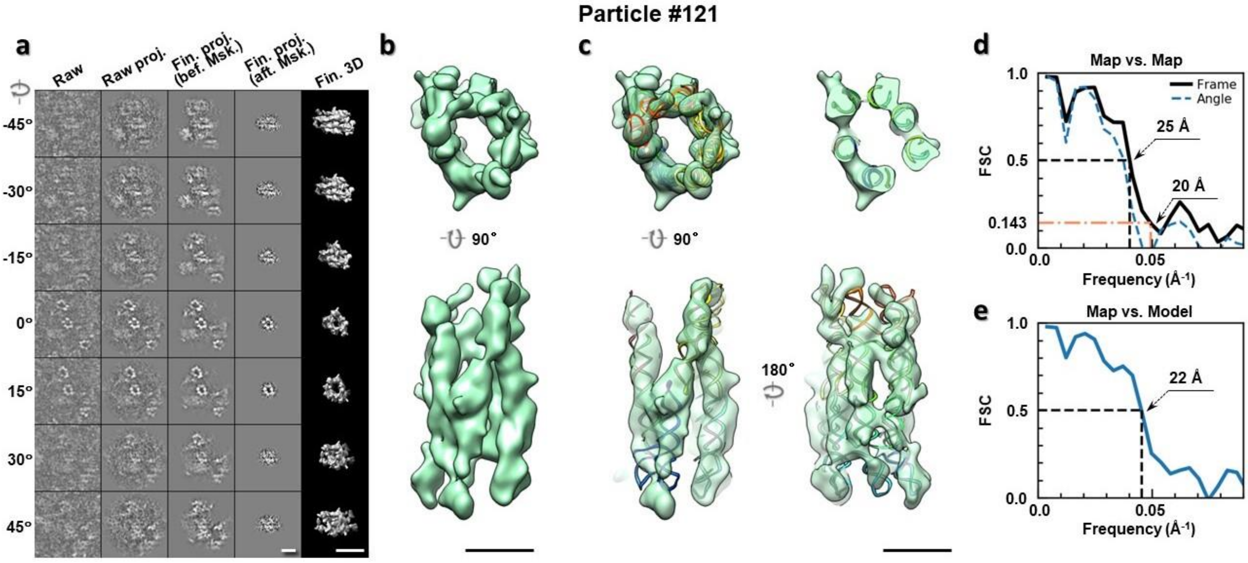

**Supplementary Fig. 126: IPET 3D reconstruction of an individual 6HBC RNA particle #121.** **a,** Seven representative tilt images of a single particle are shown in the first column from the left. The tilt images are aligned to a common center using IPET through iterative refinements. The projections of the raw, intermediate, and final 3D reconstructions at the corresponding angles are displayed in the next four columns. **b,** Two orthogonal views of the final 3D map; **c,** Overlap of the map with the fitted model; **d,** FSC analyses of the final map resolution using two methods, "map-map FSC", in which each map is reconstructed from one half of the images, based on the even vs. odd indices of frames or tilt angles. respectively, **e,** FSC analysis of map-model, where the model map is generated by the fitted model after being low-pass filtered to 8 Å. The resolutions are assessed based on the frequencies of the FSC curve falls at 0.5. Scale bars represent 10 nm in a, and 5 nm in b and c.

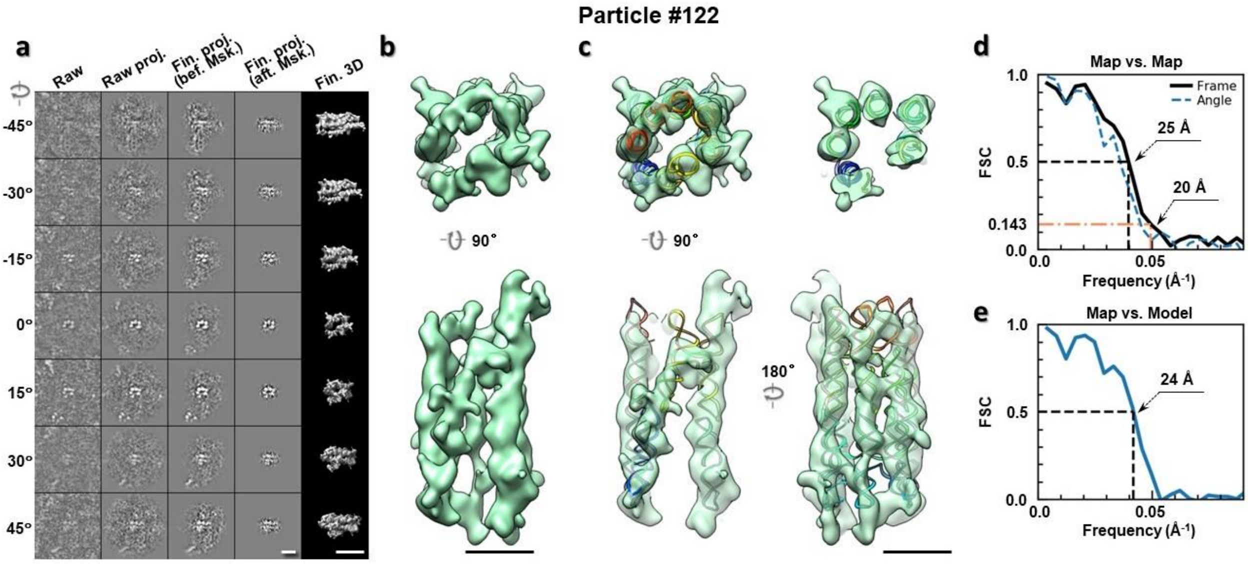

**Supplementary Fig. 127: IPET 3D reconstruction of an individual 6HBC RNA particle #122. a,** Seven representative tilt images of a single particle are shown in the first column from the left. The tilt images are aligned to a common center using IPET through iterative refinements. The projections of the raw, intermediate, and final 3D reconstructions at the corresponding angles are displayed in the next four columns. **b,** Two orthogonal views of the final 3D map; **c,** Overlap of the map with the fitted model; **d,** FSC analyses of the final map resolution using two methods, "map-map FSC", in which each map is reconstructed from one half of the images, based on the even vs. odd indices of frames or tilt angles. respectively, **e,** FSC analysis of map-model, where the model map is generated by the fitted model after being low-pass filtered to 8 Å. The resolutions are assessed based on the frequencies of the FSC curve falls at 0.5. Scale bars represent 10 nm in a, and 5 nm in b and c.

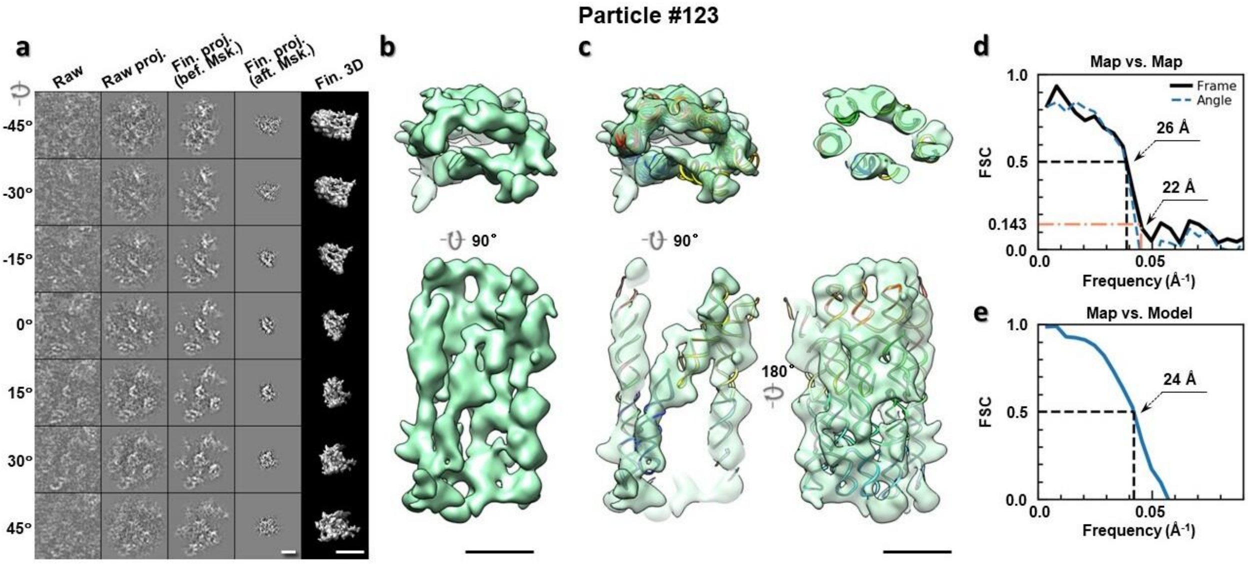

**Supplementary Fig. 128: IPET 3D reconstruction of an individual 6HBC RNA particle #123. a,** Seven representative tilt images of a single particle are shown in the first column from the left. The tilt images are aligned to a common center using IPET through iterative refinements. The projections of the raw, intermediate, and final 3D reconstructions at the corresponding angles are displayed in the next four

columns. **b,** Two orthogonal views of the final 3D map; **c,** Overlap of the map with the fitted model; **d,** FSC analyses of the final map resolution using two methods, "map-map FSC", in which each map is reconstructed from one half of the images, based on the even vs. odd indices of frames or tilt angles. respectively, **e,** FSC analysis of map-model, where the model map is generated by the fitted model after being low-pass filtered to 8 Å. The resolutions are assessed based on the frequencies of the FSC curve falls at 0.5. Scale bars represent 10 nm in a, and 5 nm in b and c.

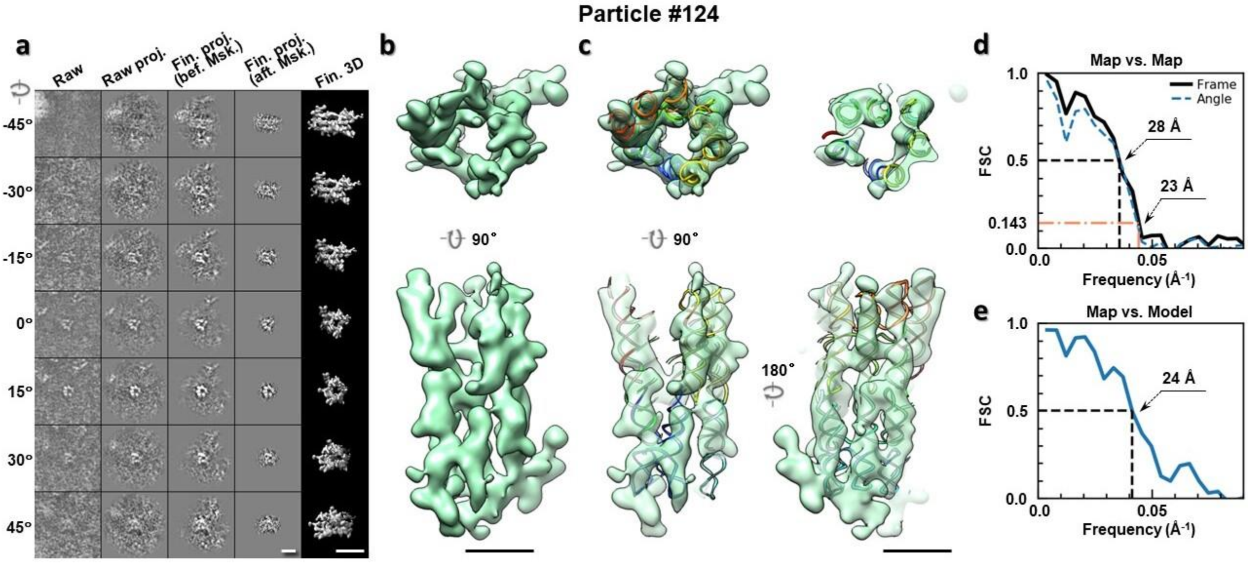

**Supplementary Fig. 129: IPET 3D reconstruction of an individual 6HBC RNA particle #124. a,** Seven representative tilt images of a single particle are shown in the first column from the left. The tilt images are aligned to a common center using IPET through iterative refinements. The projections of the raw, intermediate, and final 3D reconstructions at the corresponding angles are displayed in the next four columns. **b,** Two orthogonal views of the final 3D map; **c,** Overlap of the map with the fitted model; **d,** FSC analyses of the final map resolution using two methods, "map-map FSC", in which each map is reconstructed from one half of the images, based on the even vs. odd indices of frames or tilt angles. respectively, **e,** FSC analysis of map-model, where the model map is generated by the fitted model after being low-pass filtered to 8 Å. The resolutions are assessed based on the frequencies of the FSC curve falls at 0.5. Scale bars represent 10 nm in a, and 5 nm in b and c.

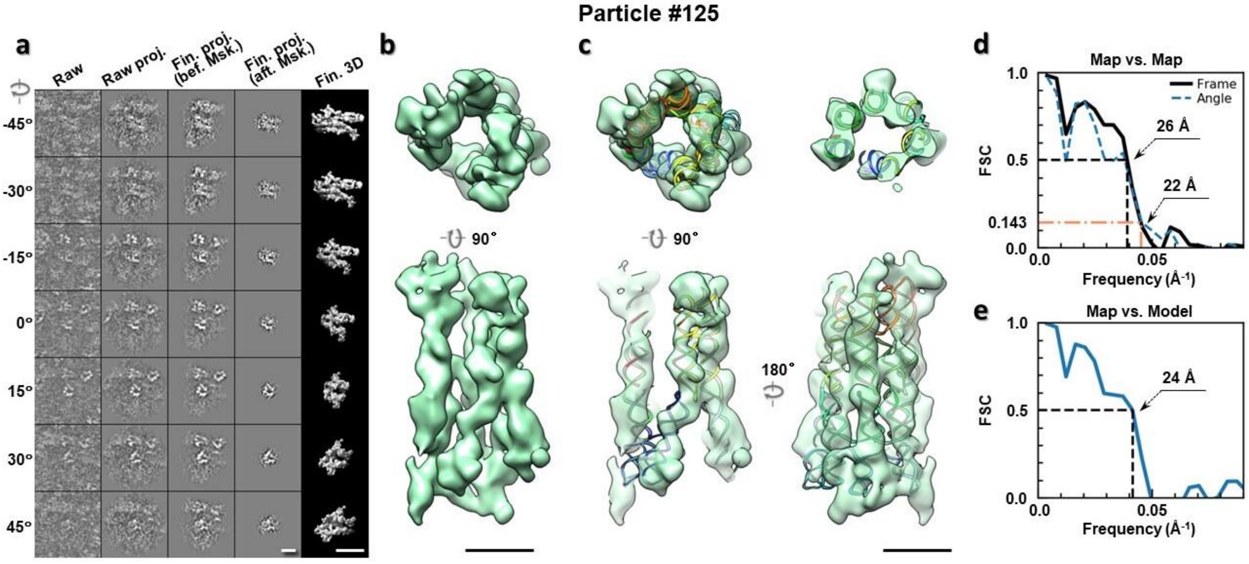

**Supplementary Fig. 130: IPET 3D reconstruction of an individual 6HBC RNA particle #125. a,** Seven representative tilt images of a single particle are shown in the first column from the left. The tilt images are aligned to a common center using IPET through iterative refinements. The projections of the raw, intermediate, and final 3D reconstructions at the corresponding angles are displayed in the next four columns. **b,** Two orthogonal views of the final 3D map; **c,** Overlap of the map with the fitted model; **d,** FSC analyses of the final map resolution using two methods, "map-map FSC", in which each map is reconstructed from one half of the images, based on the even vs. odd indices of frames or tilt angles. respectively, **e,** FSC analysis of map-model, where the model map is generated by the fitted model after being low-pass filtered to 8 Å. The resolutions are assessed based on the frequencies of the FSC curve falls at 0.5. Scale bars represent 10 nm in a, and 5 nm in b and c.

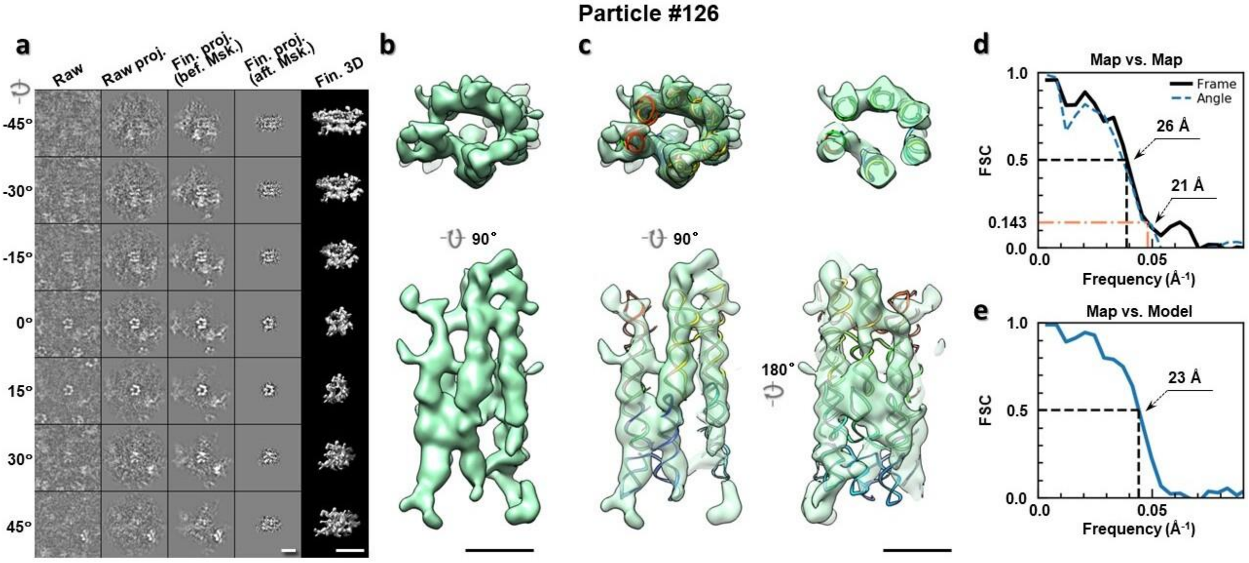

**Supplementary Fig. 131: IPET 3D reconstruction of an individual 6HBC RNA particle #126. a,** Seven representative tilt images of a single particle are shown in the first column from the left. The tilt images are aligned to a common center using IPET through iterative refinements. The projections of the raw, intermediate, and final 3D reconstructions at the corresponding angles are displayed in the next four

columns. **b,** Two orthogonal views of the final 3D map; **c,** Overlap of the map with the fitted model; **d,** FSC analyses of the final map resolution using two methods, "map-map FSC", in which each map is reconstructed from one half of the images, based on the even vs. odd indices of frames or tilt angles. respectively, **e,** FSC analysis of map-model, where the model map is generated by the fitted model after being low-pass filtered to 8 Å. The resolutions are assessed based on the frequencies of the FSC curve falls at 0.5. Scale bars represent 10 nm in a, and 5 nm in b and c.

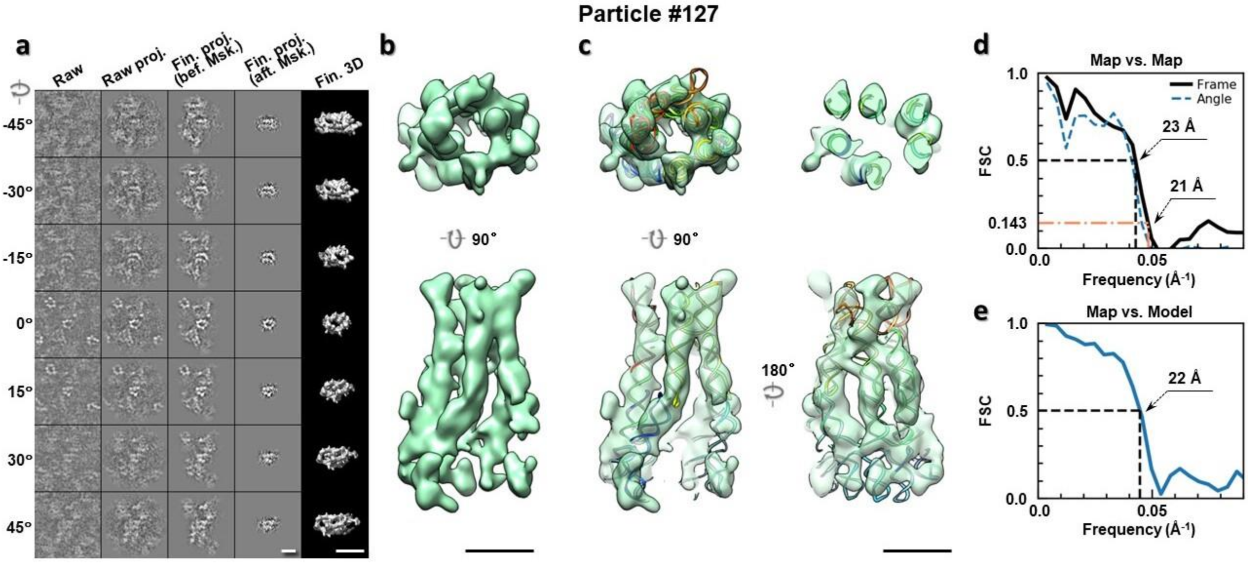

**Supplementary Fig. 132: IPET 3D reconstruction of an individual 6HBC RNA particle #127. a,** Seven representative tilt images of a single particle are shown in the first column from the left. The tilt images are aligned to a common center using IPET through iterative refinements. The projections of the raw, intermediate, and final 3D reconstructions at the corresponding angles are displayed in the next four columns. **b,** Two orthogonal views of the final 3D map; **c,** Overlap of the map with the fitted model; **d,** FSC analyses of the final map resolution using two methods, "map-map FSC", in which each map is reconstructed from one half of the images, based on the even vs. odd indices of frames or tilt angles. respectively, **e,** FSC analysis of map-model, where the model map is generated by the fitted model after being low-pass filtered to 8 Å. The resolutions are assessed based on the frequencies of the FSC curve falls at 0.5. Scale bars represent 10 nm in a, and 5 nm in b and c.

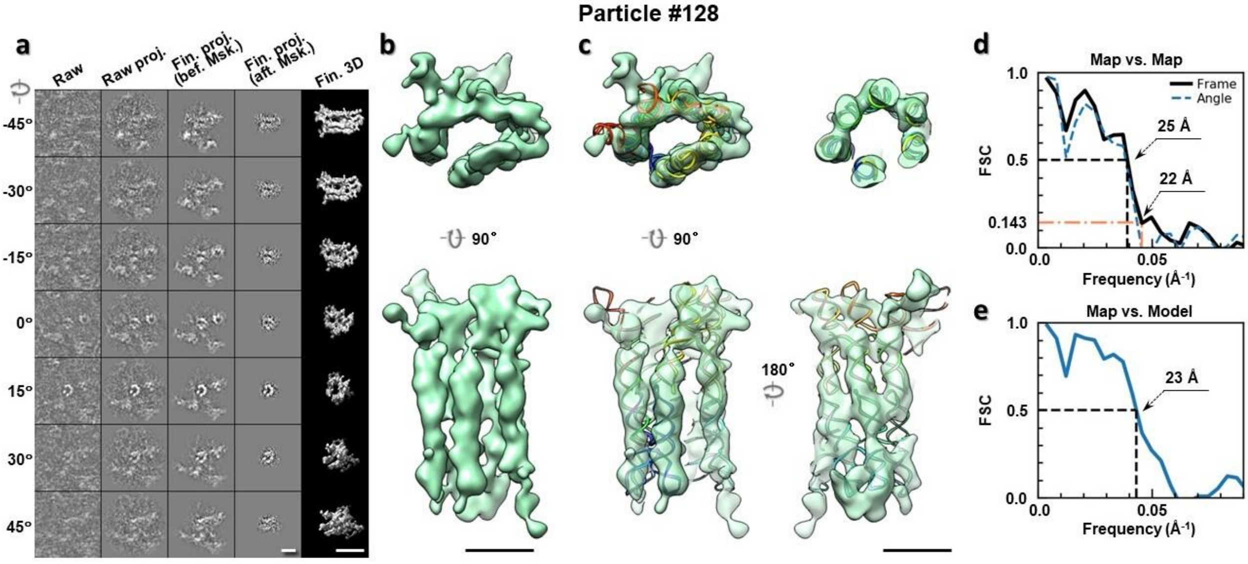

**Supplementary Fig. 133: IPET 3D reconstruction of an individual 6HBC RNA particle #128. a,** Seven representative tilt images of a single particle are shown in the first column from the left. The tilt images are aligned to a common center using IPET through iterative refinements. The projections of the raw, intermediate, and final 3D reconstructions at the corresponding angles are displayed in the next four columns. **b,** Two orthogonal views of the final 3D map; **c,** Overlap of the map with the fitted model; **d,** FSC analyses of the final map resolution using two methods, "map-map FSC", in which each map is reconstructed from one half of the images, based on the even vs. odd indices of frames or tilt angles. respectively, **e,** FSC analysis of map-model, where the model map is generated by the fitted model after being low-pass filtered to 8 Å. The resolutions are assessed based on the frequencies of the FSC curve falls at 0.5. Scale bars represent 10 nm in a, and 5 nm in b and c.

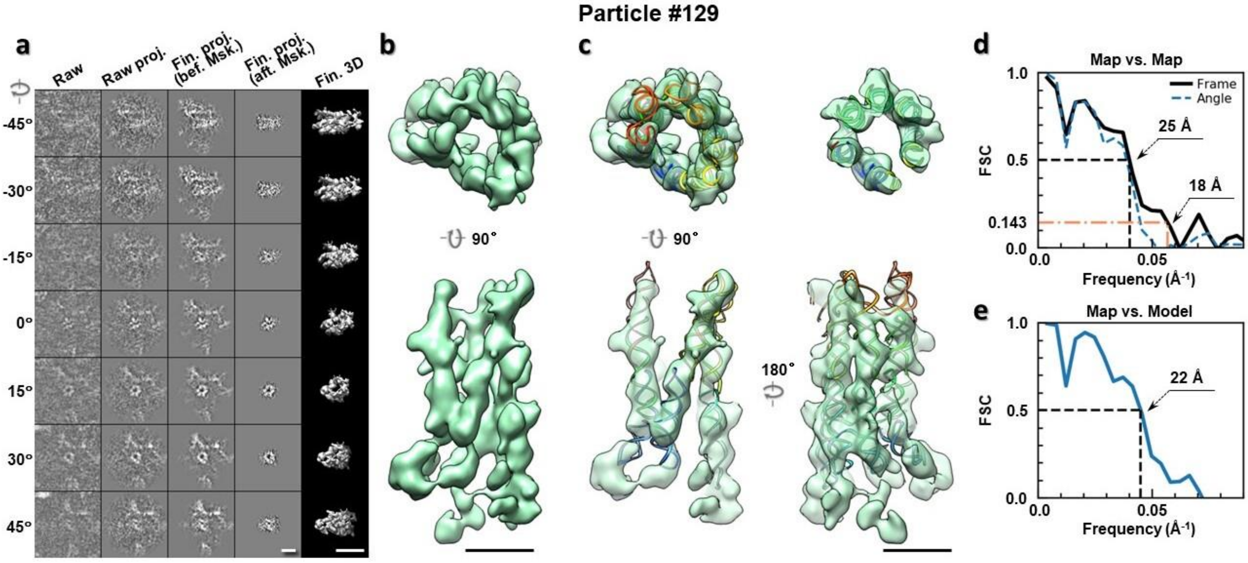

**Supplementary Fig. 134: IPET 3D reconstruction of an individual 6HBC RNA particle #129. a,** Seven representative tilt images of a single particle are shown in the first column from the left. The tilt images are aligned to a common center using IPET through iterative refinements. The projections of the raw, intermediate, and final 3D reconstructions at the corresponding angles are displayed in the next four

columns. **b,** Two orthogonal views of the final 3D map; **c,** Overlap of the map with the fitted model; **d,** FSC analyses of the final map resolution using two methods, "map-map FSC", in which each map is reconstructed from one half of the images, based on the even vs. odd indices of frames or tilt angles. respectively, **e,** FSC analysis of map-model, where the model map is generated by the fitted model after being low-pass filtered to 8 Å. The resolutions are assessed based on the frequencies of the FSC curve falls at 0.5. Scale bars represent 10 nm in a, and 5 nm in b and c.

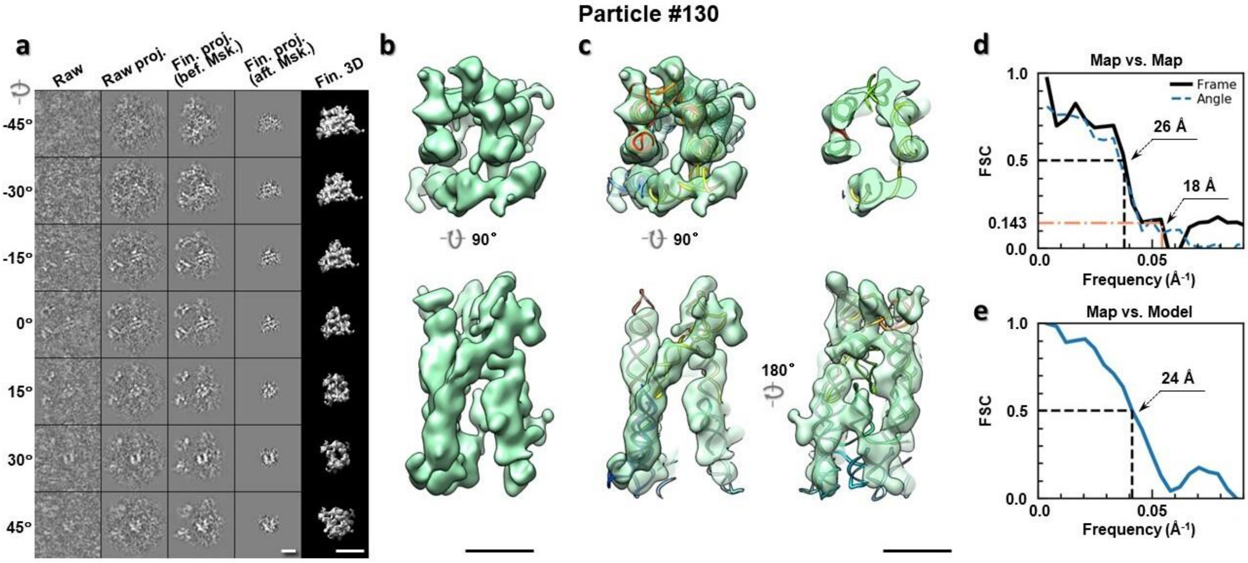

**Supplementary Fig. 135: IPET 3D reconstruction of an individual 6HBC RNA particle #130. a,** Seven representative tilt images of a single particle are shown in the first column from the left. The tilt images are aligned to a common center using IPET through iterative refinements. The projections of the raw, intermediate, and final 3D reconstructions at the corresponding angles are displayed in the next four columns. **b,** Two orthogonal views of the final 3D map; **c,** Overlap of the map with the fitted model; **d,** FSC analyses of the final map resolution using two methods, "map-map FSC", in which each map is reconstructed from one half of the images, based on the even vs. odd indices of frames or tilt angles. respectively, **e,** FSC analysis of map-model, where the model map is generated by the fitted model after being low-pass filtered to 8 Å. The resolutions are assessed based on the frequencies of the FSC curve falls at 0.5. Scale bars represent 10 nm in a, and 5 nm in b and c.

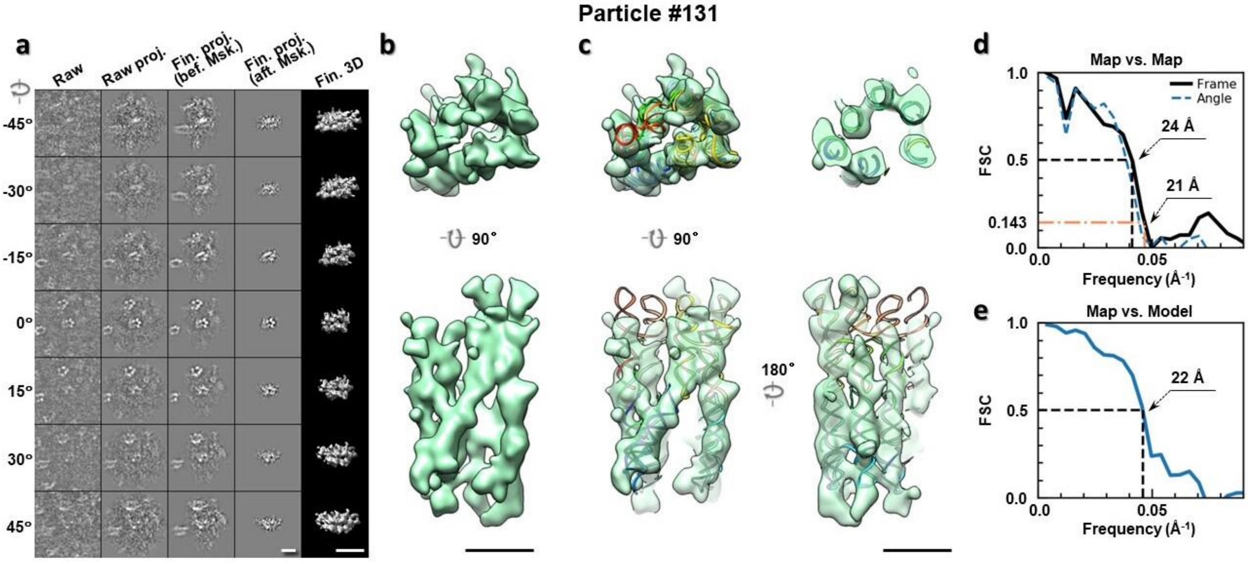

**Supplementary Fig. 136: IPET 3D reconstruction of an individual 6HBC RNA particle #131. a,** Seven representative tilt images of a single particle are shown in the first column from the left. The tilt images are aligned to a common center using IPET through iterative refinements. The projections of the raw, intermediate, and final 3D reconstructions at the corresponding angles are displayed in the next four columns. **b,** Two orthogonal views of the final 3D map; **c,** Overlap of the map with the fitted model; **d,** FSC analyses of the final map resolution using two methods, "map-map FSC", in which each map is reconstructed from one half of the images, based on the even vs. odd indices of frames or tilt angles. respectively, **e,** FSC analysis of map-model, where the model map is generated by the fitted model after being low-pass filtered to 8 Å. The resolutions are assessed based on the frequencies of the FSC curve falls at 0.5. Scale bars represent 10 nm in a, and 5 nm in b and c.

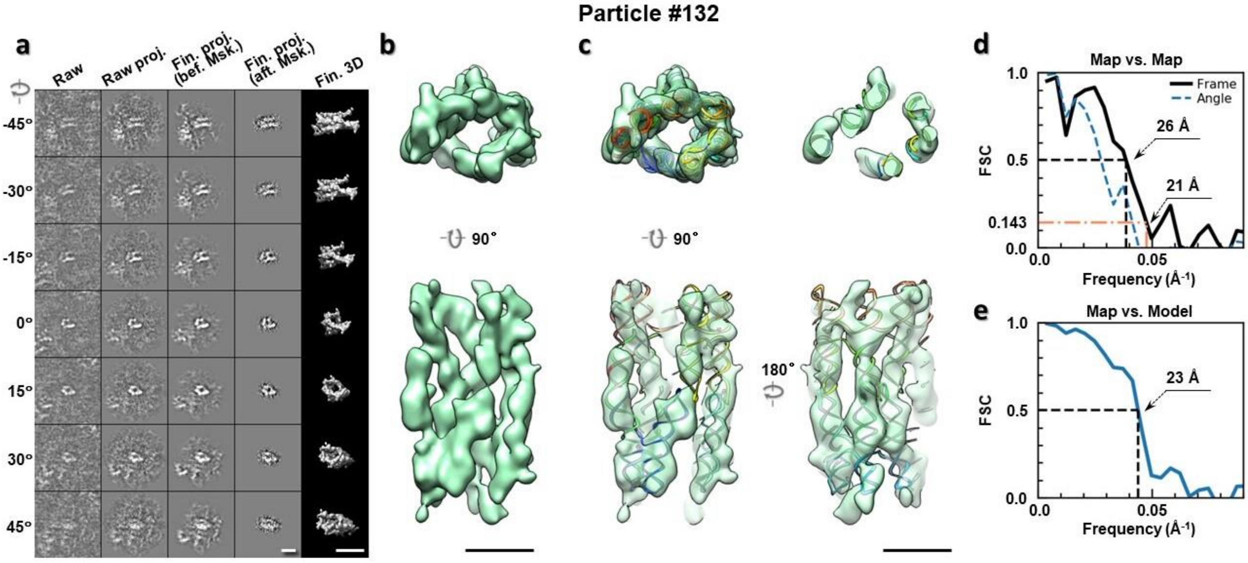

**Supplementary Fig. 137: IPET 3D reconstruction of an individual 6HBC RNA particle #132. a,** Seven representative tilt images of a single particle are shown in the first column from the left. The tilt images are aligned to a common center using IPET through iterative refinements. The projections of the raw, intermediate, and final 3D reconstructions at the corresponding angles are displayed in the next four

columns. **b,** Two orthogonal views of the final 3D map; **c,** Overlap of the map with the fitted model; **d,** FSC analyses of the final map resolution using two methods, "map-map FSC", in which each map is reconstructed from one half of the images, based on the even vs. odd indices of frames or tilt angles. respectively, **e,** FSC analysis of map-model, where the model map is generated by the fitted model after being low-pass filtered to 8 Å. The resolutions are assessed based on the frequencies of the FSC curve falls at 0.5. Scale bars represent 10 nm in a, and 5 nm in b and c.

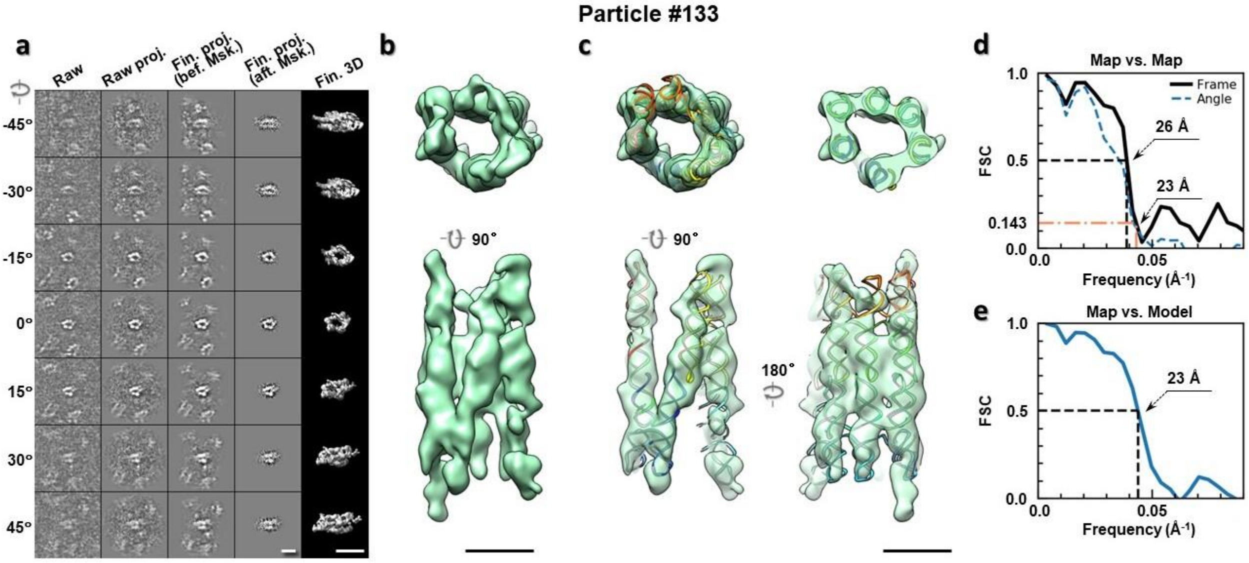

**Supplementary Fig. 138: IPET 3D reconstruction of an individual 6HBC RNA particle #133.** **a,** Seven representative tilt images of a single particle are shown in the first column from the left. The tilt images are aligned to a common center using IPET through iterative refinements. The projections of the raw, intermediate, and final 3D reconstructions at the corresponding angles are displayed in the next four columns. **b,** Two orthogonal views of the final 3D map; **c,** Overlap of the map with the fitted model; **d,** FSC analyses of the final map resolution using two methods, "map-map FSC", in which each map is reconstructed from one half of the images, based on the even vs. odd indices of frames or tilt angles. respectively, **e,** FSC analysis of map-model, where the model map is generated by the fitted model after being low-pass filtered to 8 Å. The resolutions are assessed based on the frequencies of the FSC curve falls at 0.5. Scale bars represent 10 nm in a, and 5 nm in b and c.

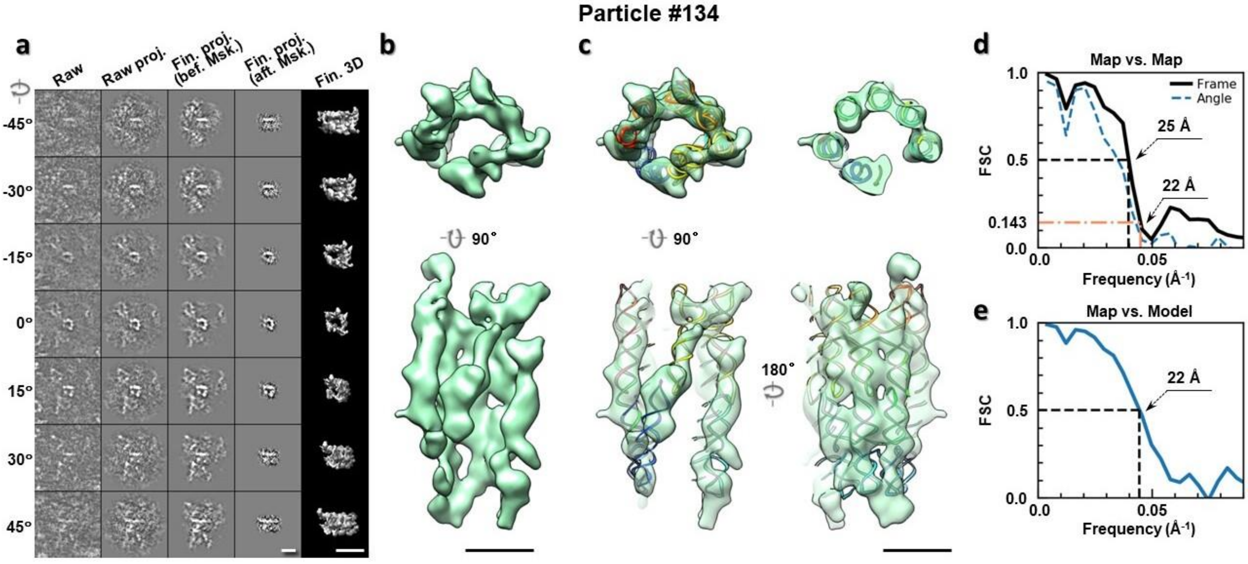

**Supplementary Fig. 139: IPET 3D reconstruction of an individual 6HBC RNA particle #134. a,** Seven representative tilt images of a single particle are shown in the first column from the left. The tilt images are aligned to a common center using IPET through iterative refinements. The projections of the raw, intermediate, and final 3D reconstructions at the corresponding angles are displayed in the next four columns. **b,** Two orthogonal views of the final 3D map; **c,** Overlap of the map with the fitted model; **d,** FSC analyses of the final map resolution using two methods, "map-map FSC", in which each map is reconstructed from one half of the images, based on the even vs. odd indices of frames or tilt angles. respectively, **e,** FSC analysis of map-model, where the model map is generated by the fitted model after being low-pass filtered to 8 Å. The resolutions are assessed based on the frequencies of the FSC curve falls at 0.5. Scale bars represent 10 nm in a, and 5 nm in b and c.

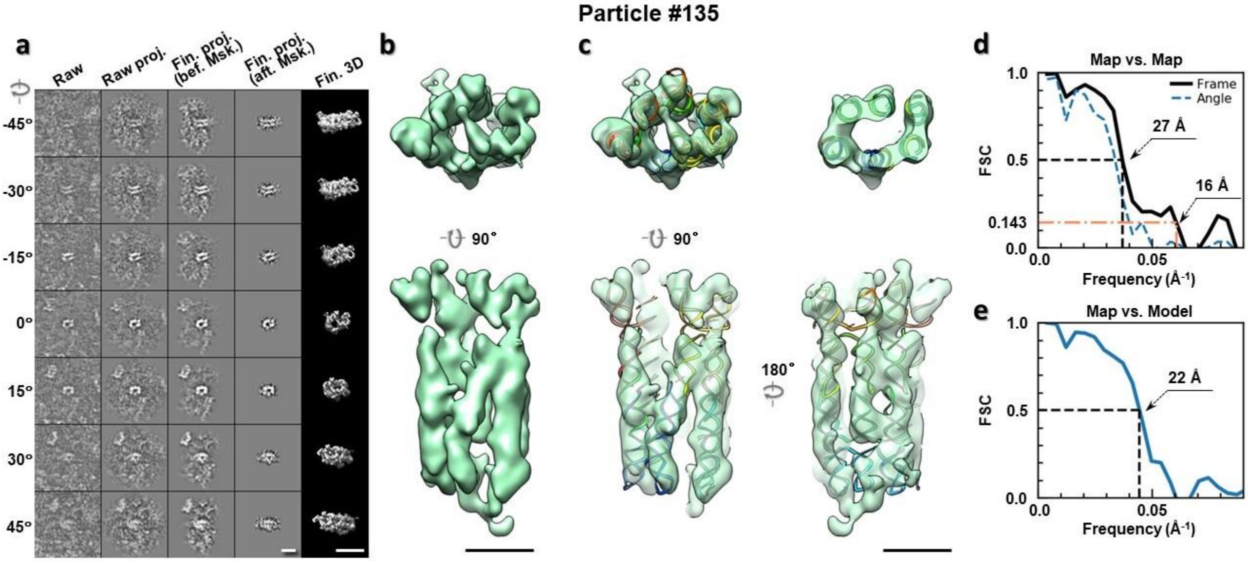

**Supplementary Fig. 140: IPET 3D reconstruction of an individual 6HBC RNA particle #135. a,** Seven representative tilt images of a single particle are shown in the first column from the left. The tilt images are aligned to a common center using IPET through iterative refinements. The projections of the raw, intermediate, and final 3D reconstructions at the corresponding angles are displayed in the next four

columns. **b,** Two orthogonal views of the final 3D map; **c,** Overlap of the map with the fitted model; **d,** FSC analyses of the final map resolution using two methods, "map-map FSC", in which each map is reconstructed from one half of the images, based on the even vs. odd indices of frames or tilt angles. respectively, **e,** FSC analysis of map-model, where the model map is generated by the fitted model after being low-pass filtered to 8 Å. The resolutions are assessed based on the frequencies of the FSC curve falls at 0.5. Scale bars represent 10 nm in a, and 5 nm in b and c.

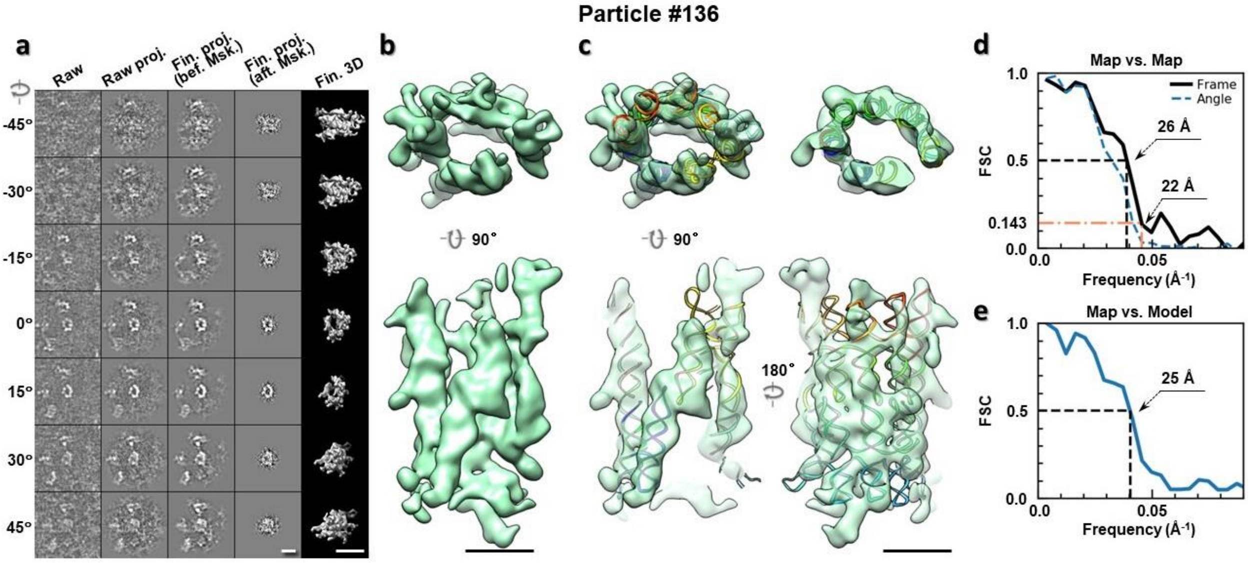

**Supplementary Fig. 141: IPET 3D reconstruction of an individual 6HBC RNA particle #136.** **a,** Seven representative tilt images of a single particle are shown in the first column from the left. The tilt images are aligned to a common center using IPET through iterative refinements. The projections of the raw, intermediate, and final 3D reconstructions at the corresponding angles are displayed in the next four columns. **b,** Two orthogonal views of the final 3D map; **c,** Overlap of the map with the fitted model; **d,** FSC analyses of the final map resolution using two methods, "map-map FSC", in which each map is reconstructed from one half of the images, based on the even vs. odd indices of frames or tilt angles. respectively, **e,** FSC analysis of map-model, where the model map is generated by the fitted model after being low-pass filtered to 8 Å. The resolutions are assessed based on the frequencies of the FSC curve falls at 0.5. Scale bars represent 10 nm in a, and 5 nm in b and c.

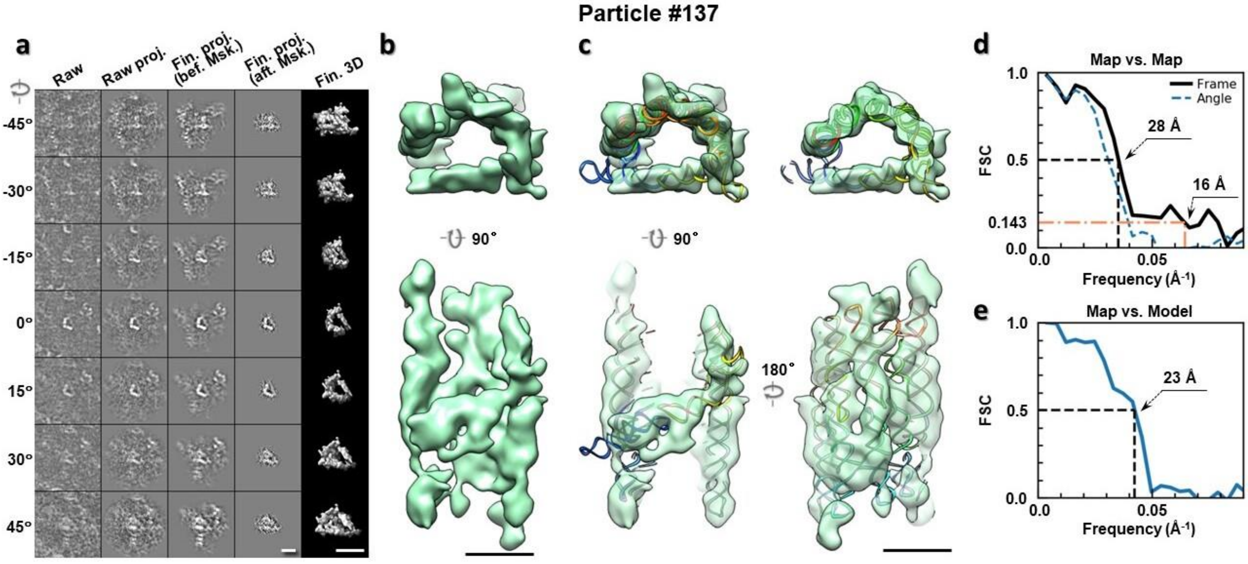

**Supplementary Fig. 142: IPET 3D reconstruction of an individual 6HBC RNA particle #137. a,** Seven representative tilt images of a single particle are shown in the first column from the left. The tilt images are aligned to a common center using IPET through iterative refinements. The projections of the raw, intermediate, and final 3D reconstructions at the corresponding angles are displayed in the next four columns. **b,** Two orthogonal views of the final 3D map; **c,** Overlap of the map with the fitted model; **d,** FSC analyses of the final map resolution using two methods, "map-map FSC", in which each map is reconstructed from one half of the images, based on the even vs. odd indices of frames or tilt angles. respectively, **e,** FSC analysis of map-model, where the model map is generated by the fitted model after being low-pass filtered to 8 Å. The resolutions are assessed based on the frequencies of the FSC curve falls at 0.5. Scale bars represent 10 nm in a, and 5 nm in b and c.

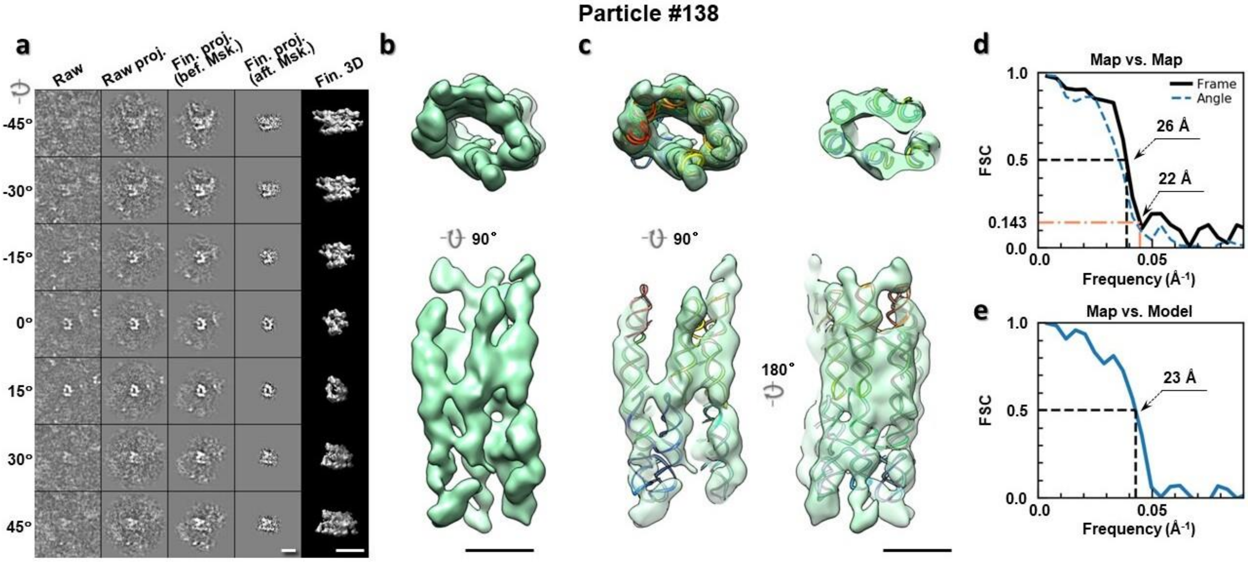

**Supplementary Fig. 143: IPET 3D reconstruction of an individual 6HBC RNA particle #138. a,** Seven representative tilt images of a single particle are shown in the first column from the left. The tilt images are aligned to a common center using IPET through iterative refinements. The projections of the raw, intermediate, and final 3D reconstructions at the corresponding angles are displayed in the next four

columns. **b,** Two orthogonal views of the final 3D map; **c,** Overlap of the map with the fitted model; **d,** FSC analyses of the final map resolution using two methods, "map-map FSC", in which each map is reconstructed from one half of the images, based on the even vs. odd indices of frames or tilt angles. respectively, **e,** FSC analysis of map-model, where the model map is generated by the fitted model after being low-pass filtered to 8 Å. The resolutions are assessed based on the frequencies of the FSC curve falls at 0.5. Scale bars represent 10 nm in a, and 5 nm in b and c.

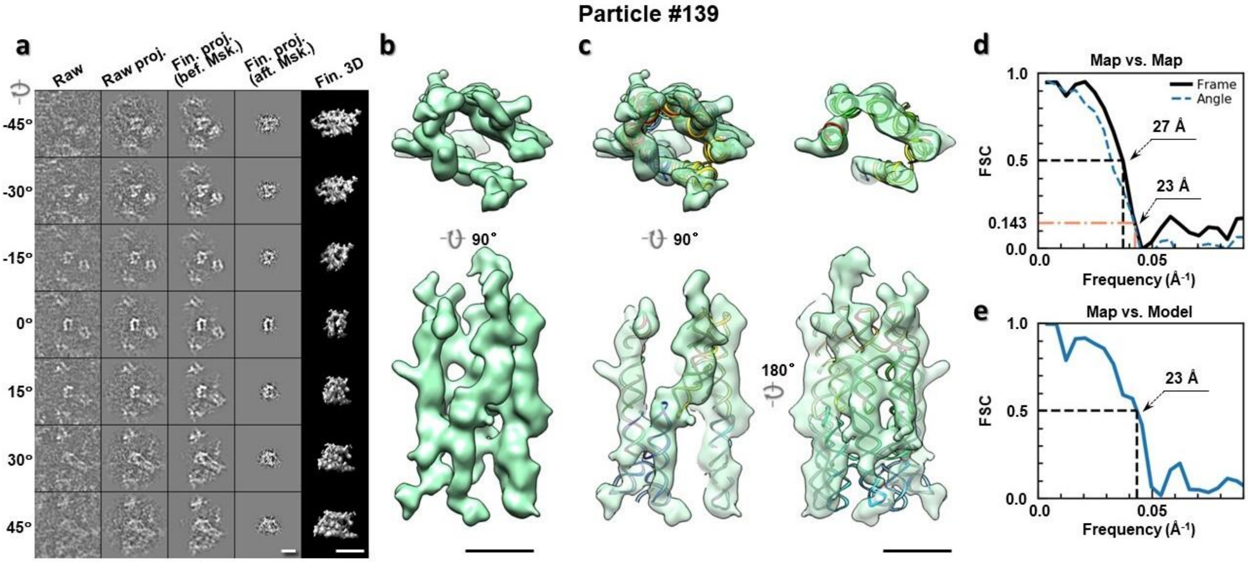

**Supplementary Fig. 144: IPET 3D reconstruction of an individual 6HBC RNA particle #139. a,** Seven representative tilt images of a single particle are shown in the first column from the left. The tilt images are aligned to a common center using IPET through iterative refinements. The projections of the raw, intermediate, and final 3D reconstructions at the corresponding angles are displayed in the next four columns. **b,** Two orthogonal views of the final 3D map; **c,** Overlap of the map with the fitted model; **d,** FSC analyses of the final map resolution using two methods, "map-map FSC", in which each map is reconstructed from one half of the images, based on the even vs. odd indices of frames or tilt angles. respectively, **e,** FSC analysis of map-model, where the model map is generated by the fitted model after being low-pass filtered to 8 Å. The resolutions are assessed based on the frequencies of the FSC curve falls at 0.5. Scale bars represent 10 nm in a, and 5 nm in b and c.

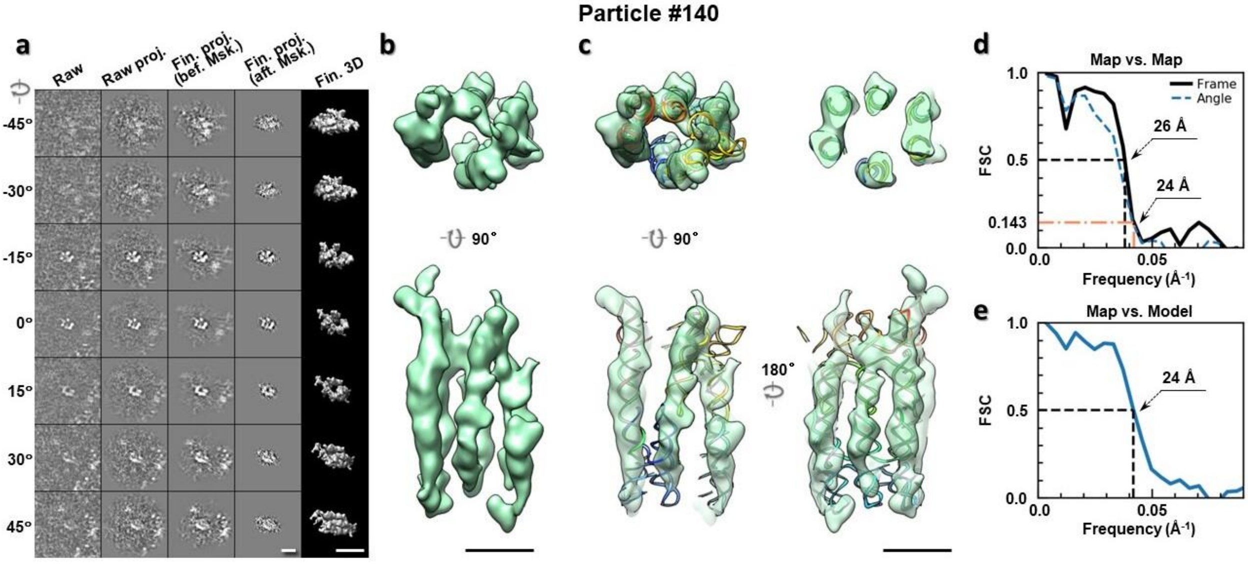

**Supplementary Fig. 145: IPET 3D reconstruction of an individual 6HBC RNA particle #140. a,** Seven representative tilt images of a single particle are shown in the first column from the left. The tilt images are aligned to a common center using IPET through iterative refinements. The projections of the raw, intermediate, and final 3D reconstructions at the corresponding angles are displayed in the next four columns. **b,** Two orthogonal views of the final 3D map; **c,** Overlap of the map with the fitted model; **d,** FSC analyses of the final map resolution using two methods, "map-map FSC", in which each map is reconstructed from one half of the images, based on the even vs. odd indices of frames or tilt angles. respectively, **e,** FSC analysis of map-model, where the model map is generated by the fitted model after being low-pass filtered to 8 Å. The resolutions are assessed based on the frequencies of the FSC curve falls at 0.5. Scale bars represent 10 nm in a, and 5 nm in b and c.

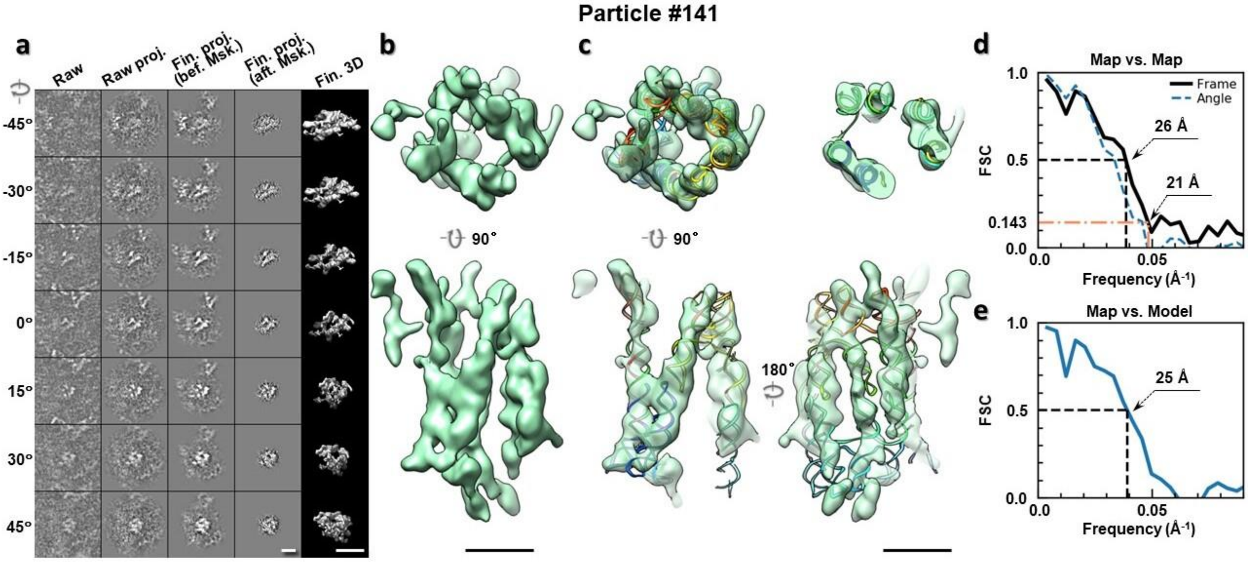

**Supplementary Fig. 146: IPET 3D reconstruction of an individual 6HBC RNA particle #141. a,** Seven representative tilt images of a single particle are shown in the first column from the left. The tilt images are aligned to a common center using IPET through iterative refinements. The projections of the raw, intermediate, and final 3D reconstructions at the corresponding angles are displayed in the next four

columns. **b,** Two orthogonal views of the final 3D map; **c,** Overlap of the map with the fitted model; **d,** FSC analyses of the final map resolution using two methods, "map-map FSC", in which each map is reconstructed from one half of the images, based on the even vs. odd indices of frames or tilt angles. respectively, **e,** FSC analysis of map-model, where the model map is generated by the fitted model after being low-pass filtered to 8 Å. The resolutions are assessed based on the frequencies of the FSC curve falls at 0.5. Scale bars represent 10 nm in a, and 5 nm in b and c.

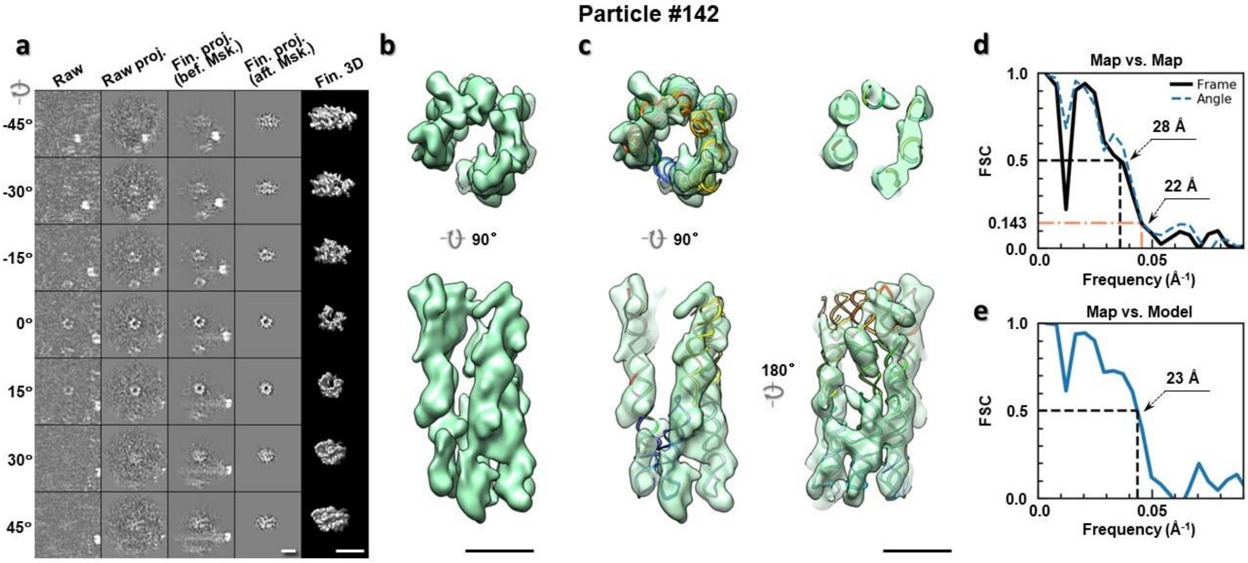

**Supplementary Fig. 147: IPET 3D reconstruction of an individual 6HBC RNA particle #142.** **a,** Seven representative tilt images of a single particle are shown in the first column from the left. The tilt images are aligned to a common center using IPET through iterative refinements. The projections of the raw, intermediate, and final 3D reconstructions at the corresponding angles are displayed in the next four columns. **b,** Two orthogonal views of the final 3D map; **c,** Overlap of the map with the fitted model; **d,** FSC analyses of the final map resolution using two methods, "map-map FSC", in which each map is reconstructed from one half of the images, based on the even vs. odd indices of frames or tilt angles. respectively, **e,** FSC analysis of map-model, where the model map is generated by the fitted model after being low-pass filtered to 8 Å. The resolutions are assessed based on the frequencies of the FSC curve falls at 0.5. Scale bars represent 10 nm in a, and 5 nm in b and c.

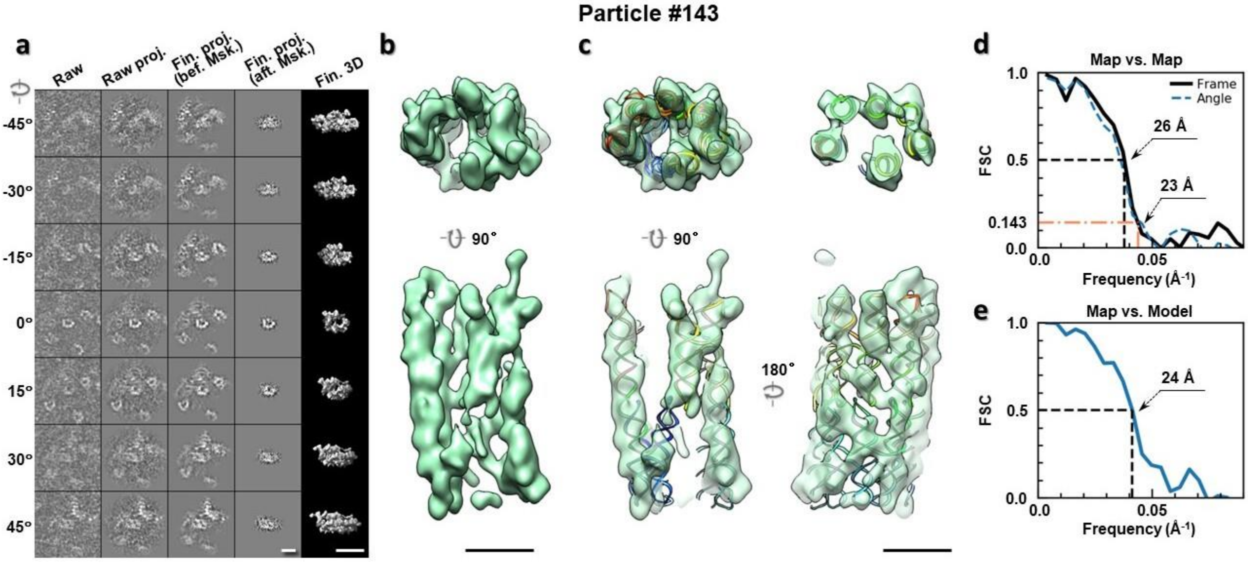

**Supplementary Fig. 148: IPET 3D reconstruction of an individual 6HBC RNA particle #143. a,** Seven representative tilt images of a single particle are shown in the first column from the left. The tilt images are aligned to a common center using IPET through iterative refinements. The projections of the raw, intermediate, and final 3D reconstructions at the corresponding angles are displayed in the next four columns. **b,** Two orthogonal views of the final 3D map; **c,** Overlap of the map with the fitted model; **d,** FSC analyses of the final map resolution using two methods, "map-map FSC", in which each map is reconstructed from one half of the images, based on the even vs. odd indices of frames or tilt angles. respectively, **e,** FSC analysis of map-model, where the model map is generated by the fitted model after being low-pass filtered to 8 Å. The resolutions are assessed based on the frequencies of the FSC curve falls at 0.5. Scale bars represent 10 nm in a, and 5 nm in b and c.

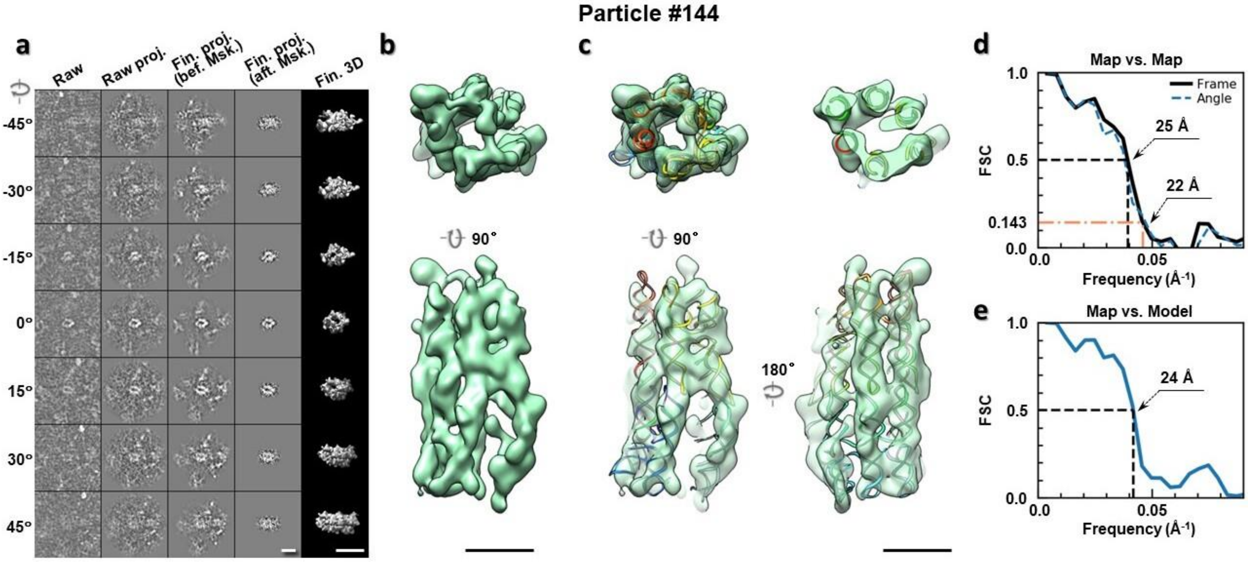

**Supplementary Fig. 149: IPET 3D reconstruction of an individual 6HBC RNA particle #144. a,** Seven representative tilt images of a single particle are shown in the first column from the left. The tilt images are aligned to a common center using IPET through iterative refinements. The projections of the raw, intermediate, and final 3D reconstructions at the corresponding angles are displayed in the next four

columns. **b,** Two orthogonal views of the final 3D map; **c,** Overlap of the map with the fitted model; **d,** FSC analyses of the final map resolution using two methods, "map-map FSC", in which each map is reconstructed from one half of the images, based on the even vs. odd indices of frames or tilt angles. respectively, **e,** FSC analysis of map-model, where the model map is generated by the fitted model after being low-pass filtered to 8 Å. The resolutions are assessed based on the frequencies of the FSC curve falls at 0.5. Scale bars represent 10 nm in a, and 5 nm in b and c.

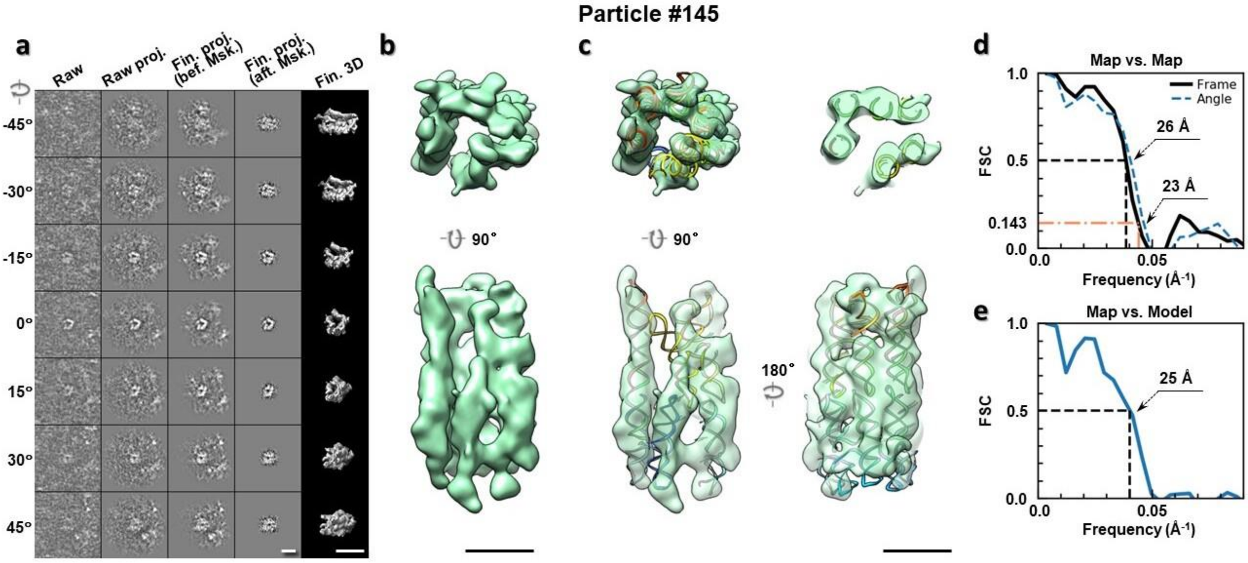

**Supplementary Fig. 150: IPET 3D reconstruction of an individual 6HBC RNA particle #145. a,** Seven representative tilt images of a single particle are shown in the first column from the left. The tilt images are aligned to a common center using IPET through iterative refinements. The projections of the raw, intermediate, and final 3D reconstructions at the corresponding angles are displayed in the next four columns. **b,** Two orthogonal views of the final 3D map; **c,** Overlap of the map with the fitted model; **d,** FSC analyses of the final map resolution using two methods, "map-map FSC", in which each map is reconstructed from one half of the images, based on the even vs. odd indices of frames or tilt angles. respectively, **e,** FSC analysis of map-model, where the model map is generated by the fitted model after being low-pass filtered to 8 Å. The resolutions are assessed based on the frequencies of the FSC curve falls at 0.5. Scale bars represent 10 nm in a, and 5 nm in b and c.

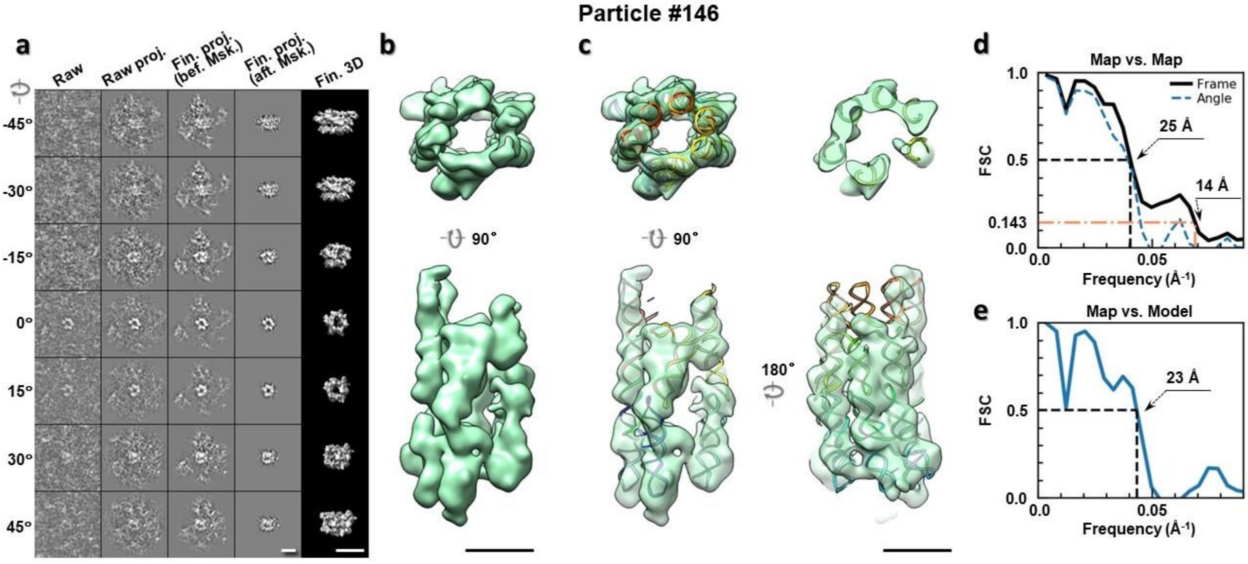

**Supplementary Fig. 151: IPET 3D reconstruction of an individual 6HBC RNA particle #146. a,** Seven representative tilt images of a single particle are shown in the first column from the left. The tilt images are aligned to a common center using IPET through iterative refinements. The projections of the raw, intermediate, and final 3D reconstructions at the corresponding angles are displayed in the next four columns. **b,** Two orthogonal views of the final 3D map; **c,** Overlap of the map with the fitted model; **d,** FSC analyses of the final map resolution using two methods, "map-map FSC", in which each map is reconstructed from one half of the images, based on the even vs. odd indices of frames or tilt angles. respectively, **e,** FSC analysis of map-model, where the model map is generated by the fitted model after being low-pass filtered to 8 Å. The resolutions are assessed based on the frequencies of the FSC curve falls at 0.5. Scale bars represent 10 nm in a, and 5 nm in b and c.

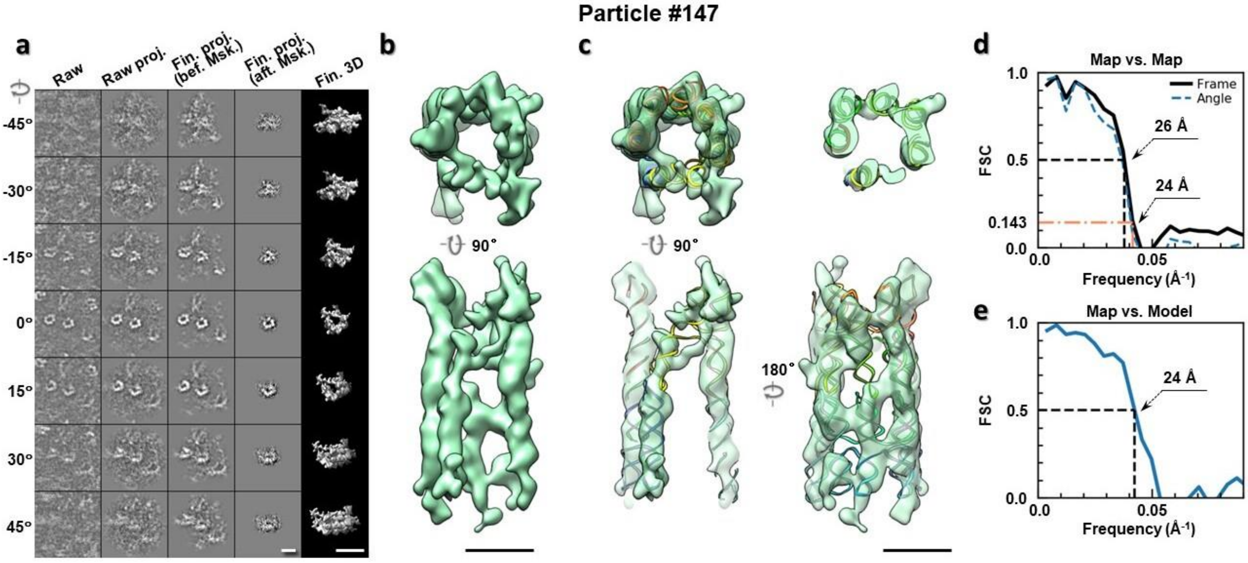

**Supplementary Fig. 152: IPET 3D reconstruction of an individual 6HBC RNA particle #147. a,** Seven representative tilt images of a single particle are shown in the first column from the left. The tilt images are aligned to a common center using IPET through iterative refinements. The projections of the raw, intermediate, and final 3D reconstructions at the corresponding angles are displayed in the next four

columns. **b,** Two orthogonal views of the final 3D map; **c,** Overlap of the map with the fitted model; **d,** FSC analyses of the final map resolution using two methods, "map-map FSC", in which each map is reconstructed from one half of the images, based on the even vs. odd indices of frames or tilt angles. respectively, **e,** FSC analysis of map-model, where the model map is generated by the fitted model after being low-pass filtered to 8 Å. The resolutions are assessed based on the frequencies of the FSC curve falls at 0.5. Scale bars represent 10 nm in a, and 5 nm in b and c.

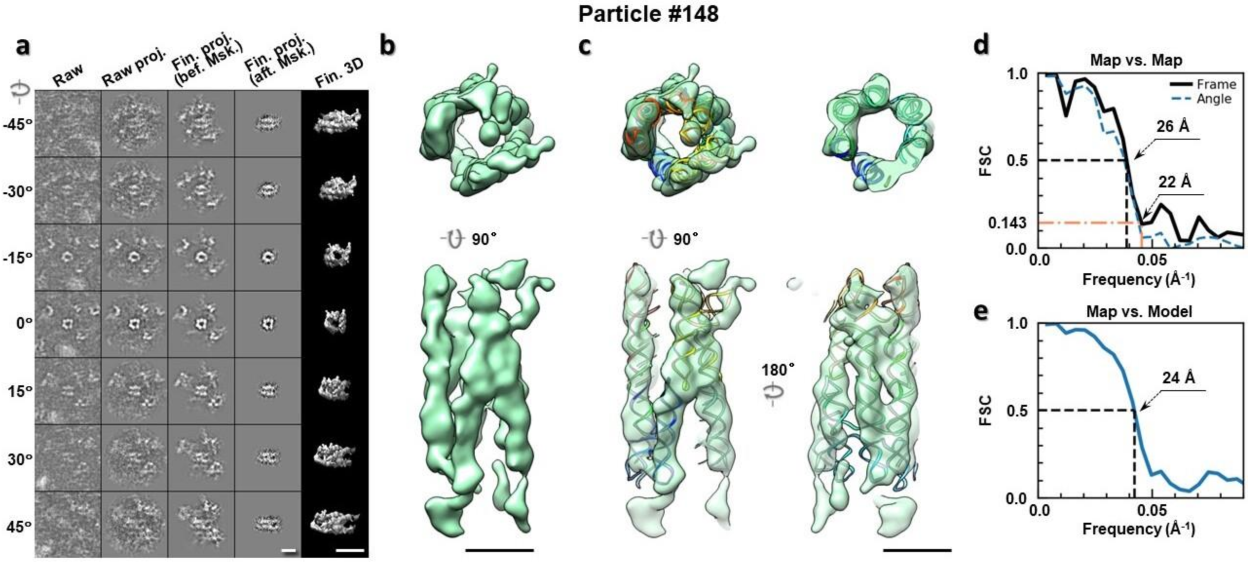

**Supplementary Fig. 153: IPET 3D reconstruction of an individual 6HBC RNA particle #148.** **a,** Seven representative tilt images of a single particle are shown in the first column from the left. The tilt images are aligned to a common center using IPET through iterative refinements. The projections of the raw, intermediate, and final 3D reconstructions at the corresponding angles are displayed in the next four columns. **b,** Two orthogonal views of the final 3D map; **c,** Overlap of the map with the fitted model; **d,** FSC analyses of the final map resolution using two methods, "map-map FSC", in which each map is reconstructed from one half of the images, based on the even vs. odd indices of frames or tilt angles. respectively, **e,** FSC analysis of map-model, where the model map is generated by the fitted model after being low-pass filtered to 8 Å. The resolutions are assessed based on the frequencies of the FSC curve falls at 0.5. Scale bars represent 10 nm in a, and 5 nm in b and c.

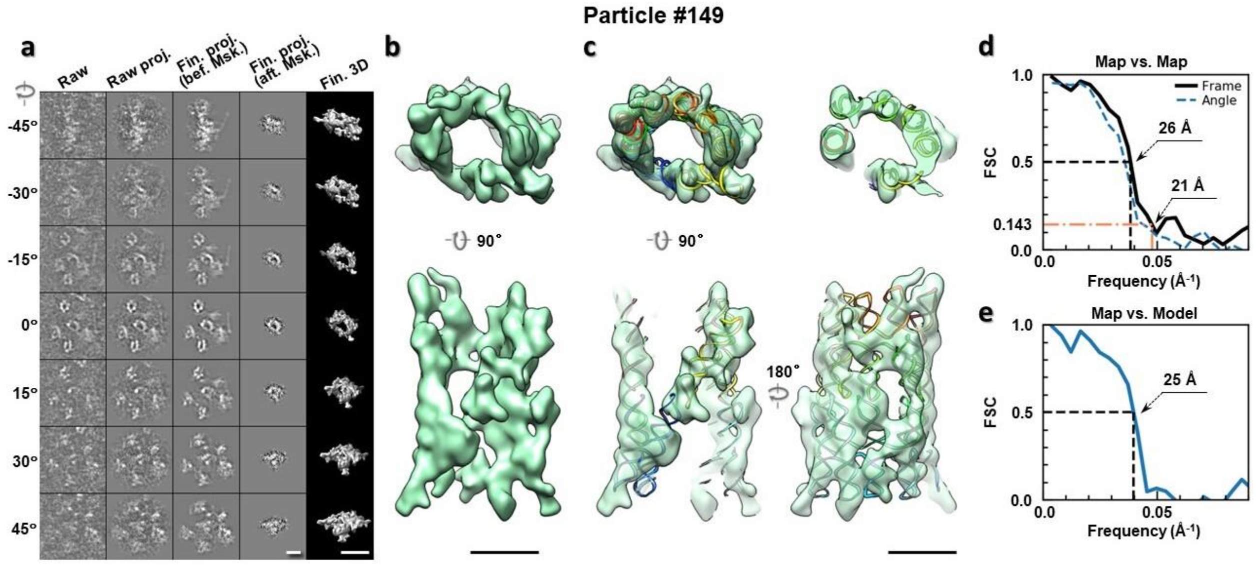

**Supplementary Fig. 154: IPET 3D reconstruction of an individual 6HBC RNA particle #149. a,** Seven representative tilt images of a single particle are shown in the first column from the left. The tilt images are aligned to a common center using IPET through iterative refinements. The projections of the raw, intermediate, and final 3D reconstructions at the corresponding angles are displayed in the next four columns. **b,** Two orthogonal views of the final 3D map; **c,** Overlap of the map with the fitted model; **d,** FSC analyses of the final map resolution using two methods, "map-map FSC", in which each map is reconstructed from one half of the images, based on the even vs. odd indices of frames or tilt angles. respectively, **e,** FSC analysis of map-model, where the model map is generated by the fitted model after being low-pass filtered to 8 Å. The resolutions are assessed based on the frequencies of the FSC curve falls at 0.5. Scale bars represent 10 nm in a, and 5 nm in b and c.

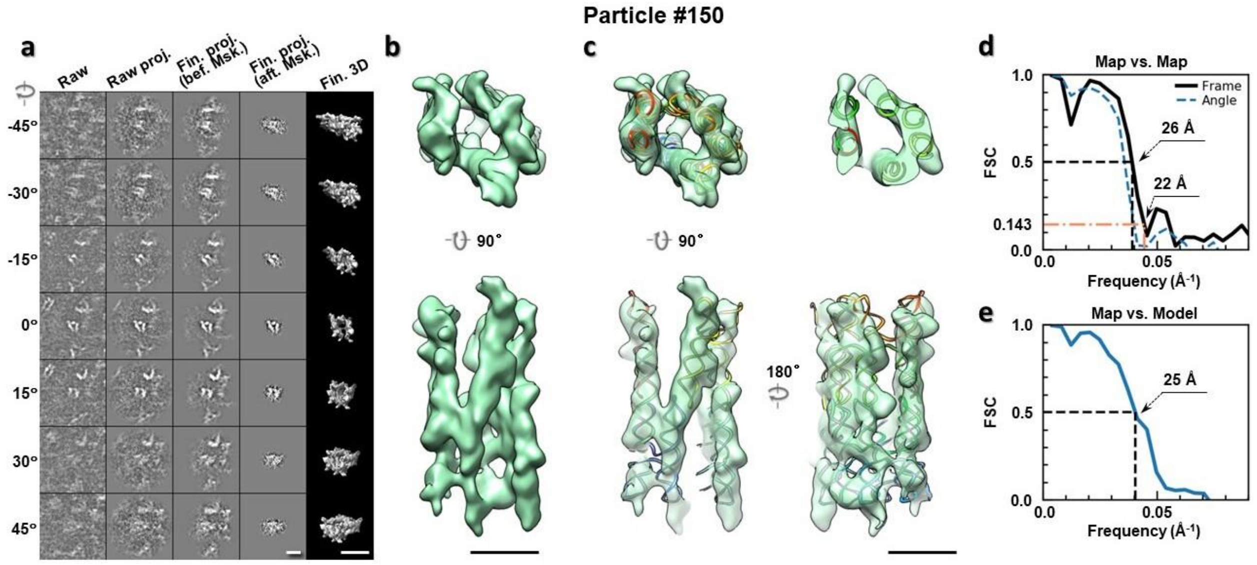

**Supplementary Fig. 155: IPET 3D reconstruction of an individual 6HBC RNA particle #150. a,** Seven representative tilt images of a single particle are shown in the first column from the left. The tilt images are aligned to a common center using IPET through iterative refinements. The projections of the raw, intermediate, and final 3D reconstructions at the corresponding angles are displayed in the next four

columns. **b,** Two orthogonal views of the final 3D map; **c,** Overlap of the map with the fitted model; **d,** FSC analyses of the final map resolution using two methods, "map-map FSC", in which each map is reconstructed from one half of the images, based on the even vs. odd indices of frames or tilt angles. respectively, **e,** FSC analysis of map-model, where the model map is generated by the fitted model after being low-pass filtered to 8 Å. The resolutions are assessed based on the frequencies of the FSC curve falls at 0.5. Scale bars represent 10 nm in a, and 5 nm in b and c.

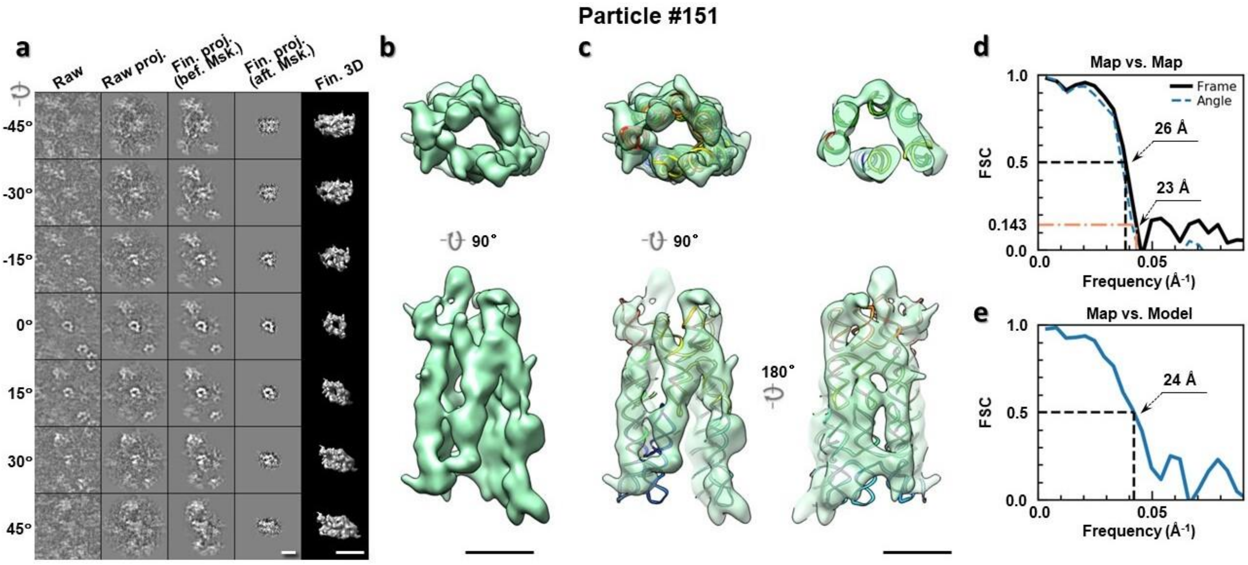

**Supplementary Fig. 156: IPET 3D reconstruction of an individual 6HBC RNA particle #151. a,** Seven representative tilt images of a single particle are shown in the first column from the left. The tilt images are aligned to a common center using IPET through iterative refinements. The projections of the raw, intermediate, and final 3D reconstructions at the corresponding angles are displayed in the next four columns. **b,** Two orthogonal views of the final 3D map; **c,** Overlap of the map with the fitted model; **d,** FSC analyses of the final map resolution using two methods, "map-map FSC", in which each map is reconstructed from one half of the images, based on the even vs. odd indices of frames or tilt angles. respectively, **e,** FSC analysis of map-model, where the model map is generated by the fitted model after being low-pass filtered to 8 Å. The resolutions are assessed based on the frequencies of the FSC curve falls at 0.5. Scale bars represent 10 nm in a, and 5 nm in b and c.

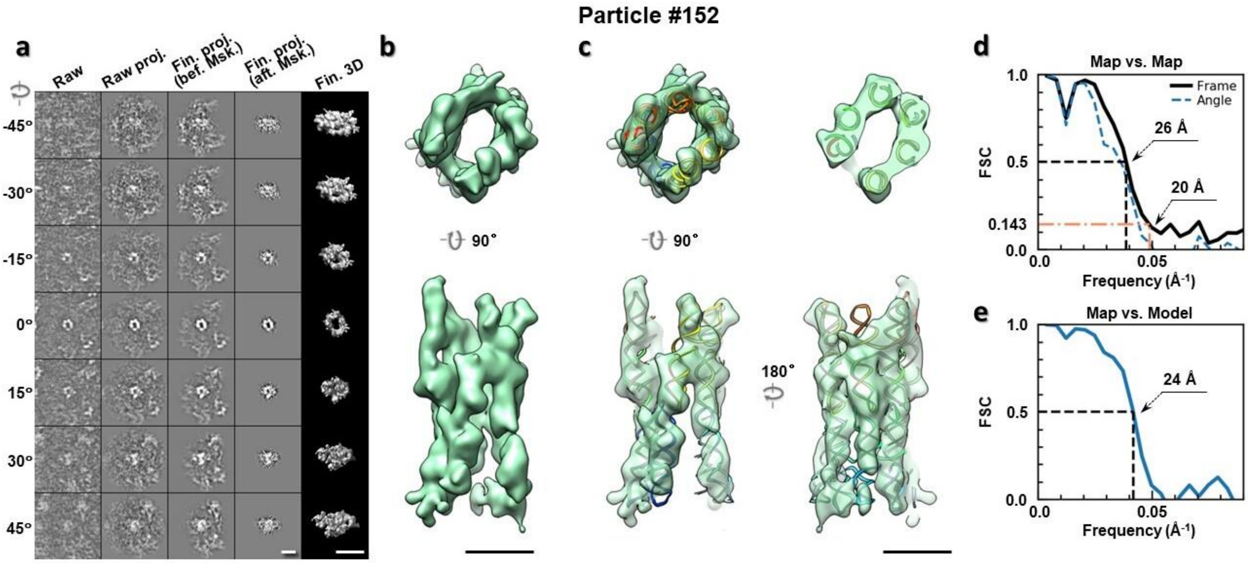

**Supplementary Fig. 157: IPET 3D reconstruction of an individual 6HBC RNA particle #152. a,** Seven representative tilt images of a single particle are shown in the first column from the left. The tilt images are aligned to a common center using IPET through iterative refinements. The projections of the raw, intermediate, and final 3D reconstructions at the corresponding angles are displayed in the next four columns. **b,** Two orthogonal views of the final 3D map; **c,** Overlap of the map with the fitted model; **d,** FSC analyses of the final map resolution using two methods, "map-map FSC", in which each map is reconstructed from one half of the images, based on the even vs. odd indices of frames or tilt angles. respectively, **e,** FSC analysis of map-model, where the model map is generated by the fitted model after being low-pass filtered to 8 Å. The resolutions are assessed based on the frequencies of the FSC curve falls at 0.5. Scale bars represent 10 nm in a, and 5 nm in b and c.

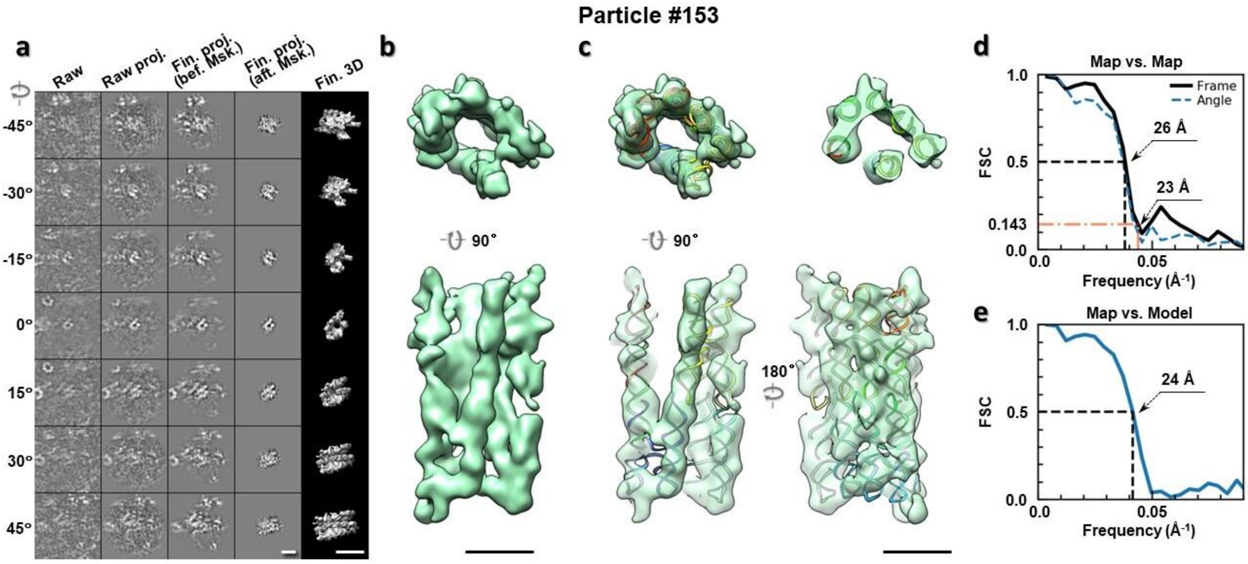

**Supplementary Fig. 158: IPET 3D reconstruction of an individual 6HBC RNA particle #153. a,** Seven representative tilt images of a single particle are shown in the first column from the left. The tilt images are aligned to a common center using IPET through iterative refinements. The projections of the raw, intermediate, and final 3D reconstructions at the corresponding angles are displayed in the next four

columns. **b,** Two orthogonal views of the final 3D map; **c,** Overlap of the map with the fitted model; **d,** FSC analyses of the final map resolution using two methods, "map-map FSC", in which each map is reconstructed from one half of the images, based on the even vs. odd indices of frames or tilt angles. respectively, **e,** FSC analysis of map-model, where the model map is generated by the fitted model after being low-pass filtered to 8 Å. The resolutions are assessed based on the frequencies of the FSC curve falls at 0.5. Scale bars represent 10 nm in a, and 5 nm in b and c.

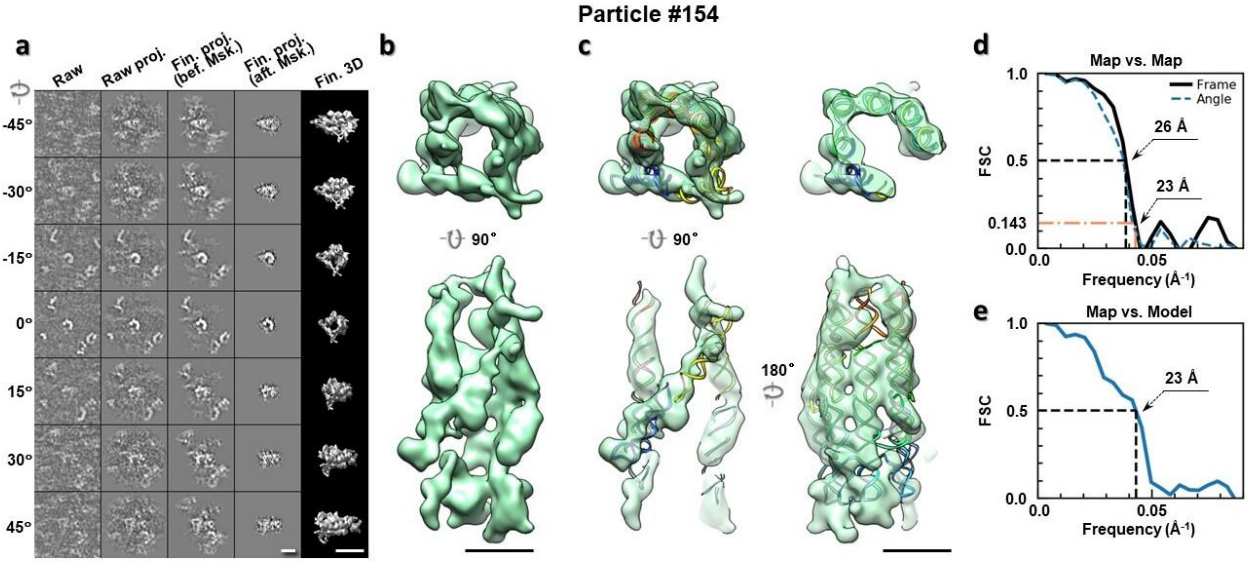

**Supplementary Fig. 159: IPET 3D reconstruction of an individual 6HBC RNA particle #154. a,** Seven representative tilt images of a single particle are shown in the first column from the left. The tilt images are aligned to a common center using IPET through iterative refinements. The projections of the raw, intermediate, and final 3D reconstructions at the corresponding angles are displayed in the next four columns. **b,** Two orthogonal views of the final 3D map; **c,** Overlap of the map with the fitted model; **d,** FSC analyses of the final map resolution using two methods, "map-map FSC", in which each map is reconstructed from one half of the images, based on the even vs. odd indices of frames or tilt angles. respectively, **e,** FSC analysis of map-model, where the model map is generated by the fitted model after being low-pass filtered to 8 Å. The resolutions are assessed based on the frequencies of the FSC curve falls at 0.5. Scale bars represent 10 nm in a, and 5 nm in b and c.

### Particle #155

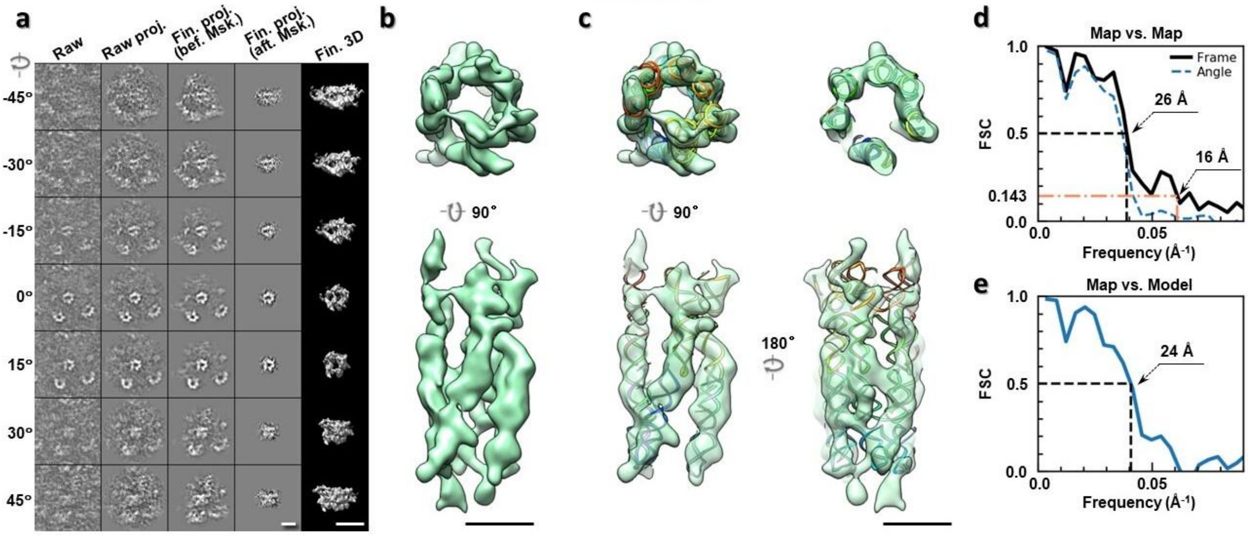

**Supplementary Fig. 160: IPET 3D reconstruction of an individual 6HBC RNA particle #155. a,** Seven representative tilt images of a single particle are shown in the first column from the left. The tilt images are aligned to a common center using IPET through iterative refinements. The projections of the raw, intermediate, and final 3D reconstructions at the corresponding angles are displayed in the next four columns. **b,** Two orthogonal views of the final 3D map; **c,** Overlap of the map with the fitted model; **d,** FSC analyses of the final map resolution using two methods, "map-map FSC", in which each map is reconstructed from one half of the images, based on the even vs. odd indices of frames or tilt angles. respectively, **e,** FSC analysis of map-model, where the model map is generated by the fitted model after being low-pass filtered to 8 Å. The resolutions are assessed based on the frequencies of the FSC curve falls at 0.5. Scale bars represent 10 nm in a, and 5 nm in b and c.

### Particle #156

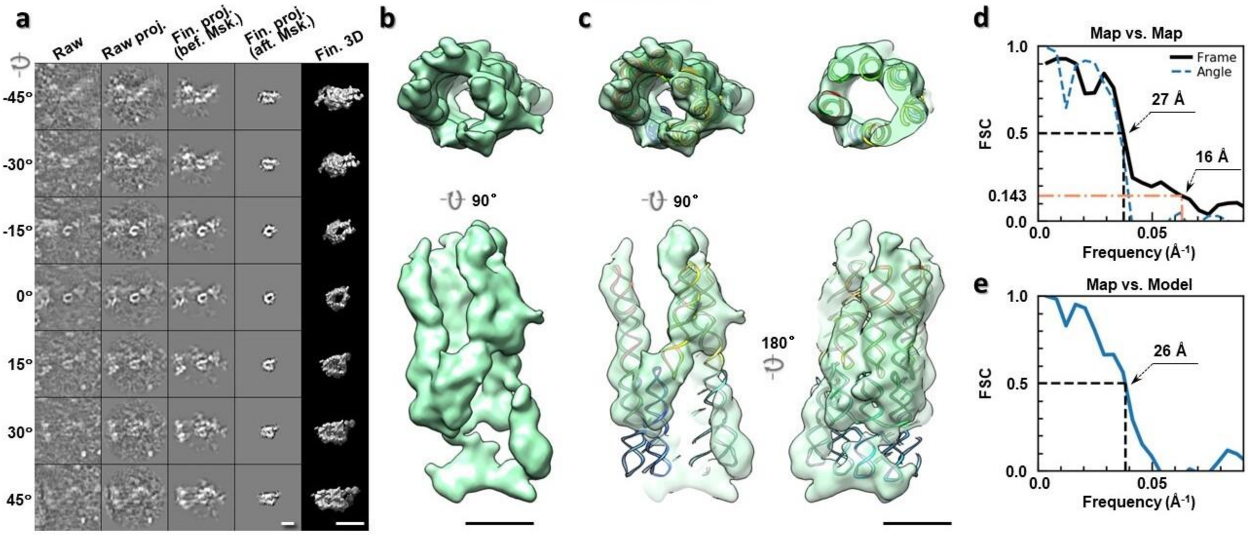

**Supplementary Fig. 161: IPET 3D reconstruction of an individual 6HBC RNA particle #156. a,** Seven representative tilt images of a single particle are shown in the first column from the left. The tilt images are aligned to a common center using IPET through iterative refinements. The projections of the raw, intermediate, and final 3D reconstructions at the corresponding angles are displayed in the next four

columns. **b,** Two orthogonal views of the final 3D map; **c,** Overlap of the map with the fitted model; **d,** FSC analyses of the final map resolution using two methods, "map-map FSC", in which each map is reconstructed from one half of the images, based on the even vs. odd indices of frames or tilt angles. respectively, **e,** FSC analysis of map-model, where the model map is generated by the fitted model after being low-pass filtered to 8 Å. The resolutions are assessed based on the frequencies of the FSC curve falls at 0.5. Scale bars represent 10 nm in a, and 5 nm in b and c.

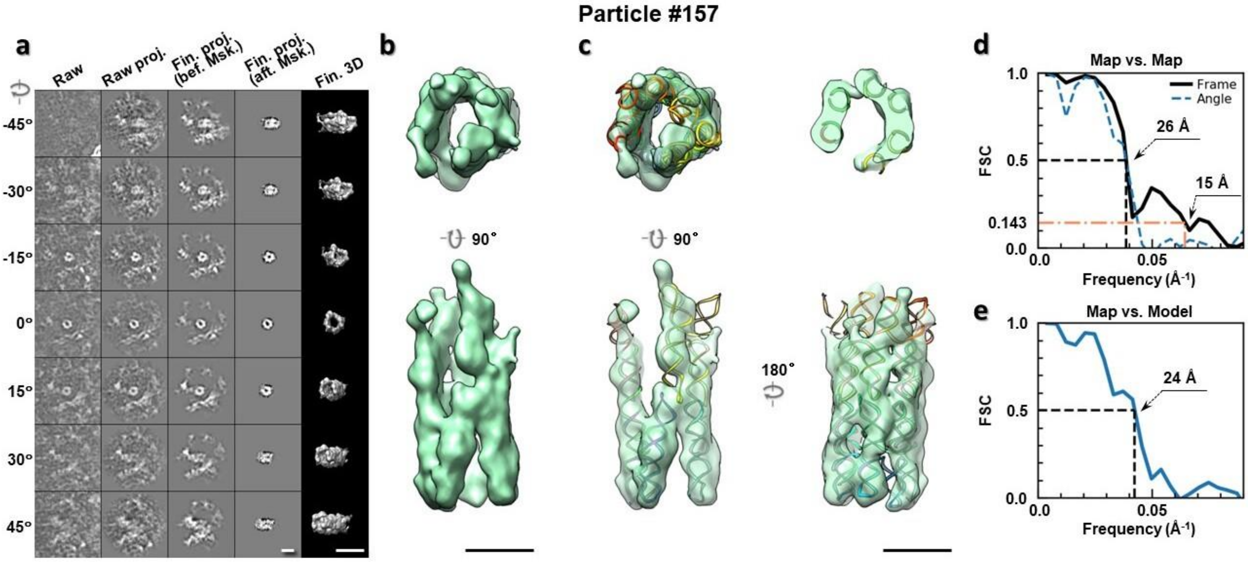

**Supplementary Fig. 162: IPET 3D reconstruction of an individual 6HBC RNA particle #157. a,** Seven representative tilt images of a single particle are shown in the first column from the left. The tilt images are aligned to a common center using IPET through iterative refinements. The projections of the raw, intermediate, and final 3D reconstructions at the corresponding angles are displayed in the next four columns. **b,** Two orthogonal views of the final 3D map; **c,** Overlap of the map with the fitted model; **d,** FSC analyses of the final map resolution using two methods, "map-map FSC", in which each map is reconstructed from one half of the images, based on the even vs. odd indices of frames or tilt angles. respectively, **e,** FSC analysis of map-model, where the model map is generated by the fitted model after being low-pass filtered to 8 Å. The resolutions are assessed based on the frequencies of the FSC curve falls at 0.5. Scale bars represent 10 nm in a, and 5 nm in b and c.

Particle #158

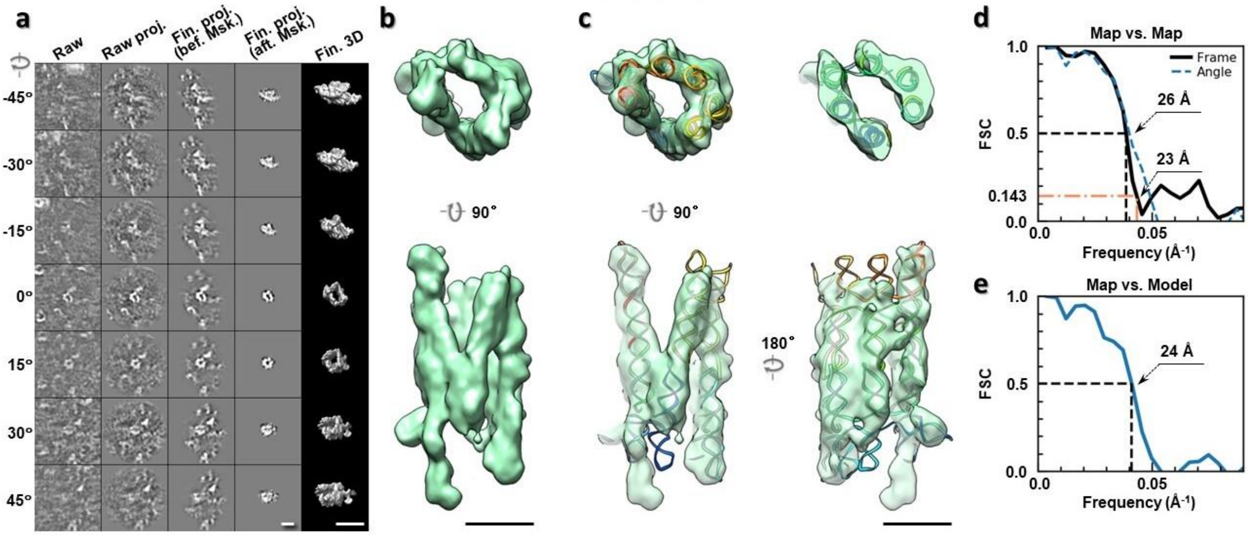

**Supplementary Fig. 163: IPET 3D reconstruction of an individual 6HBC RNA particle #158. a,** Seven representative tilt images of a single particle are shown in the first column from the left. The tilt images are aligned to a common center using IPET through iterative refinements. The projections of the raw, intermediate, and final 3D reconstructions at the corresponding angles are displayed in the next four columns. **b,** Two orthogonal views of the final 3D map; **c,** Overlap of the map with the fitted model; **d,** FSC analyses of the final map resolution using two methods, "map-map FSC", in which each map is reconstructed from one half of the images, based on the even vs. odd indices of frames or tilt angles. respectively, **e,** FSC analysis of map-model, where the model map is generated by the fitted model after being low-pass filtered to 8 Å. The resolutions are assessed based on the frequencies of the FSC curve falls at 0.5. Scale bars represent 10 nm in a, and 5 nm in b and c.

Particle #159

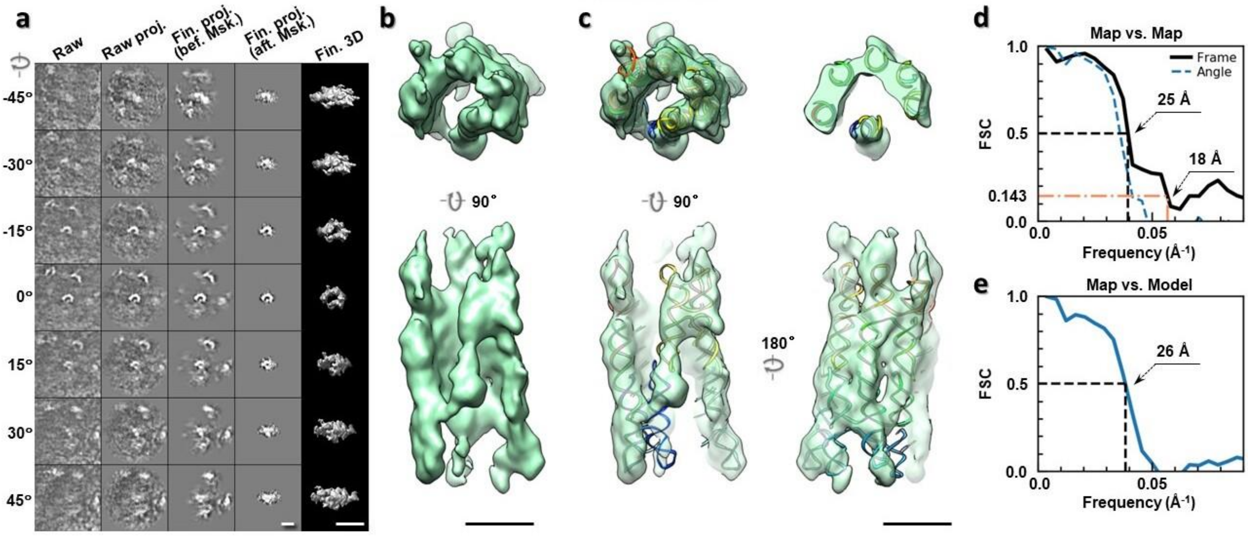

**Supplementary Fig. 164: IPET 3D reconstruction of an individual 6HBC RNA particle #159. a,** Seven representative tilt images of a single particle are shown in the first column from the left. The tilt images are aligned to a common center using IPET through iterative refinements. The projections of the raw, intermediate, and final 3D reconstructions at the corresponding angles are displayed in the next four

columns. **b,** Two orthogonal views of the final 3D map; **c,** Overlap of the map with the fitted model; **d,** FSC analyses of the final map resolution using two methods, "map-map FSC", in which each map is reconstructed from one half of the images, based on the even vs. odd indices of frames or tilt angles. respectively, **e,** FSC analysis of map-model, where the model map is generated by the fitted model after being low-pass filtered to 8 Å. The resolutions are assessed based on the frequencies of the FSC curve falls at 0.5. Scale bars represent 10 nm in a, and 5 nm in b and c.

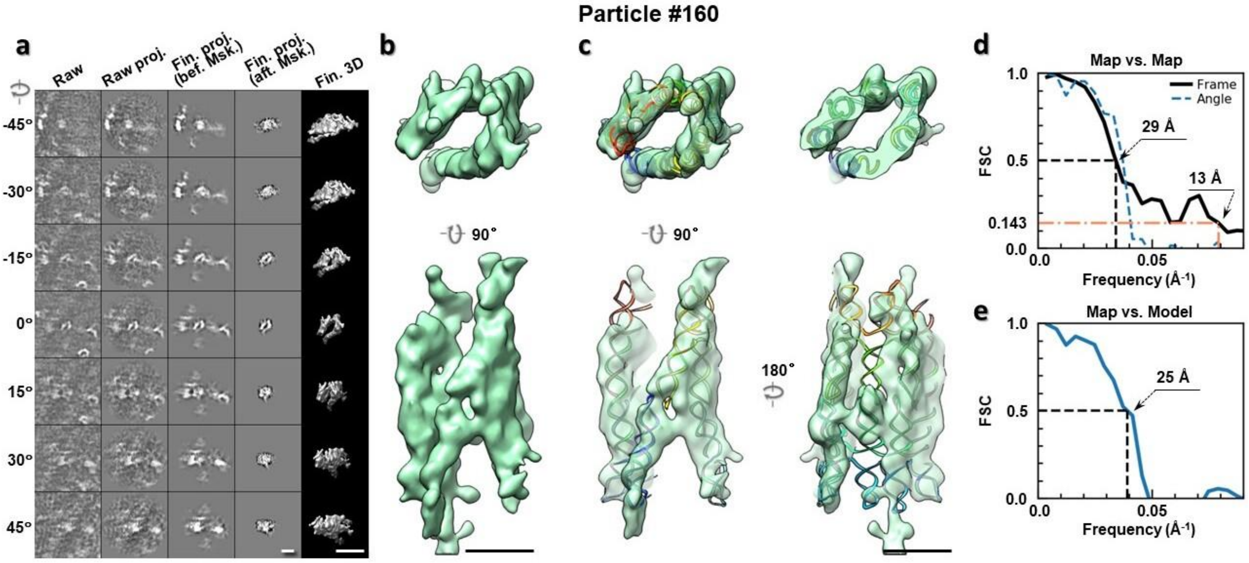

**Supplementary Fig. 165: IPET 3D reconstruction of an individual 6HBC RNA particle #160. a,** Seven representative tilt images of a single particle are shown in the first column from the left. The tilt images are aligned to a common center using IPET through iterative refinements. The projections of the raw, intermediate, and final 3D reconstructions at the corresponding angles are displayed in the next four columns. **b,** Two orthogonal views of the final 3D map; **c,** Overlap of the map with the fitted model; **d,** FSC analyses of the final map resolution using two methods, "map-map FSC", in which each map is reconstructed from one half of the images, based on the even vs. odd indices of frames or tilt angles. respectively, **e,** FSC analysis of map-model, where the model map is generated by the fitted model after being low-pass filtered to 8 Å. The resolutions are assessed based on the frequencies of the FSC curve falls at 0.5. Scale bars represent 10 nm in a, and 5 nm in b and c.

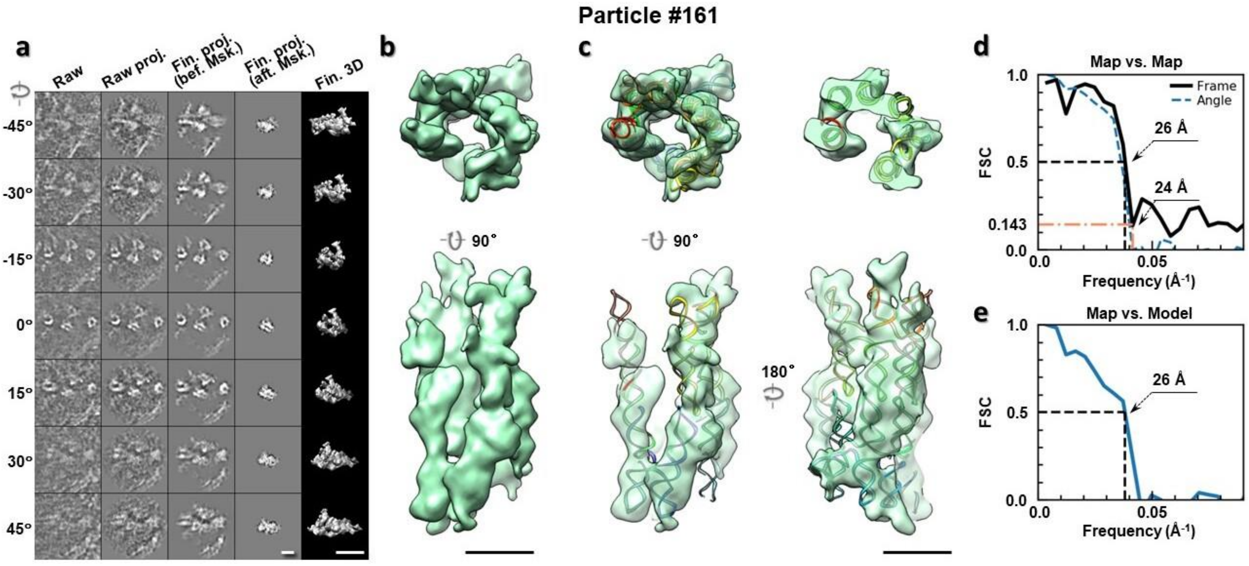

**Supplementary Fig. 166: IPET 3D reconstruction of an individual 6HBC RNA particle #161. a,** Seven representative tilt images of a single particle are shown in the first column from the left. The tilt images are aligned to a common center using IPET through iterative refinements. The projections of the raw, intermediate, and final 3D reconstructions at the corresponding angles are displayed in the next four columns. **b,** Two orthogonal views of the final 3D map; **c,** Overlap of the map with the fitted model; **d,** FSC analyses of the final map resolution using two methods, "map-map FSC", in which each map is reconstructed from one half of the images, based on the even vs. odd indices of frames or tilt angles. respectively, **e,** FSC analysis of map-model, where the model map is generated by the fitted model after being low-pass filtered to 8 Å. The resolutions are assessed based on the frequencies of the FSC curve falls at 0.5. Scale bars represent 10 nm in a, and 5 nm in b and c.

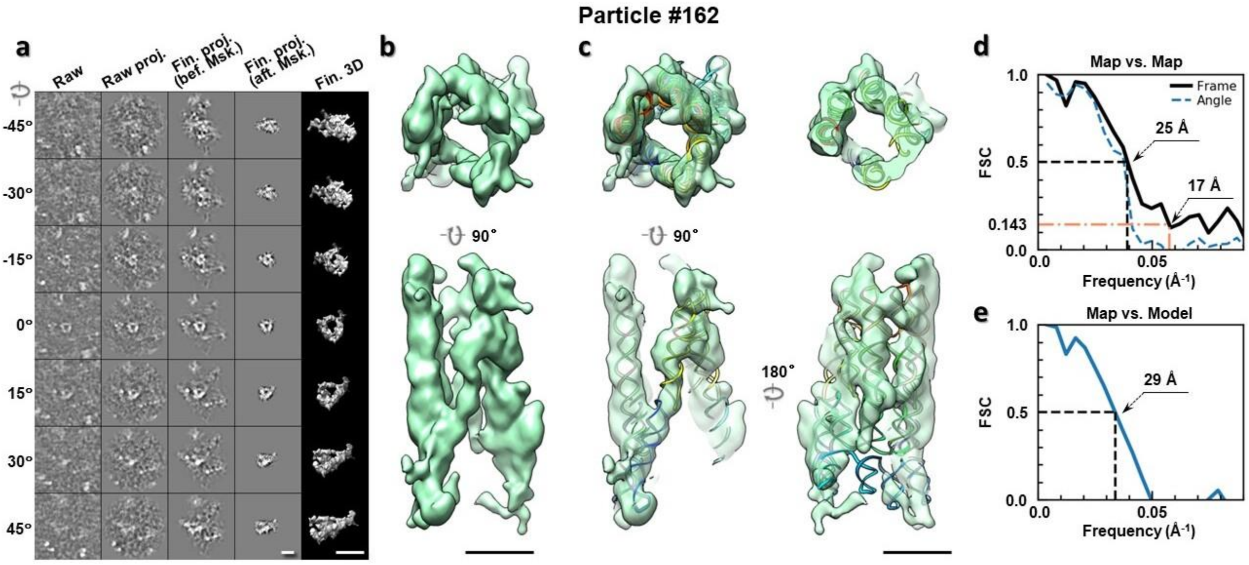

**Supplementary Fig. 167: IPET 3D reconstruction of an individual 6HBC RNA particle #162. a,** Seven representative tilt images of a single particle are shown in the first column from the left. The tilt images are aligned to a common center using IPET through iterative refinements. The projections of the raw, intermediate, and final 3D reconstructions at the corresponding angles are displayed in the next four

columns. **b,** Two orthogonal views of the final 3D map; **c,** Overlap of the map with the fitted model; **d,** FSC analyses of the final map resolution using two methods, "map-map FSC", in which each map is reconstructed from one half of the images, based on the even vs. odd indices of frames or tilt angles. respectively, **e,** FSC analysis of map-model, where the model map is generated by the fitted model after being low-pass filtered to 8 Å. The resolutions are assessed based on the frequencies of the FSC curve falls at 0.5. Scale bars represent 10 nm in a, and 5 nm in b and c.

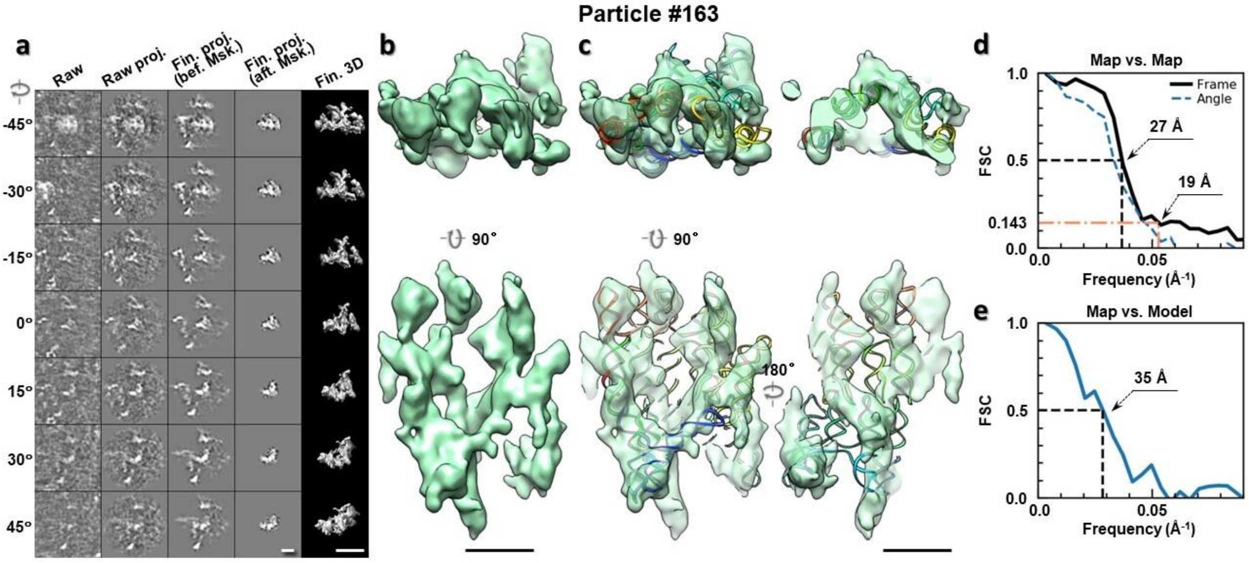

**Supplementary Fig. 168: IPET 3D reconstruction of an individual 6HBC RNA particle #163. a,** Seven representative tilt images of a single particle are shown in the first column from the left. The tilt images are aligned to a common center using IPET through iterative refinements. The projections of the raw, intermediate, and final 3D reconstructions at the corresponding angles are displayed in the next four columns. **b,** Two orthogonal views of the final 3D map; **c,** Overlap of the map with the fitted model; **d,** FSC analyses of the final map resolution using two methods, "map-map FSC", in which each map is reconstructed from one half of the images, based on the even vs. odd indices of frames or tilt angles. respectively, **e,** FSC analysis of map-model, where the model map is generated by the fitted model after being low-pass filtered to 8 Å. The resolutions are assessed based on the frequencies of the FSC curve falls at 0.5. Scale bars represent 10 nm in a, and 5 nm in b and c.

Particle #164

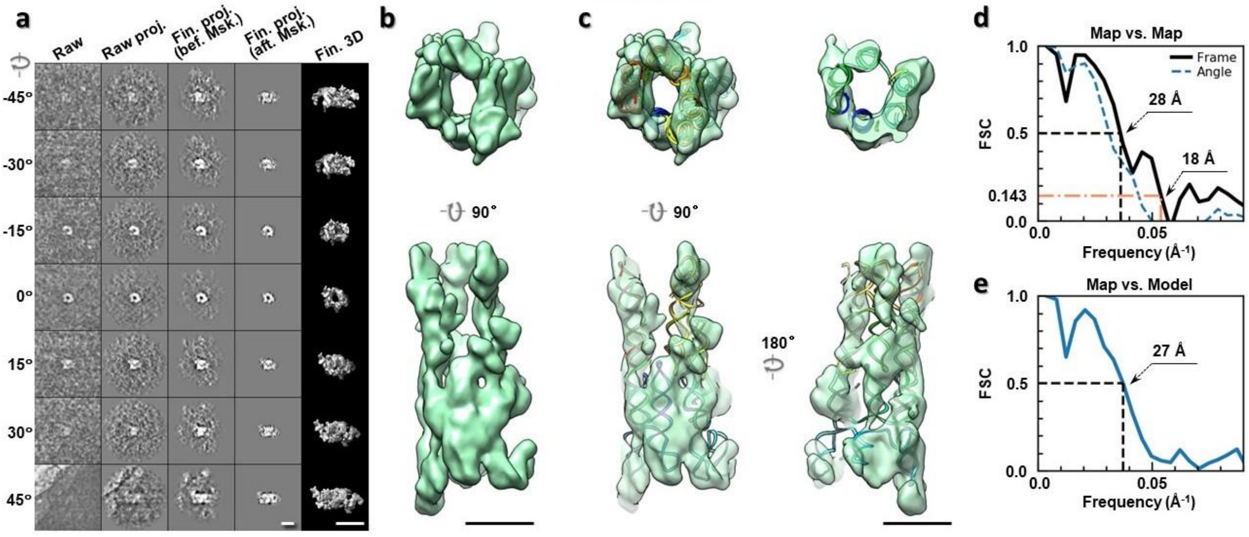

**Supplementary Fig. 169: IPET 3D reconstruction of an individual 6HBC RNA particle #164. a,** Seven representative tilt images of a single particle are shown in the first column from the left. The tilt images are aligned to a common center using IPET through iterative refinements. The projections of the raw, intermediate, and final 3D reconstructions at the corresponding angles are displayed in the next four columns. **b,** Two orthogonal views of the final 3D map; **c,** Overlap of the map with the fitted model; **d,** FSC analyses of the final map resolution using two methods, "map-map FSC", in which each map is reconstructed from one half of the images, based on the even vs. odd indices of frames or tilt angles. respectively, **e,** FSC analysis of map-model, where the model map is generated by the fitted model after being low-pass filtered to 8 Å. The resolutions are assessed based on the frequencies of the FSC curve falls at 0.5. Scale bars represent 10 nm in a, and 5 nm in b and c.

Particle #165

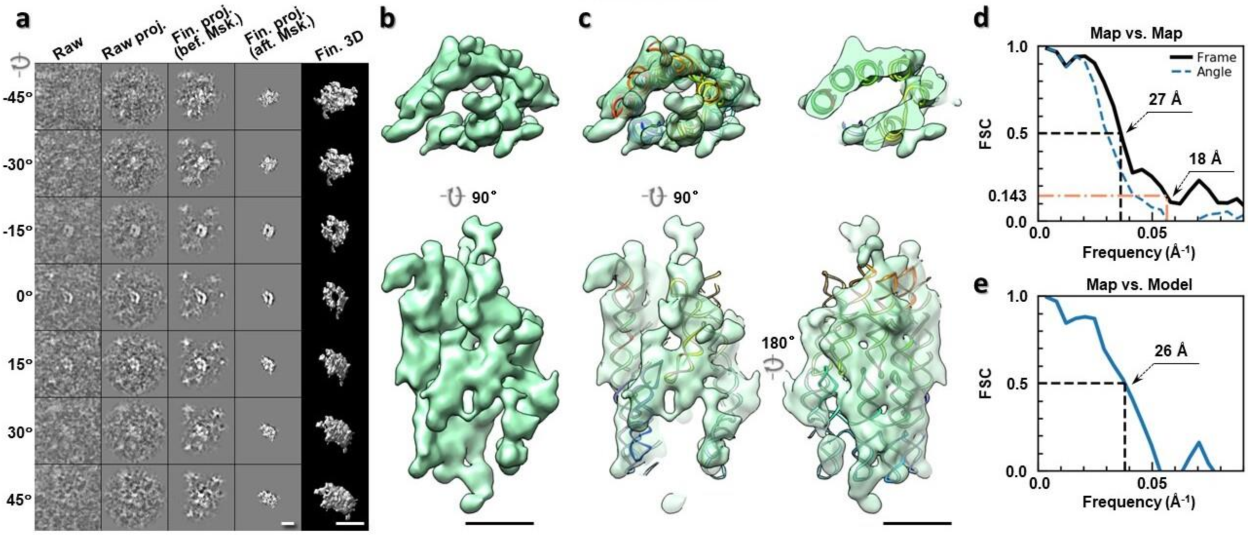

**Supplementary Fig. 170: IPET 3D reconstruction of an individual 6HBC RNA particle #165. a,** Seven representative tilt images of a single particle are shown in the first column from the left. The tilt images are aligned to a common center using IPET through iterative refinements. The projections of the raw, intermediate, and final 3D reconstructions at the corresponding angles are displayed in the next four

columns. **b,** Two orthogonal views of the final 3D map; **c,** Overlap of the map with the fitted model; **d,** FSC analyses of the final map resolution using two methods, "map-map FSC", in which each map is reconstructed from one half of the images, based on the even vs. odd indices of frames or tilt angles. respectively, **e,** FSC analysis of map-model, where the model map is generated by the fitted model after being low-pass filtered to 8 Å. The resolutions are assessed based on the frequencies of the FSC curve falls at 0.5. Scale bars represent 10 nm in a, and 5 nm in b and c.

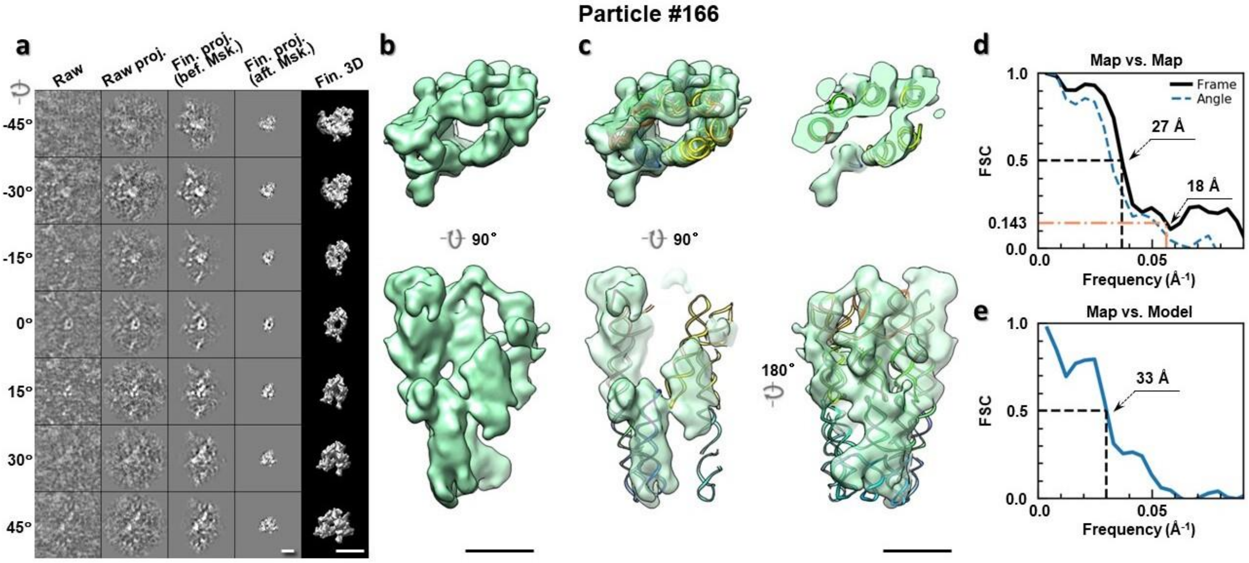

**Supplementary Fig. 171: IPET 3D reconstruction of an individual 6HBC RNA particle #166. a,** Seven representative tilt images of a single particle are shown in the first column from the left. The tilt images are aligned to a common center using IPET through iterative refinements. The projections of the raw, intermediate, and final 3D reconstructions at the corresponding angles are displayed in the next four columns. **b,** Two orthogonal views of the final 3D map; **c,** Overlap of the map with the fitted model; **d,** FSC analyses of the final map resolution using two methods, "map-map FSC", in which each map is reconstructed from one half of the images, based on the even vs. odd indices of frames or tilt angles. respectively, **e,** FSC analysis of map-model, where the model map is generated by the fitted model after being low-pass filtered to 8 Å. The resolutions are assessed based on the frequencies of the FSC curve falls at 0.5. Scale bars represent 10 nm in a, and 5 nm in b and c.

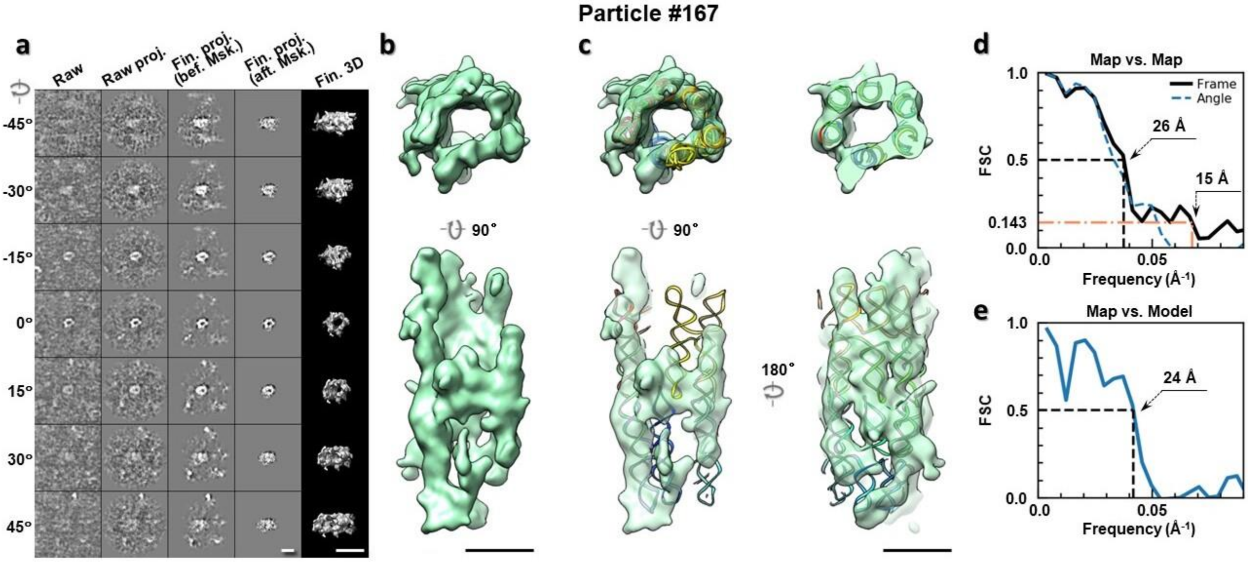

**Supplementary Fig. 172: IPET 3D reconstruction of an individual 6HBC RNA particle #167. a,** Seven representative tilt images of a single particle are shown in the first column from the left. The tilt images are aligned to a common center using IPET through iterative refinements. The projections of the raw, intermediate, and final 3D reconstructions at the corresponding angles are displayed in the next four columns. **b,** Two orthogonal views of the final 3D map; **c,** Overlap of the map with the fitted model; **d,** FSC analyses of the final map resolution using two methods, "map-map FSC", in which each map is reconstructed from one half of the images, based on the even vs. odd indices of frames or tilt angles. respectively, **e,** FSC analysis of map-model, where the model map is generated by the fitted model after being low-pass filtered to 8 Å. The resolutions are assessed based on the frequencies of the FSC curve falls at 0.5. Scale bars represent 10 nm in a, and 5 nm in b and c.

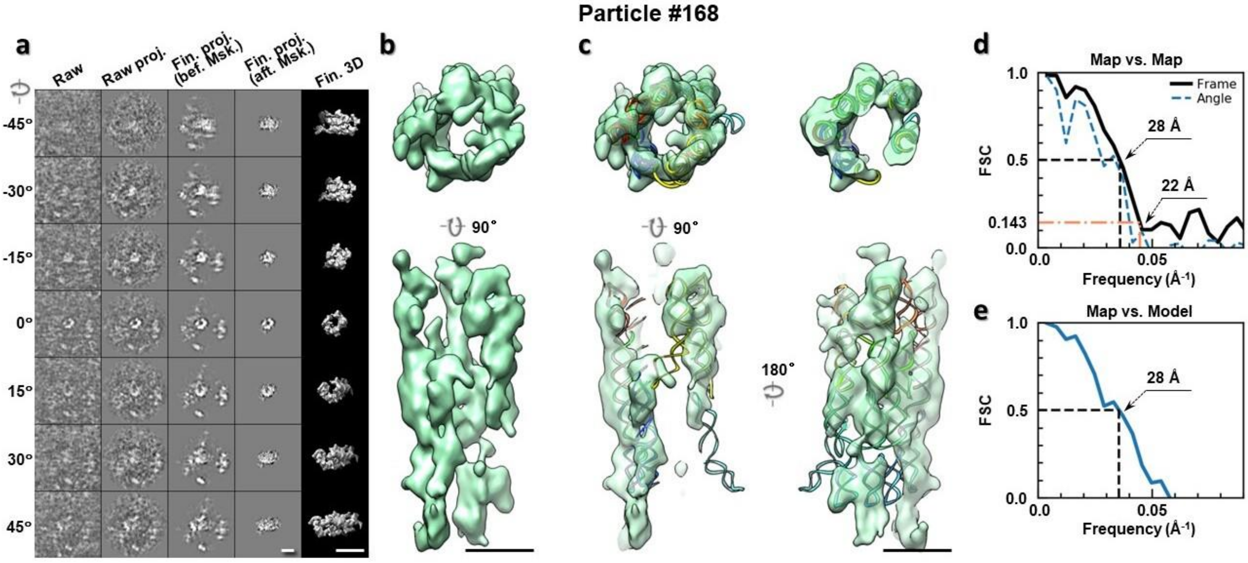

**Supplementary Fig. 173: IPET 3D reconstruction of an individual 6HBC RNA particle #168. a,** Seven representative tilt images of a single particle are shown in the first column from the left. The tilt images are aligned to a common center using IPET through iterative refinements. The projections of the raw, intermediate, and final 3D reconstructions at the corresponding angles are displayed in the next four

columns. **b,** Two orthogonal views of the final 3D map; **c,** Overlap of the map with the fitted model; **d,** FSC analyses of the final map resolution using two methods, "map-map FSC", in which each map is reconstructed from one half of the images, based on the even vs. odd indices of frames or tilt angles. respectively, **e,** FSC analysis of map-model, where the model map is generated by the fitted model after being low-pass filtered to 8 Å. The resolutions are assessed based on the frequencies of the FSC curve falls at 0.5. Scale bars represent 10 nm in a, and 5 nm in b and c.

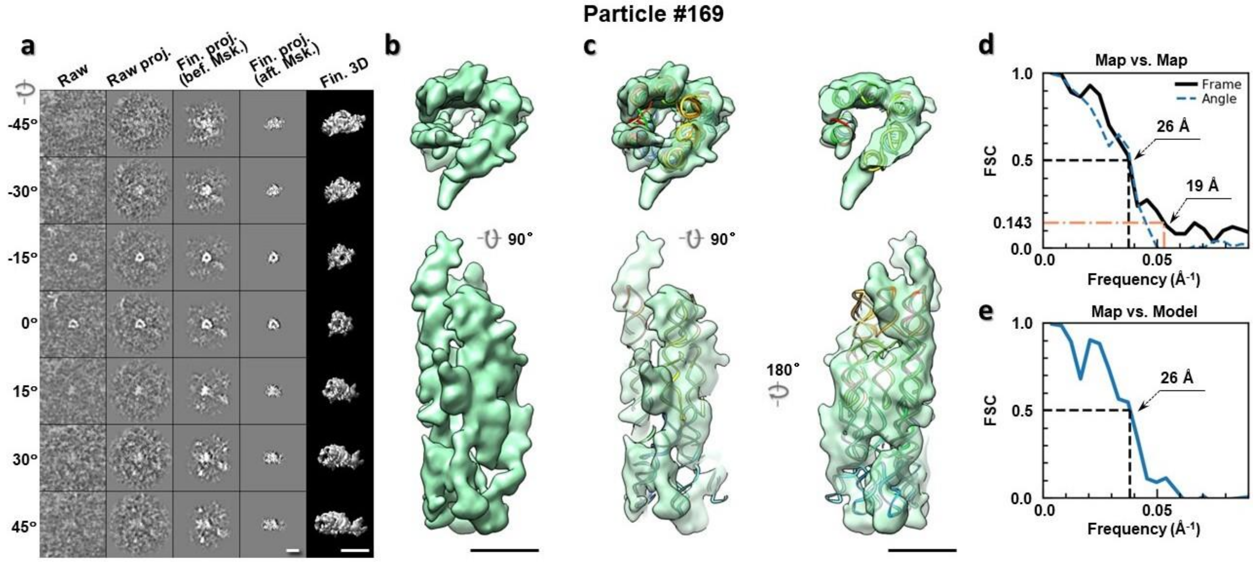

**Supplementary Fig. 174: IPET 3D reconstruction of an individual 6HBC RNA particle #169.** **a,** Seven representative tilt images of a single particle are shown in the first column from the left. The tilt images are aligned to a common center using IPET through iterative refinements. The projections of the raw, intermediate, and final 3D reconstructions at the corresponding angles are displayed in the next four columns. **b,** Two orthogonal views of the final 3D map; **c,** Overlap of the map with the fitted model; **d,** FSC analyses of the final map resolution using two methods, "map-map FSC", in which each map is reconstructed from one half of the images, based on the even vs. odd indices of frames or tilt angles. respectively, **e,** FSC analysis of map-model, where the model map is generated by the fitted model after being low-pass filtered to 8 Å. The resolutions are assessed based on the frequencies of the FSC curve falls at 0.5. Scale bars represent 10 nm in a, and 5 nm in b and c.

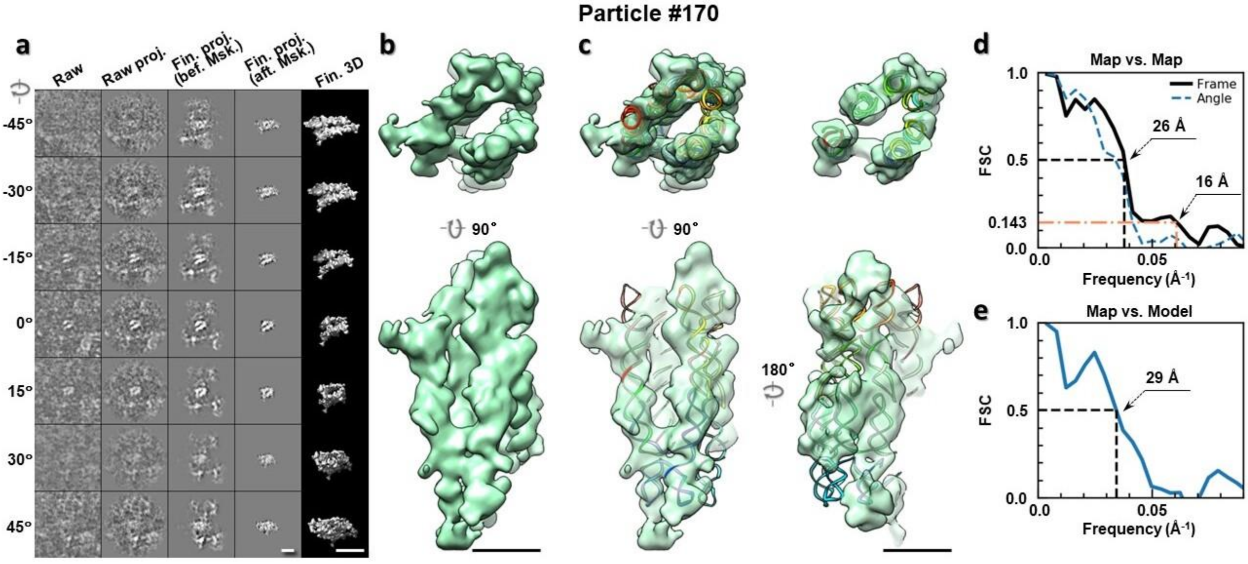

**Supplementary Fig. 175: IPET 3D reconstruction of an individual 6HBC RNA particle #170. a,** Seven representative tilt images of a single particle are shown in the first column from the left. The tilt images are aligned to a common center using IPET through iterative refinements. The projections of the raw, intermediate, and final 3D reconstructions at the corresponding angles are displayed in the next four columns. **b,** Two orthogonal views of the final 3D map; **c,** Overlap of the map with the fitted model; **d,** FSC analyses of the final map resolution using two methods, "map-map FSC", in which each map is reconstructed from one half of the images, based on the even vs. odd indices of frames or tilt angles. respectively, **e,** FSC analysis of map-model, where the model map is generated by the fitted model after being low-pass filtered to 8 Å. The resolutions are assessed based on the frequencies of the FSC curve falls at 0.5. Scale bars represent 10 nm in a, and 5 nm in b and c.
